# Algal kainoid synthases exhibit substrate-dependent hydroxylation and cyclization activities

**DOI:** 10.1101/2023.07.10.548461

**Authors:** Austin R. Hopiavuori, Shaun M. K. McKinnie

## Abstract

Fe^II^/α-ketoglutarate-dependent dioxygenases (Fe/αKG) are a large enzyme family that functionalize C-H bonds on diverse organic substrates. Although Fe/αKG homologs catalyze an array of chemically useful reactions, hydroxylation typically predominates. Microalgal DabC uniquely forms a novel C-C bond to construct the bioactive pyrrolidine ring in domoic acid biosynthesis. However, this kainoid synthase exclusively performs a stereospecific hydroxylation reaction on its *cis* substrate regioisomer. Mechanistic and kinetic analyses with native and alternative substrates identified a 20-fold rate increase in DabC radical cyclization over β–hydroxylation, with no observable 1,5-hydrogen atom transfer. Moreover, this dual activity was conserved among macroalgal RadC1 and KabC homologs and provided insight into substrate recognition and reactivity trends. Investigation of this substrate-dependent chemistry improves our understanding of Fe/αKG enzymes and their biocatalytic application.

---

The non-heme iron/α-ketoglutarate-dependent dioxygenases (Fe/αKGs) are a large and diverse enzyme family widely distributed throughout Nature in both primary and secondary metabolism.^[1,2]^ Fe/αKGs catalyze a variety of valuable transformations at unactivated C-H bonds including halogenations, desaturations, epoxidations, ring formations, expansions and contractions, however hydroxylation reactions are the most prevalent.^[1,2]^ Structurally, Fe/αKGs have a conserved double-stranded β-helix (DSBH) fold with a Fe(II)-binding facial triad. In the presence of atmospheric oxygen and αKG co-substrate, these enzymes generate a high-energy Fe(IV)-oxo species capable of radically abstracting a hydrogen from the targeted carbon center of the substrate.^[1,2]^ Outside their important regulatory roles on macromolecular substrates and in primary metabolism,^[1]^ Fe/αKG hydroxylases are important for the biosynthesis of natural products in diverse biological species. A number of dioxygenases act on amino acid substrates, which can be further functionalized and/or incorporated into complex natural product syntheses (Figures 1A, S1).^[3]^ Notable examples include bacterial VioC and TauD, and fungal IboH which stereospecifically hydroxylate L-arginine, taurine, and L-glutamic acid respectively (Figure 1A).^[4–6]^ One rare reaction that this family catalyzes is C-C bond formation; of the hundreds of characterized Fe/αKG enzymes, only a few have been confirmed to function as C-C bond forming cyclases (Figures 1A, S1).^[7]^ Cyclization reactions are often critical in natural product synthesis in that they restrict compounds into entropically favorable binding orientations that improves target specificity and potency. Recent examples of Fe/αKGs from diverse biological organisms that catalyze C-C bond cyclization include: BelL and HrmJ from belactosin A and hormaomycin biosyntheses in Streptomyces bacteria that catalyze the *trans*-nitrocyclopropanation of 6-nitro-L-norleucine with opposite diastereomeric outcomes;^[8]^ and 2-ODD from the plant podophyllotoxin biosynthetic pathway that assembles the core tetracyclic scaffold from (-)-yatein en route to the valuable chemotherapeutic etoposide (Figures 1A, S1).^[9,10]^ Notably, algal kainoid synthase DabC from the diatom *Pseudo-nitzschia multiseries* and homologs RadC1 and KabC from red macroalgae *Chondria armata* and *Digenea simplex* cyclize *N*-prenylated L-glutamic acids to construct the pyrrolidine moieties of domoic acid (DA) and kainic acid (KA) (Figure 1B).^[11–13]^

**Figure 1.**
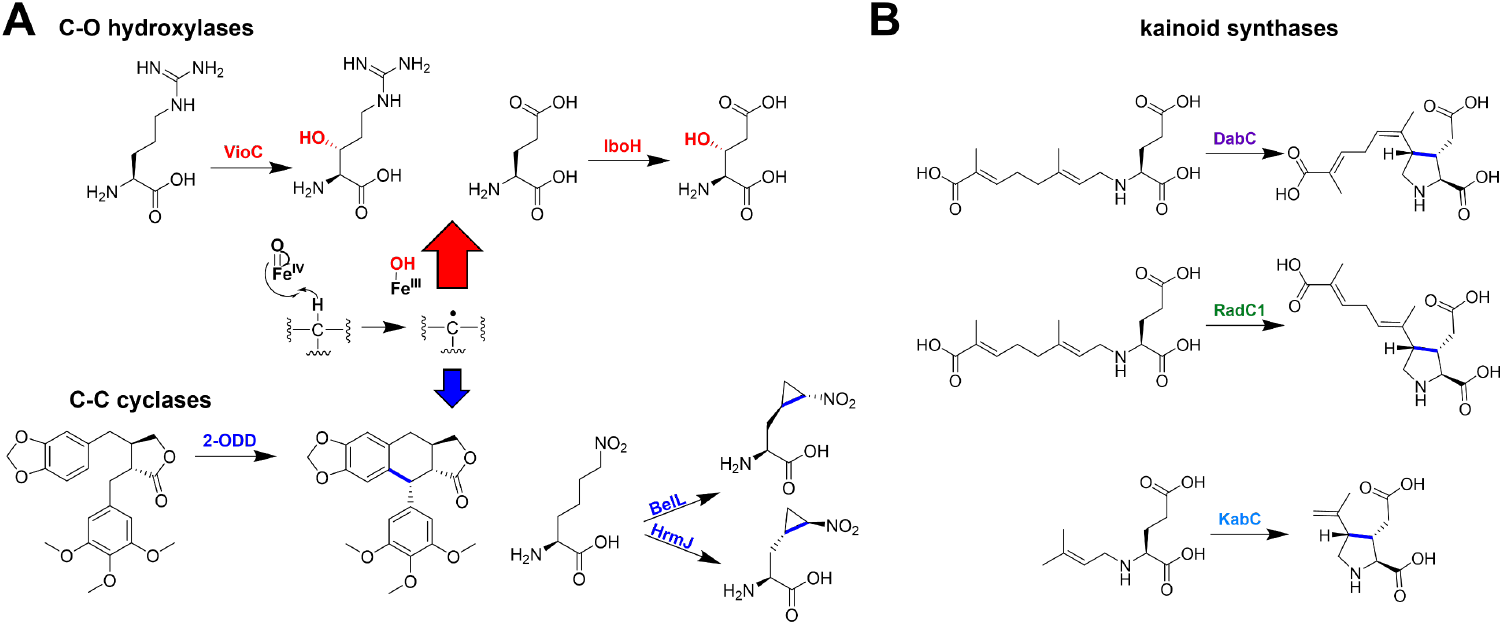
**A**. Select examples of Fe/αKG-catalyzed hydroxylation (top, red) and C-C cyclization (bottom, blue) reactions from various natural product biosynthetic pathways. **B**. Reactions of micro- and macroalgal kainoid synthases DabC, RadC1 and KabC.

DA is a potent neurotoxic natural product that agonistically binds ionotropic glutamate receptors (iGluRs) in the central nervous system, causing neuronal excitotoxicity through enhanced Ca^2+^ accumulation.^[14]^ The release of DA during marine harmful algal blooms has had an increasingly devastating environmental impact on wildlife; moreover, human exposure via consuming contaminated seafood causes amnesic shellfish poisoning.^[15]^ Seaweed-derived KA has a reduced agonistic effect and is employed experimentally to investigate epilepsy and other neurological disorders.^[12]^ The key structural feature imparting improved iGluR agonism is the pyrrolidine formed following the action of DabC and homologs. This entropically constrained kainoid ring stabilizes the glutamic acid moiety in an advantageous binding orientation, rationalizing its improved potency relative to its endogenous ligand.^[14]^ It is not clear how kainoid synthases divert Fe/αKG hydroxylation enzymology to preferentially build this bioactive C-C bond forming cyclization, leading to the basis of our current investigation. Mechanistic and structural insights into DabC and homologs not only improve our basic science understanding of Fe/αKG enzymology, but concurrently enable their downstream biocatalytic application towards diverse and putatively neuroactive kainoid scaffolds.

We initially aimed to investigate the range of non-native substrates DabC can recognize and transform, and adapted established protocols for heterologous DabC expression in *Escherichia coli* and purification via Ni-NTA affinity chromatography. Validated DabC substrate *trans*-7’-carboxygeranyl-*N-*L-glutamic acid (**1-*trans***) was synthesized alongside its minor *cis*-regioisomer byproduct (*cis-*7’-carboxyneryl-*N*-L-glutamic acid; **1-*cis***) following reductive amination of L-Glu with 7-carboxygeranial.^[11]^ In the presence of previously identified cofactors and co-substrates (Fe^2+^, αKG, L-ascorbate, and O2), DabC efficiently converted **1-*trans*** to major product isodomoic acid A (**2**) following *in vitro* assays and ultrahigh performance liquid chromatography – mass spectrometry (UPLC-MS) analyses.^[11]^ However, when employing regioisomer **1-*cis*** under identical reaction conditions, negligible cyclic and/or dehydrogenated product [M-2H] extracted ion chromatograms (EICs) were detected. Instead, a new putatively hydroxylated [M+O] product **3** was exclusively observed (Figure 2A). When a purified mixture of both substrate isomers (**1-mix)** was used for *in vitro* DabC assays, both cyclic **2** and hydroxylated **3** products were observed in comparable ratios to the individually purified regioisomers, thus simplifying our downstream workflow. A DabC reaction with **1-mix** was performed on 25 mg scale, isolating **3** after reversed-phase high performance liquid chromatography (RP-HPLC). Following nuclear magnetic resonance (NMR) spectroscopy and high-resolution mass spectrometry (HRMS), **3** was confirmed to be hydroxylated at the β-position of glutamic acid while maintaining the *cis-*configuration at the allyl amine. Intriguingly, *N-*geranyl-(3*R*)-hydroxy-L-glutamic acid was isolated in microgram quantities from domoic acid-producing *C. armata*;^[16]^ comparison of the glutamate moiety ^1^H and ^13^C NMR chemical shifts between **3** and this red algal metabolite supported that the β-hydroxy group was installed at the pro-*R* hydrogen (Tables S1, S2).^[16]^ To unequivocally assign the stereochemistry of **3**, 3*R*-hydroxy-L-glutamic acid was enantioselectively synthesized from L-malic acid,^[17,18]^ then condensed with 7‘-carboxygeranial via reductive amination. Regioisomers were separated by RP-HPLC and enzymatic **3** proved to be identical to synthetic 7’-carboxy-*N*-neryl-(3*R*)-hydroxy-L-glutamic acid by UPLC-MS retention time (Figure 2A) and NMR (Figure 2B, Tables S1, S2). This validates that Fe/αKG homolog DabC is capable of both hydroxylation and cyclization biochemistries with relatively conservative substrate alterations.

**Figure 2.**
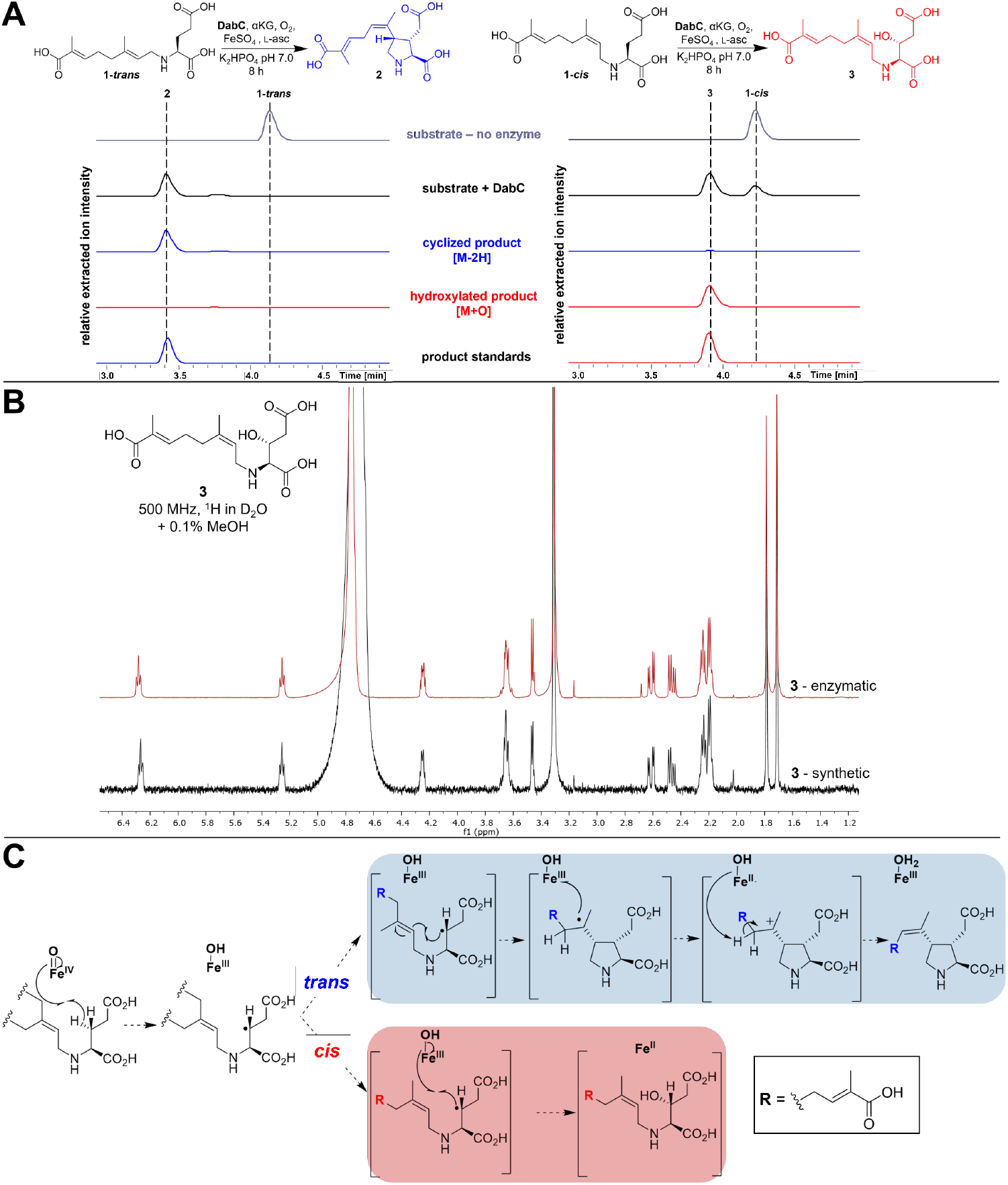
DabC preferentially hydroxylates the *cis* regioisomer of its native substrate. **A**. *In vitro* analytical DabC assays using **1-*trans*** (left) or **1-*cis*** (right) substrates. Relative intensities of negative mode extracted ion chromatograms were extracted from UPLC-MS traces (± 0.3 *m/z*) and show that **1-*trans*** ([M-H]^-^ 312.15 *m/z*) is converted to cyclic product **2** ([M-H]^-^ 310.12 *m/z*) whereas **1-*cis*** ([M-H]^-^ 312.15 *m/z*) instead forms β-hydroxylated product **3** ([M-H]^-^ 328.14 *m/z*). **B**. ^1^H NMR comparison of synthetic **3** (black) and enzymatic product **3** (red). **C**. Mechanistic proposal for divergent DabC activities with *cis* (red) and *trans* (blue) substrates.

To rationalize these divergent enzymatic functions and products, we propose that DabC initially abstracts the pro-*R* glutamic acid β-hydrogen on both substrates (Figure 2C). However, the mechanism then deviates depending on allyl amine stereochemistry. Non-native **1-*cis*** undergoes a canonical Fe/αKG hydroxyl rebound from the iron cofactor to generate stereoselectively β-hydroxylated **3**. In contrast, **1-*trans*** completes a radical 5*-exo-trig* cyclization to construct a novel carbon-carbon bond and set the last of the three contiguous stereocenters of the kainoid ring. Electron transfer to the Fe(III)-OH intermediate followed by regioselective deprotonation adjacent to the tertiary carbocation completes the ‘interrupted desaturase’ mechanism in the production of **2** (Figure 2C).^[11,19]^ To experimentally corroborate this mechanism, a glutamate-backbone *d5*-labeled mixture of both native substrate isomers (***d5*-1-mix**) was prepared. The overnight incubation of ***d5*-1-mix** with DabC indicated the loss of one deuterium within both cyclic (***d4*-2**) and hydroxylated (***d4*-3**) products by UPLC-MS (Figure S2), supporting that a single C-H radical abstraction occurs on the glutamate backbone without any adjacent desaturation in either mechanism. To assess DabC stereospecificity, the substrate enantiomer mixture was synthesized (**D-1-mix**) and incubated with DabC. No definitive cyclic, desaturated, or hydroxylated products could be detected by UPLC-MS, suggesting that DabC possesses a high degree of stereospecificity for both enzymatic functions (Figure S3).

To probe the catalytic differences between the DabC cyclization and hydroxylation reactions, *in vitro* kinetics assays were conducted with **1-*trans*** and **1-*cis***. Conditions were initially optimized with **1-mix** to improve overall conversion efficiency of DabC for both products in comparison to a chloramphenicol (CAM) internal standard. Buffer pH (Figure S4), cofactor (Figures S5, S6) and co-substrate concentrations (Figure S7) were independently assessed over a range of values using 5 mol% DabC and 14 h reaction conditions; individual end points that displayed the highest relative conversion for **2** and **3** were selected for *in vitro* kinetic assays. Optimized assay conditions (pH 6.0, 1.0 mM iron (II) sulfate, 50 mM α-ketoglutarate, 0.1 mM L-ascorbate) were compared to literature, and the hydroxylation reaction showed the most notable conversion improvement (Figure S8). After preparing standard curves for major products **2** and **3** (Figure S9), DabC steady-state kinetics experiments were individually performed with ***1-trans*** and ***1-cis*** substrates under a range of enzyme:substrate ratios, holding all other co-substrate and cofactor concentrations constant. Both regioisomers were individually fit to Michaelis-Menten kinetic parameters, revealing that DabC has a nearly 20-fold greater apparent kcat/KM for the cyclization of **1-*trans*** relative to the hydroxylation of **1-*cis*** (Figure 3). The binding affinity for **1-*trans*** was surprisingly not substantially different than **1-*cis*** (93 and 255 μM respectively); instead, the rate of catalysis (kcat) was the most significant contributor to the difference in catalytic efficiency between substrates. The 7-fold increase in turnover rate for **2** vs. **3** production (1700 and 250 min^-1^) was intriguing given the mechanistic complexity of **1-*trans*** carbon-carbon bond formation, electron transfer, and desaturation compared to the **1-*cis*** hydroxyl rebound.

**Figure 3.**
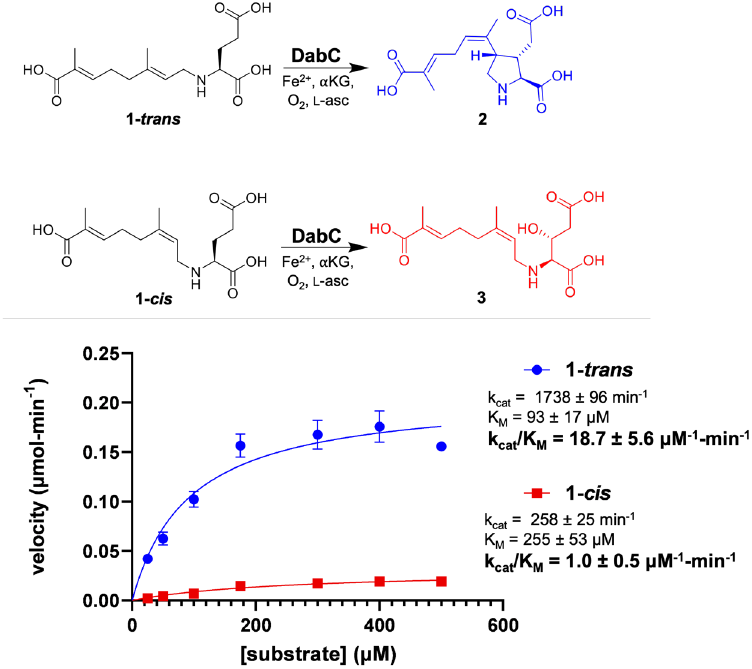
Michaelis-Menten kinetic parameters for the DabC-catalyzed cyclization reaction of **1-*trans*** (blue) compared to the β-hydroxylation reaction with **1-*cis*** (red).

We next wanted to assess how substrate analogs affect the extent of DabC conversion for both cyclization and hydroxylation. Previous efforts identified a pronounced reduction in kainoid product formation when interrogating DabC with *N-*geranylated L-glutamic acids lacking the 7’-carboxylic acid moiety.^[11,20]^ We wanted to see if modifications at the distal end would have a comparable impact on β-hydroxylation activity. Literature protocols were adapted to synthesize 7’-hydroxy (**4**) and 7’-methyl (**5**) modified *trans* (*N*-geranyl L-glutamic acid; L-NGG) and *cis* (*N*-neryl L-glutamic acid; L-NNG) substrate scaffolds.^[11]^ *In vitro* DabC assays followed by UPLC-MS analyses confirmed that **4-*cis*** and **5*-cis*** substrate analogs were preferentially hydroxylated (Figures S10, S11). The extent of conversion was assessed by comparing the intensity of the EIC for the major enzymatic product to the original substrate; this corroborated the 7’-substitution recognition trends, where a decrease in both cyclization and hydroxylation activities followed decreasing 7’-position oxidation (Figures 4A, S9). The recent characterization of red macroalgal homolog RadC1 (63% DabC sequence identity), which preferentially cyclizes **1-*trans*** to isodomoic acid B (Figure 1B), provided an opportunity to assess whether kainoid synthases from biologically diverse organisms exhibit similar 7’-position reactivity trends. After the heterologous expression, purification, and *in vitro* interrogation with our focused 7’-modified L-NGG and L-NNG substrate panel, *C. armata* RadC1 also exhibited substrate-dependent biochemistries. All examined *trans* isomers (**1-*trans*, 4-*trans*, 5-*trans***) were converted by RadC1 into putative cyclic and/or dehydrogenated [M-2] products, while *cis* isomers (**1-*cis*, 4-*cis*, 5-*cis***) were hydroxylated following UPLC-MS analyses (Figures 4A, S12). Two major differences were observed: RadC1 preferred both 7’-methyl-substrates (**5-*trans*, 5-*cis***) under our optimized *in vitro* conditions; and **4-*cis*** was effectively hydroxylated by DabC (62 ± 2%) but less tolerated by RadC1 (8 ± 2%) (Figures 4A, S11, S12). While the structural basis of 7’-recognition is still under investigation, we were encouraged to see that similar reactivity patterns emerged for both homologs.

**Figure 4.**
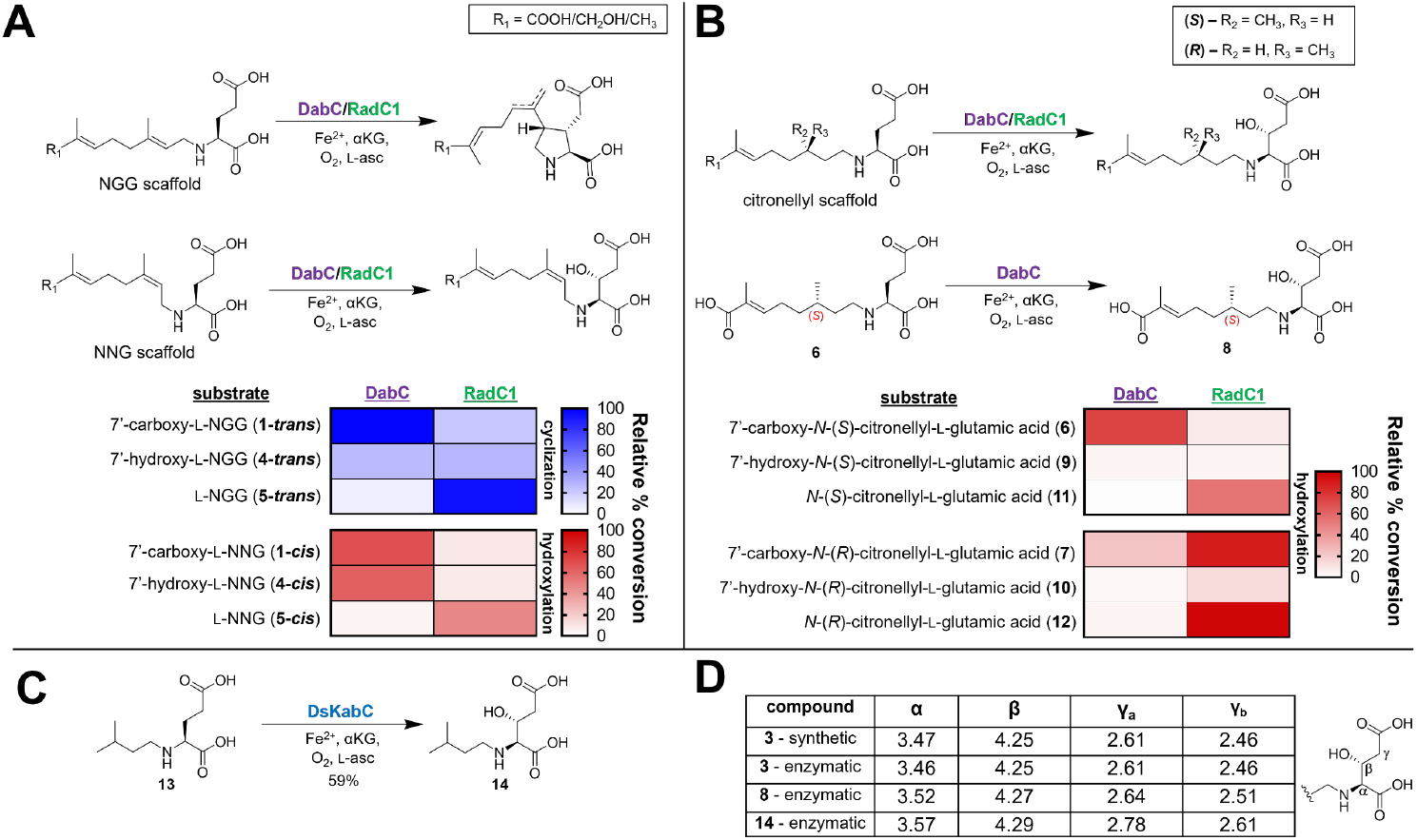
Hydroxylation activity on alternative substrates is conserved across kainoid synthase homologs DabC, RadC1, and KabC. **A**. General reaction schemes for *in vitro* reactions of NGG and NNG substrate analogs by DabC and RadC1, with heat maps indicating the relative % conversion to cyclic (blue) or β-hydroxylated products (red). **B**. General reaction scheme for *in vitro* reactions of citronellyl substrate analogs by DabC and RadC1 with scaled up DabC production of **8** shown underneath. Heat maps indicating the relative % conversion of *N*-citronellyl analogs to β-hydroxylated products. **C**. *In vitro* β-hydroxylation of *N*-isopentyl-L-glutamic acid (13) by DsKabC. **D**. Comparison of ^1^H NMR shifts for the glutamic acid moiety in isolated β-hydroxylated products.

The 5-*exo*-*trig* cyclization mechanism catalyzed by DabC and its homologs involves the radical interception of the substrate 2’ allyl amine.^[11,19]^ We hypothesized that the removal of this alkene would divert kainoid synthases to exclusively hydroxylate substrates. Given the utility of intramolecular 1,5-hydrogen atom transfer (1,5-HAT) in chemosynthetic^[21]^ and Fe/αKG biocatalytic reactions,^[22]^ and propensity of DabC to cyclize **1-*trans***, it was unclear if regioselective β-hydroxylation or 1,5-HAT would predominate with saturated analogs (Figure S13). We leveraged the available citronellol enantiomers to divergently synthesize two novel 7’-carboxy-*N*-citronellyl-L-glutamic acid substrates with *S* (**6**) or *R* (**7**) 3’-methyl stereochemistries. *In vitro* interrogation of DabC and RadC1 with **6** and **7** exclusively showed a new hydroxylated product in the presence of all cofactors and co-substrates (Figures 4B, S14, S15). To validate the regioselectivity of product hydroxylation, the DabC assay with **6** was scaled up, purified via RP-HPLC, and characterized as 7’-carboxy-*N*-(*3’S*)-citronellyl-3*R*-hydroxy-L-glutamic acid (**8**) following NMR and HRMS (Figure 4B). The two tested homologs had an intriguing conversion preference – DabC preferentially hydroxylated *S* isomer **6**, whereas **7** was much better tolerated by RadC1. Given the improved *in vitro* conversion of RadC1 with **5-*trans*** and **5-*cis***, we used *S* or *R* citronellal enantiomers to divergently synthesize four additional saturated analogs with 7’-hydroxy or 7’-methyl end permutations (**9 – 12**). UPLC-MS analyses of *in vitro* assays revealed that *N*-citronellyl substrate analogs were hydroxylated by both kainoid synthases, mirroring both the 7’-substitution and 3’-methyl stereochemical trends observed with **1/4/5** and **6/7** substrate analogs respectively (Figures 4B, S14, S15). Comparison of DabC and RadC1 AlphaFold models with docked **1-*trans*** identified highly conserved amino acid residues in putatively structured regions surrounding the 3’-methyl and 7’-positions; variability in isodomoic acid regioisomer production is presumably due to pocket shape and/or second shell differences as previously proposed (Figure S16).^[13]^ To extend our investigation to *D. simplex* KabC, (DsKabC; 60% DabC sequence identity), saturated analog *N-*isopentyl-L-glutamic acid (**13**) was synthesized and incubated with the KA-producing Fe/αKG homolog.^[12]^ UPLC-MS analysis confirmed the presence of a hydroxylated product, identified as *N*-isopentyl-(3*R*)-hydroxy-L-glutamic acid (**14**) following NMR and HRMS characterization (Figures 4C, S17). Hydroxylated product EICs were previously observed following DsKabC incubation with *N*-isopentyl and *N*-benzyl L-glutamic acids, however neither were structurally validated.^[19]^ Comparison of enzymatic **3, 8**, and **14** showed highly similar glutamate backbone NMR chemical shift and *J* values despite the variable *N-*substituents, supporting the relative β-position stereochemistry (Figure 4D, Tables S1, S2). Our cumulative kinetic and alternative substrate results suggest that intramolecular cyclization reaction is fastest when substrate stereochemistry permits, however, intermolecular hydroxyl rebound occurs more quickly than 1,5 HAT in saturated substrates. These results additionally confirm that the unique dual cyclization and hydroxylation activity is conserved across kainoid synthases from diverse biological species and biosynthetic pathways and exemplifies the use of non-native substrates to interrogate wild-type enzyme mechanisms.

The substrate-dependent chemoselectivity of kainoid synthases, though uncommon, has precedent within the Fe/αKG enzyme family. L-norvaline halogenase SyrB2 from *Pseudomonas syringae* is capable of hydroxylation on both L-threonine and L-aminobutyric acid.^[23]^ Fungal L-valine/L-isoleucine aziridinase TqaL similarly shows stereoselective hydroxylation activity on a simplified substrate analog.^[24,25]^ Substantial engineering efforts have improved the interconversion of select Fe/αKG hydroxylases into halogenases; broader application to other homologs and substrates is under active investigation.^[26–28]^ Given the broad utility of Fe/αKGs to regio- and stereoselectively functionalize C-H bonds on chemically complex scaffolds, members of this enzyme family have seen extensive biocatalytic application. From the syntheses of bioactive *ent*-kaurane diterpenoid^[29]^ and tropolone natural products,^[30]^ the preparation of isotopically enriched standards for the quantifiable detection of cyanotoxin cylindrospermopsin,^[31]^ or the scalable biotransformation of KA,^[12]^ chemoenzymatic methods using Fe/αKGs help circumvent synthetic challenges and improve overall efficiency in the production of natural products, pharmaceuticals, and small molecule tools. The dual functionality seen in the algal kainoid synthases showcases their flexible chemoselectivities on relatively conservative substrate analogs. Further investigation into these unique homologs will provide additional insight into the understanding, engineering, and application of the chemically useful and diverse Fe/αKG family.

## Supporting information

DabC-SI-PDF

## Acknowledgements

We gratefully acknowledge Prof. Laura M. Sanchez and Dr. Gordon T. Luu for assistance obtaining high-resolution mass spectrometry data, Dr. H.-W. Lee for maintenance of nuclear magnetic resonance spectroscopy facilities, Prof. John B. MacMillan for access to high performance liquid chromatography, and the University of California, Santa Cruz for financial support in the form of startup funding.

## Table of Contents Entry

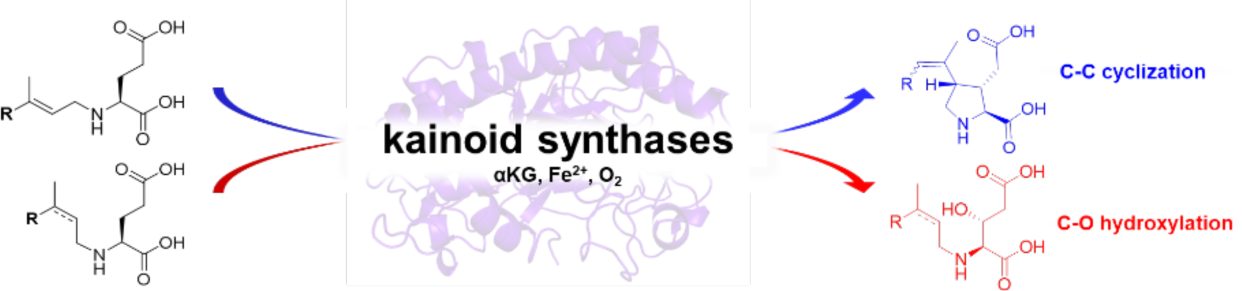

*Trans* isomer – neurotoxin; *cis* isomer – adds an oxygen. Kainoid synthases are a unique family of algal Fe^II^/α-ketoglutarate-dependent dioxygenases that form a novel C-C bond within domoic and kainic acid biosyntheses; however, they instead catalyze canonical hydroxylation reactions on substrate analogs. Kinetic, mechanistic, and substrate impacts on catalysis are explored, enhancing our understanding of these unique dual-functioning enzymes.

