## Supplementary material for "Algal kainoid synthases exhibit substrate-dependent hydroxylation and cyclization activities": DabC-SI-PDF

**This PDF file includes:**

General Materials and Methods

Figures S1 to S17

Chemical Synthesis

NMR and Compound Characterization

Tables S1 to S3

References

#### Table of Contents


**General Materials & Methods** **5**

Synthetic and analytical methods **5**
LCMS & HPLC instrumentation **5**

**1. Molecular Biology/Biochemical Methods** **6**

*dabC* transformation into *E. coli* **6**
DabC expression in BL21(DE3) *E. coli* **6**
DabC FPLC affinity purification **6**
Cloning of *radC1* with N-terminal-$His_6$ tag **7**
DsKabC and RadC1 expression in BL21(DE3) *E. coli* **7**
DsKabC FPLC affinity purification **7**
RadC1 FPLC affinity purification **7**
Analytical DabC/RadC1 assays with *N*-prenylated-L-glutamic acid substrates & analogs **7**
Preparative DabC assays for product isolation and NMR characterization **8**
Enzymatic synthesis of 7'-carboxy-*N*-neryl-(3*R*)-hydroxy-L-glutamic acid (**3**) **8**
HPLC purification of 7'-carboxy-*N*-neryl-(3*R*)-hydroxy-L-glutamic acid (**3**) **8**
Enzymatic synthesis of 7'-carboxy-*N*-(3'*S*)-citronellyl-3*R*-hydroxy-L-glutamic acid (**8**) **8**
HPLC purification of 7'-carboxy-*N*-(3'*S*)-citronellyl-3*R*-hydroxy-L-glutamic acid (**8**) **9**
Enzymatic synthesis of *N*-isopentyl-(3*R*)-hydroxy-L-glutamic acid (**14**) **9**
HPLC purification of *N*-isopentyl-(3*R*)-hydroxy-L-glutamic acid (**14**) **9**
DabC optimization assays **9**
Establishment of standard curve for DabC kinetics assays **10**
DabC kinetics assays **10**

**2. Supplementary Figures & Tables** **11**

Fig. S1. Notable examples of native hydroxylation and C-C cyclization reactions catalyzed by non-heme $Fe^{II}$/$\alpha$KG-dependent dioxygenases **11**
Fig. S2. *In vitro* DabC assay with ***d₅*-1-mix** substrate **12**
Fig. S3. *In vitro* DabC assay with **D-1-mix** substrate **13**
Fig. S4. *In vitro* DabC pH optimization of **1-mix** cyclization and hydroxylation **14**
Fig. S5. *In vitro* DabC $FeSO_4$ optimization of **1-mix** cyclization and hydroxylation **15**
Fig. S6. *In vitro* DabC $\alpha$KG optimization of **1-mix** cyclization and hydroxylation **16**
Fig. S7. *In vitro* DabC L-ascorbate optimization of **1-mix** cyclization and hydroxylation **17**
Fig. S8. *In vitro* DabC assay of **1-mix** cyclization and hydroxylation pre- and **18**

post-optimization
Fig. S9. Standard curves of DabC products **2** and **3** from UPLC-MS analyses **19**
Fig. S10. DabC and RadC1 activity assays on 7’-modified *trans* and *cis* substrate analogs comparing cyclized and hydroxylated product mass EICs. **20**
Fig. S11. DabC activity assay - *trans* vs *cis* substrate analogs **23**
Fig. S12. RadC1 activity assay - *trans* vs *cis* substrate analogs **25**
Fig. S13. Alternative HAT mechanistic proposal for saturated substrate hydroxylation **27**
Fig. S14. DabC activity assay - *R* vs *S* citronellyl substrate analogs **28**
Fig. S15. RadC1 activity assay - *R* vs *S* citronellyl substrate analogs **30**
Fig. S16 DabC/RadC1 AlphaFold model comparison and **1-*trans*** substrate docking **32**
Fig. S17. *In vitro* DsKabC hydroxylation of saturated substrate isopentyl-*N*-L-glutamic acid (**13**) **34**

#### 3. Chemical Synthesis **35**

Synthesis of 7’-carboxy- L-NGG (**1-*trans***) and 7’-carboxy-L-NNG (**1-*cis***) **35**
Synthesis of **D-1-mix** and ***d*$_5$-1-mix** **36**
Synthesis of 7’-hydroxy- L-NGG (**4-*trans***) and 7’-hydroxy-L-NNG (**4-*cis***) **37**
Synthesis of L-NGG (**5-*trans***) and L-NNG (**5-*cis***) **38**
Synthesis of 7’-carboxy-(*S*)-citronellyl-*N*-L-glutamic acid (**6**) and 7’-carboxy-(*R*)-citronellyl-L-glutamic acid (**7**) **40**
Synthesis of 7’-hydroxy-(*S*)-citronellyl-*N*-L-glutamic acid (**9**) and 7’-hydroxy-(*R*)-citronellyl-L-glutamic acid (**10**) **43**
Synthesis of (*S*)-citronellyl-*N*-L-glutamic acid (**11**) and (*R)*-citronellyl-*N*-L-glutamic acid (**12**) **45**
Synthesis of 3*R*-hydroxy-L-glutamic acid (**SI-9**) **46**
Synthesis of 7’-carboxy-*N*-neryl-(3*R*)-hydroxy-L-glutamic acid (**3**) **47**
Synthesis of *N*-isopentyl-L-glutamic acid (**13**) **48**

#### 4. Tables

Table S1. Glutamic acid moiety $^{1}$H NMR shifts of β-hydroxylated molecules in this study **49**
Table S2. Glutamic acid moiety $^{13}$C NMR shifts of β-hydroxylated molecules in this study **50**
Table S3. Primers used in this study **50**

#### 5. NMR & Compound Characterization 51


7’-carboxy-L-NGG – synthetic (**1-*trans***) 51
7’-carboxy-L-NNG – synthetic (**1-*cis***) 54
isodomoic acid A – enzymatic (**2**) 57
7’-carboxy-*N*-neryl-(3*R*)-hydroxy-L-glutamic acid – synthetic (**3**) 58
7’-carboxy-*N*-neryl-(3*R*)-hydroxy-L-glutamic acid – enzymatic (**3**) 61
7’-hydroxy- L-NGG – synthetic (**4-*trans***) 64
7’-hydroxy- L-NNG – synthetic (**4-*cis***) 67
L-NGG – synthetic (**5-*trans***) 70
L-NNG – synthetic (**5-*cis***) 73
7’-carboxy-*N*-(*S*)-citronellyl-L-glutamic acid – synthetic (**6**) 76
7’-carboxy-*N*-(*R*)-citronellyl-L-glutamic acid – synthetic (**7**) 79
7’-carboxy-*N*-(3’*S*)-citronellyl-3*R*-hydroxy-L-glutamic acid – enzymatic (**8**) 82
7’-hydroxy-*N*-(*S*)-citronellyl-L-glutamic acid – synthetic (**9**) 85
7’-hydroxy-*N*-(*R*)-citronellyl-L-glutamic acid – synthetic (**10**) 88
(*S*)-citronellyl-*N*-L-glutamic acid – synthetic (**11**) 91
(*R)*-citronellyl-*N*-L-glutamic acid – synthetic (**12**) 94
*N*-isopentyl-L-glutamic acid – synthetic (**13**) 97
*N*-isopentyl-(3*R*)-hydroxy-L-glutamic acid – enzymatic (**14)** 100
3*R*-hydroxy-L-glutamic acid – synthetic (**SI-9**) 103


#### 6. References 106

#### General Materials & Methods

All chemicals, including solvents and media components, were used as received from commercial suppliers (Millipore Sigma, Thermo Fisher Scientific). Reagents were used without further purification unless otherwise noted.

##### Synthetic and analytical methods

Anhydrous reactions were done in oven-dried glassware and carried out under an argon atmosphere. Reactions were monitored using thin-layer chromatography (TLC, Merck, 60 $F_{254}$). TLC plates were visualized with a UV lamp at 254 nm and stained with a potassium permanganate staining solution (1.5 g $KMnO_4$, 10 g $K_2CO_3$, 1.25 mL 10% NaOH in 200 mL water). TLC $R_f$ values were rounded to the nearest 0.05. Silica ((Alfa Aesar, 60 (215-400 mesh)) was used when flash column chromatography was employed to purify compounds. Rotary evaporation was used to concentrate samples under reduced pressure. Additionally, polar final products were concentrated and dried via lyophilization (Labconco FreeZone 4.5 Liter). Nuclear magnetic resonance (NMR) spectra were obtained using an Avance III HD spectrometer (Bruker) equipped with a BBFO SmartProbe at 500 MHz ($^1$H NMR) or 125 MHz ($^{13}$C NMR) using $CDCl_3$ or $D_2O$ as solvents. Chemical shifts (δ) are reported in ppm and referenced either to methanol (δ = 3.31 ppm for $^1$H, δ = 49.0 ppm for $^{13}$C NMR) or formic acid (δ = 8.26 ppm for $^1$H, δ = 166.3 ppm for $^{13}$C NMR) as an internal standard for samples in $D_2O$ or the $CDCl_3$ solvent signal (δ = 7.26 ppm for $^1$H, δ = 77.2 ppm for $^{13}$C NMR) for samples in $CDCl_3$. NMR data are reported as follows: s = singlet, d = doublet, t = triplet, q = quartet, m = multiplet, *J* = coupling constant in Hz.

##### LCMS & HPLC instrumentation

General LCMS measurements were measured on a Bruker Elute UHPLC system coupled with a Bruker amazon SL ESI-Ion Trap mass spectrometer in negative mode. Compounds were separated via reversed-phase chromatography on a Bruker Intensity Solo C18(2), 2 µm- 2 x 100 mm column with the eluents water + 0.1% formic acid (Solvent A) and acetonitrile + 0.1% formic acid (Solvent B). The LC method uses a flow rate of 0.3 mL/min and the following gradient: 10% to 20% B over 3 minutes, 20% to 45% B over 3 min, 45% - 100% B over 2 min, hold at 100% B for 2 min, 100% - 10% B over 1 min, hold at 10% B for 2 min.

Preparative and semi-preparative HPLC purification was performed using a Shimadzu Prominence preparative HPLC system with a SPD-20A model UV/Vis detector. Analytical HPLC purification was performed using an Agilent 1260 Infinity system with a G1314B model VWD. The monitored wavelength was 210 nm for all runs. High-performance flash chromatography was performed using a Biotage Isolera Prime system with a dual variable UV detector (200 - 400 nm) monitored at 210 nm and 254 nm.

#### Molecular Biology/Biochemical Methods

*dabC* transformation into *E. coli*

A pET28a(+) plasmid containing the N-terminal hexahistidine ($His_6$) tagged *Pseudo-nitzschia multiseries dabC* gene (*dabC*) was generated as previously described in the literature[1] and transformed into *E. coli* DH10β chemically competent cells for storage and BL21(DE3) for expression. The transformation by heat shock proceeded according to the following protocol: 0.5 µL of plasmid was added to the chemical competent cells and maintained on ice for 30 minutes. After this, the cells were heated to 42 °C for 45 sec and placed in the ice again for 3 min; 700 µL of LB medium was added in the tube and the cells were incubated for 1 h at 37 °C and 200 rpm of agitation. After this step, the cells were plated on LB agar plates supplemented with kanamycin. The plates were incubated at 37 °C, overnight. An inoculum with colonies was grown overnight in the same condition of LB agar plates at 37 °C and 200 rpm of agitation and purified following the protocol of Plasmid DNA Purification QIAprep Spin Miniprep Kit (QIAGEN). After plasmid purification, concentrations were measured by NanoDrop UV-vis spectrophotometry and stored at -20 °C.

DabC expression in BL21(DE3) *E. coli*

Either a colony of DabC transformed BL21(DE3) cells or a glycerol stock of said cells was used to inoculate 30 mL of LB media supplemented with kanamycin (50 µM). The culture was grown overnight at 37 °C, 200 rpm. 10 mL of this overnight culture was added per liter of Terrific Broth supplemented with kanamycin (50 µM). Each liter of culture was grown in a shaking incubator at 37 °C, 200 rpm until $OD_{600}$ reached ~0.6. The flasks were then cooled to 18 °C over 45 - 60 min at which point 0.5 mM IPTG was added to induce protein expression. The cultures were allowed to grow at 18 °C and 200 rpm overnight (16 - 18 h) and pelleted in a centrifuge at approx. 3500 *x g* for 30 min. Supernatant was discarded and the cell pellet was resuspended in 25 mL of lysis buffer (1 M NaCl, 20 mM Tris-HCl pH 8.0, 30 mM imidazole, and 10% glycerol) and stored at -70 °C. If cells were to be immediately purified, the pellet was resuspended in buffer A (1 M NaCl, 20 mM Tris-HCl pH 8.0, 30 mM imidazole) and put on ice.

DabC FPLC affinity purification

DabC resuspended cell pellet was lysed on ice via sonication at 40% amplitude alternating between 15 sec on and 45 sec off (5 minutes total 'on' time). The lysate was pelleted via centrifugation at 4 °C and ~16,000 *x g* for 40 min. The cleared lysate was loaded onto a 5 mL HisTrap FF column (GE Healthcare Life Sciences) that was pre-equilibrated with at least 5 CV of buffer A. After loading, the column was washed with buffer A until the UV absorbance plateaued and reached approximately 50 mAU. The column was then washed with 10% buffer B for 5 - 10 CV to remove any loosely bound proteins. DabC was then eluted using a linear gradient of 10% to 100% buffer B over 60 mL while collecting 5 mL fractions. The flowrate through the column was 2 mL/min throughout the purification process. Fractions were assessed for purity using SDS-PAGE. Fractions containing DabC had 2 mM EDTA added to remove any bound metals and were combined and further purified through a PD-10 desalting column (GE Healthcare Life

Sciences) pre-equilibrated with GF buffer (20 mM HEPES pH 8.0, 300 mM KCl) following the default gravity filtration protocol. For enzyme kinetics assays, the protein was further purified on a Superdex 75 HiLoad 16/600 size exclusion column (GE Healthcare Life Sciences) equilibrated in GF buffer. 400 µL of protein was injected onto the column and eluted with GF buffer at a rate of 1 mL/min. The protein was concentrated by Amicon ultra centrifugal filters (30 kDa molecular weight cut-off). Enzyme concentration was determined via Bradford assay and pure enzyme was either used immediately for assays or aliquoted and stored at -70 °C.

Cloning of *radC1* with N-terminal-$His_6$ tag

The enzyme coding sequence for *radC1* was optimized for expression in *E. coli* and synthesized by Twist Bioscience. The synthetic *radC1* gene was sub cloned into the pET28a(+) kanamycin resistant expression vector containing an N-terminal hexahistidine ($His_6$) tag using the primers listed in Table S1. Amplification of the *rad*C1 gene insert was done using Primestar HS DNA polymerase (Takara Bio USA Inc.). To prepare linearized pET28a(+) backbone, a restriction enzyme digest was performed on empty pET28a(+) with the restriction enzymes NcoI and XhoI (New England Biolabs) at 37 °C for 30 min, then 80 °C for 20 min to heat inactivate the restriction enzymes. Once the gene insert and linear pET28a(+) were prepared, a Gibson assembly was performed using HiFi Assembly polymerase mix (New England Biolabs) and incubated at 55 °C for 30 minutes, and transformed into *E. coli* DH10β chemically competent cells. The *radC1* plasmid was further amplified and purified in the same manner as described for *dabC* above, and the sequence of *radC1* was confirmed via Sanger sequencing (Azenta Life Sciences). Once the sequence was confirmed, the plasmid was further transformed into *E. coli* BL21(DE3) cells for protein expression as previously described for *dabC*.

DsKabC and RadC1 expression in BL21(DE3) *E. coli*

Expression of DsKabC was carried out as was described previously for DabC. Expression of RadC1 was carried out as was described previously for DabC except 1.0 mM of IPTG was used instead of 0.5 mM.

DsKabC FPLC affinity purification

Purification of DsKabC was carried out as previously described for DabC without modification.

RadC1 FPLC affinity purification

Purification of RadC1 was carried out as previously described for DabC with the following modifications: Sonication was done for 20 min alternating between 15 sec on and 45 sec off, with gentle mixing after 15 min. Buffer A used was 20 mM Tris-HCl pH 8.0, 500 mM NaCl, 10 mM imidazole. Buffer B used was 20 mM Tris-HCl pH 8.0, 500 mM NaCl, 250 mM imidazole.

Analytical DabC/RadC1 assays with *N*-prenylated-L-glutamic acid substrates & analogs

Analytical DabC enzyme assays were conducted in 50 mM potassium phosphate (KPi) buffer (pH 7.0) with 1 mM L-ascorbate, 6.25 mM $\alpha$-ketoglutarate, and 50 µM $FeSO_4$ with

1 mM of substrate and 25 μM of purified DabC or RadC1. Enzyme followed by $FeSO_4$ were the last additions to assay and the reaction was allowed to run for 8 h. The total volume for the reaction was 100 μL. The reaction was then quenched with 0.01 mM chloramphenicol in methanol (1 eq.) and centrifuged at ~ 21,000 *x g* for 5 min. The supernatant was then subjected to analysis by UPLC-MS.

Preparative DabC assays for product isolation and NMR characterization

Preparative assays were conducted in the same manner as the small-scale assays except the reaction volume was increased to either 0.5 mL, 1.0 mL or 10 mL. Multiple reactions were set up simultaneously, the number of which was dependent on the desired quantity of starting material for the reaction.

Enzymatic synthesis of 7'-carboxy-*N*-neryl-(3*R*)-hydroxy-L-glutamic acid (**3**)

Preparative DabC assays were conducted as previously described in 8 x 10 mL volumes using a total of 0.025 g **1-mix** as a substrate with light stirring. The reactions were quenched as previously described after 18 h, centrifuged to pellet precipitated protein, concentrated *in vacuo*, and subjected to RP-HPLC purification.

HPLC purification of 7'-carboxy-*N*-neryl-(3*R*)-hydroxy-L-glutamic acid (**3**)

Compound **3** was purified via preparative RP-HPLC (Phenomenex Kinetex 5 μm EVO C18 100 Å 250 x 21.2 mm) at a flow rate of 8.5 mL/min using the following method: 10% B (5 min), 10 - 27% B (15 min), 27 - 95% B (2 min), 95% B (4 min), 95 - 10% B (1 min), 10% B (4 min), where A = 0.1% aqueous formic acid, and B = 0.1% formic acid in acetonitrile. The peak containing **3** was manually collected (retention time 16.5 min), concentrated *in vacuo* and lyophilized. The sample was then further purified via semi-preparative RP-HPLC (Phenomenex Luna 5 μm C18(2) 100 Å 250 x 10 mm) at a flow rate of 3.0 mL/min using 10.5 % acetonitrile in 0.1% aqueous formic acid. Compound **3** was manually collected (retention time 35.5 min), concentrated *in vacuo* and lyophilized to afford the product **3** as a white solid (0.001 g, 0.003 mmol, 4%). $^1$H NMR (500 MHz, $D_2O$ + 0.1% MeOH): δ 6.28 (t, *J* = 6.9 Hz, 1H), 5.26 (t, *J* = 7.5 Hz, 1H), 4.25 (dt, *J* = 7.4, 4.9 Hz, 1H), 3.70 – 3.60 (m, 2H), 3.46 (d, *J* = 5.5 Hz, 1H), 2.61 (dd, *J* = 15.9, 4.5 Hz, 1H), 2.46 (dd, *J* = 15.9, 7.4 Hz, 1H), 2.28 – 2.15 (m, 4H), 1.79 (s, 3H), 1.71 (s, 3H); $^{13}$C NMR (125 MHz, $D_2O$ + 0.01% MeOH): δ 178.9, 178.0, 172.0, 147.0, 136.5, 133.7, 114.2, 67.6, 65.2, 44.3, 42.2, 30.6, 26.3, 22.9, 13.0; HRMS (ESI) Calculated for $C_{15}H_{24}NO_7$ 330.1547, found 330.1555 $(M+H)^+$.

Enzymatic synthesis of 7'-carboxy-*N*-(3'*S*)-citronellyl-3*R*-hydroxy-L-glutamic acid (**8**)

Preparative DabC assays were conducted as previously described in 8 x 0.5 mL volumes using a total of 1.3 mg **6** as a substrate with light stirring. The reactions were quenched as previously described after 18 h, centrifuged to pellet precipitated protein, concentrated *in vacuo*, and subjected to RP-HPLC purification.

HPLC purification of 7'-carboxy-*N*-(3'*S*)-citronellyl-3*R*-hydroxy-L-glutamic acid (**8**)
Compound **8** was purified via semi-preparative RP-HPLC (Phenomenex Luna 5 μm C18(2) 100 Å 250 x 10 mm) at a flow rate of 3.0 mL/min using 16% acetonitrile in 0.1 % formic acid. The peak containing **8** was manually collected (retention time 15.1 min), concentrated *in vacuo* and lyophilized to afford the product **8** as a white solid (0.001 g, 0.003 mmol, 75%). $^1$H NMR (500 MHz, $D_2O$ + 0.1% MeOH): δ 6.35 – 6.30 (m, 1H), 4.27 (dt, *J* = 7.2, 4.9 Hz, 1H), 3.52 (d, *J* = 5.3 Hz, 1H), 3.13 – 2.97 (m, 2H), 2.64 (dd, *J* = 16.0, 4.6 Hz, 1H), 2.51 (dd, *J* = 16.0, 7.0 Hz, 1H), 2.19 – 2.07 (m, *J* = 7.1 Hz, 2H), 1.73 (s, 3H), 1.59 – 1.48 (m, 2H), 1.46 – 1.38 (m, 1H), 1.31 – 1.22 (m, 1H), 0.90 (d, *J* = 6.0 Hz, 3H); $^{13}$C NMR (125 MHz, $D_2O$ + 0.1% MeOH): δ 178.7, 178.3, 172.0, 137.9, 132.7, 67.6, 67.0, 49.0, 46.1, 34.0, 32.6, 29.9, 25.4, 18.4, 13.0; HRMS (ESI) Calculated for $C_{15}H_{26}NO_7$ 332.1704, found 332.1714 $(M+H)^+$.

Enzymatic synthesis of *N*-isopentyl-(3*R*)-hydroxy-L-glutamic acid (**14**)
Preparative DsKabC assays for the enzymatic synthesis of **14** were conducted in 50 mM KPi buffer (pH 8.0) with 1 mM L-ascorbate, 6.25 mM α-ketoglutarate, and 50 μM $FeSO_4$ with 1 mM of substrate and 50 μM of purified DsKabC. Enzyme followed by $FeSO_4$ were the last additions to assay. A total of 23 x 1 mL reactions using a total of 4.3 mg **13** as a substrate were set up with light stirring. The reactions were quenched with equimolar methanol after 15 h, centrifuged to pellet precipitated protein, concentrated *in vacuo*, and subjected to RP-HPLC purification.

HPLC purification of *N*-isopentyl-(3*R*)-hydroxy-L-glutamic acid (**14**)
Compound **14** was purified via semi-preparative RP-HPLC (Phenomenex Luna 5 μm C18(2) 100 Å 250 x 10 mm) at a flow rate of 3.0 mL/min using the following method: 5% B (4 min), 5 - 15% B (14 min), 15 - 95% B (1 min), 95% B (4 min), 95 - 5% B (1 min), 5% B (4 min), where A = 0.1% aqueous formic acid, and B = 0.1% formic acid in acetonitrile. The peak containing **14** was manually collected (retention time 15.5 min), concentrated *in vacuo* and lyophilized to afford the product **14** as a white solid (0.003 g, 0.012 mmol, 59%). $^1$H NMR (500 MHz, $D_2O$ + 0.1% MeOH): δ 4.29 (td, *J* = 7.7, 3.9 Hz, 1H), 3.57 (d, *J* = 7.2 Hz, 1H), 3.06 (td, *J* = 8.0, 2.6 Hz, 2H), 2.78 (dd, *J* = 16.3, 3.9 Hz, 1H), 2.61 (dd, *J* = 16.3, 8.2 Hz, 1H), 1.69 – 1.60 (m, 1H), 1.60-1.54 (m, 2H), 0.88 (dd, *J* = 6.6, 3.8 Hz, 6H); $^{13}$C NMR (125 MHz, $D_2O$ + 0.1% MeOH): δ 175.4, 171.3, 66.7, 66.7, 46.2, 39.9, 34.2, 25.4, 21.6, 21.4; HRMS (ESI) Calculated for $C_{10}H_{20}NO_5$ 234.1336, found 234.1336 $(M+H)^+$.

DabC optimization assays
DabC optimization assays (Figs. S4 – S7) were performed in the manner described above for analytical DabC assays with *N*-prenylated-L-glutamic acid substrates & analogs with the following modifications: 0.1 mM of substrate **1-mix**, 5 μM DabC, and 150 μL total reaction volumes were used. Reactions were left for ~14 h prior to quenching. All reaction component concentrations were held constant other than the component being optimized

in each assay. For the final assay comparing optimized conditions to literature conditions (Fig. S8), the concentration of each component showing the greatest product formation in each individual optimization assay was used (KPi pH 6.0, 0.1 mM L-ascorbate, 50 mM α-ketoglutarate disodium salt, 1.0 mM $FeSO_4$). The literature conditions used for comparison were the following: KPi pH 8.0, 1.0 mM L-ascorbate, 6.25 mM α-ketoglutarate disodium salt, 0.05 mM $FeSO_4$. Reactions were quenched after 1 h.

<u>Establishment of standard curve for DabC kinetics assays</u>

To establish standard curves for both DabC products for subsequent kinetics assays, 50 µL samples with were prepared in triplicate with the determined optimal DabC conditions (KPi pH 6.0, 0.1 mM L-ascorbate, 50 mM α-ketoglutarate, 1.0 mM $FeSO_4$), except the DabC enzyme was excluded. Each sample contained either **2** or **3** in the following concentrations: 0.005 mM, 0.01 mM, 0.05 mM, 0.1 mM, 0.5 mM, 1.0 mM. To each sample was then added 50 µL 0.01 mM chloramphenicol in methanol (1 eq.). Reactions were then centrifuged at ~ 21,000 *x g* for 5 min. The supernatant was then subjected to analysis by LCMS.

<u>DabC kinetics assays</u>

DabC kinetics assays for both **1-*trans*** and **1-*cis*** substrates were set up concurrently and in triplicate. Total reaction volumes were 200 µL and contained 50 mM KPi pH 6.0, 0.1 mM L-ascorbate, 50 mM α-ketoglutarate disodium salt, 1.0 mM $FeSO_4$, 1 µM DabC, and either **1-*trans*** and **1-*cis*** in each of the following concentrations: 25 µM, 50 µM, 100 µM, 175 µM, 300 µM, 400 µM, 500 µM. Four concentrations were assayed at a time in 500 µL PCR tubes (Eppendorf). At 5, 10, and 20 min timepoints, 60 µL of each reaction was quenched with 1 eq. of 0.01 mM chloramphenicol in methanol. Reactions were started and quenched via multichannel pipette to ensure consistent total reaction times for each sample. Replicates were staggered such that they were started and quenched 1 min apart from each other. All reactions were centrifuged in a benchtop microcentrifuge (USA Scientific) at 2000 *x g*. The supernatant was then subjected to analysis by UPLC-MS.

#### Supplementary Figures & Tables

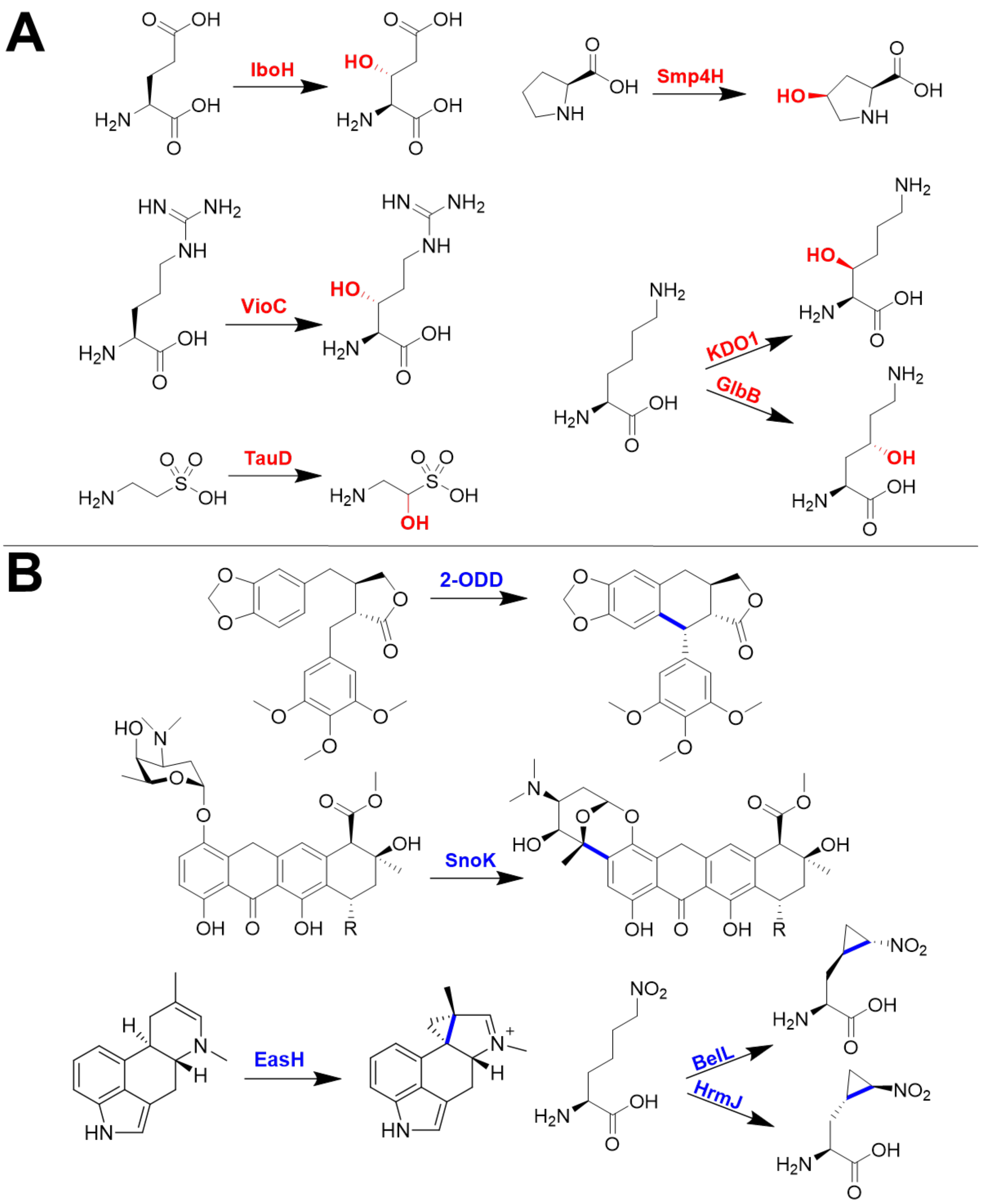


**Figure S1.** Notable examples of native hydroxylation (A.) and C-C cyclization (B.) reactions catalyzed by non-heme $Fe^{II}$/ $\alpha$KG-dependent dioxygenases. **A.** IboH [2], VioC [3], TauD [4], KDO1 [5], GlbB [6]. **B.** 2-ODD [7], SnoK [8], EasH [9], BelL[10], HrmJ [10].

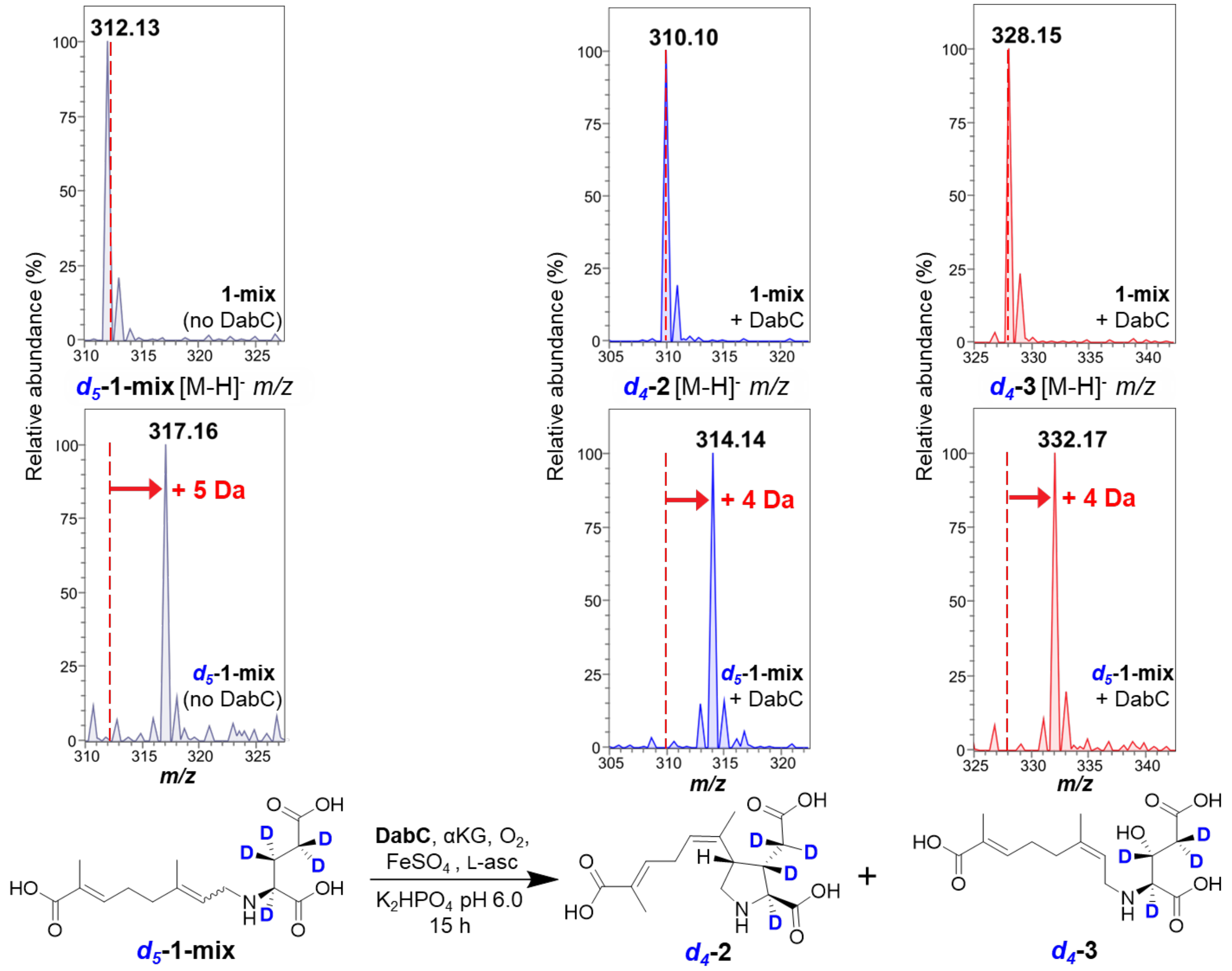


**Figure S2.** *In vitro* DabC assay with ***d*₅-1-mix** substrate. *d*5-labeling of DabC substrates shows radical abstraction of glutamic acid moiety for cyclization and hydroxylation. Analytical DabC assays were set up as previously described with ***d*₅-1-mix** the substrate. Relative intensities of negative mode extracted ion chromatograms were extracted from UPLC-MS traces. Results confirm that ***d*₅-1-mix** ([M-H]$^-$ 317.16 *m/z*) is converted to cyclized product ***d*₄-2** ([M-H]$^-$ 314.14 *m/z*) as well as β-hydroxylated product ***d*₄-3** ([M-H]$^-$ 332.17 *m/z*). This indicates a mass loss of one deuterium from both cyclized and hydroxylated products, supporting the mechanistic radical abstraction from the beta carbon of the L-glutamic acid moiety for both *cis* and *trans* substrates.

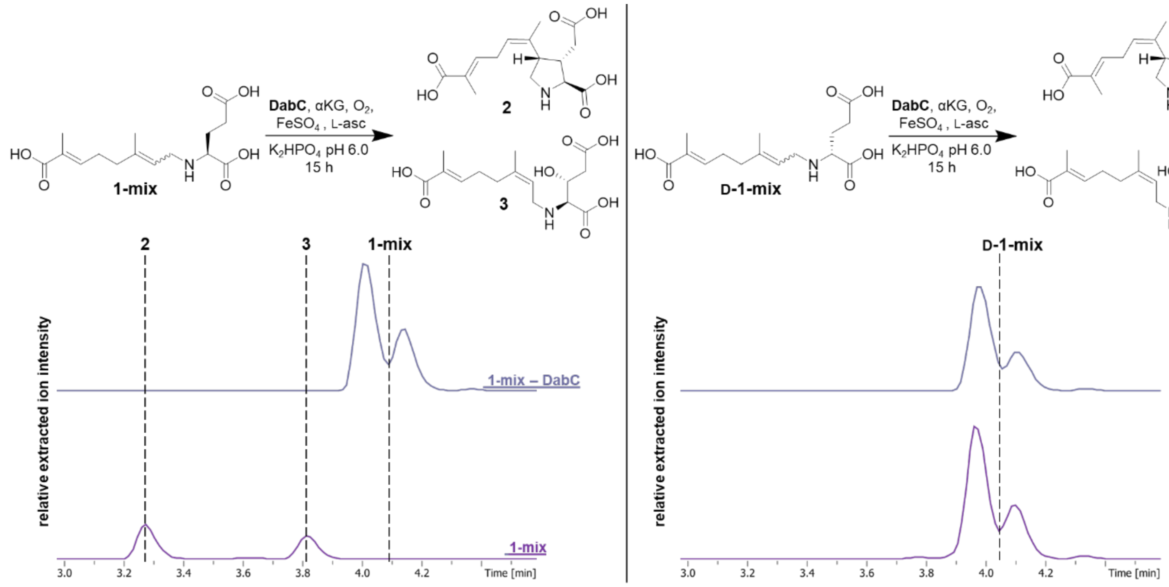


**Figure S3.** *In vitro* DabC assay with **D-1-mix** substrate. D-glutamic acid substrate analog shows stereospecificity for cyclization and hydroxylation. Analytical DabC assays were set up as previously described with either **1-mix** (left) or **D-1-mix** (right) as the substrate in the absence (top) or presence (bottom) of enzyme. Relative intensities of negative mode extracted ion chromatograms were extracted from UPLC-MS traces (312.15, 310.12, 328.14 ± 0.3 *m/z*). Results confirm that **1-mix** ($[M-H]^-$ 312.15 *m/z*) is converted to cyclized product **2** ($[M-H]^-$ 310.12 *m/z*) and β-hydroxylated product **3** ($[M-H]^-$ 328.14 *m/z*), whereas **D-1-mix** does not show definitive conversion to any cyclized or hydroxylated products.

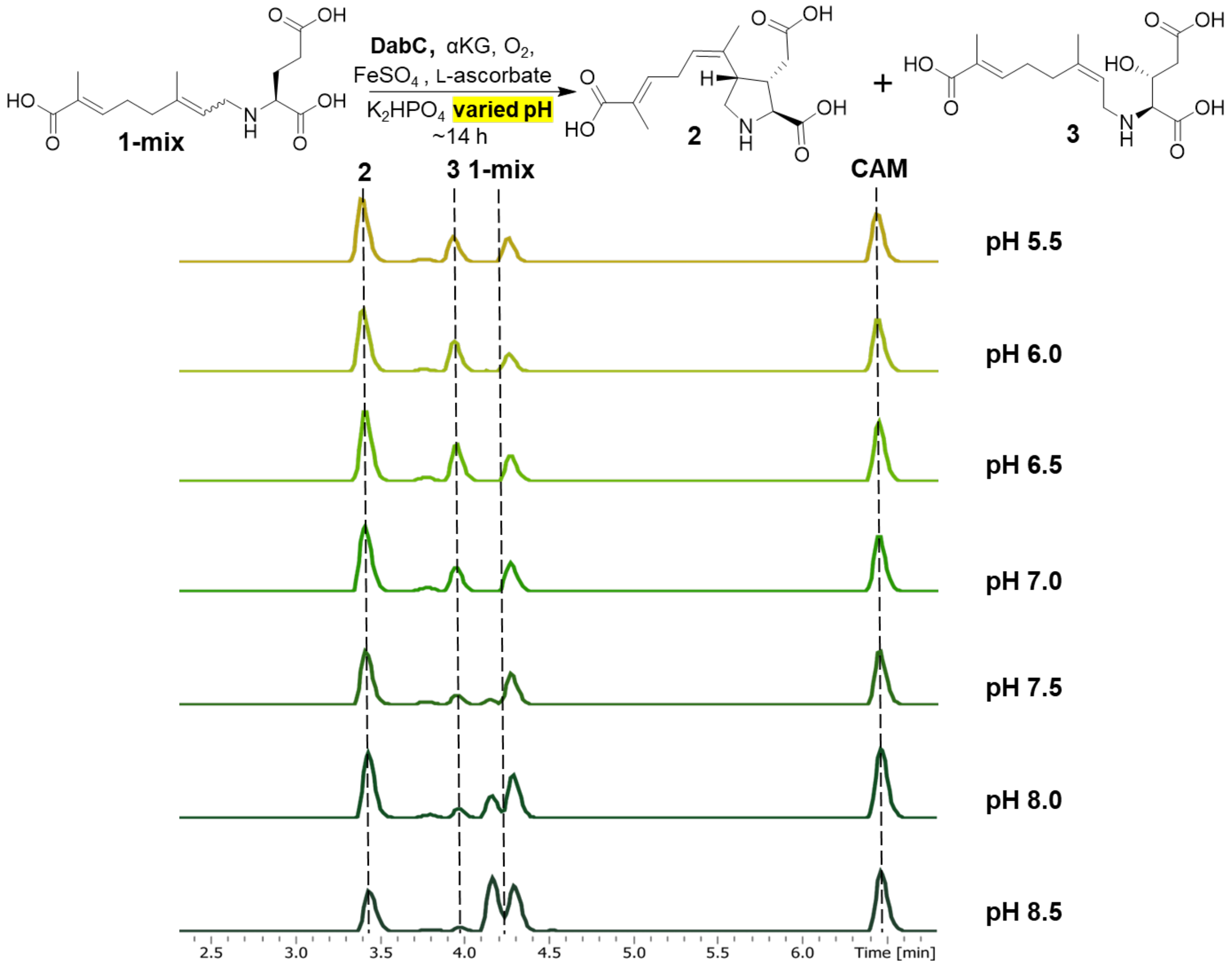


**Figure S4**. *In vitro* DabC pH optimization of **1-mix** cyclization and hydroxylation. Analytical optimization assays were carried out as described previously at a range of pHs from 5.5 – 8.5. Relative intensities of negative mode extracted ion chromatograms were extracted from UPLC-MS traces for substrate **1-mix**, products **2** and **3**, as well as the chloramphenicol standard (CAM) (312.15, 310.12, 328.14, 321.01 ± 0.3 *m/z* respectively). A pH of 6.0 was determined to be optimal and implemented for subsequent kinetics assays.

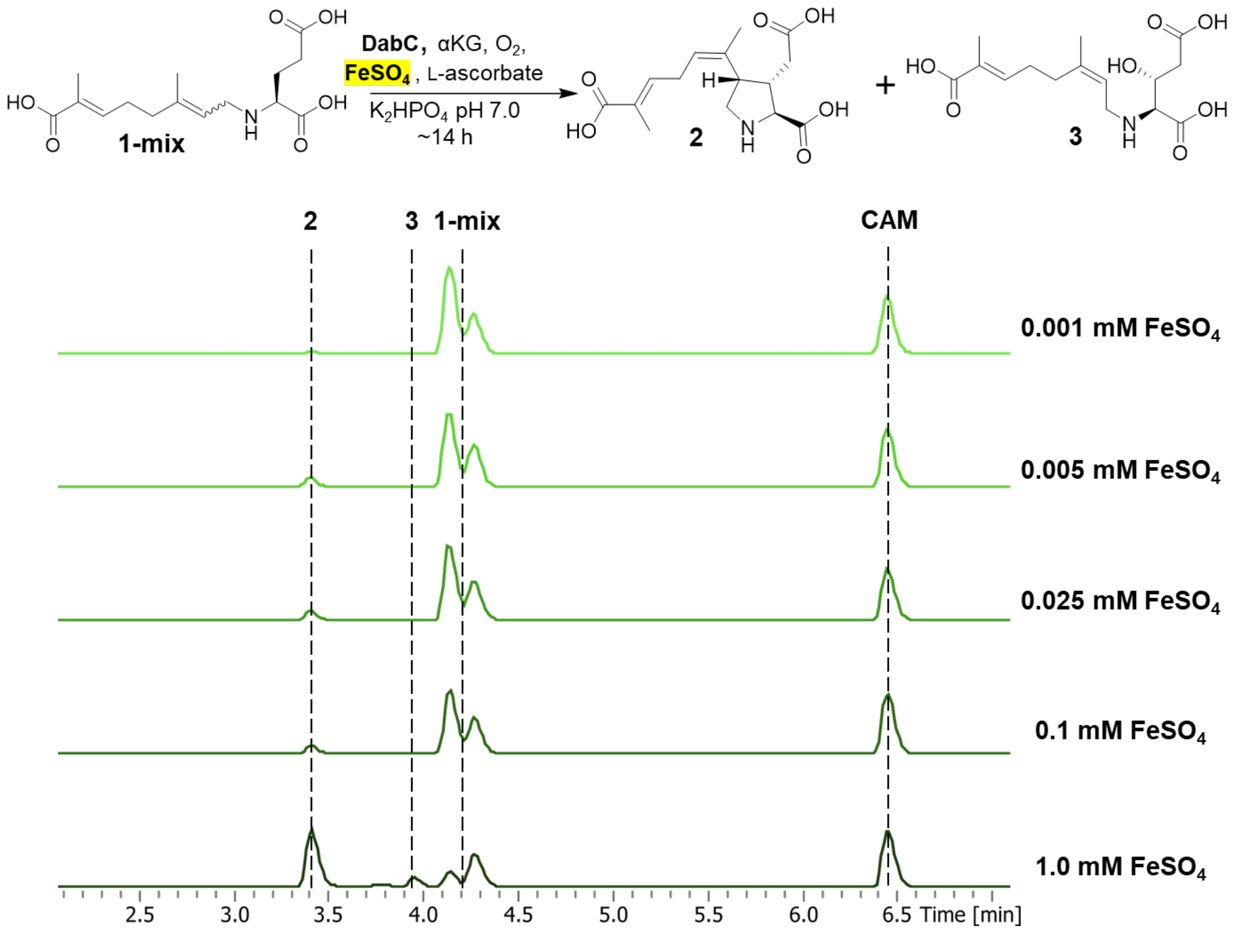


**Figure S5**. *In vitro* DabC $FeSO_4$ optimization of **1-mix** cyclization and hydroxylation. Analytical optimization assays were carried out as described previously over a range of $FeSO_4$ concentrations from 0.001 – 1.0 mM. Relative intensities of negative mode extracted ion chromatograms were extracted from UPLC-MS traces for substrate **1-mix**, products **2** and **3**, as well as the chloramphenicol standard (CAM) (312.15, 310.12, 328.14, 321.01 ± 0.3 *m/z* respectively). A concentration of 1.0 mM $FeSO_4$ was determined to be optimal and implemented for subsequent kinetics assays.

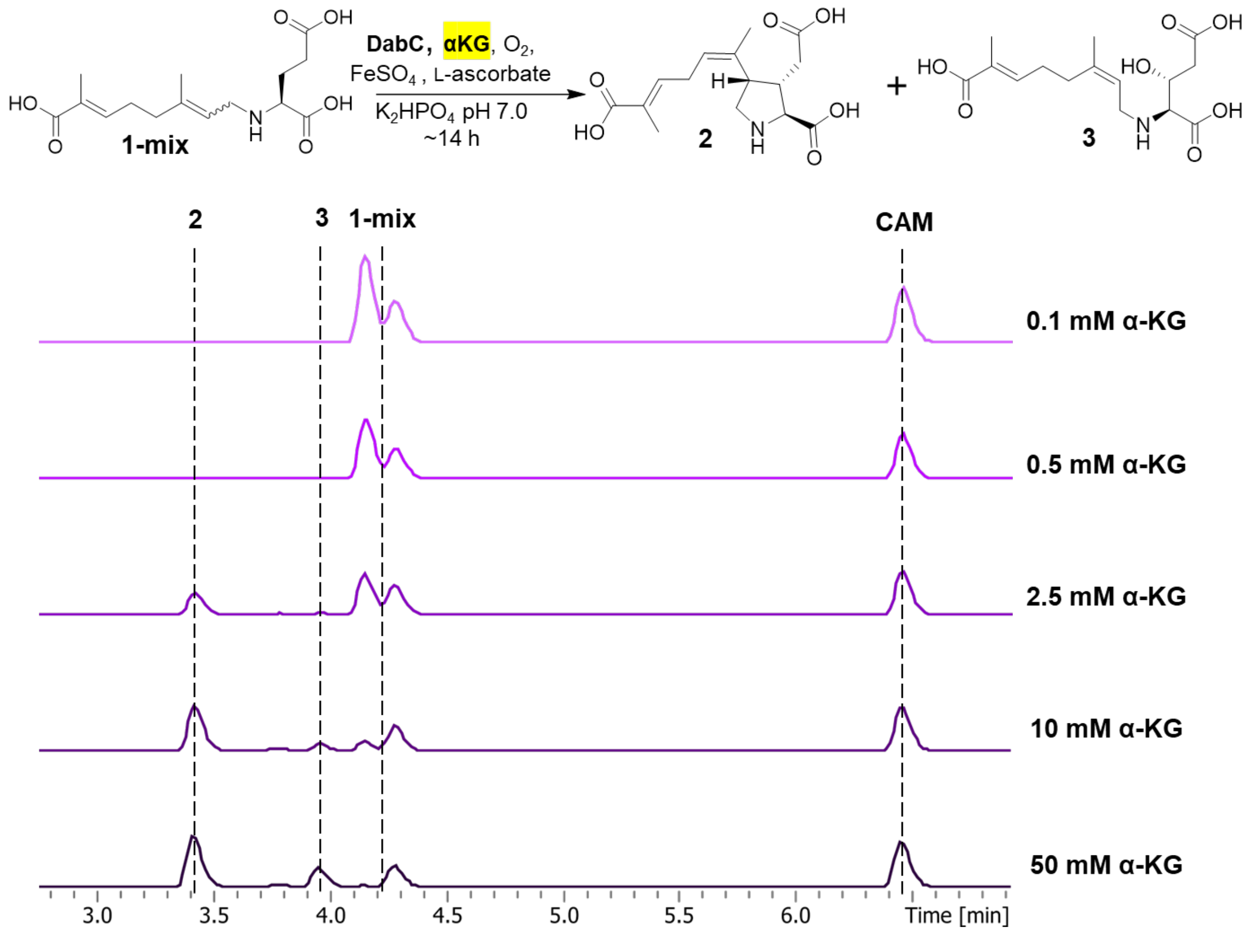


**Figure S6**. *In vitro* DabC αKG optimization of **1-mix** cyclization and hydroxylation. Analytical optimization assays were carried out as described previously over a range of αKG concentrations from 0.1 – 50 mM. Relative intensities of negative mode extracted ion chromatograms were extracted from UPLC-MS traces for substrate **1-mix**, products **2** and **3**, as well as the chloramphenicol standard (CAM) (312.15, 310.12, 328.14, 321.01 ± 0.3 *m/z* respectively). A concentration of 50 mM αKG was determined to be optimal and implemented for subsequent kinetics assays.

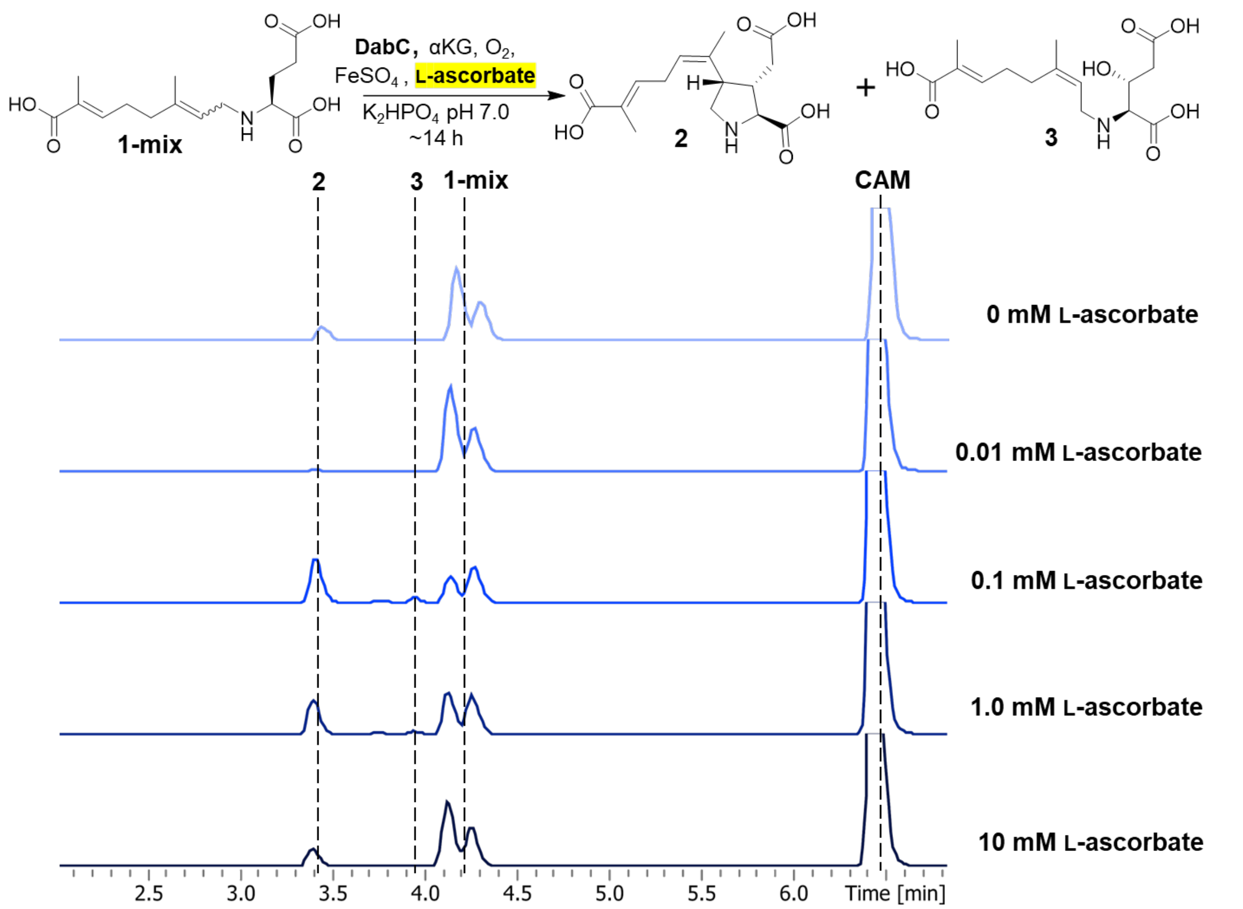


**Figure S7**. *In vitro* DabC L-ascorbate optimization of **1-mix** cyclization and hydroxylation. Analytical optimization assays were carried out as described previously over a range of L-ascorbate concentrations from 0 – 10 mM. Relative intensities of negative mode extracted ion chromatograms were extracted from UPLC-MS traces for substrate **1-mix**, products **2** and **3**, as well as the chloramphenicol standard (CAM) (312.15, 310.12, 328.14, 321.01 ± 0.3 *m/z* respectively). A concentration of 0.1 mM L-ascorbate was determined to be optimal and implemented for subsequent kinetics assays.

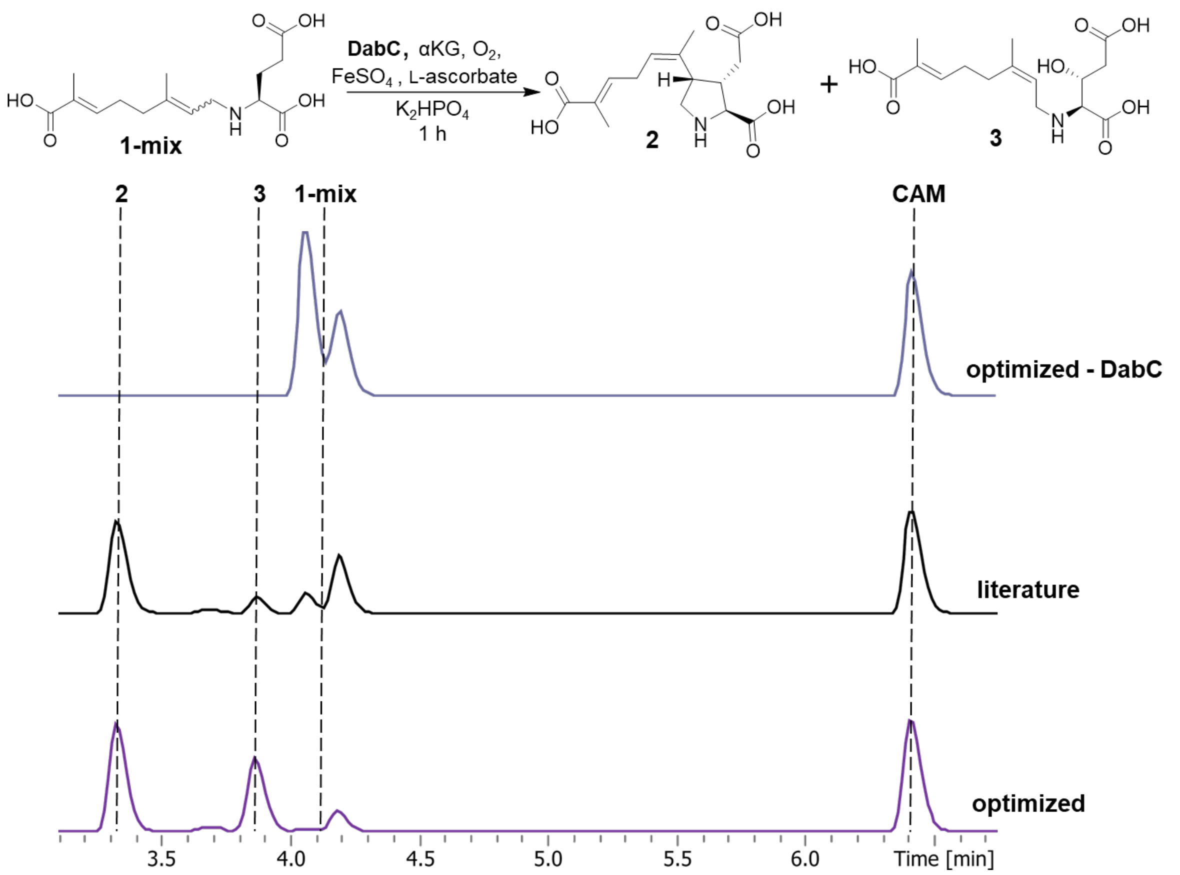


**Figure S8**. *In vitro* DabC assay of **1-mix** cyclization and hydroxylation pre- and post-optimization. Analytical optimization assays were carried out as described previously comparing the optimized assay conditions (KPi pH 6.0, 0.1 mM L-ascorbate, 50 mM α-ketoglutarate disodium salt, 1.0 mM $FeSO_4$) to the conditions reported in the literature (KPi pH 8.0, L-ascorbate, 50 mM α-ketoglutarate disodium salt, 1.0 mM $FeSO_4$). Relative intensities of negative mode extracted ion chromatograms were extracted from UPLC-MS traces for substrate **1-mix**, products **2** and **3**, as well as the chloramphenicol standard (CAM) (312.15, 310.12, 328.14, 321.01 ± 0.3 *m/z* respectively). The results indicate an increase in conversion of **1-mix** to products **2** and **3** in assays with optimized conditions relative to literature conditions.

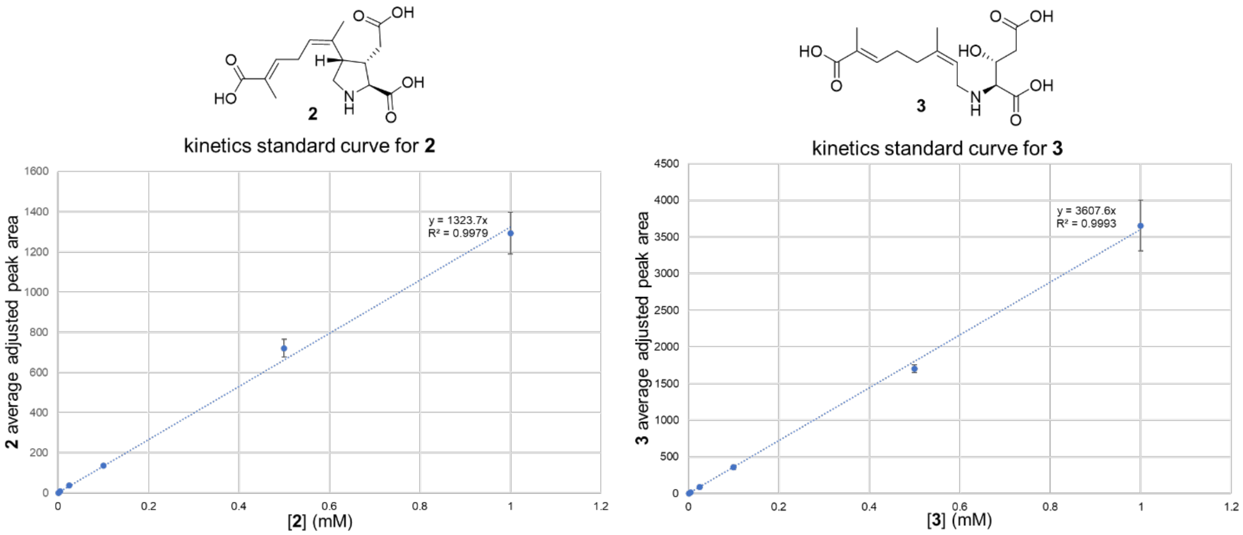


**Figure S9**. Standard curves of DabC products **2** and **3** from UPLC-MS analyses**.** Purified products were ran in triplicate at the following concentrations: 0.005 mM, 0.01 mM, 0.05 mM, 0.1 mM, 0.5 mM, 1.0 mM. Both product peak areas were individually adjusted using the peak area of a chloramphenicol internal standard and plotted to produce a standard curve for quantification of *in vitro* enzyme kinetic assays.

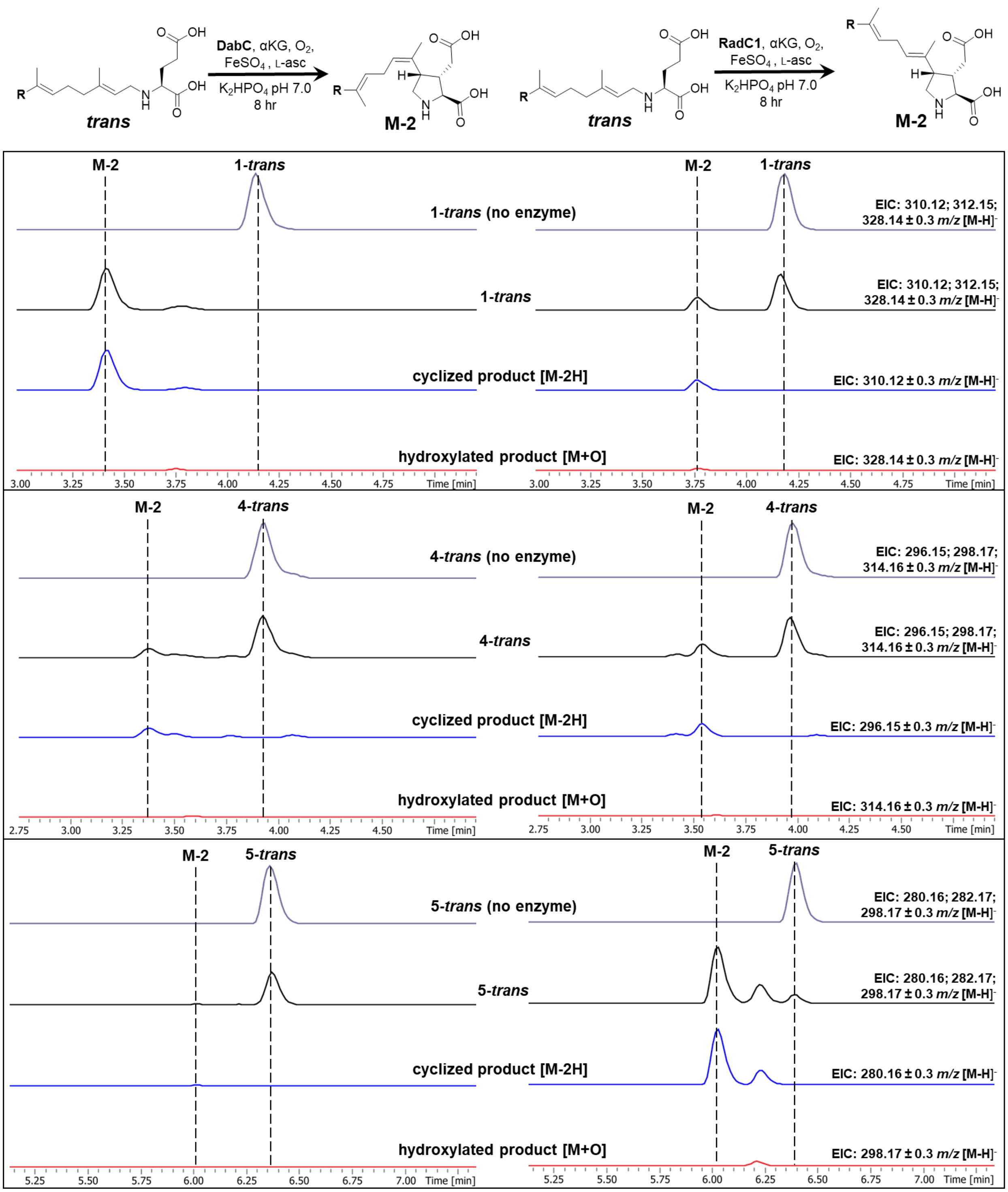
trans
DabC, αKG, O2, FeSO4, L-asc
K2HPO4 pH 7.0
8 hr
M-2
RadC1, αKG, O2, FeSO4, L-asc
K2HPO4 pH 7.0
8 hr
M-2
1-trans (no enzyme)
1-trans
cyclized product [M-2H]
hydroxylated product [M+O]
EIC: 310.12; 312.15; 328.14 ± 0.3 m/z [M-H]-
EIC: 310.12 ± 0.3 m/z [M-H]-
EIC: 328.14 ± 0.3 m/z [M-H]-
4-trans (no enzyme)
4-trans
EIC: 296.15; 298.17; 314.16 ± 0.3 m/z [M-H]-
EIC: 296.15 ± 0.3 m/z [M-H]-
EIC: 314.16 ± 0.3 m/z [M-H]-
5-trans (no enzyme)
5-trans
EIC: 280.16; 282.17; 298.17 ± 0.3 m/z [M-H]-
EIC: 280.16 ± 0.3 m/z [M-H]-
EIC: 298.17 ± 0.3 m/z [M-H]-
Time [min]

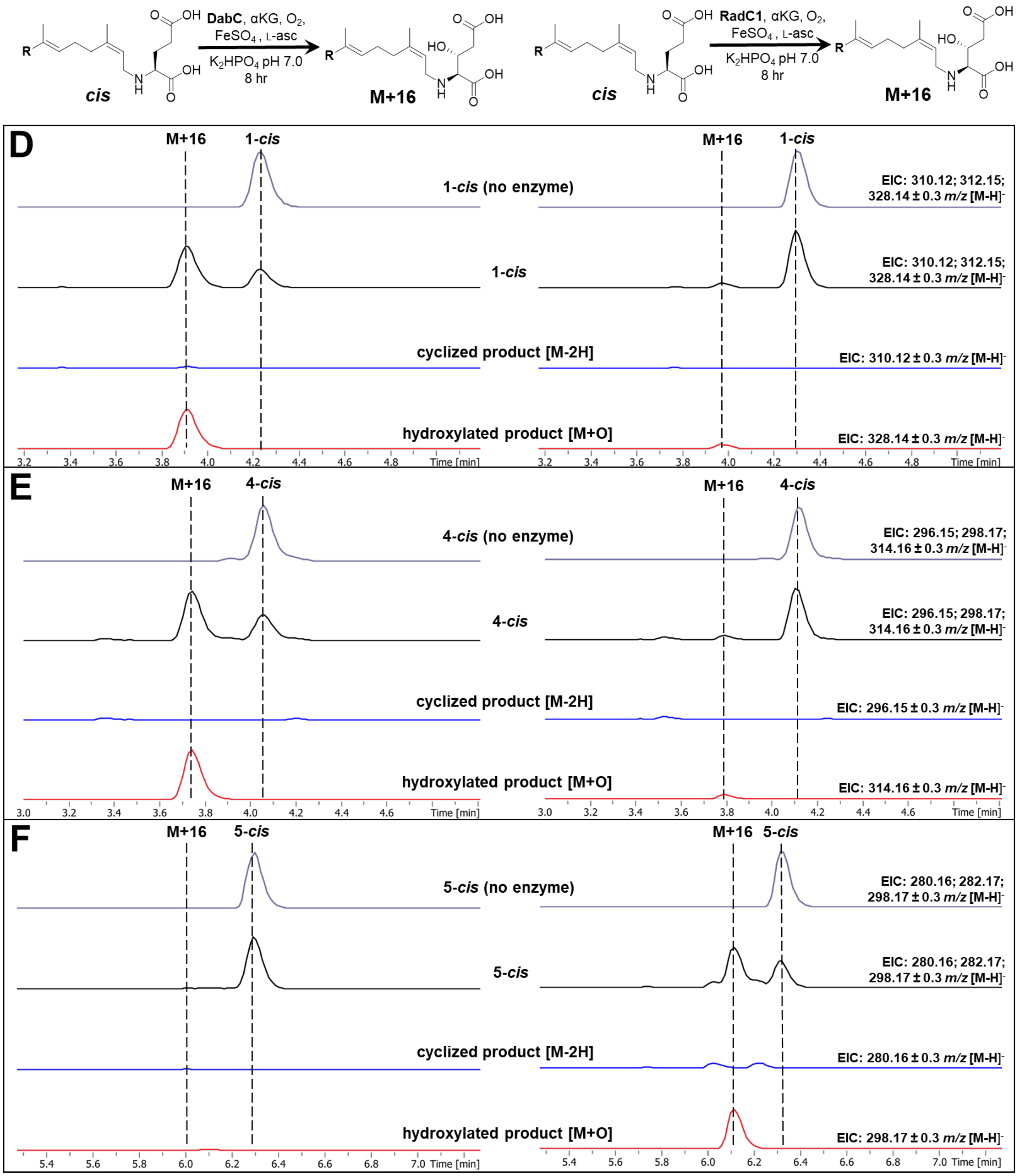

DabC, αKG, O2, FeSO4, L-asc
K2HPO4 pH 7.0
8 hr
cis
M+16
RadC1, αKG, O2, FeSO4, L-asc
K2HPO4 pH 7.0
8 hr
D
M+16
1-cis
1-cis (no enzyme)
1-cis
cyclized product [M-2H]
hydroxylated product [M+O]
EIC: 310.12; 312.15; 328.14 ± 0.3 m/z [M-H]-
EIC: 310.12 ± 0.3 m/z [M-H]-
EIC: 328.14 ± 0.3 m/z [M-H]-
Time [min]
E
M+16
4-cis
4-cis (no enzyme)
4-cis
cyclized product [M-2H]
hydroxylated product [M+O]
EIC: 296.15; 298.17; 314.16 ± 0.3 m/z [M-H]-
EIC: 296.15 ± 0.3 m/z [M-H]-
EIC: 314.16 ± 0.3 m/z [M-H]-
Time [min]
F
M+16
5-cis
5-cis (no enzyme)
5-cis
cyclized product [M-2H]
hydroxylated product [M+O]
EIC: 280.16; 282.17; 298.17 ± 0.3 m/z [M-H]-
EIC: 280.16 ± 0.3 m/z [M-H]-
EIC: 298.17 ± 0.3 m/z [M-H]-
Time [min]

**Figure S10**. DabC and RadC1 activity assays on 7’-modified *trans* and *cis* substrate analogs comparing cyclized and hydroxylated product mass EICs. Reactions were set up as previously described using *N*-prenylated glutamic acid analogs (1.0 mM) and incubated for 8 h at room temperature. Negative mode LCMS chromatograms of substrate specificities with various *N*-prenylated glutamic acid analogs are shown. Relative intensities were extracted for substrate and all potential product masses followed by cyclization and hydroxylation product masses individually. (**A**: **1-*trans*** substrate - EIC: 310.12; 312.15; 328.14 ± 0.3 *m/z* $[M-H]^-$; **B**: **4-*trans*** - EIC: 296.15; 298.17; 314.16 ± 0.3 *m/z* $[M-H]^-$ ; **C**: **5-*trans*** - EIC: 280.16; 282.17; 298.17 ± 0.3 *m/z* $[M-H]^-$ **D**: **1-*cis*** - EIC: 310.12; 312.15; 328.14 ± 0.3 *m/z* $[M-H]^-$; **E**: **4-*cis*** - EIC: 296.15; 298.17; 314.16 ± 0.3 *m/z* $[M-H]^-$ ; **F**: **5-*cis*** - EIC: 280.16; 282.17; 298.17 ± 0.3 *m/z* $[M-H]^-$).

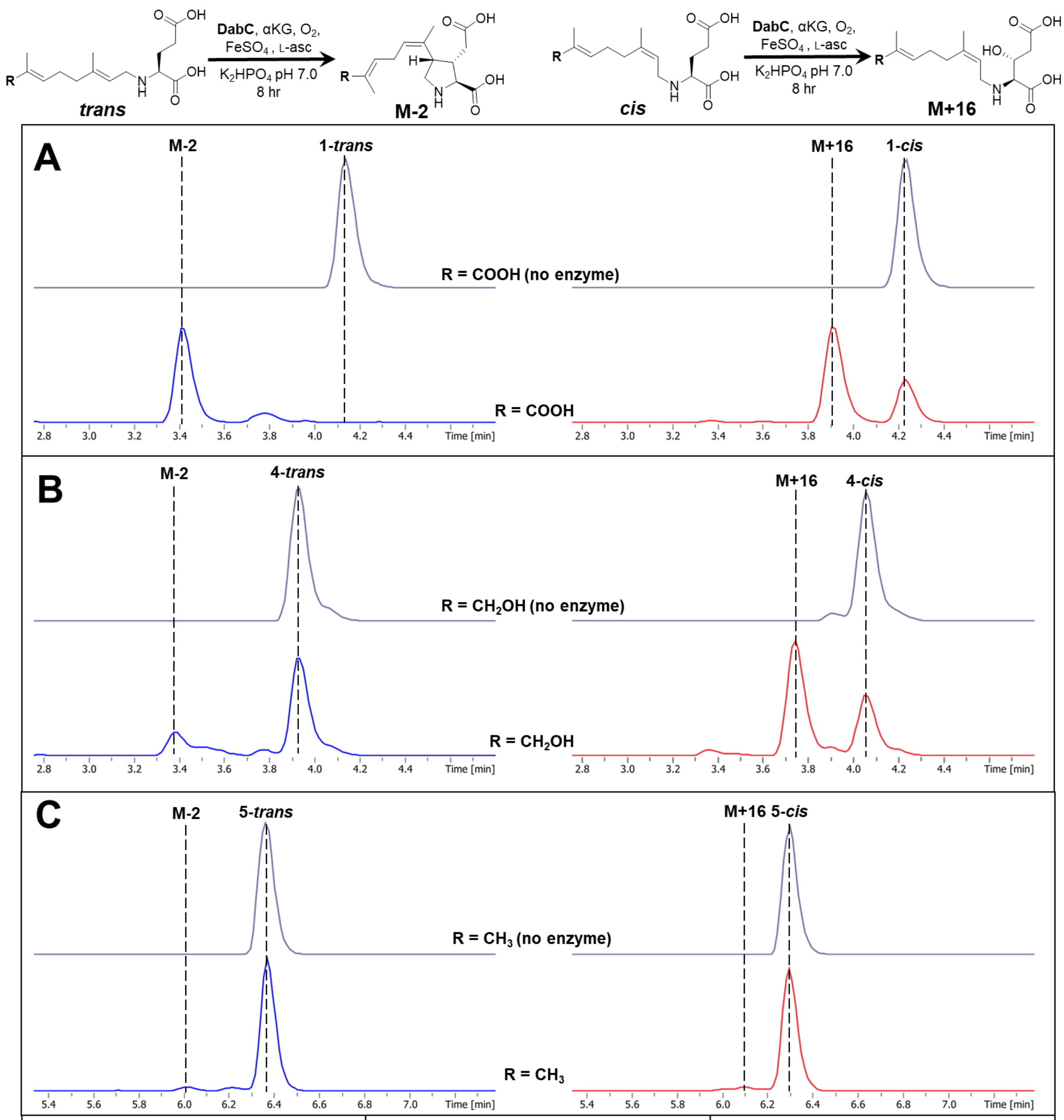


| % conversion to cyclized product | substrate R group | % conversion to hydroxylated product |
|---|---|---|
| **98** ± 2% | COOH | **68** ± 3% |
| **27** ± 2% | $CH_2OH$ | **62** ± 3% |
| **6** ± 0.4% | $CH_3$ | **4** ± 0.2% |

**Fig. S11.** DabC activity assay - *trans* vs *cis* substrate analogs. DabC reactions were set up as previously described in triplicate using *N*-prenylated glutamic acid analogs (1.0 mM) and incubated for 8 h at room temperature. Negative mode LCMS chromatograms of DabC substrate specificities with various *N*-prenylated glutamic acid analogs are shown. Relative intensities were extracted for substrate and all potential product masses (**A** - EIC: 310.12; 312.15; 328.14 ± 0.3 *m/z* $[M-H]^-$); **B** - EIC: 296.15; 298.17; 314.16 ± 0.3 *m/z* $[M-H]^-$ ; **C** - EIC: 280.16; 282.17; 298.17 ± 0.3 *m/z* $[M-H]^-$)

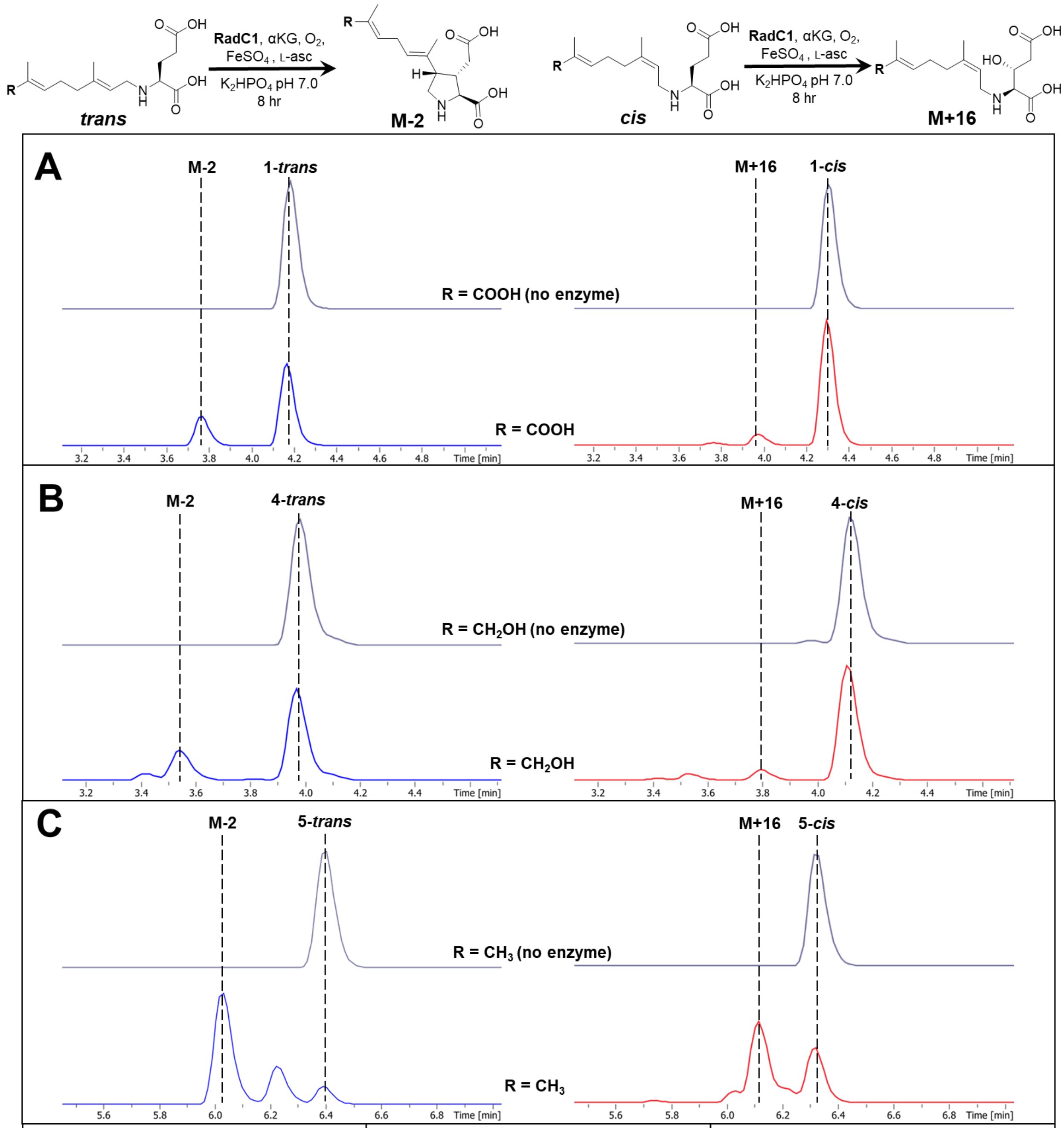


| % conversion to cyclized product | substrate R group | % conversion to hydroxylated product |
|---|---|---|
| **21** ± 3% | COOH | **9** ± 0.8% |
| **28** ± 2% | $CH_2OH$ | **8** ± 1.8% |
| **94** ± 3% | $CH_3$ | **47** ± 9% |

**Fig. S12.** RadC1 activity assay - *trans* vs *cis* substrate analogs. RadC1 reactions were set up as previously described in triplicate using *N*-prenylated glutamic acid analogs (1.0 mM) and incubated for 8 h at room temperature. Negative mode LCMS chromatograms of RadC1 substrate specificities with various *N*-prenylated glutamic acid analogs are shown. Relative intensities were extracted for substrate and all potential product masses (**A** - EIC: 310.12; 312.15; 328.14 ± 0.3 *m/z* $[M-H]^-$); **B** - EIC: 296.15; 298.17; 314.16 ± 0.3 *m/z* $[M-H]^-$ ; **C** - EIC: 280.16; 282.17; 298.17 ± 0.3 *m/z* $[M-H]^-$)

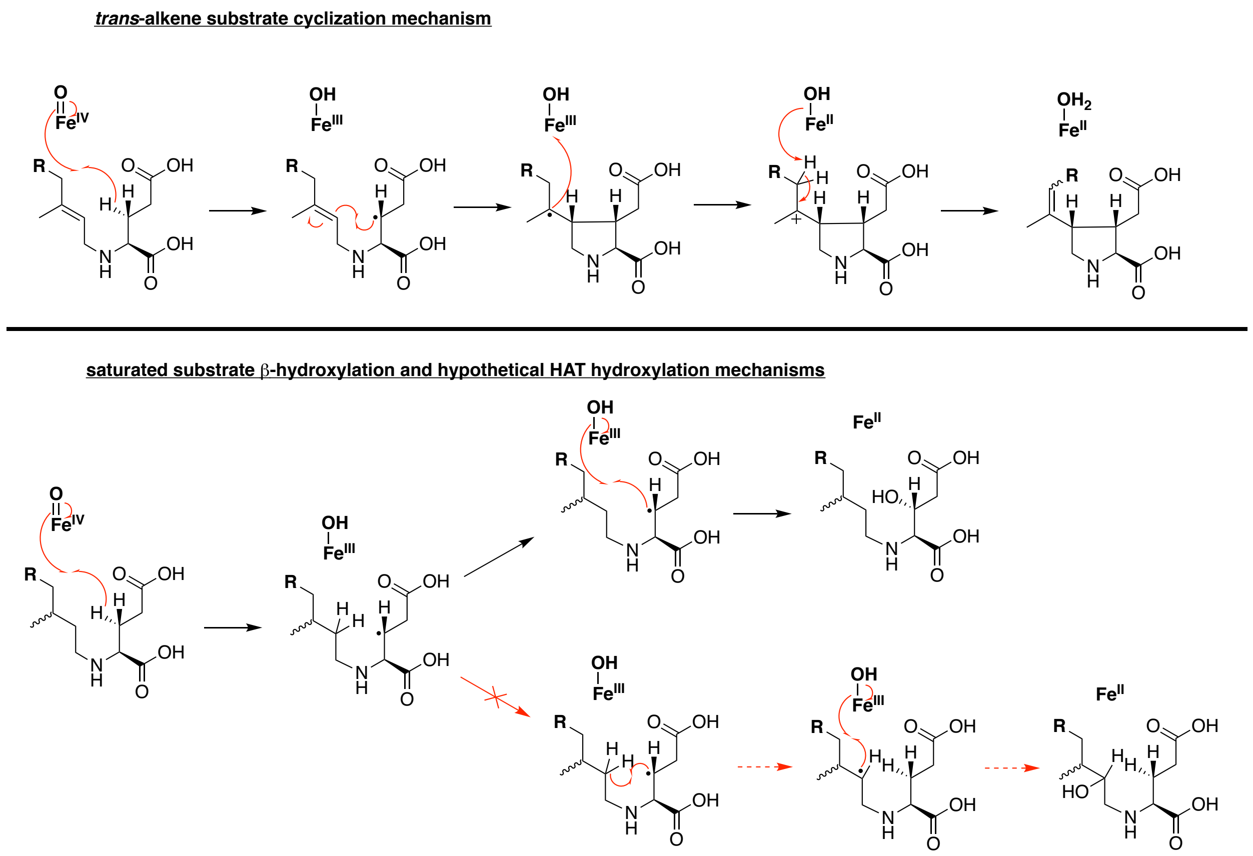


**Figure S13.** Alternative HAT mechanistic proposal for saturated substrate hydroxylation. Cyclization mechanism for the native *trans*-substrates by DabC and homologs (top) compared to the with the direct β-hydroxylation (middle). The theoretical 1,5-HAT transfer mechanism prior to hydroxyl rebound (bottom) does not occur experimentally with citronellyl substrate analogs within our limits of detection.

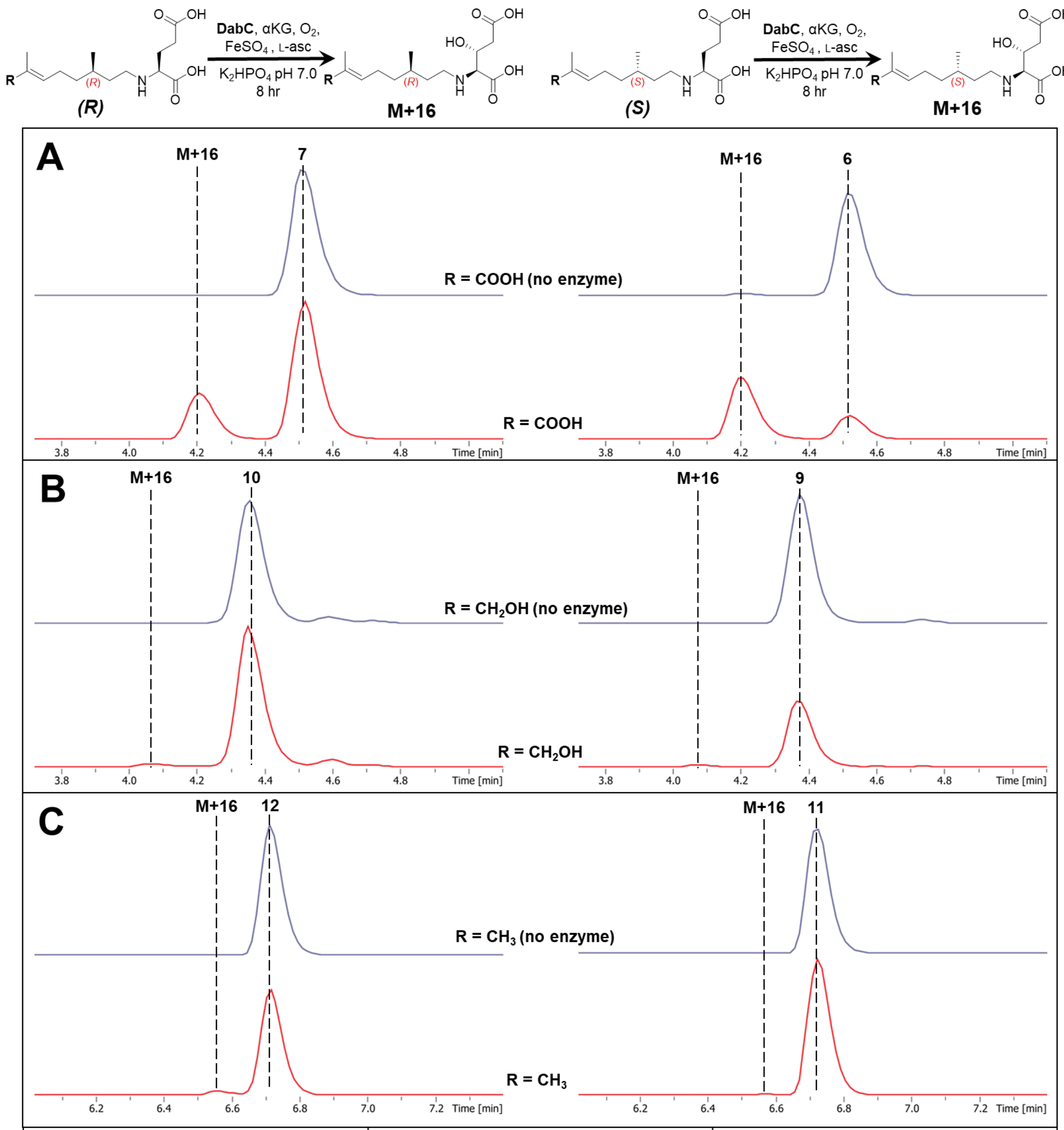


| % conversion to hydroxylated product | substrate R group | % conversion to hydroxylated product |
|---|---|---|
| **23** ± 2% | COOH | **73** ± 1% |
| **3** ± 0.1% | $CH_2OH$ | **4** ± 0.2% |
| **4** ± 0.1% | $CH_3$ | trace |

**Fig. S14.** DabC activity assay - *R* vs *S* citronellyl substrate analogs. DabC reactions were set up as previously described in triplicate using *N*-prenylated citronellyl glutamic acid analogs (1.0 mM) and incubated for 8 h at room temperature. Negative mode LCMS chromatograms of DabC substrate specificities with various *N*-prenylated citronellyl glutamic acid analogs are shown. Relative intensities were extracted for substrate and all potential product masses (**A** - EIC: 312.15; 314.16; 330.16 ± 0.3 *m/z* $[M-H]^-$); **B** - EIC: 298.17; 300.18; 316.18 ± 0.3 *m/z* $[M-H]^-$ ; **C** - EIC: 282.17; 284.19; 300.18 ± 0.3 *m/z* $[M-H]^-$).

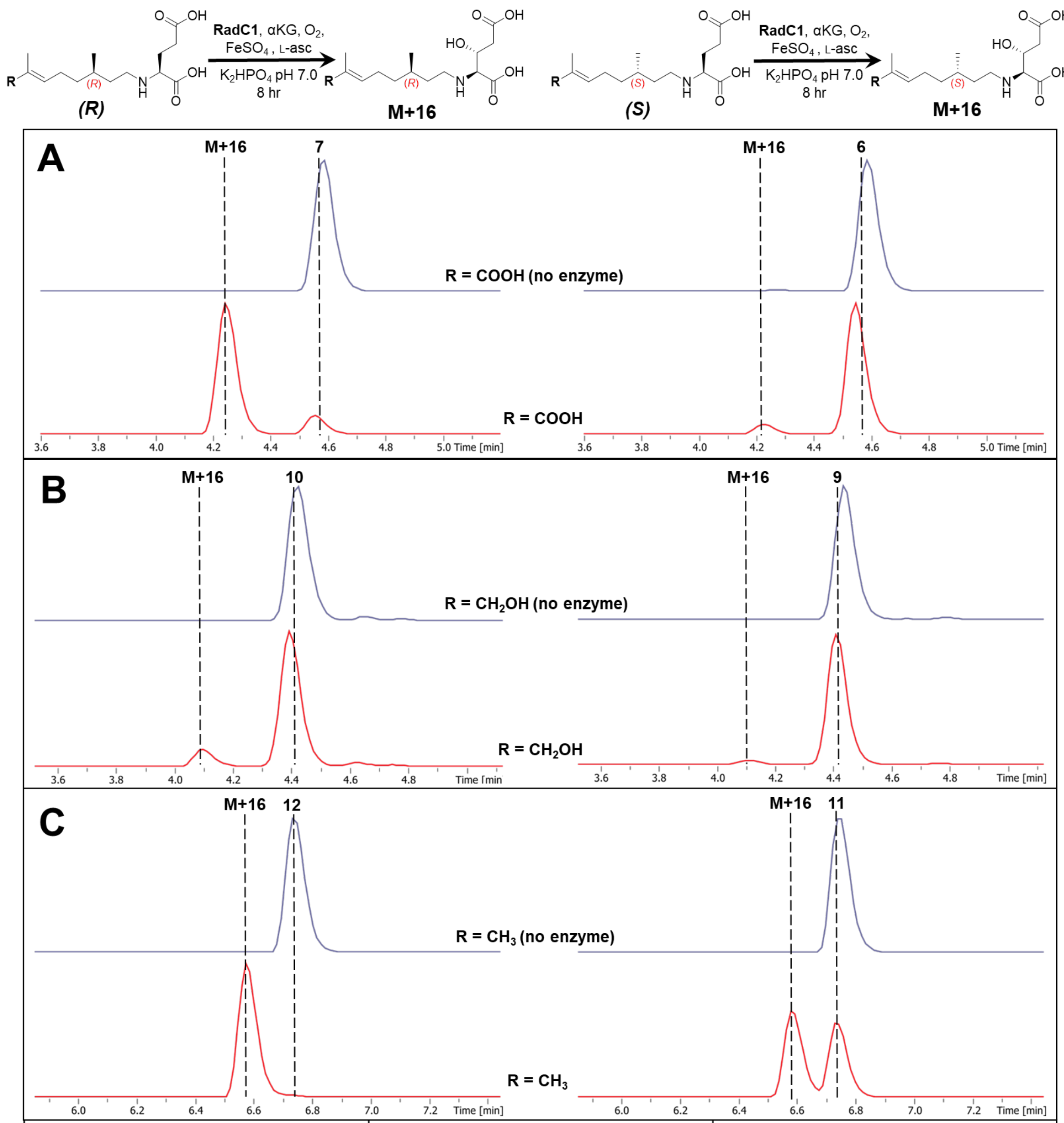


| % conversion to hydroxylated product | substrate R group | % conversion to hydroxylated product |
|---|---|---|
| **89** ± 4% | COOH | **8** ± 0.8% |
| **14** ± 3% | $CH_2OH$ | **4** ± 0.2% |
| **98** ± 1% | $CH_3$ | **54** ± 2% |

**Fig. S15.** RadC1 activity assay - *R* vs *S* citronellyl substrate analogs. RadC1 reactions were set up as previously described in triplicate using *N*-prenylated citronellyl glutamic acid analogs (1.0 mM) and incubated for 8 h at room temperature. Negative mode LCMS chromatograms of RadC1 substrate specificities with various *N*-prenylated citronellyl glutamic acid analogs are shown. Relative intensities were extracted for substrate and all potential product masses (**A** - EIC: 312.15; 314.16; 330.16 ± 0.3 *m/z* $[M-H]^-$); **B** - EIC: 298.17; 300.18; 316.18 ± 0.3 *m/z* $[M-H]^-$ ; **C** - EIC: 282.17; 284.19; 300.18 ± 0.3 *m/z* $[M-H]^-$).

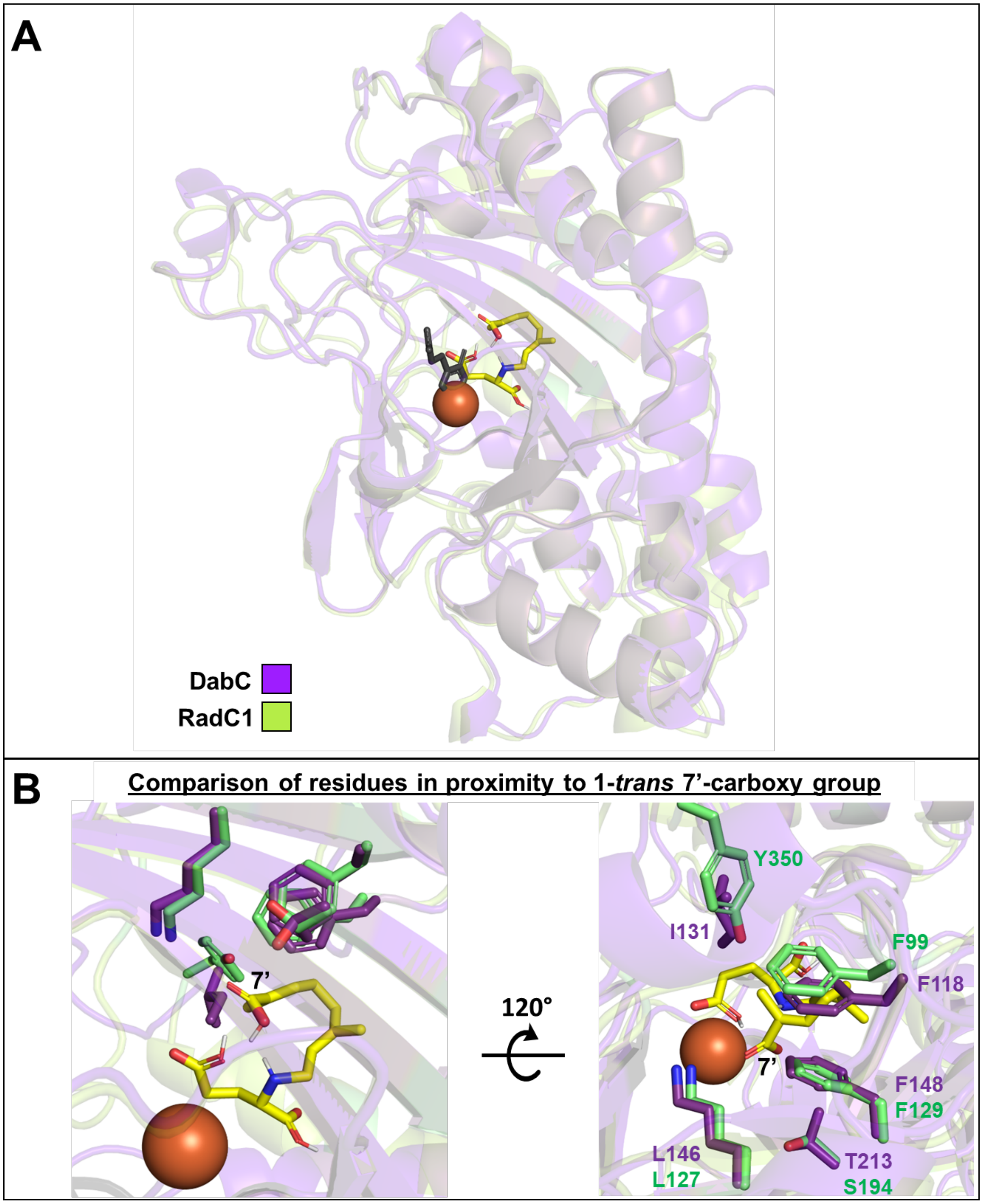
A
DabC
RadC1
B
Comparison of residues in proximity to 1-*trans* 7'-carboxy group
7'
120°
Y350
I131
F99
F118
7'
F148
F129
L146
L127
T213
S194

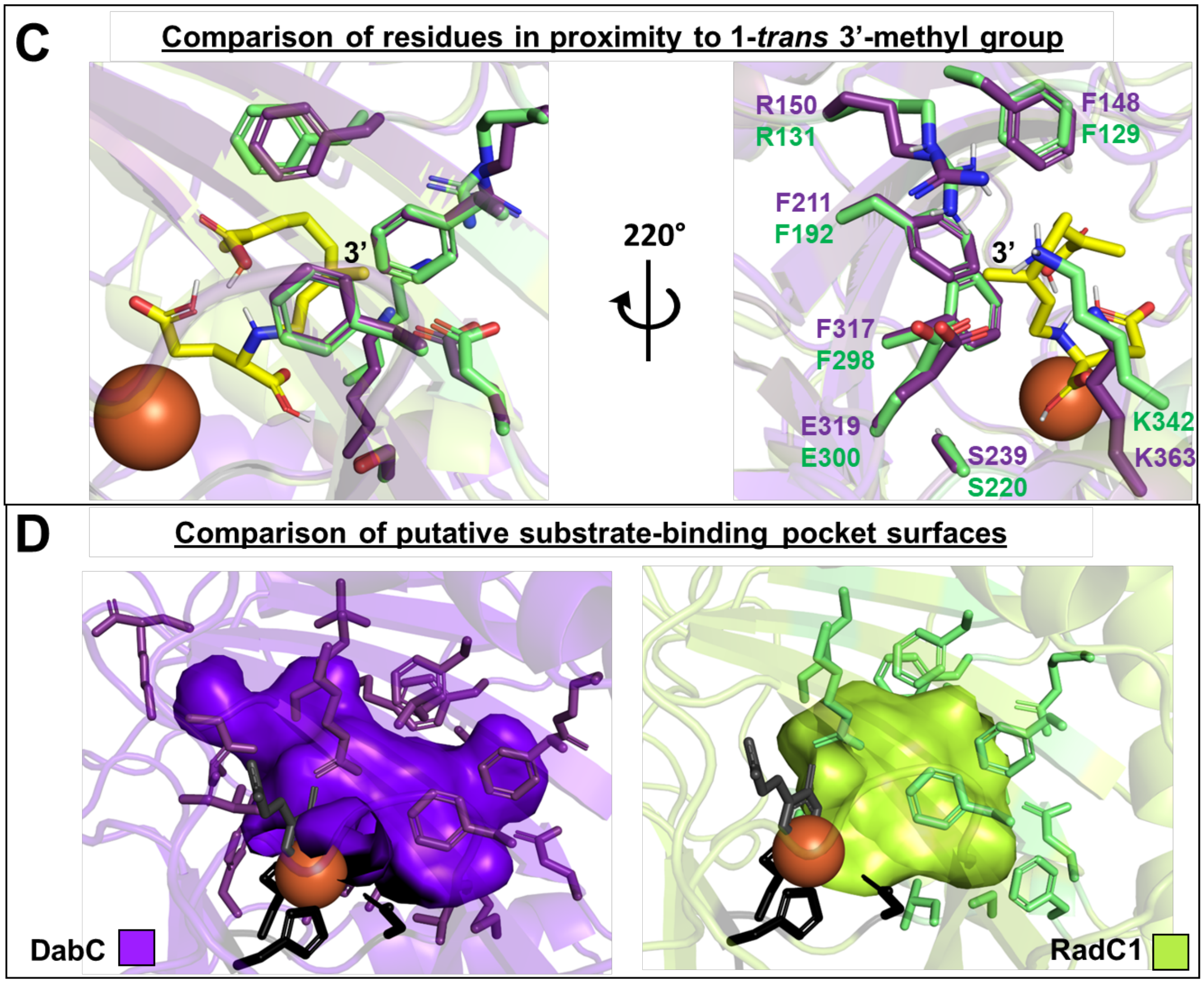


**Figure S16.** DabC/RadC1 AlphaFold model comparison and **1-*trans*** substrate docking. **A**. Overlay of DabC (purple) and RadC1 (green) structural models generated with AlphaFold, with docked **1-*trans*** substrate (yellow) displayed along with iron co-factor (orange) and $\alpha$-ketoglutarate co-substrate (grey). **B.** Overlay of DabC and RadC1 with indicated residues in 5Å proximity to the putative 7' carboxy group of **1-*trans.*** **C.** Overlay of DabC and RadC1 with indicated residues in 5Å proximity to the putative 3' methyl group of **1-*trans.*** **D.** Visualization of putative substrate-binding pocket surfaces in DabC (left) and RadC1 (right) with conserved facial triad residues shown (black). All images were generated using PyMol, enzyme models were generated using AlphaFold, substrate model of **1-*trans*** was generated with Avogadro, substrate docking was performed with AutoDock.

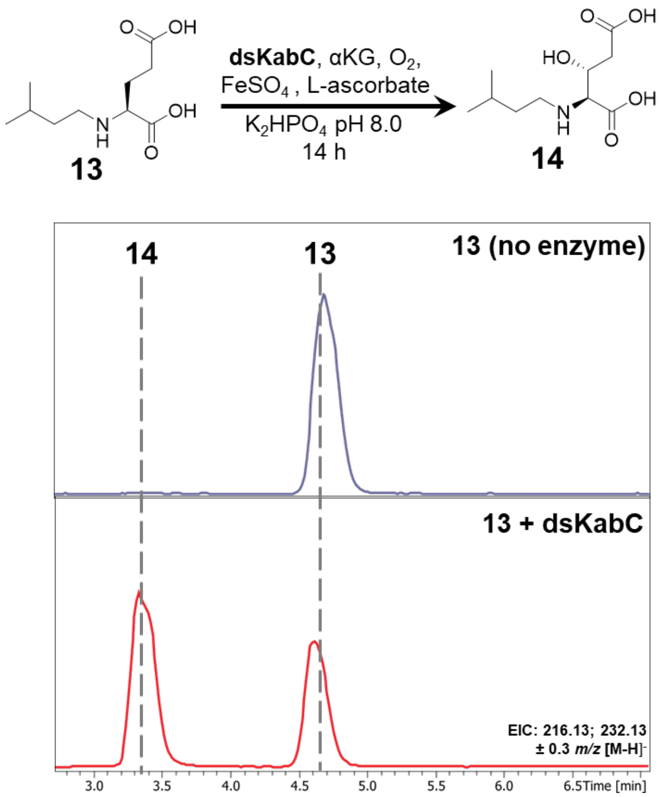


**Fig. S17.** *In vitro* DsKabC hydroxylation of saturated substrate isopentyl-*N*-L-glutamic acid (**13**). dsKabC reactions were set up in the same manner as was previously described for DabC small scale enzyme assays using **13** (1.0 mM) as the substrate and incubated for 14 h at room temperature. Negative mode LCMS chromatograms of reactions without and with dsKabC enzyme are shown. Relative intensities were extracted for substrate and hydroxylated product **14** masses (EIC: 216.13; 232.13 ± 0.3 *m/z* $[M\text{-}H]^-$)

Chemical Synthesis

Synthesis of 7'-carboxy-L-NGG (**1-*trans***) and 7'-carboxy- L-NNG (**1-*cis***)

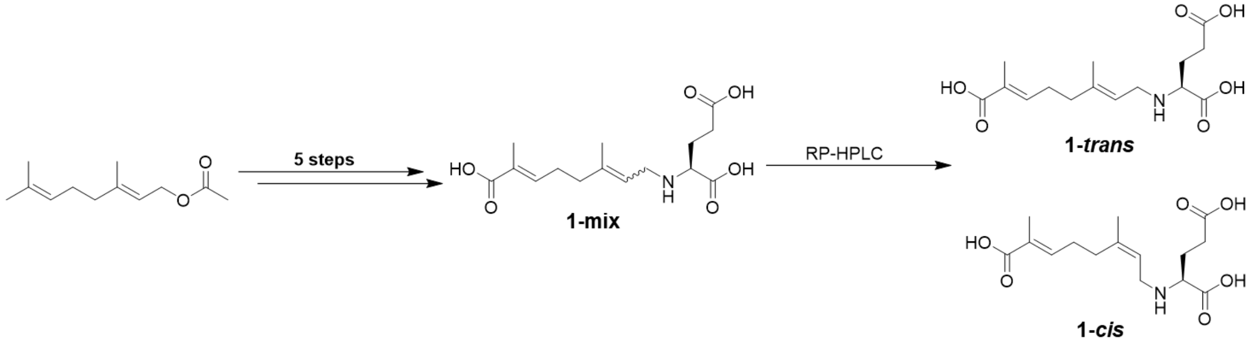


Compound **1** was synthesized as previously reported[11] to afford a white solid (0.073 g, 0.23 mmol, 2% over five steps) as a mixture of regioisomers (**1-mix**). Regioisomers were separated via preparative RP-HPLC (Phenomenex Kinetex 5 μm EVO C18 100 Å 250 x 21.2 mm) at a flow rate of 8 mL/min using 11.5% acetonitrile in 0.1% aqueous formic acid. The compounds were manually collected with the major **1-*trans*** isomer eluting at 22.0 min and the minor **1-*cis*** isomer eluting at 23.8 min.

7'-carboxy- L-NGG (**1-*trans***)

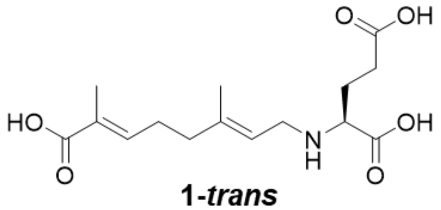

**1-*trans***

$^1$H NMR (500 MHz, $D_2O$ + 0.01% MeOH): δ 6.72 (t, *J* = 7.2 Hz, 1H), 5.24 (t, *J* = 7.9 Hz, 2H), 3.83 (dd, *J* = 7.9, 5.3 Hz, 1H), 3.73 (d, *J* = 7.7 Hz, 2H), 2.55 (q, *J* = 7.2 Hz, 2H), 2.44 – 2.33 (m, 2H), 2.29 – 2.08 (m, 4H), 1.77 (s, 3H), 1.69 (s, 3H) ; $^{13}$C NMR (125 MHz, $D_2O$ + 0.01% MeOH): 176.9, 172.8, 172.1, 147.8, 144.6, 128.3, 113.9, 58.6, 44.5, 38.1, 30.1, 26.7, 25.0, 16.2, 12.3; HRMS (ESI) Calculated for $C_{15}H_{24}NO_6$ 314.1604, found 314.1610 $(M+H)^+$

7'-carboxy -L-NNG (**1-*cis***)

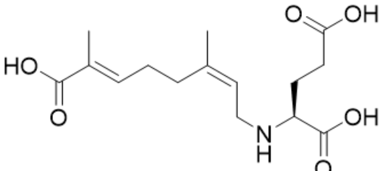

**1-*cis***

$^{1}H$ NMR (500 MHz, $D_2O$ + 0.1% MeOH): 6.68 (t, *J* = 7.6 Hz, 1H), 5.28 (t, *J* = 8.1 Hz, 1H), 3.67 - 3.59 (m, 2H), 3.57 (t, *J* = 6.4 Hz, 1H), 2.49 – 2.38 (m, 2H), 2.38 – 2.30 (m, 2H), 2.27 – 2.21 (m, 2H), 2.13 – 1.98 (m, 2H), 1.78 (s, 3H), 1.76 (s, 3H); $^{13}C$ NMR (125 MHz, $D_2O$) δ 178.2, 173.1, 172.8, 146.2, 142.7, 128.8, 114.5, 60.8, 44.1, 30.1, 26.5, 25.5, 22.7, 20.5, 11.9; HRMS (ESI) Calculated for $C_{15}H_{24}NO_6$ 314.1604, found 314.1608 $(M+H)^+$

Characterization of **1-mix**

$^{1}H$ NMR (500 MHz, $D_2O$ + 0.1% MeOD): δ 6.85 – 6.74 (m, 1H), 5.28 (t, *J* = 8.0 Hz, 1H), 3.82 – 3.67 (m, 3H), 2.63 – 2.51 (m, 2H), 2.45 – 2.36 (m, 2H), 2.31 – 2.25 (m, 2H), 2.25 – 2.08 (m, 2H), 1.81 (m, 3H), 1.73 (s, 3H). Calculated for $C_{15}H_{22}NO_6$ 312.14, found 312.14 $(M-H)^-$.

Synthesis of **D-1-mix** and ***d*5-1-mix**

Compounds **D-1-mix** and ***d*5-1-mix** were synthesized via modification of the literature procedure used for the synthesis of compound **1** starting from 0.025 g of 7-carboxygeranial and substituting either D-glutamic acid or $d_5$-L-glutamic acid in place of L-glutamic acid.[11] The crude reaction mixtures were subsequently purified via a Biotage Isolera Prime flash chromatography system using a Sfar Ultra C18 D 6g pre-packed column at a flow rate of 6 mL/min with the following method: 10% B (2.5 column volumes (CV)), 10 - 30% B (9 CV), 30 - 45% B (2 CV), 45 - 95% B (1 CV), 95% B (3 CV), 95 - 10% B (1 CV), 10% B (2 CV), where A = 0.1 % aqueous formic acid, and B = 0.1% formic acid in acetonitrile.

**D-1-mix**

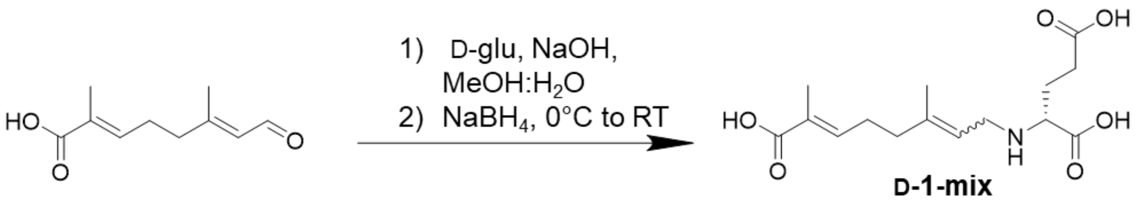


**D-1-mix**

Compound **D-1-mix** was isolated as a white solid (0.003 g, 0.01 mmol, 7%) as a mixture of regioisomers and characterization was consistent with that of compound **1**. $^1$H NMR (500 MHz, $D_2O$ + 0.1% MeOD): δ 6.85 – 6.74 (m, 1H), 5.34 – 5.27 (m, 1H), 3.75 – 3.61 (m, 3H), 2.56 – 2.50 (m, 2H), 2.44 – 2.37 (m, 2H), 2.34 – 2.25 (m, 2H), 2.21 – 2.03 (m, 2H), 1.83 – 1.80 (m, 3H), 1.73 (s, 3H). MS (ESI) Calculated for $C_{15}H_{22}NO_6$ 312.14, found 312.12 $(M-H)^-$.

***d*<sub></sub>**

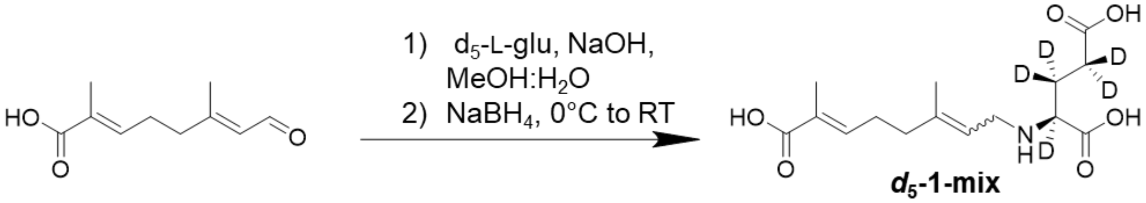


Compound ***d*$_5$-1-mix** was isolated as a white solid (0.0123 g, 0.04 mmol, 28%) as a mixture of regioisomers and characterization was consistent with that of compound **1** but with the expected absence of *d*$_5$-glutamic acid signals. $^1$H NMR (500 MHz, $D_2O$ + 0.1% MeOD): δ 6.82 (m, 1H), 5.29 (m, 1H), 3.73 – 3.65 (m, 2H), 2.45 – 2.36 (m, 2H), 2.34 – 2.25 (m, 2H), 1.84 – 1.80 (m, 3H), 1.73 (s, 3H). MS (ESI) Calculated for $C_{15}H_{17}D_5NO_6$ 317.18, found 317.16 $(M-H)^-$.

Synthesis of 7'-hydroxy -L-NGG (**4-*trans***) and 7'-hydroxy- L-NNG (**4-*cis***)

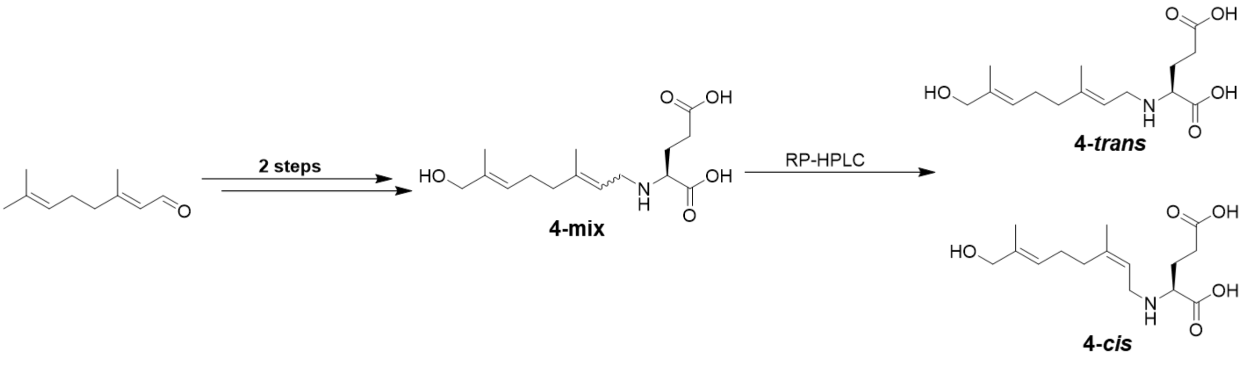


Compound **4** was synthesized as previously reported [11] to afford a white solid (0.066 g, 0.22 mmol, 14% over 2 steps) as a mixture of regioisomers. Isomers were separated via preparative RP-HPLC (Phenomenex Kinetex 5 µm EVO C18 100 Å 250 x 21.2 mm) at a flow rate of 8 mL/min using 9.5% acetonitrile in 0.1% aqueous formic acid. The compounds were manually collected with the *trans* isomer eluting at 22.5 min and the *cis* isomer eluting at 25.0 min.

7'-hydroxy- L-NGG (**4-*trans***)

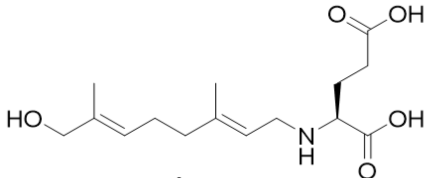

**4-*trans***

$^{1}$H NMR (500 MHz, $D_2O$): δ 5.44 (t, *J* = 7.0 Hz, 1H), 5.30 (t, *J* = 7.9 Hz, 1H), 3.99 (s, 2H), 3.72 (m, 2H), 3.65 (t, *J* = 6.3 Hz, 1H), 2.54 (m, 2H), 2.16 (t, *J* = 6.9 Hz, 2H), 2.06 – 2.27 (m, 6H), 1.75 (s, 3H), 1.68 (s, 3H); $^{13}$C NMR (125 MHz, $D_2O$): δ 177.4, 173.0, 147.4, 134.8, 126.1, 113.0, 67.5, 60.0, 44.0, 38.4, 30.7, 25.1, 25.0, 15.7, 13.0; HRMS (ESI) Calculated for $C_{15}H_{26}NO_5$ 300.1805, found 300.1808 $(M+H)^+$

7'-hydroxy-L-NNG (**4-*cis***)

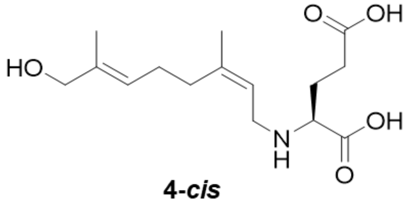

**4-*cis***

$^{1}$H NMR (500 MHz, $D_2O$ + 0.01% MeOH): δ 5.38 (t, *J* = 7.1 Hz, 1H), 5.25 (t, *J* = 8.0 1H), 3.92 (s, 2H), 3.68 – 3.59 (m, 2H), 3.57 (t, *J* = 6.7 Hz, 1H), 2.49 – 2.36 (m, 2H), 2.20 – 2.12 (m, 4H), 2.11 – 1.98 (m, 2H), 1.78 (s, 3H), 1.60 (s, 3H). $^{13}$C NMR (125 MHz, $D_2O$ + 0.01% MeOH) δ 178.5, 173.3, 147.1, 135.4, 125.7, 113.9, 67.6, 60.7, 44.2, 31.7, 31.1, 25.6, 25.4, 22.8, 13.1; HRMS (ESI) Calculated for $C_{15}H_{26}NO_5$ 300.1805, found 300.1823 $(M+H)^+$

Synthesis of L-NGG (**5-*trans***) and L-NNG (**5-*cis***)

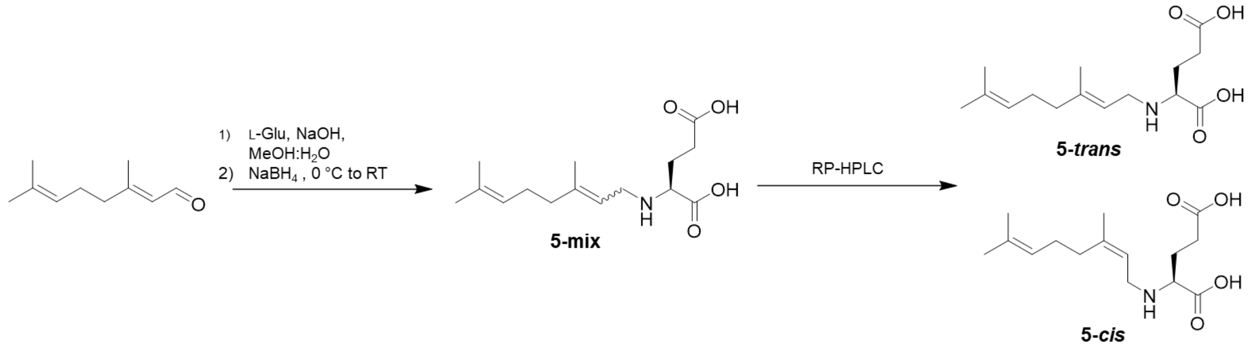


Compound **5** was synthesized as previously reported[11] to afford a white solid (0.045 g, 22%) as a mixture of regioisomers. Isomers were separated via preparative RP-HPLC (Phenomenex Kinetex 5 µm EVO C18 100 Å 250 x 21.2 mm) at a flow rate of 8 mL/min using 20% acetonitrile in 0.1% aqueous formic acid. The compounds were manually collected with the *cis* isomer eluting at 28.0 min and the *trans* isomer eluting at 30.5 min.

L-NGG (**5-*trans***)

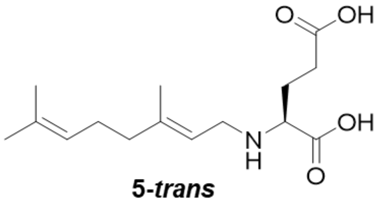

**5-*trans***

$^{1}$H NMR (500 MHz, $CD_3OD$): δ 5.30 (t, *J* = 7.6 Hz, 1H), 5.11 (t, *J* = 6.8 Hz, 1H), 3.66 (qd, *J* = 13.4, 7.5 Hz, 2H), 3.53 (t, *J* = 6.0 Hz, 1H), 2.54 (td, *J* = 7.2, 3.8 Hz, 2H), 2.19-2.03 (m, 6H), 1.76 (s, 3H), 1.68 (s, 3H), 1.62 (s, 3H); $^{13}$C NMR (125 MHz, $CD_3OD$): δ 177.1, 172.8, 147.5, 133.0, 124.7, 115.1, 62.0, 45.4, 40.7, 31.7, 27.2 , 26.7, 25.9, 17.8, 16.7; Calculated for $C_{15}H_{26}NO_4$ 284.1856, found 284.1860 $(M+H)^+$

L-NNG (**5-*cis***)

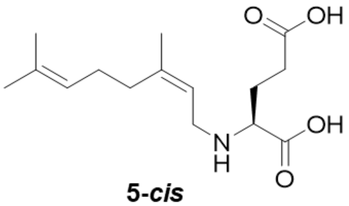

**5-*cis***

$^{1}$H NMR (500 MHz, $CD_3OD$): δ 5.30 (t, *J* = 7.6 Hz, 1H), 5.12 (t, *J* = 7.1 Hz, 1H), 3.70-3.56 (m, 2H), 3.52 (t, *J* = 6.0 Hz, 1H), 2.53 (td, J = 7.2, 3.4 Hz, 2H), 2.20-2.03 (m, 6H), 1.82 (s, 3H), 1.68 (s, 3H), 1.61 (s, 3H); $^{13}$C NMR (125 MHz, $CD_3OD$): δ 177.2, 172.8, 147.2, 133.4, 124.5, 116.0, 62.2, 45.3, 33.1, 31.8, 27.3 , 26.8, 25.9, 23.8, 17.8; HRMS (ESI) Calculated for $C_{15}H_{26}NO_4$ 284.1856, found 284.1870 $(M+H)^+$

Synthesis of 7’-carboxy-(*S*)-citronellyl-*N*-L-glutamic acid (**6**) and 7’-carboxy-(*R*)-citronellyl-L-glutamic acid (**7**)

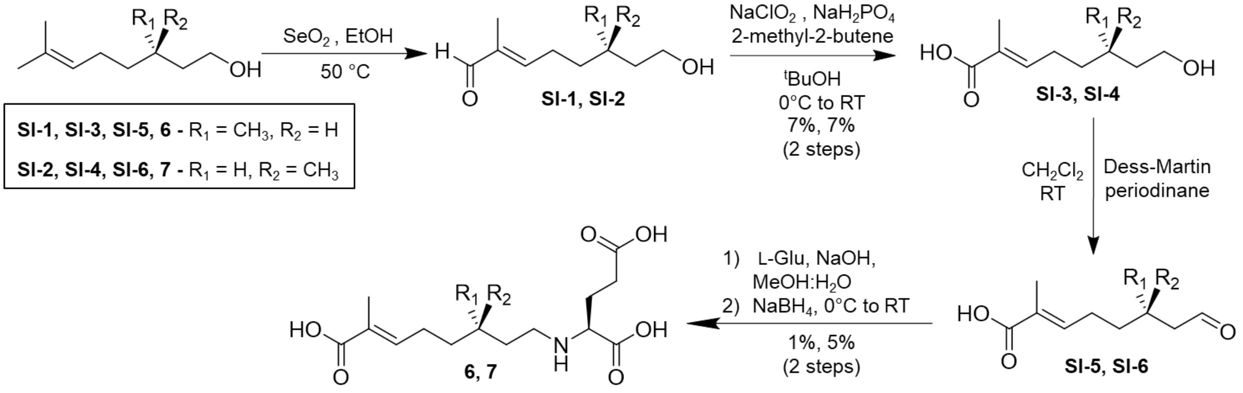


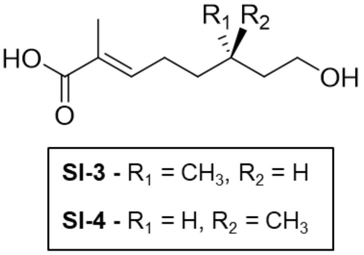


7-carboxy-(*S*)-citronellol (**SI-3**) and 7-carboxy-(*R*)-citronellol (**SI-4**)

**SI-3** and **SI-4** were synthesized via the same modified synthetic procedure.[11] Generally, to a solution of selenium dioxide (1.03 eq.) in ethanol (10 mL) refluxing at 50 °C was added a solution of either (*S*)-citronellol or (*R*)-citronellol, respectively (1.0 eq.) in ethanol (5 mL) dropwise over 1 h. The reaction was refluxed for 24 h, then warmed to room temperature and left stirring for an additional 24 h. The crude reaction was filtered through a pad of silica, concentrated i*n vacuo*, and carried forward without further purification. To a solution of crude aldehyde intermediate **SI-1** or **SI-2**, respectively, and 90% 2-methyl-2-butene (4.0 eq.) in *tert*-butanol (10 mL) at 0 °C was added an aqueous solution (10 mL) of sodium dihydrogen phosphate and 80% sodium chlorite (4.0 eq.) dropwise over 30 minutes. The reaction was stirred for 1 h at 0 °C, then warmed to room temperature and stirred for an additional 16 h. The crude reaction mixture was concentrated *in vacuo*, resuspended in water (10 mL) and extracted with diethyl ether (2 x 20 mL). The aqueous layer was then acidified to pH 2 with 1 M HCl and further extracted with diethyl ether (2 x 20 mL). Pooled organic layers were washed with brine acidified to pH < 3 with 1 M HCl, dried over magnesium sulfate, and concentrated *in vacuo*. The dried reaction mixture was then purified by silica flash chromatography (2.5% - 5% MeOH in $CHCl_3$).

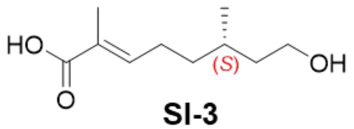

**SI-3**

7-carboxy-(*S*)-citronellol (**SI-3**)

Beginning with (*S*)-citronellol (1.000 g, 6.40 mmol) and following the previous general procedure, **SI-3** was isolated as a colorless oil (0.088 g, 0.47 mmol, 7% over two steps) and NMR characterization matched literature values.[12] $^1$H NMR (500 MHz, $CDCl_3$): δ 6.91 (t, *J* = 7.5 Hz, 1H), 3.77 – 3.67 (m, 2H), 2.31 – 2.17 (m, 2H), 1.87 (s, 3H) 1.70 – 1.60 (m, 1H), 1.55 – 1.40 (m, 2H), 1.38 – 1.30 (m, 1H), 0.96 (d, *J* = 6.5 Hz, 3H). HRMS (ESI) Calculated for $C_{10}H_{16}O_2$ 168.1149, found 168.1151 $(M+H-H_2O)^+$ (literature).[12] MS (ESI) Calculated for $C_{10}H_{17}O_3$ 185.12, found 184.93 $(M-H)^-$.

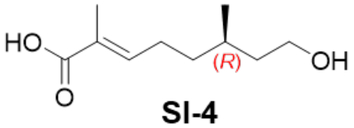

**SI-4**

7-carboxy-(*R*)-citronellol (**SI-4**)

Beginning with (*R*)-citronellol (0.500 g, 3.20 mmol) and following the previous general procedure, **SI-4** was isolated as a colorless oil (0.043 g, 0.23 mmol, 7% over two steps) and NMR characterization matched literature values.[12] $^1$H NMR (500 MHz, $CDCl_3$): δ 6.88 (t, *J* = 7.6 Hz, 1H), 3.75 – 3.64 (m, 2H), 2.28 – 2.15 (m, 2H), 1.84 (s, 3H) 1.67 – 1.58 (m, 1H), 1.52 – 1.37 (m, 2H), 1.35 – 1.27 (m, 1H), 0.93 (d, *J* = 6.4 Hz, 3H). HRMS (ESI) Calculated for $C_{10}H_{16}O_2$ 168.1149, found 168.1150 $(M+H-H_2O)^+$ (literature).[12] Calculated for $C_{10}H_{17}O_3$ 185.12, found 184.93 $(M-H)^-$.

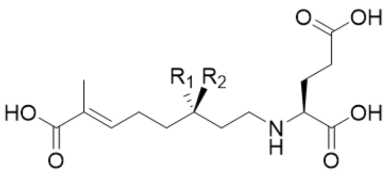

**6** - $R_1 = CH_3$, $R_2 = H$

**7** - $R_1 = H$, $R_2 = CH_3$

**6** and **7** were synthesized via the same modified synthetic procedure.[11] Generally, to a solution of either **SI-3** (0.116 g, 0.62 mmol) or **SI-4** (0.106 g, 0.57 mmol), respectively, in anhydrous $CH_2Cl_2$ at room temperature was added Dess-Martin periodinane at once and the reaction was stirred for 1 h. The reaction was quenched with 5% aqueous sodium thiosulfate (10 mL), then transferred to a separatory funnel and diluted with water (10 mL). The layers were separated and the organic layer was washed with water (20 mL). The aqueous layer was then washed with $CH_2Cl_2$ (3 x 20 mL). Pooled organic layers were washed with brine (20 mL), dried over magnesium sulfate, filtered, and concentrated *in vacuo*. The reaction mixture was then purified by silica flash chromatography (4:1 hexanes/EtOAc + 0.1% AcOH) and carried forward without further purification. To an aqueous solution (2.5 mL) of L-glutamic acid (2 eq.) and sodium hydroxide (3 eq.) was added to a solution of either semi-crude aldehyde intermediate **SI-5** or **SI-6**, respectively (1.0 eq.) in methanol (2.5 mL) and the reaction was stirred for 3 h at room temperature. The reaction was cooled to 0 °C and sodium borohydride (1.3 eq.) was added at once and stirred for 1 h. The reaction was warmed to room temperature, acidified to pH 3 with 1 M HCl, and concentrated *in vacuo* to remove the majority of the methanol. It was necessary for some methanol to remain in order to maintain product solubility. The crude reaction mixture was subsequently purified via a Biotage Isolera Prime flash chromatography system using a Sfar Ultra C18 D 6g pre-packed column at a flow rate of 6 mL/min with the following method: 10% B (2.5 column volumes (CV)), 10 - 30% B (9 CV), 30 - 45% B (2 CV), 45 - 95% B (1 CV), 95% B (3 CV), 95 - 10% B (1 CV), 10% B (2 CV), where A = 0.1 % aqueous formic acid, and B = 0.1% formic acid in acetonitrile.

<u>7'-carboxy-(*S*)-citronellyl-L-glutamic acid (**6**)</u>

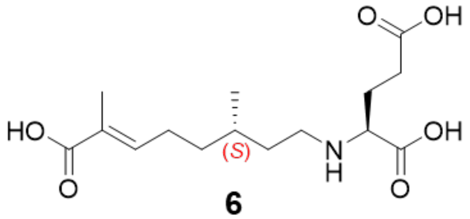


**6**

Compound **6** was isolated as a white powder (0.002 g, 0.006 mmol, 1% over two steps). $^1$H NMR (500 MHz, $CD_3OD$) δ 6.77 (t, *J* = 7.5 Hz, 1H), 3.53 (t, *J* = 6.1 Hz, 1H), 3.08 (td, *J* = 11.4, 4.8 Hz, 1H), 2.98 (ddd, *J* = 12.1, 10.4, 5.4 Hz, 1H), 2.54 (td, *J* = 7.1, 3.7 Hz, 2H), 2.28 – 2.19 (m, 2H), 2.18 – 2.04 (m, 2H), 1.81 (d, *J* = 1.3 Hz, 3H), 1.79 – 1.73 (m, 1H), 1.64 – 1.54 (m, 2H), 1.54 – 1.46 (m, 1H), 1.38 – 1.29 (m, 1H), 0.97 (d, *J* = 6.1 Hz, 3H); $^{13}$C NMR (126 MHz, $CH_3OD$) δ 178.2, 173.3, 172.1, 144.0, 129.1, 63.6, 46.5, 36.3, 34.1, 32.3, 31.6, 26.9, 26.6, 19.3, 12.5; HRMS (ESI) Calculated for $C_{15}H_{26}NO_6$ 316.1755, found 316.1769 $(M+H)^+$

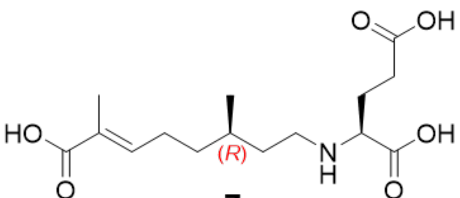


**7**

7’-carboxy-(*R*)-citronellyl-L-glutamic acid (**7**)

Compound **7** was isolated as a white powder (0.010 g, 0.03 mmol, 5% over two steps). $^{1}$H NMR (500 MHz, $CD_3OD$) δ 6.77 (t, *J* = 7.4 1H), 3.93 (dd, *J* = 7.6, 5.2 Hz, 1H), 3.17-3.05 (m, 2H), 2.58 (td, *J* = 7.2, 3.9 Hz, 2H), 2.30 – 2.19 (m, 3H), 2.19 – 2.10 (m, 1H), 1.82 (s, 3H), 1.80-1.74 (m, 1H), 1.64 – 1.53 (m, 2H), 1.53 – 1.46 (m, 1H), 1.39-1.31 (m, 1H), 0.98 (d, *J* = 6.2 Hz, 3H); $^{13}$C NMR (125 MHz, $CD_3OD$) δ 176.1, 172.9, 171.7, 143.6, 129.1, 61.1 46.4, 36.3, 34.0, 31.5, 30.5, 26.9, 25.8, 19.3, 12.4; HRMS (ESI) Calculated for $C_{15}H_{26}NO_6$ 316.1755, found 316.1763 $(M+H)^+$

Synthesis of 7’-hydroxy-(*R*)-citronellyl-*N*-L-glutamic acid (**9**) and 7’-hydroxy-(*S*)-citronellyl-L-glutamic acid (**10**)

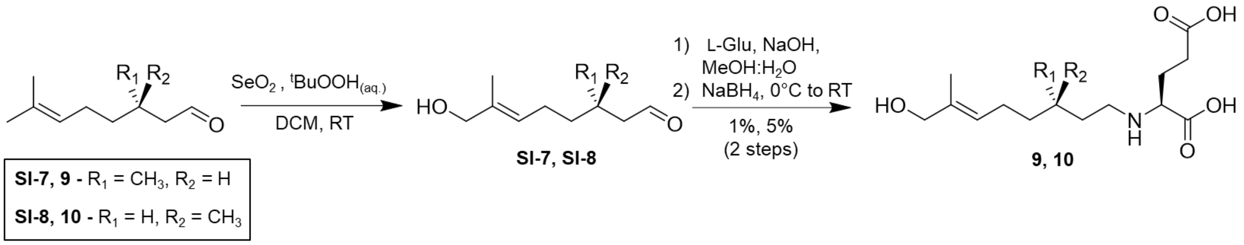


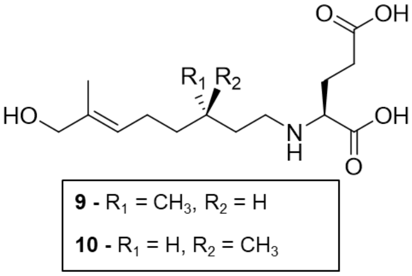


Compounds **9** and **10** were synthesized via modification of a previously reported synthetic procedure.[11] Generally, to a suspension of selenium (IV) oxide (0.1 eq.) in DCM (5 ml), a solution of citronellal (1.0 equivalent) in DCM (5 mL) was added followed by aqueous *tert*-butyl hydrogen peroxide (2.0 eq. 70% in decane). The mixture was stirred at room temperature for 22 h. The reaction was washed subsequently with saturated sodium bicarbonate (25 mL) and brine (25 mL). The organic layer was dried with magnesium sulfate, filtered, and concentrated in vacuo and purified via silica flash chromatography (1:1 hexane/EtOAc) and carried forward without further purification. An aqueous solution (2.5 mL) of L-glutamic acid (2 eq.) and sodium hydroxide (3 eq.) was added to the semi-crude solution of either **SI-7** or **SI-8**, respectively (1.0 eq.) in methanol (2.5 mL) and the reaction was stirred for 3 h at room temperature. The reaction was cooled to 0 °C and sodium borohydride (1.3 eq.) was added at once and stirred for 1 h. The reaction was warmed to room temperature and acidified to pH 3 with 1M HCl. The reaction was concentrated *in vacuo* to remove the majority of the methanol. It was necessary for some methanol to remain in order to maintain product solubility. The reaction was then purified via a Biotage Isolera Prime flash chromatography system using a Sfar C18 D 6g pre-packed column at a flow rate of 6 mL/min with the following method: 5% B (2.5 CV), 5 - 20% B (7 CV), 20 - 30% B (2 CV), 30 - 95% B (2 CV), 95% B (5 CV), 95 - 50% B (1 CV), 50% B (5 CV), where A = 0.1 % aqueous formic acid, and B = 0.1% formic acid in acetonitrile.

7'-hydroxy-(*S*)-citronellyl-L-glutamic acid (**9**)

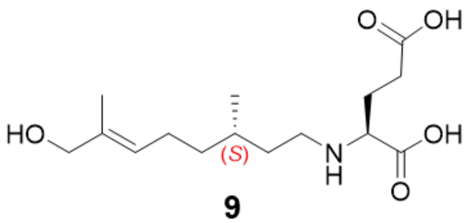


**9**

0.97 mmol *S*-citronellal was used in the aforementioned reductive amination general procedure, and **9** was isolated as a white solid (0.015 g, 0.05 mmol, 5% over two steps). $^1$H NMR (500 MHz, $CD_3OD$) δ 5.40 (t, *J* = 7.4 Hz, 1H), 3.91 (s, 2H), 3.51 (t, *J* = 6.0 Hz, 1H), 3.06 (td, *J* = 11.3, 4.7 Hz, 1H), 2.96 (td, *J* = 11.3, 5.4 Hz, 1H), 2.54 (dt, *J* = 7.4, 4.7 Hz, 2H), 2.16-2.03 (m, 4H), 1.78-1.73 (m, 1H), 1.65 (s, 3H), 1.62-1.51 (m, 2H), 1.46-1.38 (m, 1H), 1.30-1.23 (m, 1H), 0.96 (d, *J* = 6.1 Hz, 3H); $^{13}$C NMR (125 MHz, $CD_3OD$) δ 177.7, 173.1, 136.1, 126.5, 68.9, 63.6, 46.6, 37.5, 34.3, 32.2, 31.5, 26.7, 25.8, 19.5, 13.7; HRMS (ESI) Calculated for $C_{15}H_{28}NO_5$ 302.1962, found 302.1976 $(M+H)^+$

7'-hydroxy-(*R*)-citronellyl-*N*-L-glutamic acid (**10**)

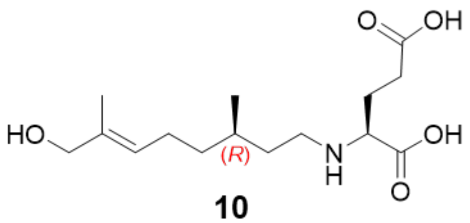


**10**

0.68 mmol *R*-citronellal was used in the aforementioned reductive amination general procedure, and **10** was isolated as a white solid (0.004 g, 0.01 mmol, 1% over two steps). $^1$H NMR (500 MHz, $CD_3OD$) δ 5.40 (t, *J* = 7.3 Hz, 1H), 3.92 (s, 2H), 3.53 (t, *J* = 6.1 Hz, 1H), 3.02 (t, *J* = 8.0 Hz, 2H), 2.61 – 2.51 (m, 2H), 2.16-2.03 (m, 4H), 1.82 – 1.70 (m, 1H), 1.66 (s, 3H), 1.61-1.48 (m, 2H), 1.47 – 1.38 (m, 1H), 1.31-1.23 (m, 1H), 0.96 (d, *J* = 6.3 Hz, 3H); $^{13}$C NMR (126 MHz, $CD_3OD$) δ 177.2, 172.9, 136.2, 126.4, 68.9, 63.4, 46.6, 37.6, 34.2, 31.7, 31.5, 26.7, 25.9, 19.5, 13.7; HRMS (ESI) Calculated for $C_{15}H_{28}NO_5$ 302.1962, found 302.1971 $(M+H)^+$

Synthesis of (*S*)-citronellyl-*N*-L-glutamic acid (**11**) and (*R*)-citronellyl-*N*-L-glutamic acid (**12**)

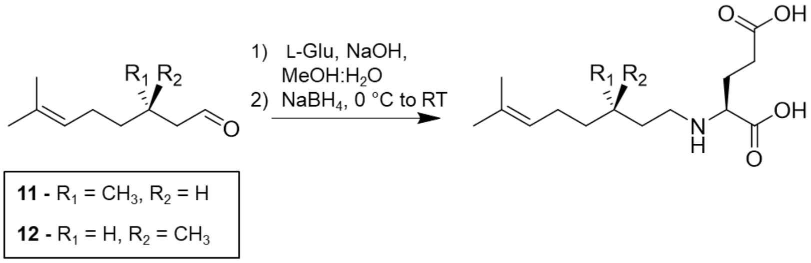


Compounds **11** and **12** were synthesized via modification of a previously reported synthetic procedure.[11] Generally, an aqueous solution (2.5 mL) of L-glutamic acid (2 eq.) and sodium hydroxide (3 eq.) was added to a solution of either *(R)*- or *(S)*-citronellal (1.0 eq.) in methanol (2.5 mL) and the reaction was stirred for 3 h at room temperature. The reaction was cooled to 0 °C and sodium borohydride (1.3 eq.) was added at once and stirred for 1 h. The reaction was warmed to room temperature and acidified to pH 3 with 1 M HCl. The reaction was concentrated *in vacuo* to remove the majority of the methanol. It was necessary for some methanol to remain in order to maintain product solubility. The reaction was then purified via a Biotage Isolera Prime flash chromatography system using a SNAP Ultra C18 60g pre-packed column at a flow rate of 50 mL/min with the following method: 10% B (2.5 CV), 10 - 30% B (2.5 CV), 30 - 45% B (5 CV), 45 - 95% B (1 CV), 95% B (2 CV), 95 - 10% B (1 CV), 10% B (2 CV), where A = 0.1 % aqueous formic acid, and B = 0.1% formic acid in acetonitrile.

(*S*)-citronellyl-*N*-L-glutamic acid (**11**)

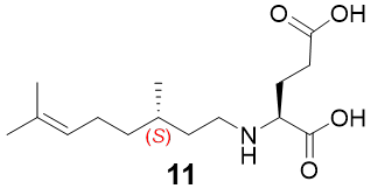


**11**

(*S*)-citronellal (0.100 g, 0.65 mmol) was used in the previous reductive amination general procedure, isolating **11** as a white solid (0.053 g, 0.15 mmol, 27%). $^1$H NMR (500 MHz, $CD_3OD$): δ 5.11 (t, *J* = 7.1 Hz, 1H), 3.54 (t, *J* = 6.1 Hz, 1H), 3.07 (td, *J* = 11.7, 4.8 Hz, 1H), 3.00-2.93 (m, 1H), 2.55 (t, *J* = 7.3 Hz, 2H), 2.18 – 1.96 (m, 4H), 1.77-1.69 (m, 1H), 1.68 (s, 3H), 1.62 (s, 3H), 1.60-1.49 (m, 2H), 1.42-1.34 (m, 1H), 1.26 – 1.17 (m, 1H), 0.95 (d, *J* = 6.3 Hz, 3H); $^{13}$C NMR (125 MHz, $CD_3OD$): δ 176.9, 172.8, 132.5, 125.4, 63.2, 46.6, 37.9, 34.3, 31.6, 31.4, 26.5, 26.3, 25.9, 19.5, 17.7; HRMS (ESI) Calculated for $C_{15}H_{28}NO_4$ 286.2013, found 286.2026 $(M+H)^+$

(*R*)-citronellyl-*N*-L-glutamic acid (**12**)

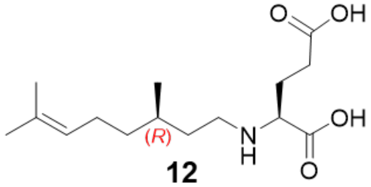


**12**

(*R*)-citronellal (0.115 g, 0.75 mmol) was used in the previous reductive amination general procedure, isolating **12** as a white solid (0.087 g, 0.25 mmol, 41%). $^1$H NMR (500 MHz, $CD_3OD$): δ 5.11 (t, *J* = 7.1 Hz, 1H), 3.53 (t, *J* = 6.1 Hz, 1H), 3.05-2.99 (m, 2H), 2.55 (t, *J* = 7.3 Hz, 2H), 2.20 – 1.95 (m, 4H), 1.78-1.70 (m, 1H), 1.68 (s, *J* = 1.4 Hz, 3H), 1.62 (s, *J* = 1.3 Hz, 3H), 1.59-1.48 (m, 2H), 1.41-1.33 (m, 1H), 1.27-1.18 (m, 1H), 0.95 (d, *J* = 6.3 Hz, 3H). $^{13}$C NMR (125 MHz, $CD_3OD$) δ 177.0, 172.8, 132.5, 125.4, 63.3, 46.6, 38.0, 34.3, 31.5, 31.5, 26.6, 26.3, 25.9, 19.5, 17.7; HRMS (ESI) Calculated for $C_{15}H_{28}NO_4$ 286.2013, found 286.2029 $(M+H)^+$

Synthesis of 3*R*-hydroxy-L-glutamic acid (**SI-9**)

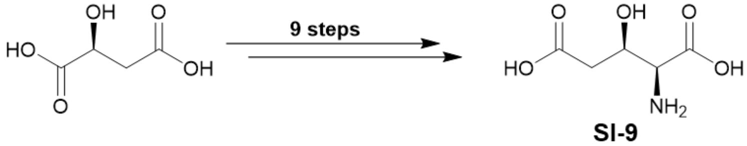


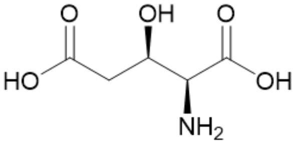


**SI-9**

Compound **SI-9** was synthesized as was previously reported for the first 8 steps from L-malic acid (25 g).[13,14] However, the final step was adjusted Dowex cation exchange purification of **SI-9** was not performed and the product was lyophilized, isolated, and carried forward without any additional purification. The product was isolated as an orange-brown solid (0.048 g, 0.294 mmol, 95% purity, 1% yield over 9 steps). $^1$H NMR (500 MHz, $D_2O$ + 0.1% FA) δ 4.62 (ddd, *J* = 9.0, 4.6, 4.6 Hz, 1H), 4.03 (d, *J* = 4.7 Hz, 1H), 2.90 (dd, *J* = 16.4, 4.4 Hz, 1H), 2.74 (dd, *J* = 16.4, 8.7 Hz, 1H); $^{13}$C NMR (125 MHz, $D_2O$ + 0.1% FA) δ 174.9, 171.4, 66.4, 58.3, 39.2; HRMS (ESI) Calculated for $C_5H_{10}NO_5$ 164.0553, found 164.0553 $(M+H)^+$

<u>Synthesis of 7'-carboxy-*N*-neryl-(3*R*)-hydroxy-L-glutamic acid (**3**)</u>

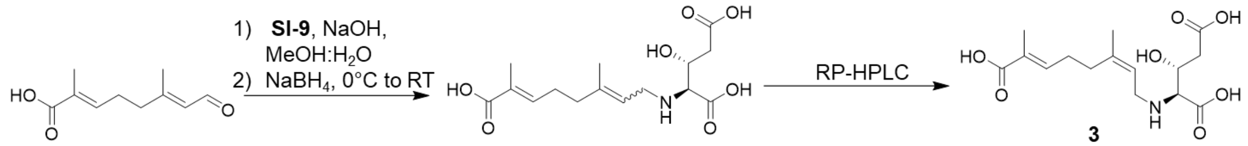


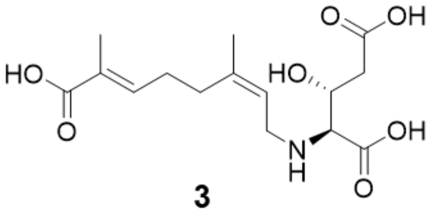

**3**

Compound **3** was synthesized via modification of a previously reported procedure.[11] Briefly, an aqueous solution of **SI-9** (0.026 g, 0.16 mmol) and sodium hydroxide (0.016 g, 0.4 mmol) in 2.0 mL of water was stirred at room temperature. A solution of 7-carboxygeranial (0.015 g, 0.08 mmol) in methanol (2.0 mL) was added dropwise and stirred for 3 h. The reaction mixture was then cooled to 0 °C and sodium borohydride (0.039 g, 1.03 mmol) was added at once and stirred for an additional 1.5 h. The reaction mixture was warmed to room temperature over 15 min, then acidified to pH 3.0 by the addition of 1 M HCl. The organic solvent was removed *in vacuo* and the crude was then purified via preparative RP-HPLC (Phenomenex Kinetex 5 µm EVO C18 100 Å 250 x 21.2 mm) at a flow rate of 7 mL/min using 12% acetonitrile in 0.1% aqueous formic acid. The peak containing **3** eluted at 21.5 min and was manually collected, concentrated *in vacuo* and lyophilized. The sample was then further purified via analytical HPLC (Phenomenex Luna 5 µm C18(2) 100 Å 150 x 4.6 mm) at a flow rate of 1 mL/min using 11.5% acetonitrile in 0.1% aqueous formic acid. The peak containing **3** eluted at 11.5 min and was manually collected, concentrated *in vacuo* and lyophilized to afford the product as a white solid (0.001 g, 0.004 mmol, 5 %) $^1$H NMR (500 MHz, $D_2O$ + 0.01 % MeOH) δ 6.27 (t, *J* = 6.7 Hz, 1H), 5.26 (t, *J* = 7.7 Hz, 1H), 4.25 (dt, *J* = 7.5, 5.3 Hz, 1H), 3.70 – 3.61 (m, 2H), 3.47 (d, *J* = 5.4 Hz, 1H), 2.61 (dd, *J* = 16.0, 4.6 Hz, 1H), 2.46 (dd, *J* = 16.1, 7.2 Hz, 1H), 2.28

– 2.16 (m, 4H), 1.79 (s, 3H), 1.71 (s, 3H); HRMS (ESI) Calculated for $C_{15}H_{24}NO_7$ 330.1547, found 330.1545 $(M+H)^+$

Synthesis of *N*-isopentyl-L-glutamic acid (**13**)

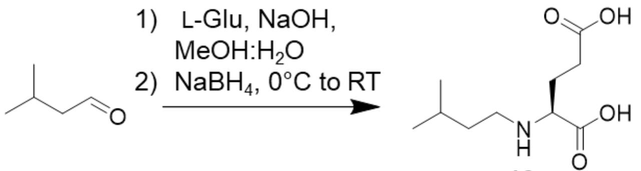

**13**

Compound **13** was synthesized via modification of a previously reported literature procedure.[15] Briefly, an aqueous solution of L-glutamate (0.100 g, 0.68 mmol) and sodium hydroxide (0.084 g, 2.11 mmol) in 2.0 mL of water was stirred at room temperature. A solution of isovaleraldehyde (0.10 mL, 0.96 mmol) in methanol (2.0 mL) was added dropwise and stirred for 5 min. The reaction mixture was then cooled to 0 °C and sodium borohydride (0.034 g, 0.89 mmol) was added at once and stirred for an additional 20 min. The reaction mixture was warmed to room temperature over 15 min, then acidified to pH 4.0 by the addition of formic acid. The organic solvent was removed *in vacuo* and the crude was then purified via a Biotage Isolera Prime flash chromatography system using a Sfar C18 D 6 g pre-packed column at a flow rate of 6 ml/min with the following method: 1% B 3 CV), 1 - 20% B (6 CV), 20 - 95% B (2 CV), 95% B (2 CV), 95% - 1% B (2 CV), 1% B (3 CV), where A = 0.1 % aqueous formic acid, and B = 0.1% formic acid in acetonitrile. The product was isolated as a white solid (0.014 g, 0.06 mmol, 10%). $^1$H NMR (500 MHz, $D_2O$) δ 3.61 (t, *J* = 6.3 Hz, 1H), 3.03 (t, *J* = 8.0 Hz, 2H), 2.52 - 2.41 (m, 2H), 2.17 - 2.01(m, 2H), 1.69 - 1.61 (m, 1H), 1.60 - 1.52 (m, 2H), 0.90 (s, 3H), 0.88 (s, 3H); $^{13}$C NMR (125 MHz, $D_2O$) δ 178.0, 173.1, 61.6, 45.3, 34.1, 30.8, 25.0, 24.9, 21.2, 21.1; HRMS (ESI) Calculated for $C_{10}H_{20}NO_4$ 218.1387, found 218.1396 $(M+H)^+$

NMR and Compound Characterization

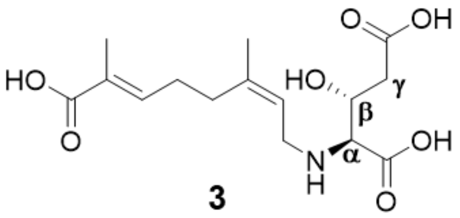

**3**

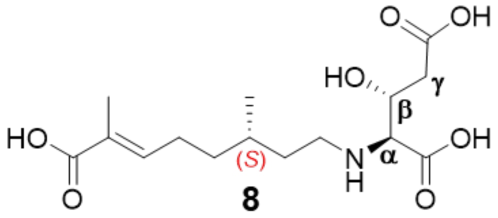

**8**

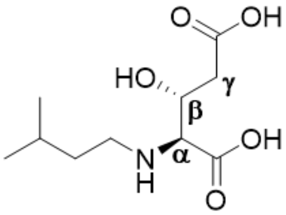

**14**

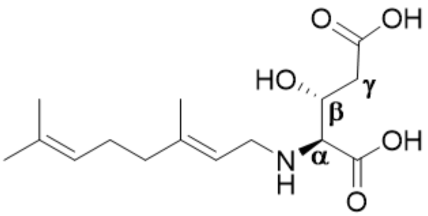

*N*-geranyl-3-(*R*)-hydroxy-L-glutamic acid

| $^1$H | 3 - synthetic | 3 – enzymatic | 8 | 14 | *N*-geranyl-3-(*R*)-hydroxy-L-glutamic acid - isolated[16] |
|---|---|---|---|---|---|
| α | 3.47 (d, *J* = 5.4 Hz, 1H) | 3.46 (d, *J* = 5.5 Hz, 1H) | 3.52 (d, *J* = 5.3 Hz, 1H) | 3.57 (d, *J* = 7.2 Hz, 1H) | 3.46 (d, *J* = 6.7 Hz, 1H) |
| β | 4.25 (dt, *J* = 7.5, 5.3 Hz, 1H) | 4.25 (dt, *J* = 7.4, 4.9 Hz, 1H) | 4.27 (dt, *J* = 7.2, 4.9 Hz, 1H) | 4.29 (td, *J* = 7.7, 3.9 Hz, 1H) | 4.31 (m, 1H) |
| $γ_a$ | 2.61 (dd, *J* = 16.0, 4.6 Hz, 1H) | 2.61 (dd, *J* = 15.9, 4.5 Hz,1H) | 2.64 (dd, *J* = 16.0, 4.6 Hz,1H) | 2.78 (dd, *J* = 16.3, 3.9 Hz,1H) | 2.82 (dd, J = 16.3, 3.7 Hz, 1H) |
| $γ_b$ | 2.46 (dd, *J* = 16.1, 7.2 Hz, 1H) | 2.46 (dd, *J* = 15.9, 7.4 Hz, 1H) | 2.51 (dd, *J* = 16.0, 7.0 Hz, 1H) | 2.61 (dd, J = 16.3, 8.2 Hz, 1H) | 2.60 (dd, J = 16.1, 7.9 Hz, 1H) |

**Table S1.** Glutamic acid moiety $^1$H NMR shifts of β-hydroxylated molecules in this study. Enzymatic products **3**, **8**, and **14** were compared to synthetic standard **3** and isolated red algal metabolite *N*-geranyl-3-(*R*)-hydroxy-L-glutamic acid.

| $^{13}C$ | **3** - synthetic | **3** - enzymatic | **8** | **14** | *N*-geranyl-3-(*R*)-hydroxy-L-glutamic acid - isolated |
|---|---|---|---|---|---|
| **α** | 65.0 | 65.2 | 66.8 | 66.7 | 66.7 |
| **β** | 67.4 | 67.6 | 67.4 | 66.7 | 68.5 |
| **γ** | 42.0 | 42.2 | 42.2 | 39.9 | 41.3 |

**Table S2.** Glutamic acid moiety $^{13}C$ NMR shifts of β-hydroxylated molecules in this study. Enzymatic products **3**, **8**, and **14** were compared to synthetic standard **3** and isolated red algal metabolite *N*-geranyl-3-(*R*)-hydroxy-L-glutamic acid.

| | |
|---|---|
| radC1-F | GGTGCCGCGCGGCAGCCATATGTTCACCATTAAGGGTACC |
| radC2-R | GGTGGTGGTGGTGCTCGAGTTAATAATATCCGTGCAGGAATTTATAC |

**Table S3.**
Primers used in this study.

#### 7’-carboxy-L-NGG - synthetic (**1-*trans***)

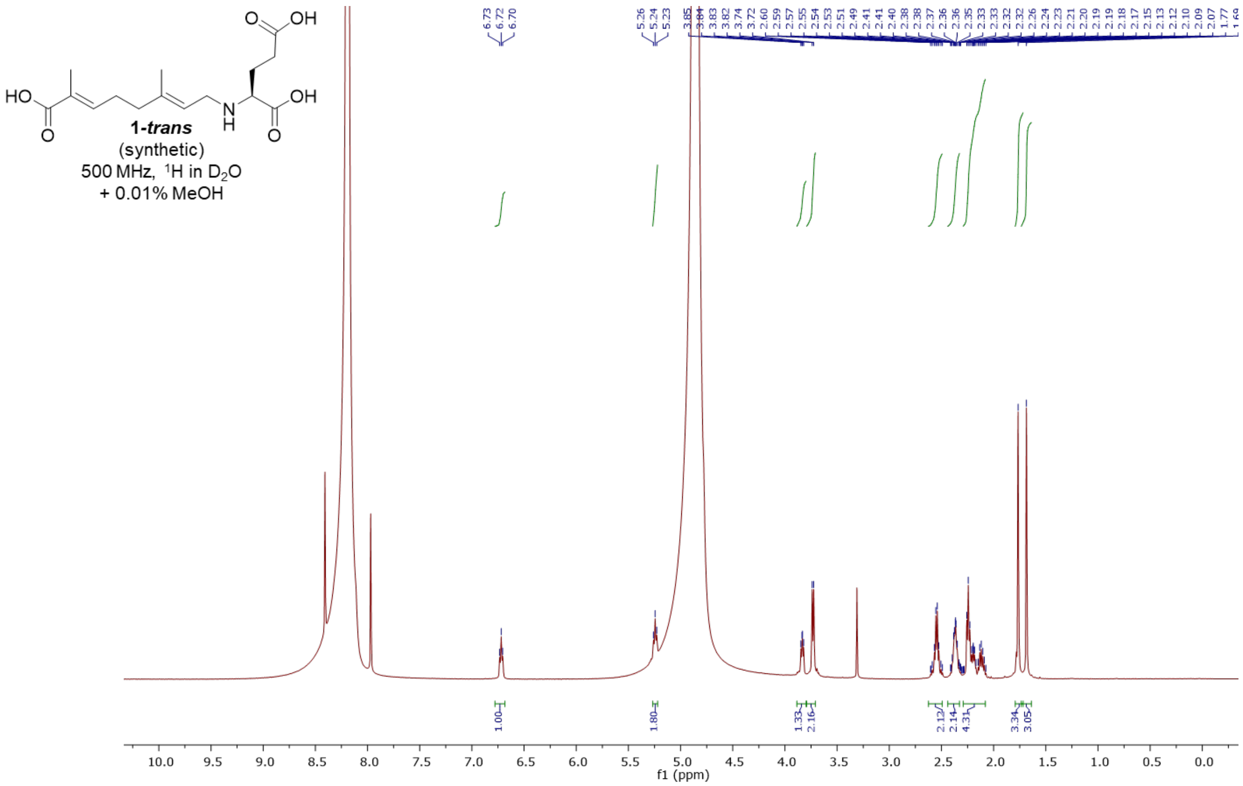


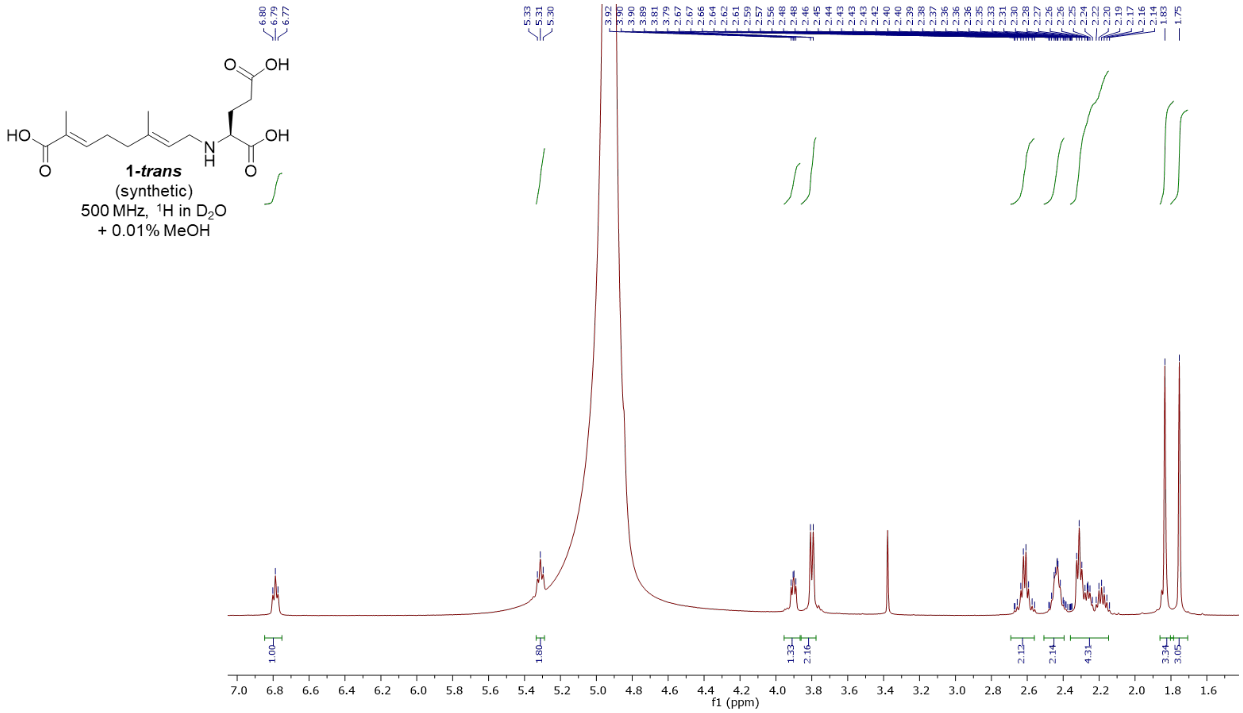

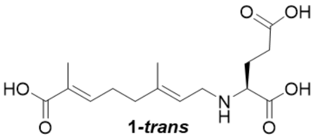

***1-trans***
(synthetic)
500 MHz, $^{13}C$ in $D_2O$
+ 0.01% MeOH

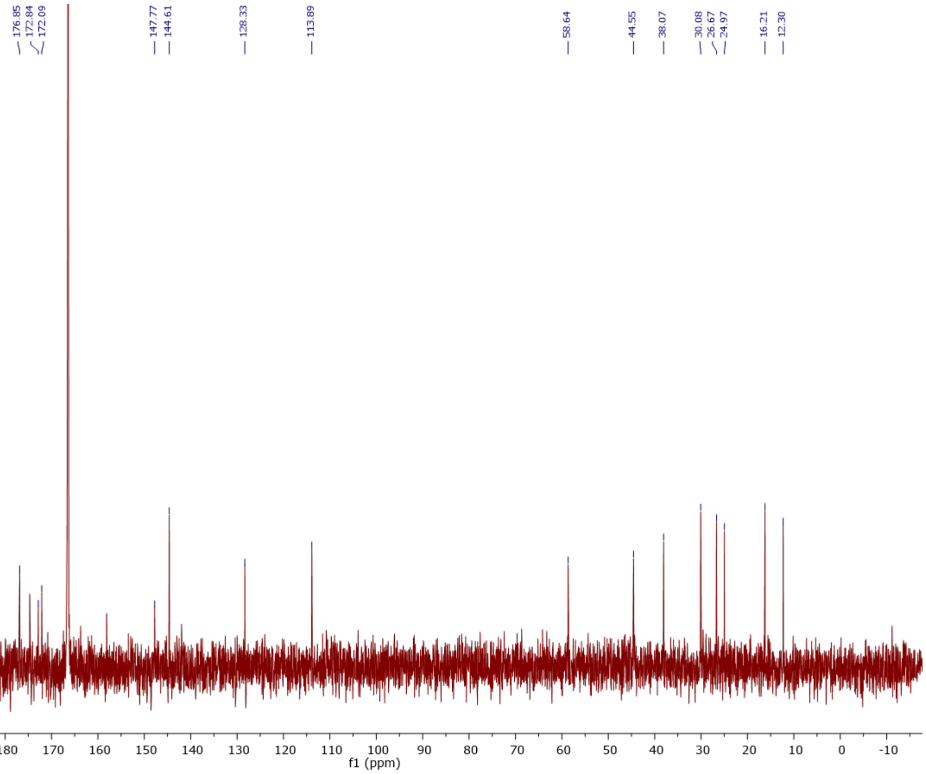


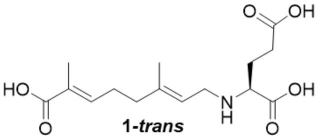

***1-trans***
(synthetic)
500 MHz, COSY in $D_2O$
+ 0.01% MeOH

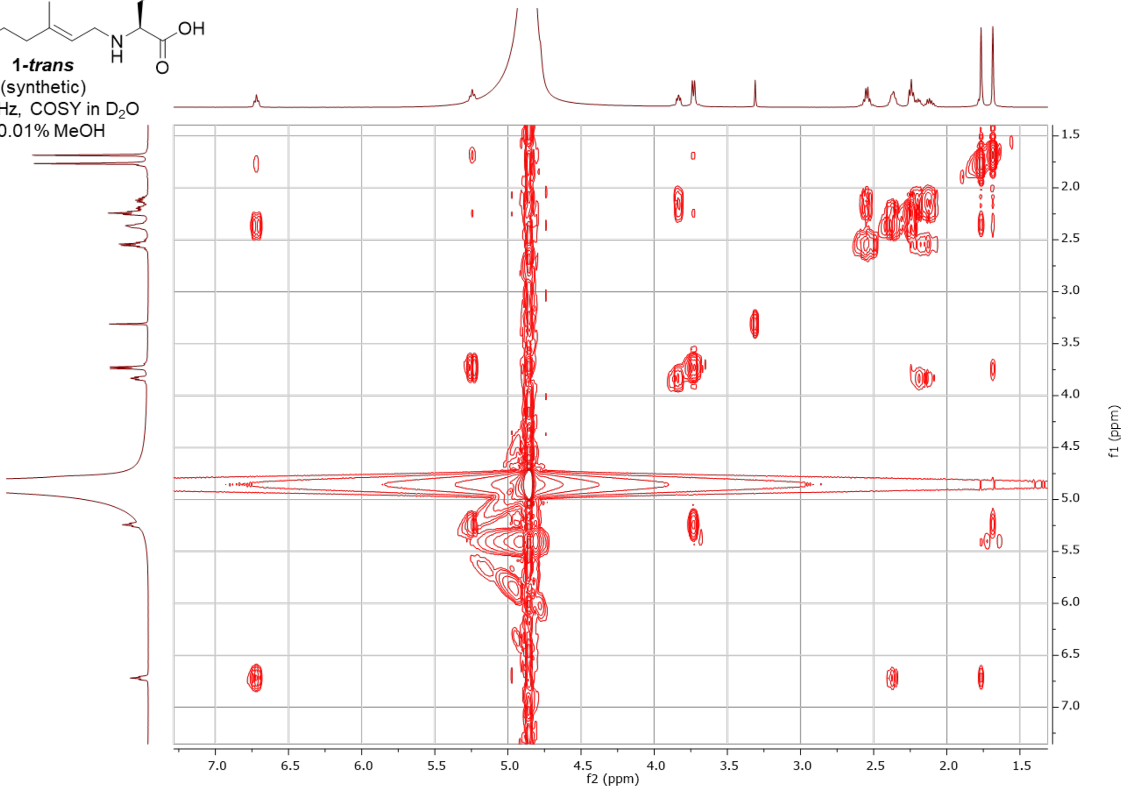

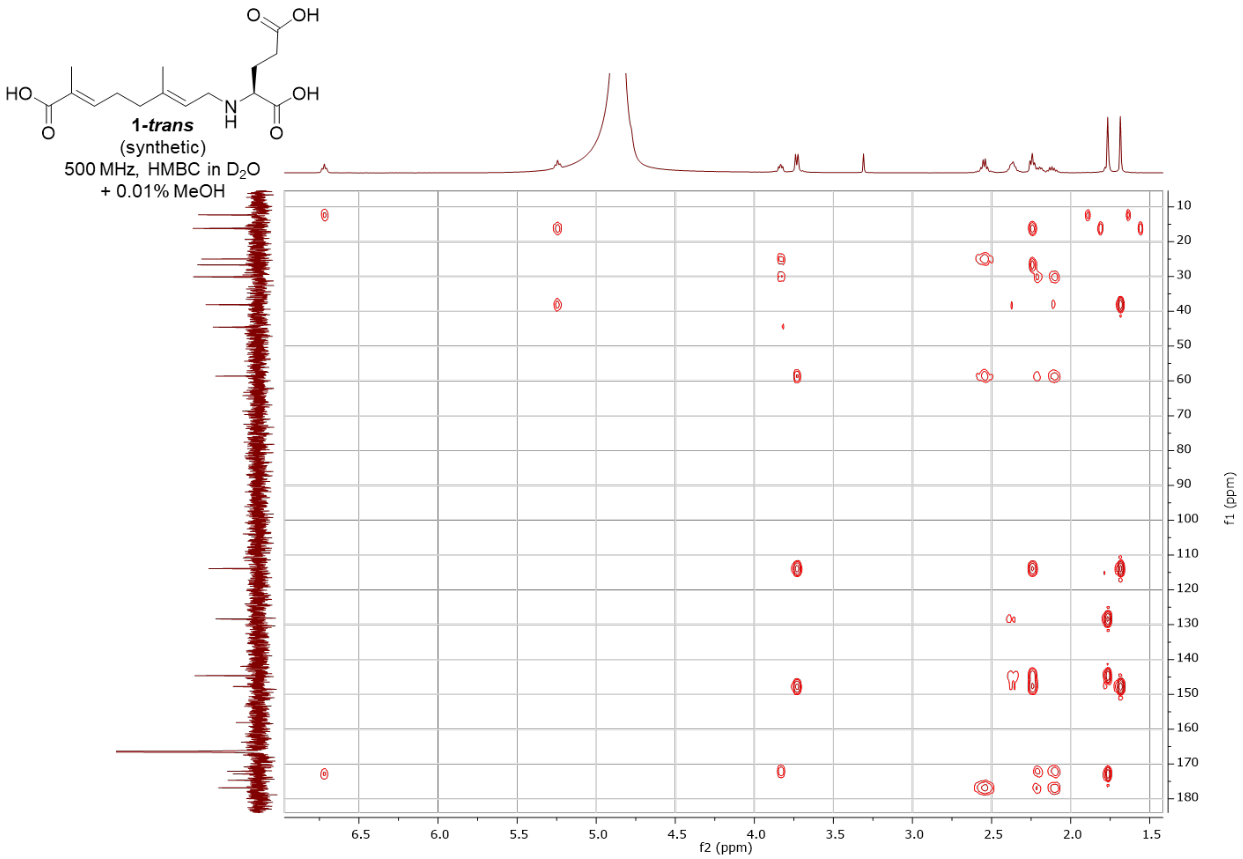
1-trans
(synthetic)
500 MHz, HMBC in $D_2O$
+ 0.01% MeOH
f2 (ppm)
f1 (ppm)

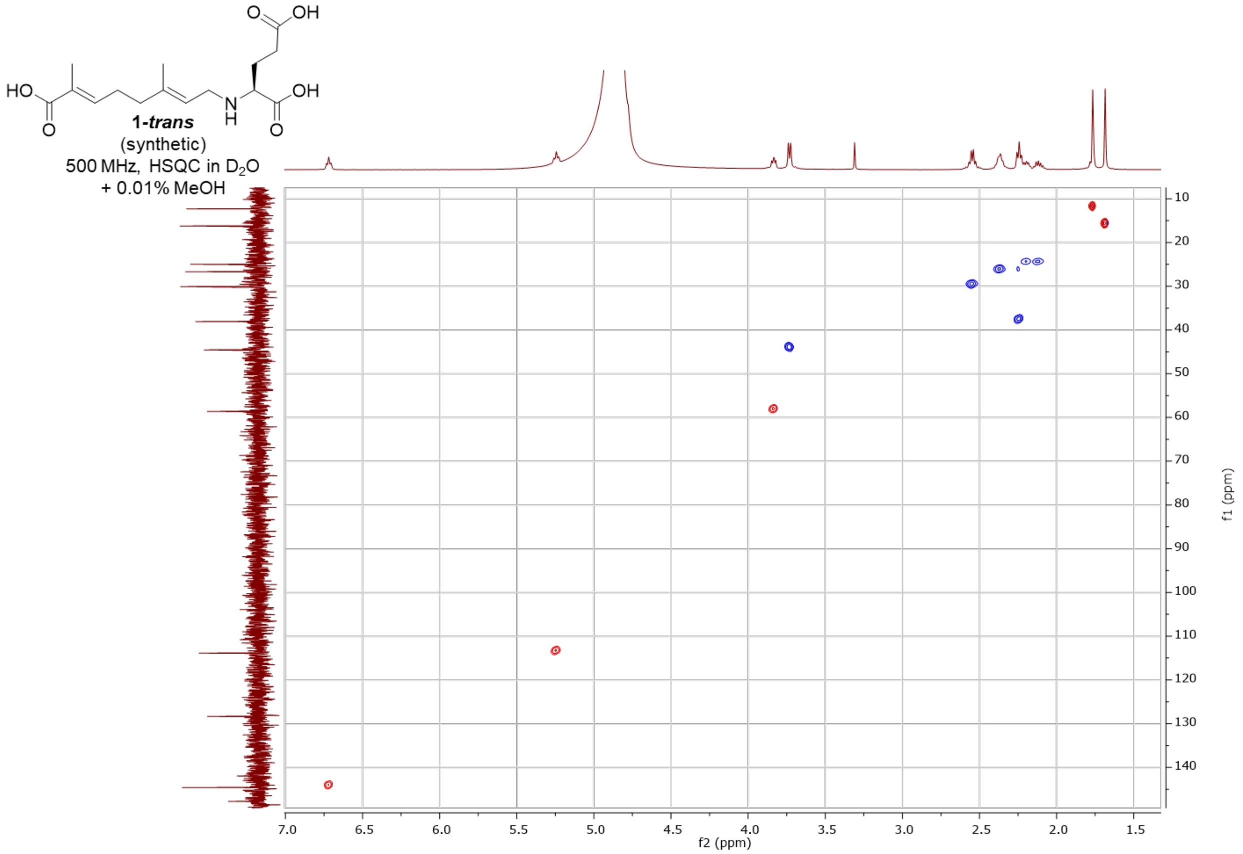
1-trans
(synthetic)
500 MHz, HSQC in $D_2O$
+ 0.01% MeOH
f2 (ppm)
f1 (ppm)

#### 7’-carboxy-L-NNG - synthetic (**1-*cis***)

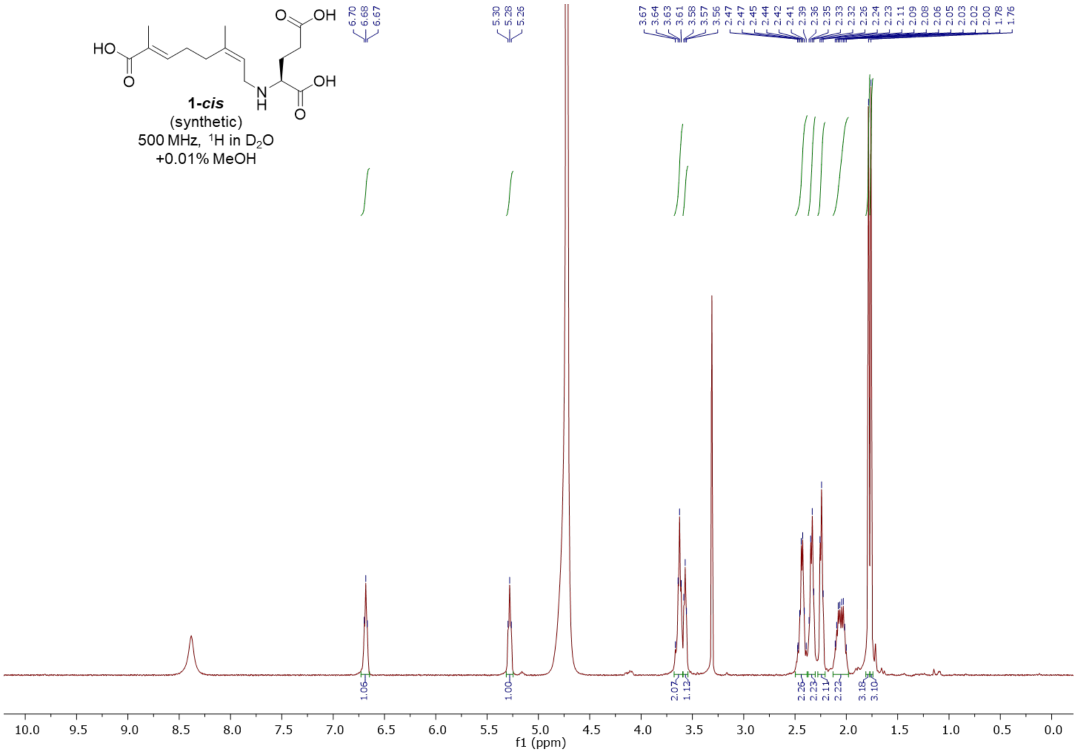


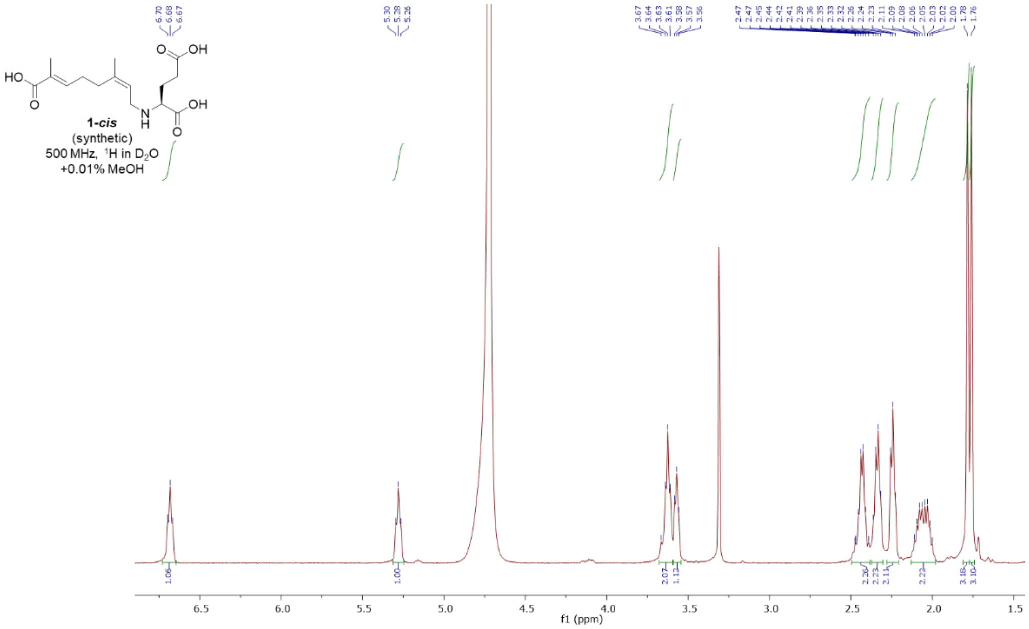

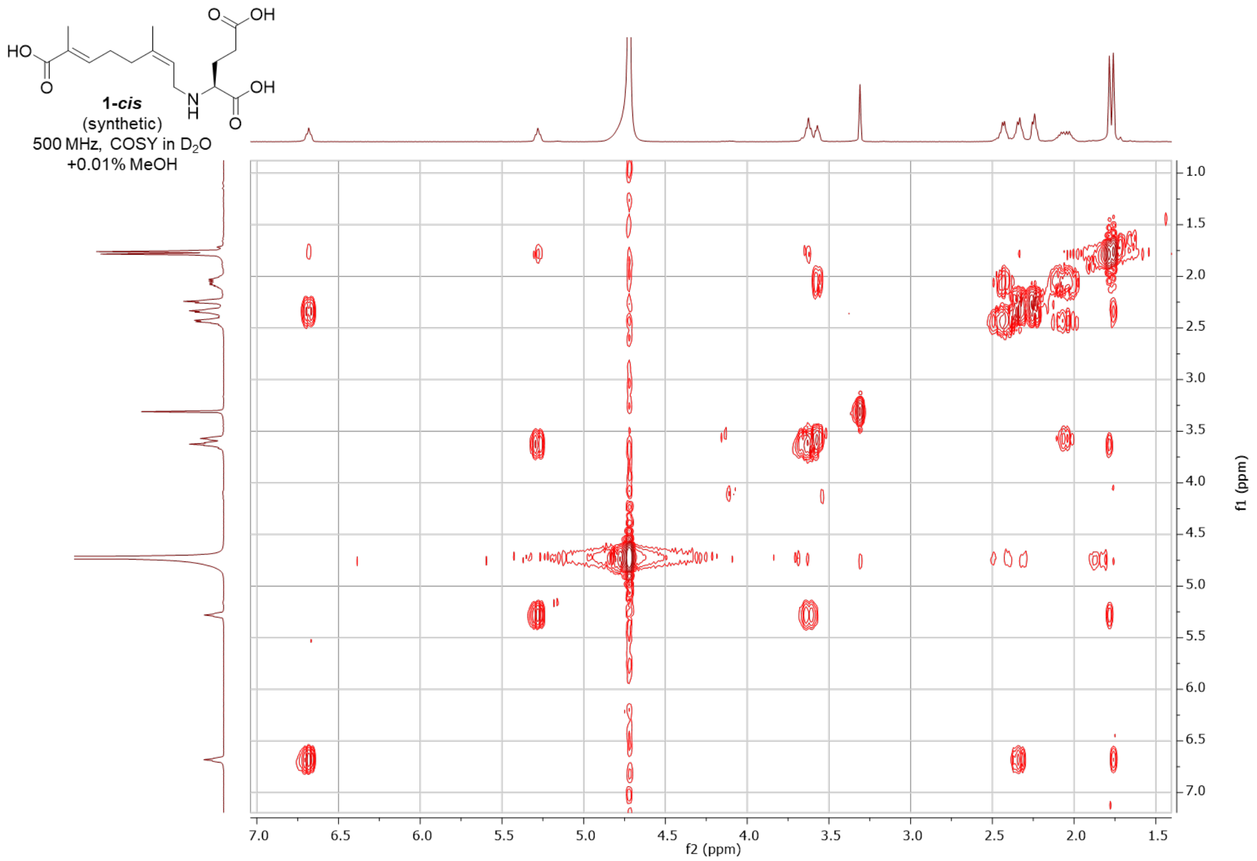

HO
O
O
OH
N
H
OH
O
1-*cis*
(synthetic)
500 MHz, COSY in $D_2O$
+0.01% MeOH
f2 (ppm)
f1 (ppm)


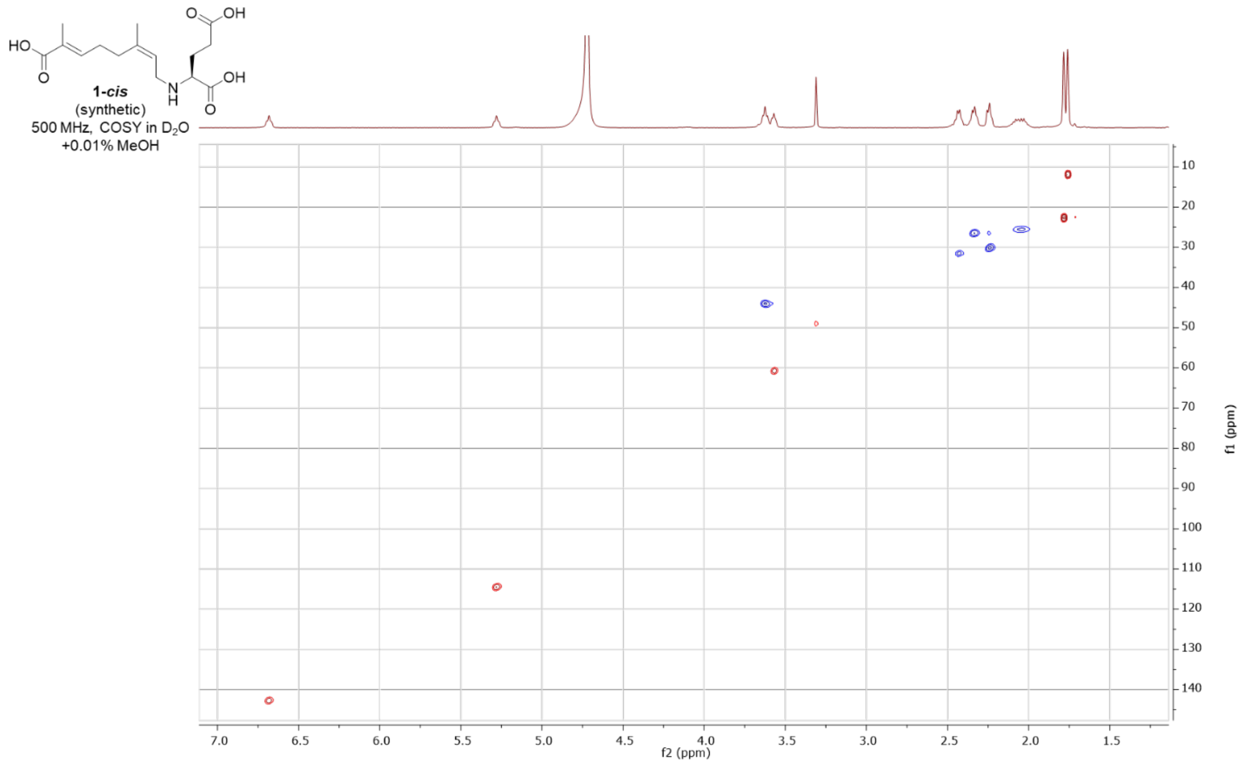

HO
O
O
OH
N
H
OH
O
1-*cis*
(synthetic)
500 MHz, COSY in $D_2O$
+0.01% MeOH
f2 (ppm)
f1 (ppm)

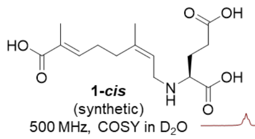


**1-*cis***
(synthetic)
500 MHz, COSY in $D_2O$
+0.01% MeOH

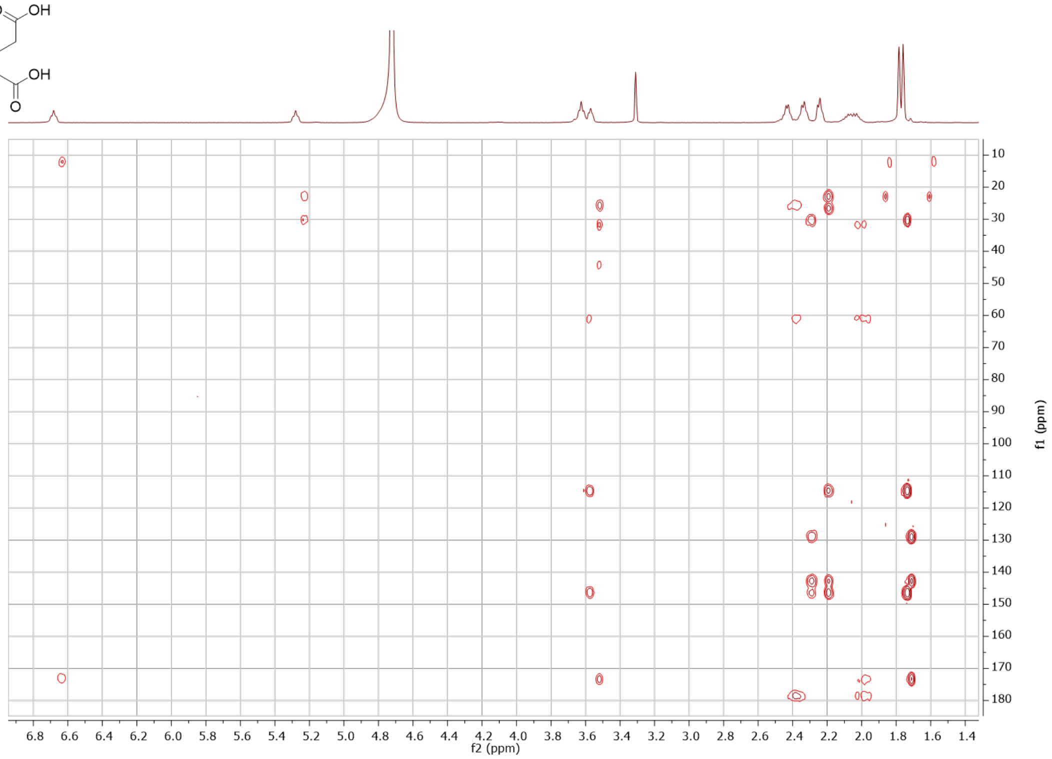

isodomoic acid A – enzymatic (**2**)

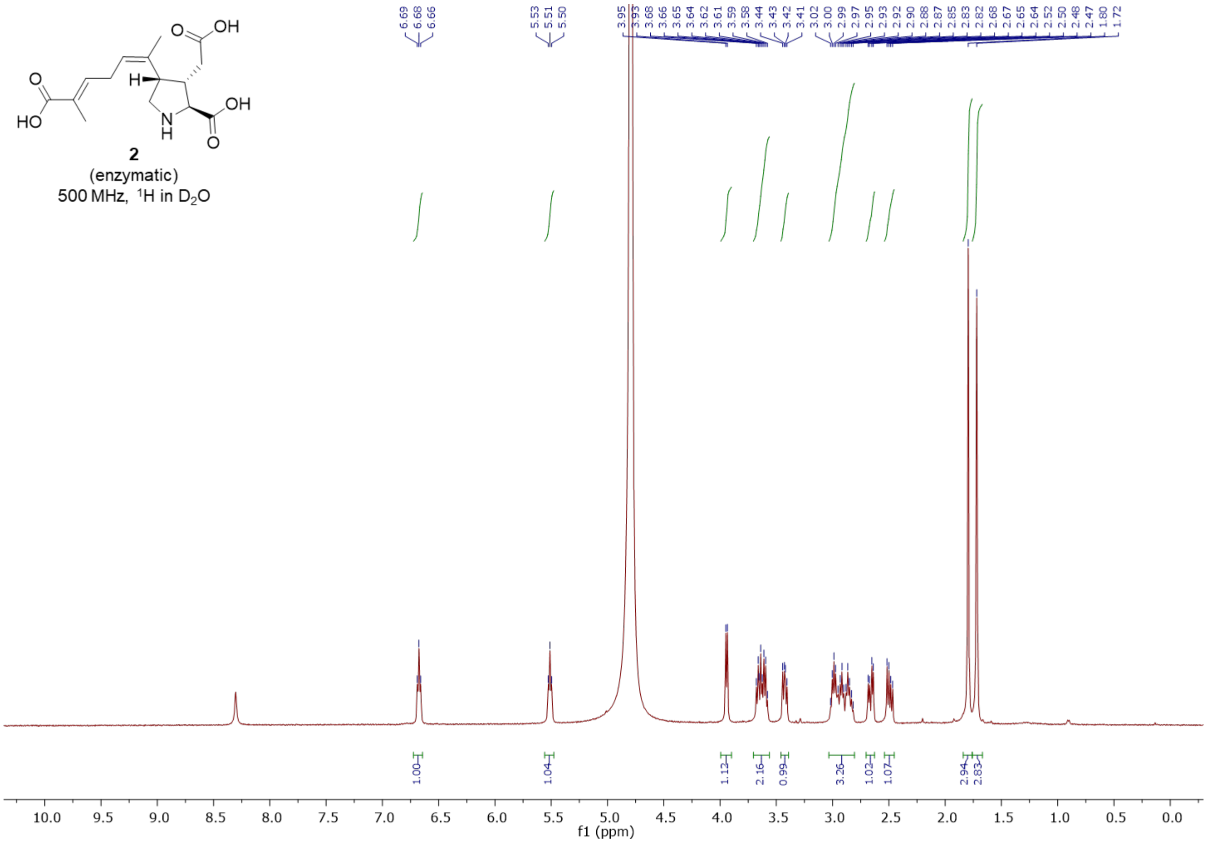


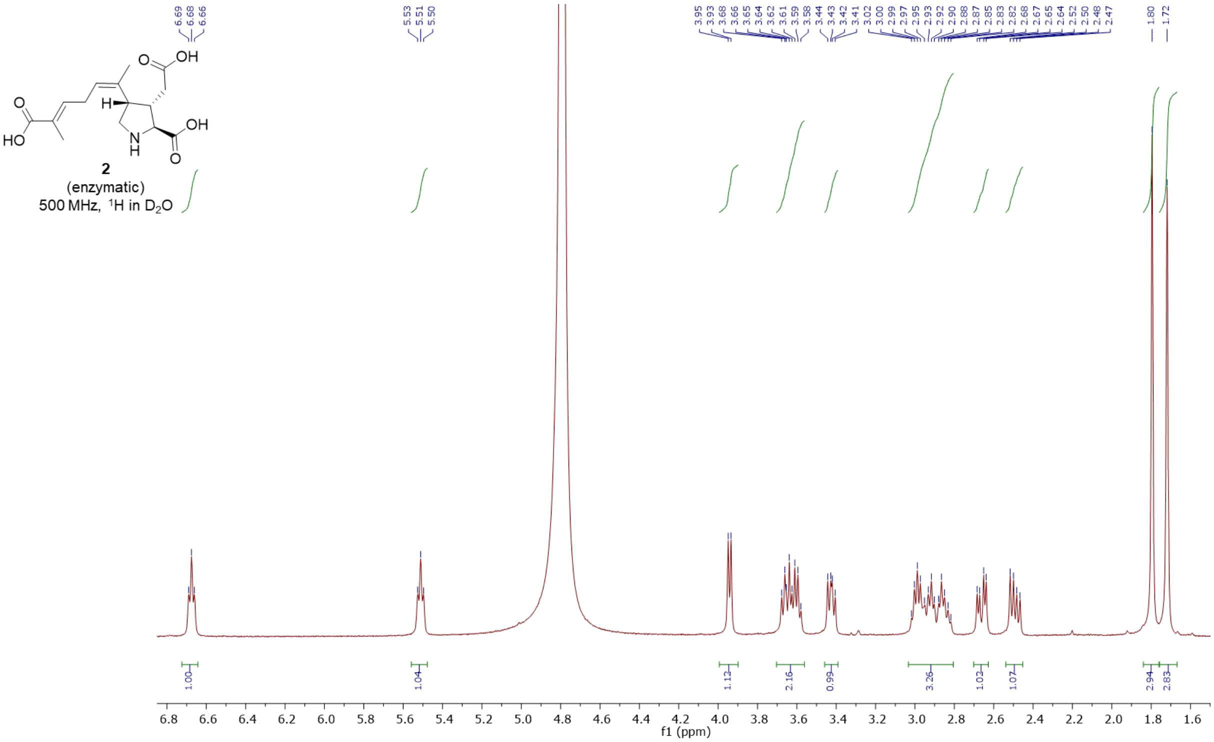

7'-carboxy-*N*-neryl-(3*R*)-hydroxy-L-glutamic acid – synthetic (**3**)

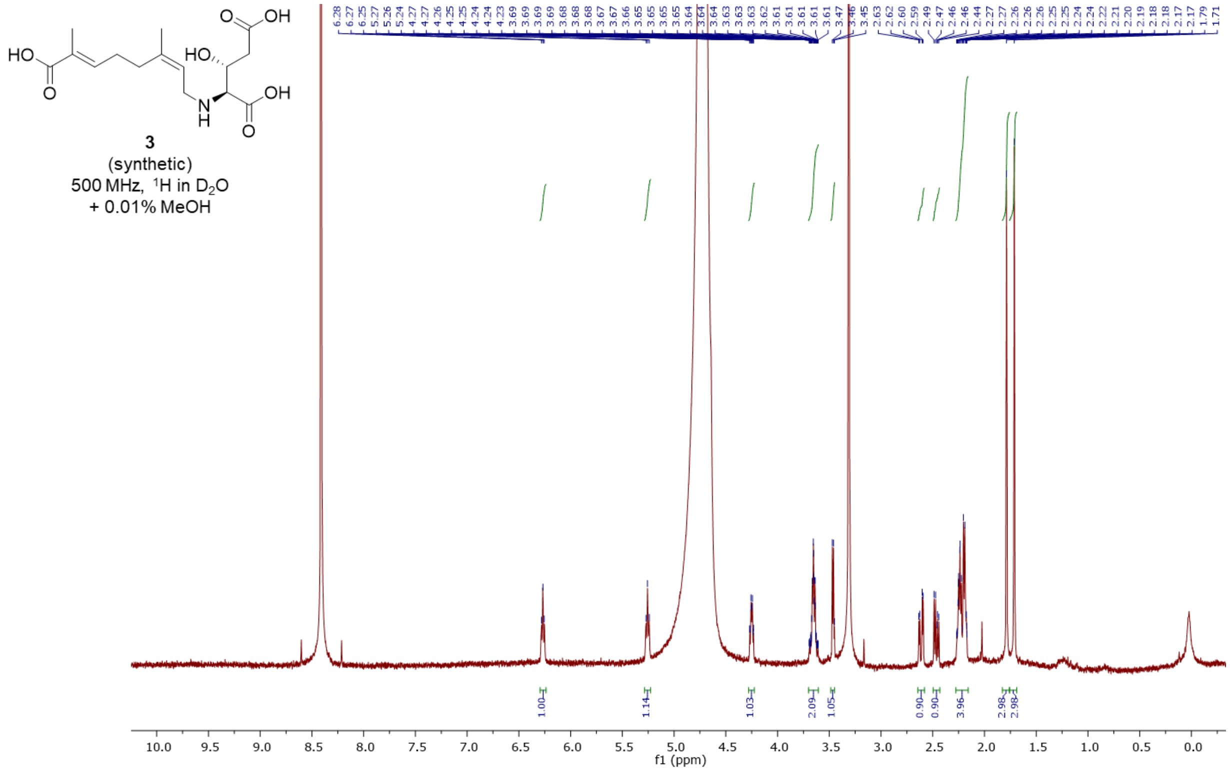


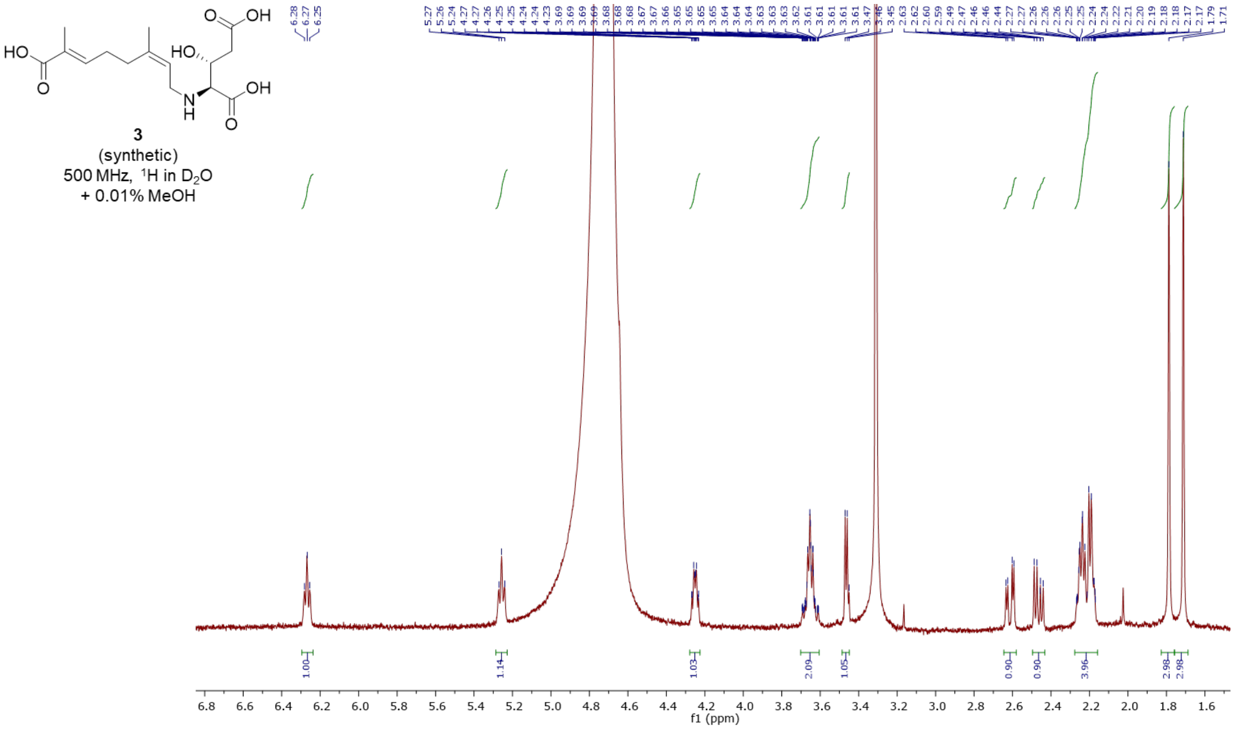

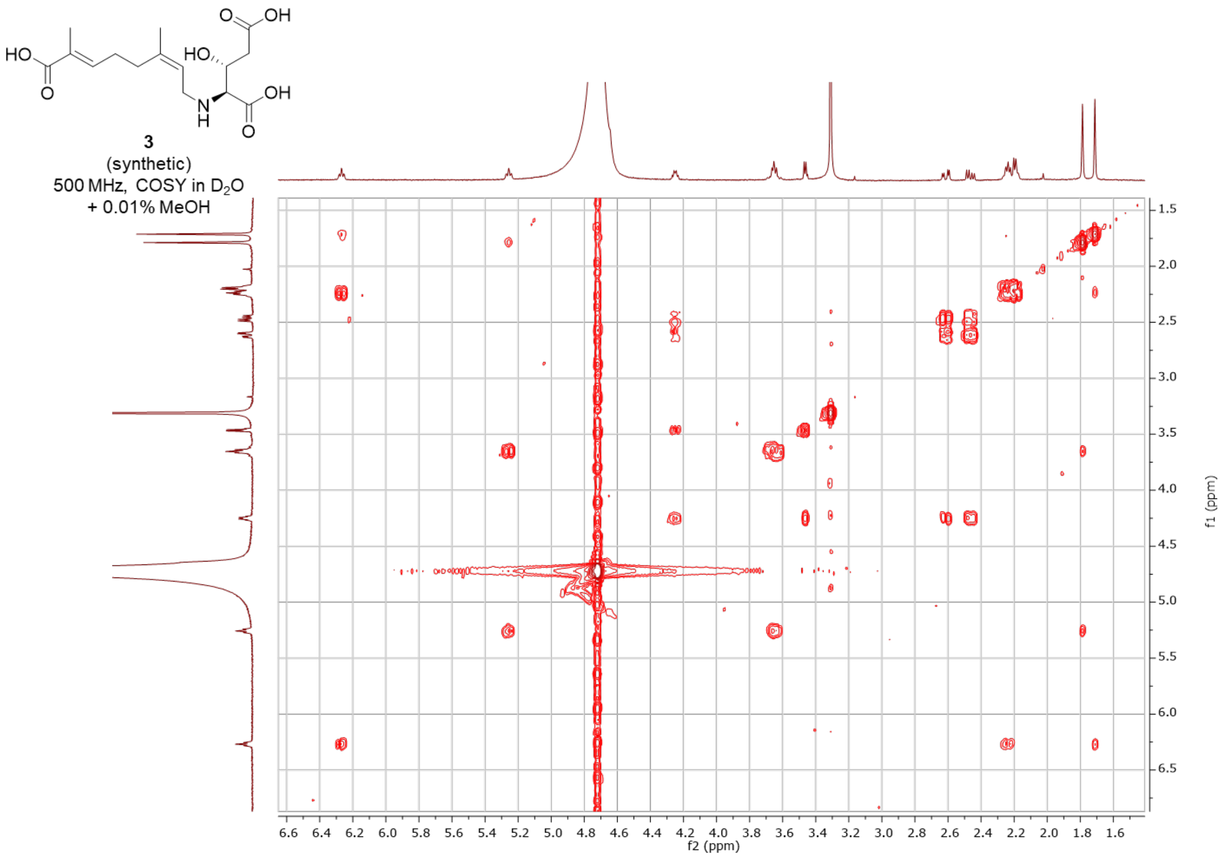


3
(synthetic)
500 MHz, COSY in $D_2O$
+ 0.01% MeOH

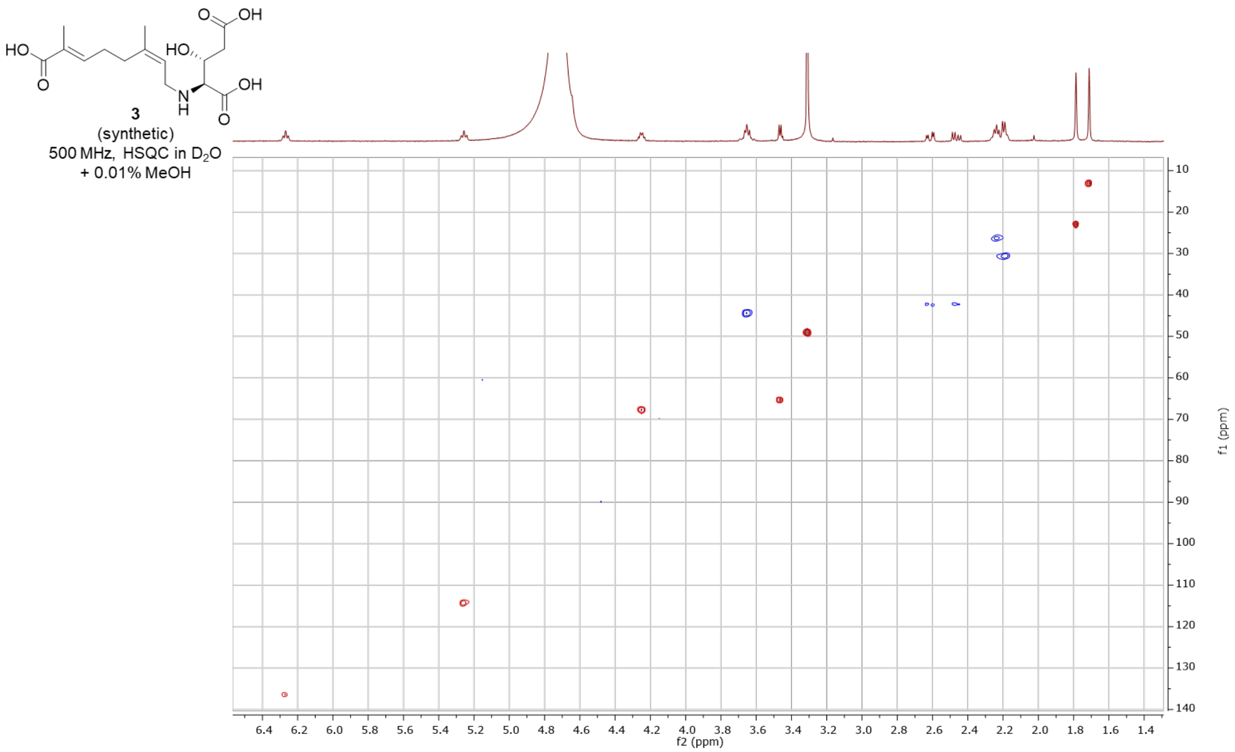


3
(synthetic)
500 MHz, HSQC in $D_2O$
+ 0.01% MeOH

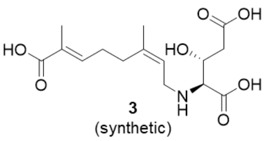


**3**
(synthetic)
500 MHz, HMBC in $D_2O$
+ 0.01% MeOH

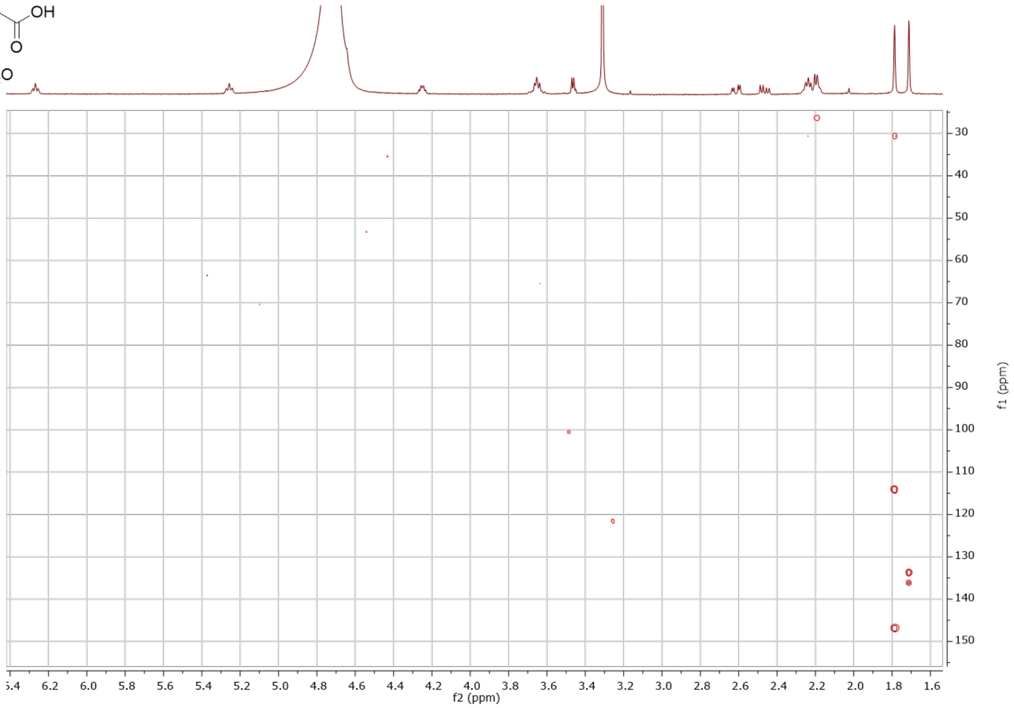

7’-carboxy-*N*-neryl-(3*R*)-hydroxy-L-glutamic acid – enzymatic (**3**)

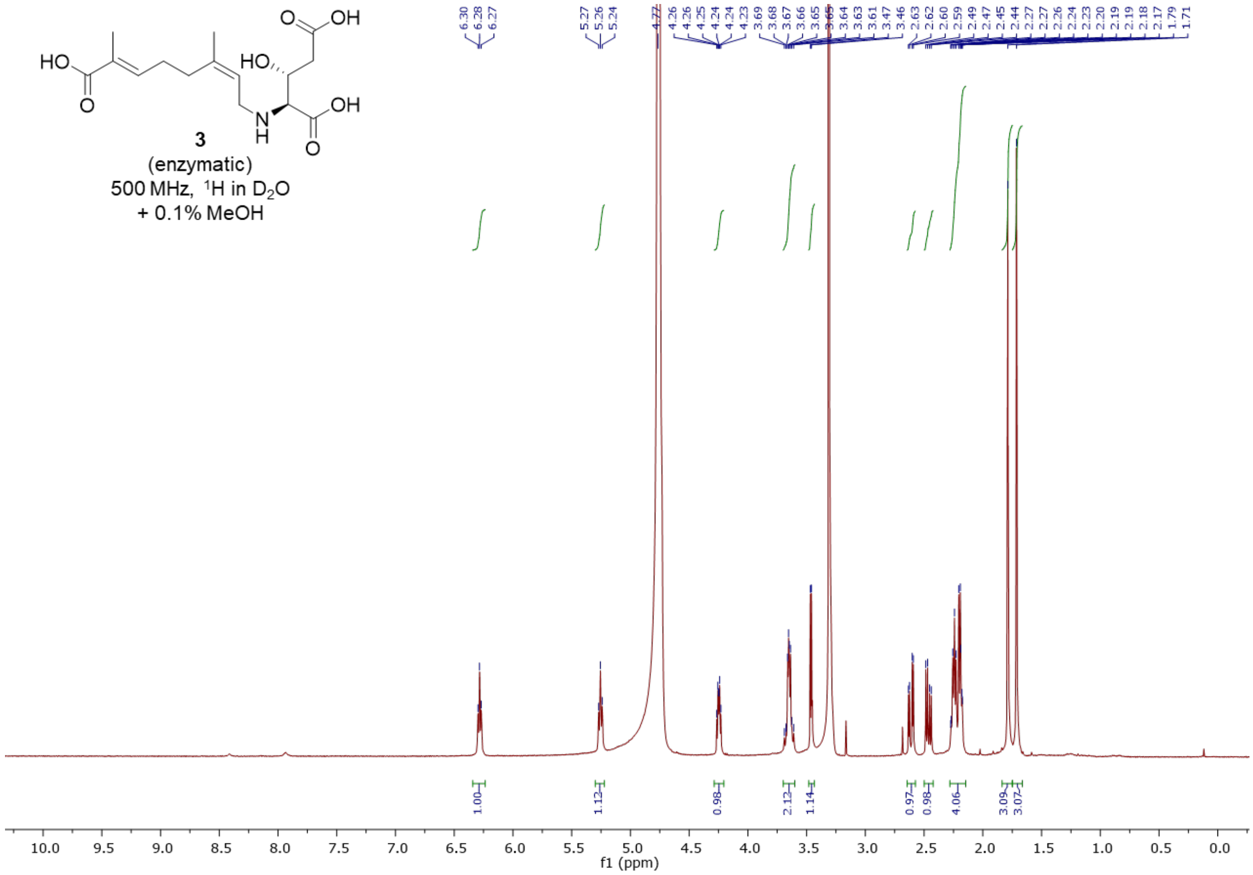


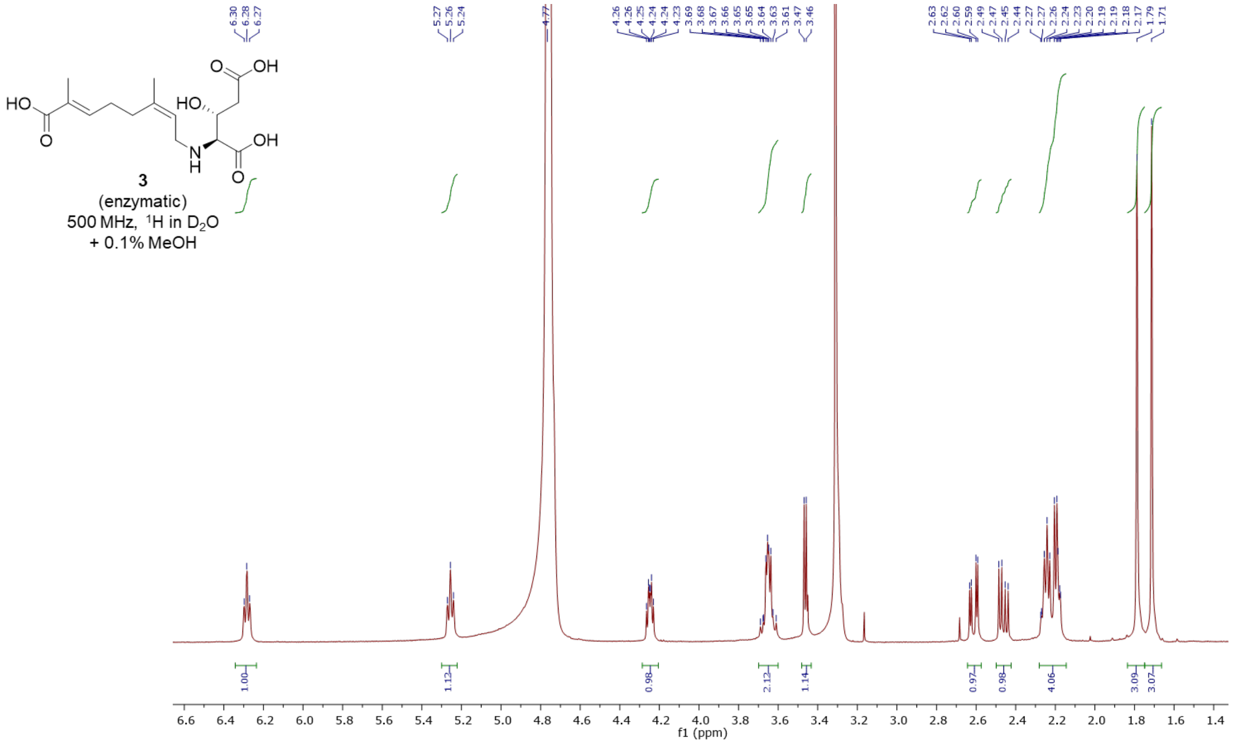

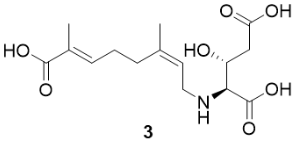


**3**
(enzymatic)
500 MHz, COSY in $D_2O$
+ 0.1% MeOH

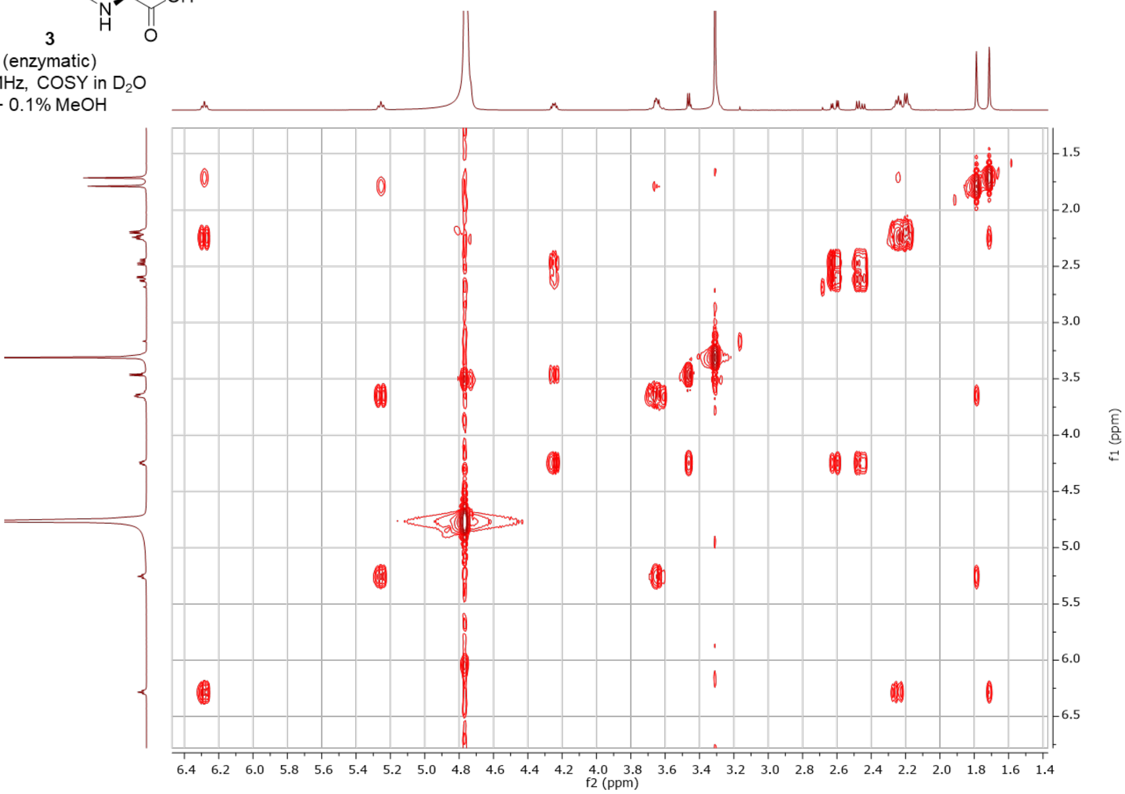


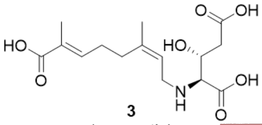


**3**
(enzymatic)
500 MHz, HSQC in $D_2O$
+ 0.1% MeOH

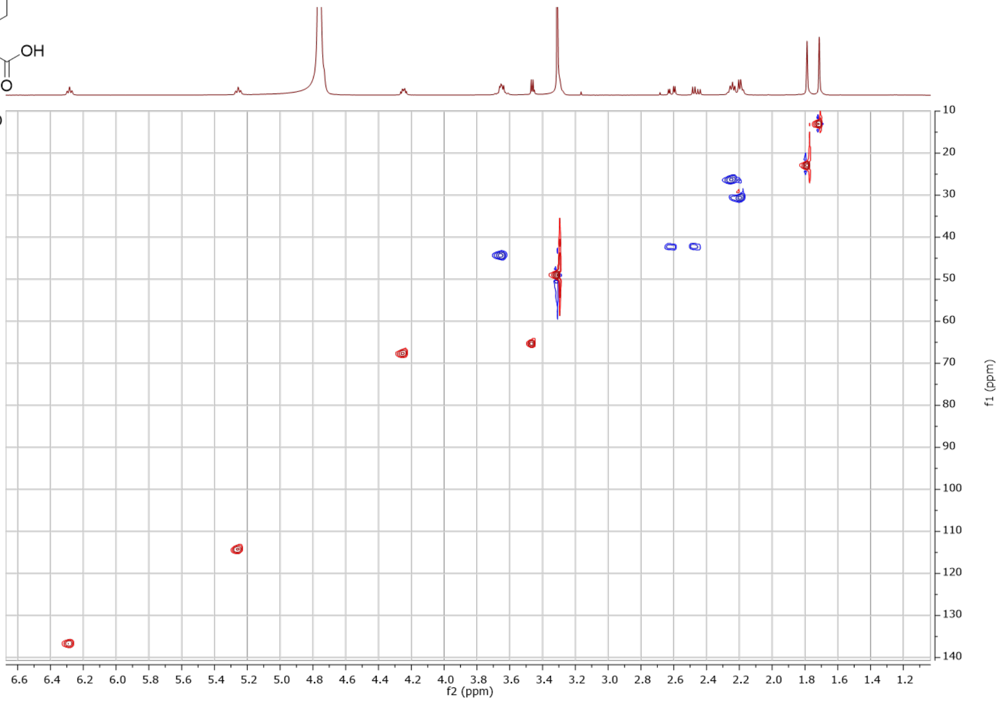

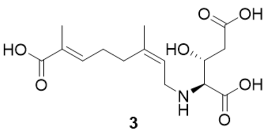


**3**
(enzymatic)
500 MHz, HMBC in $D_2O$
+ 0.1% MeOH

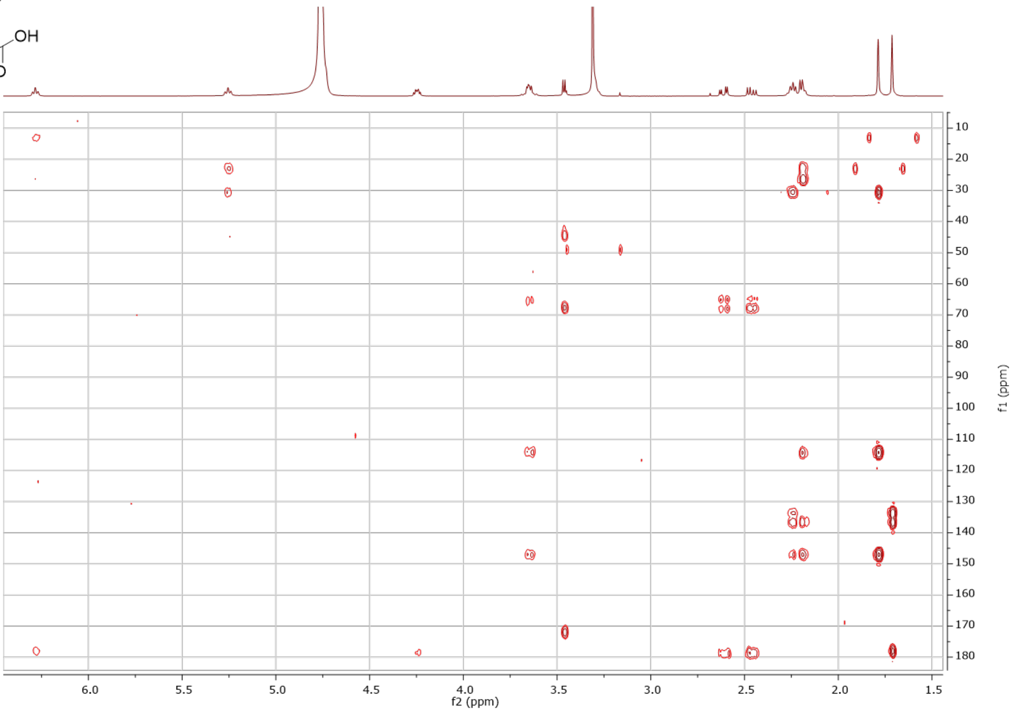


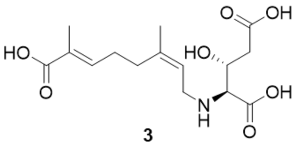


**3**
(enzymatic)
500 MHz, NOESY in $D_2O$
+ 0.1% MeOH

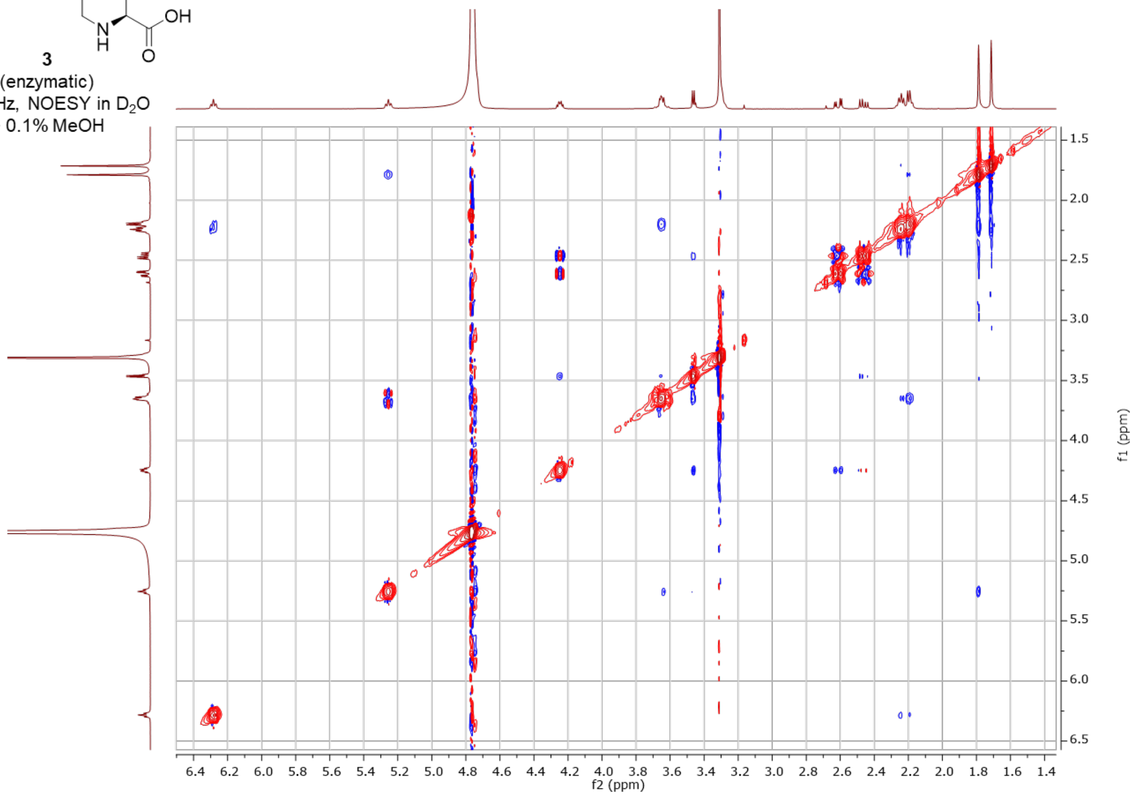

7'-hydroxy-L-NGG - synthetic (***4-trans***)

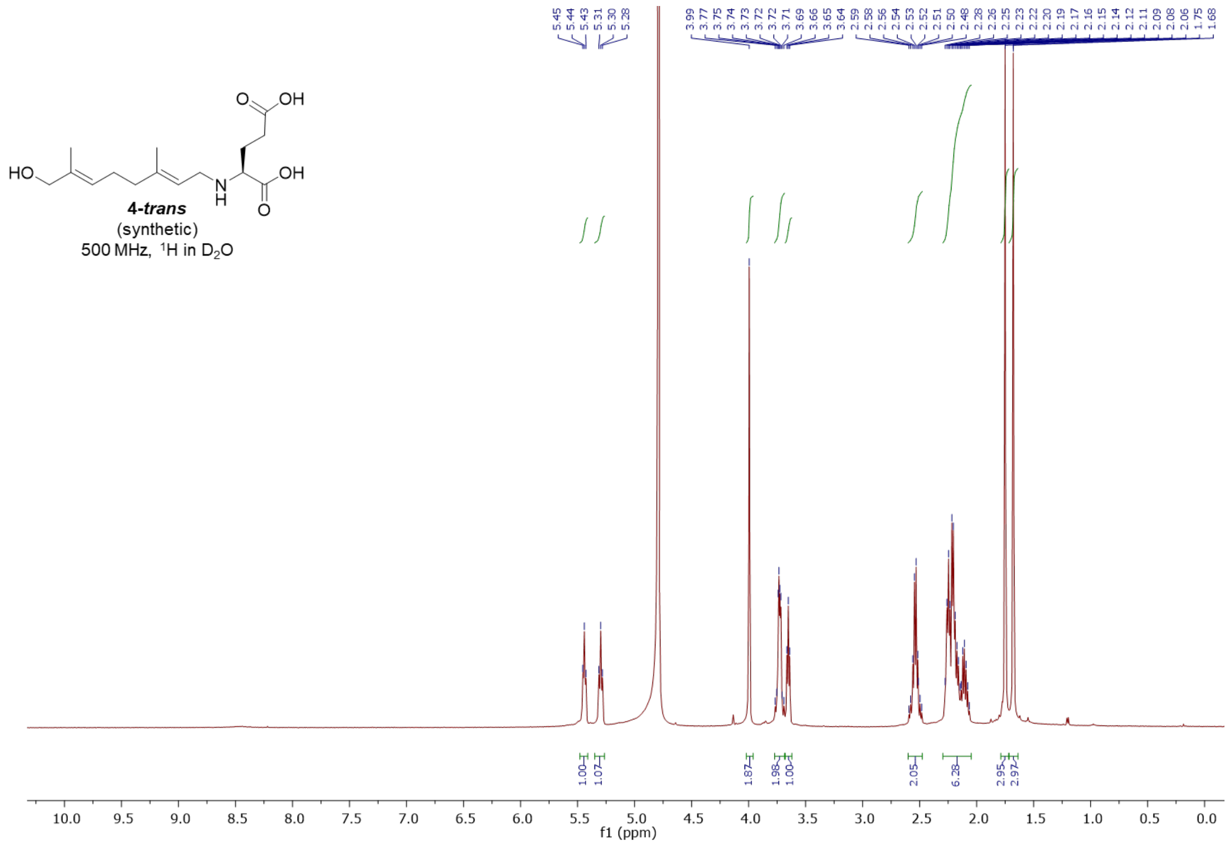


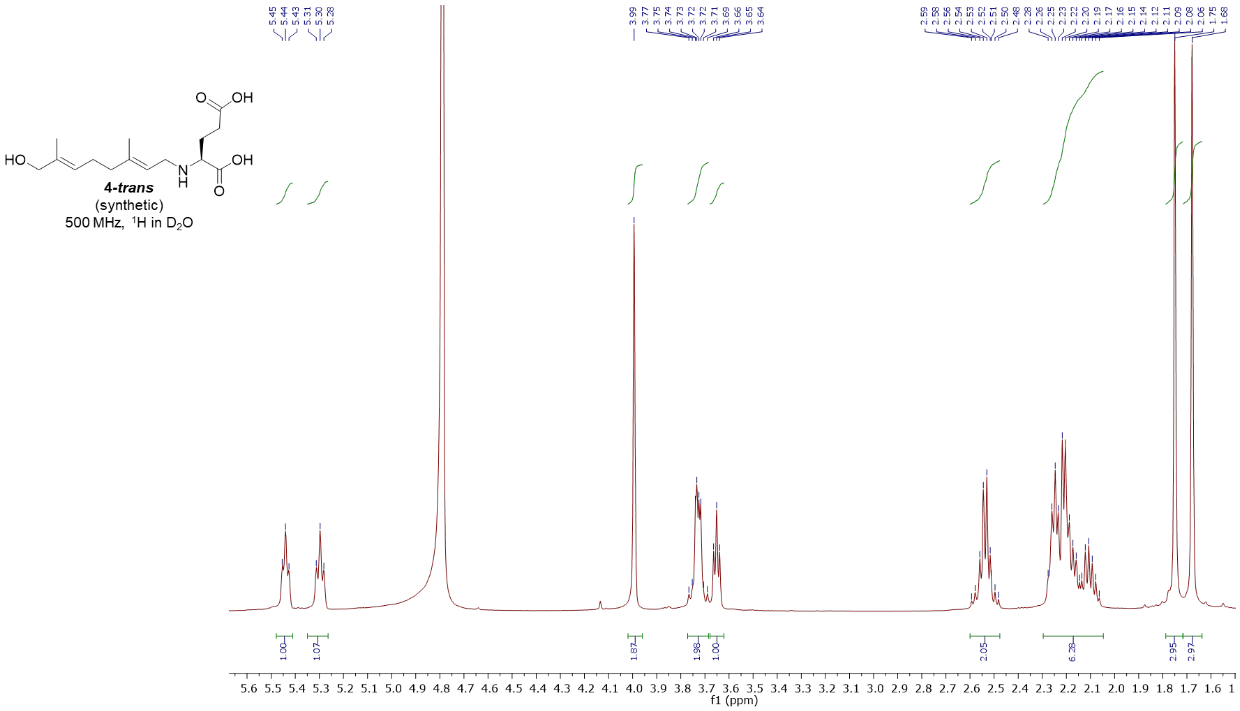

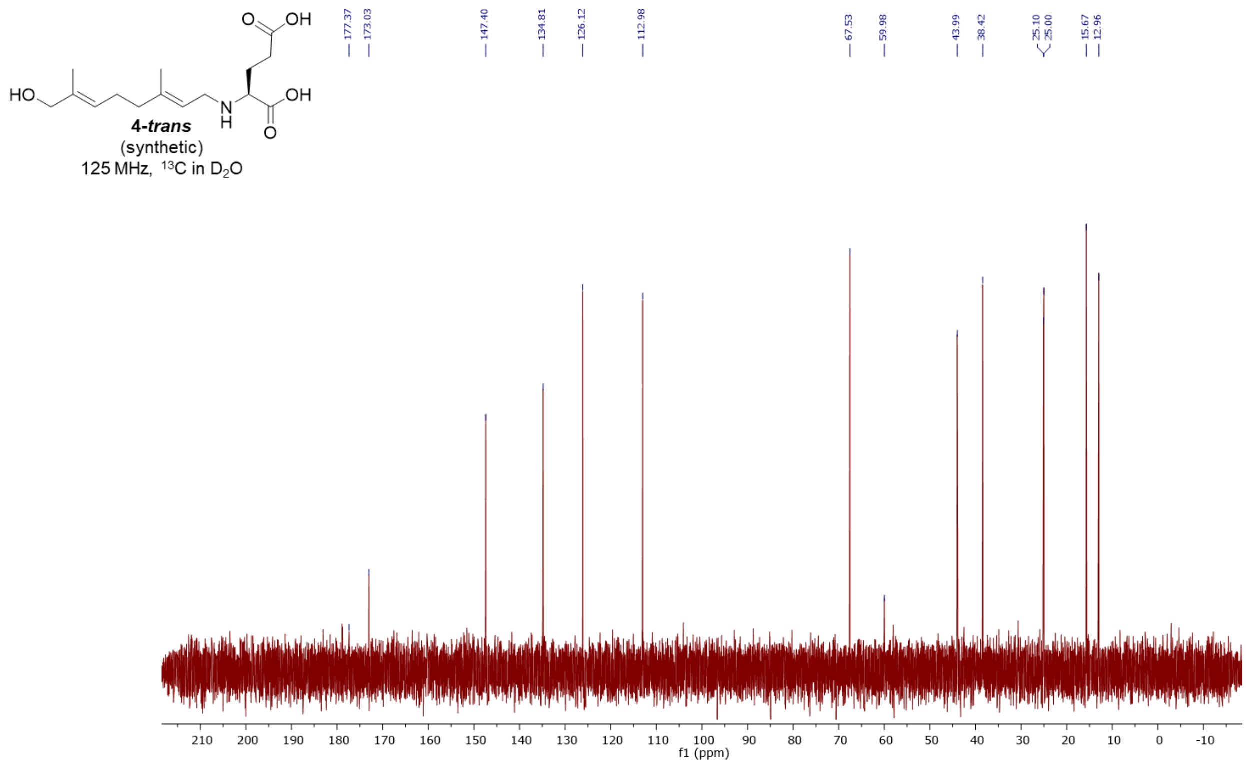
4-trans
(synthetic)
125 MHz, 13C in D2O
177.37
173.03
147.40
134.81
126.12
112.98
67.53
59.98
43.99
38.42
25.10
25.00
15.67
12.96
f1 (ppm)

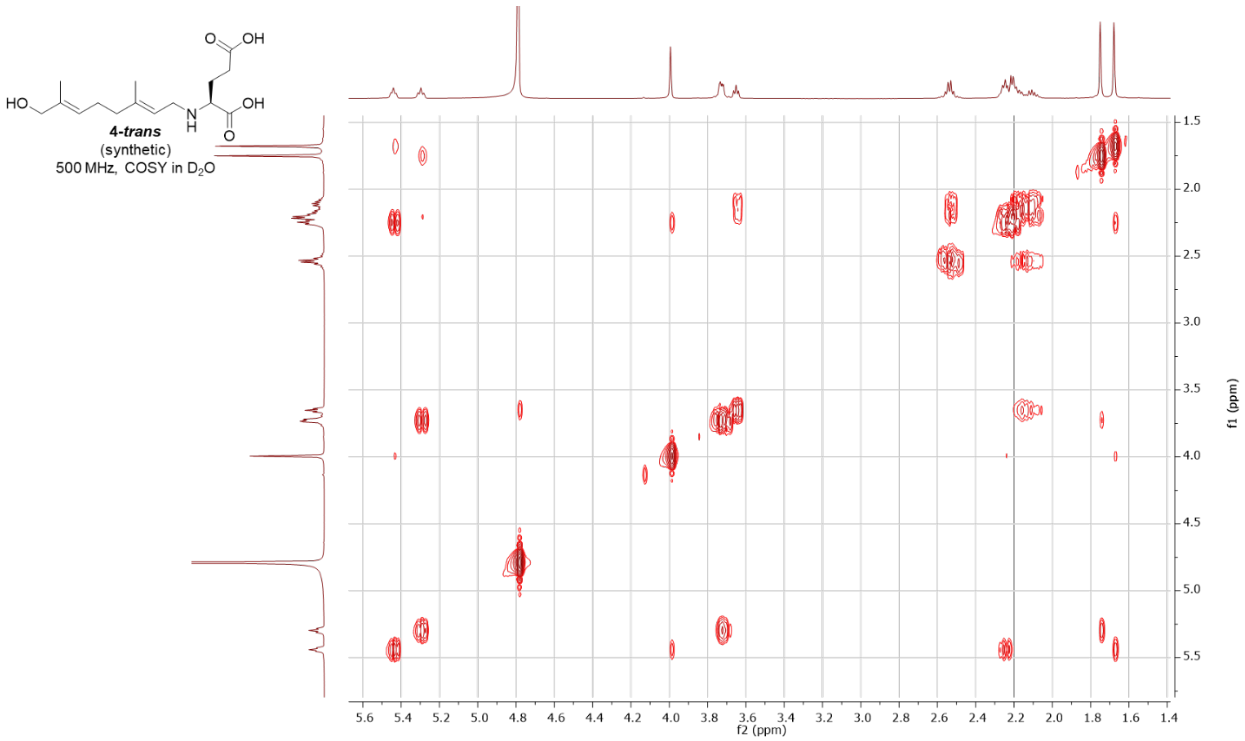
4-trans
(synthetic)
500 MHz, COSY in D2O
f2 (ppm)
f1 (ppm)

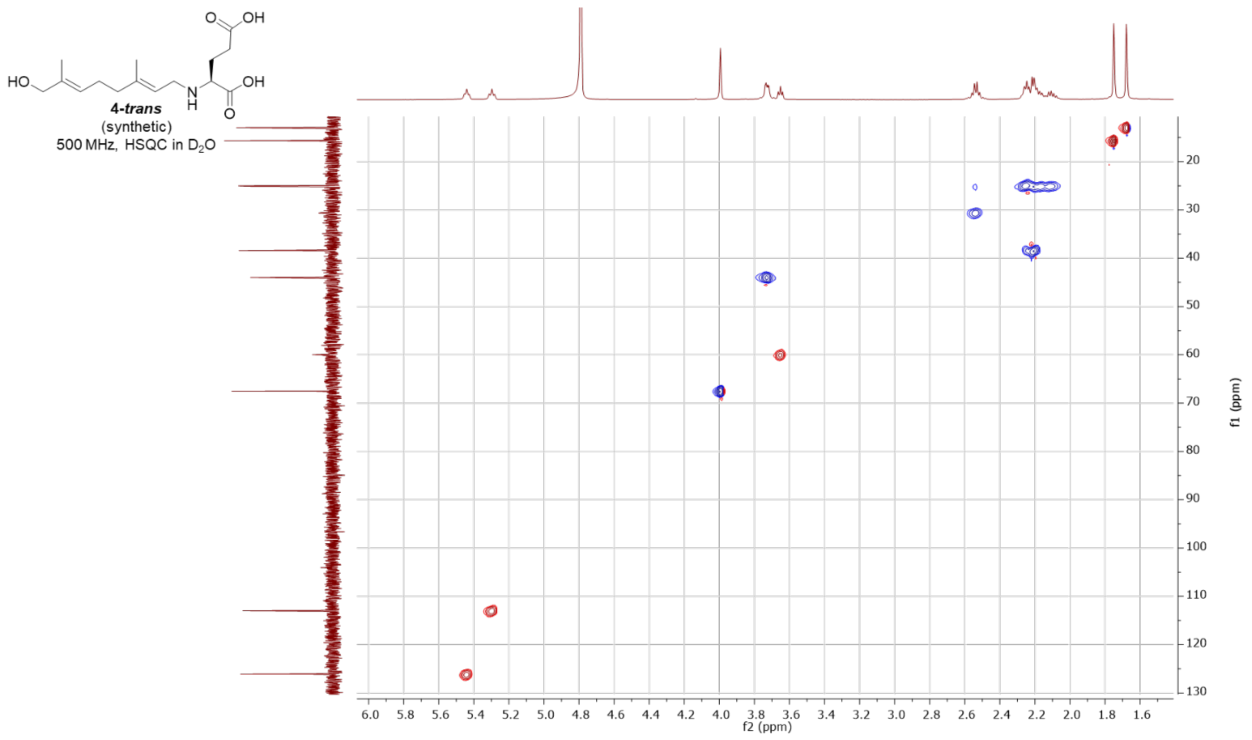
4-trans
(synthetic)
500 MHz, HSQC in $D_2O$
f2 (ppm)
f1 (ppm)

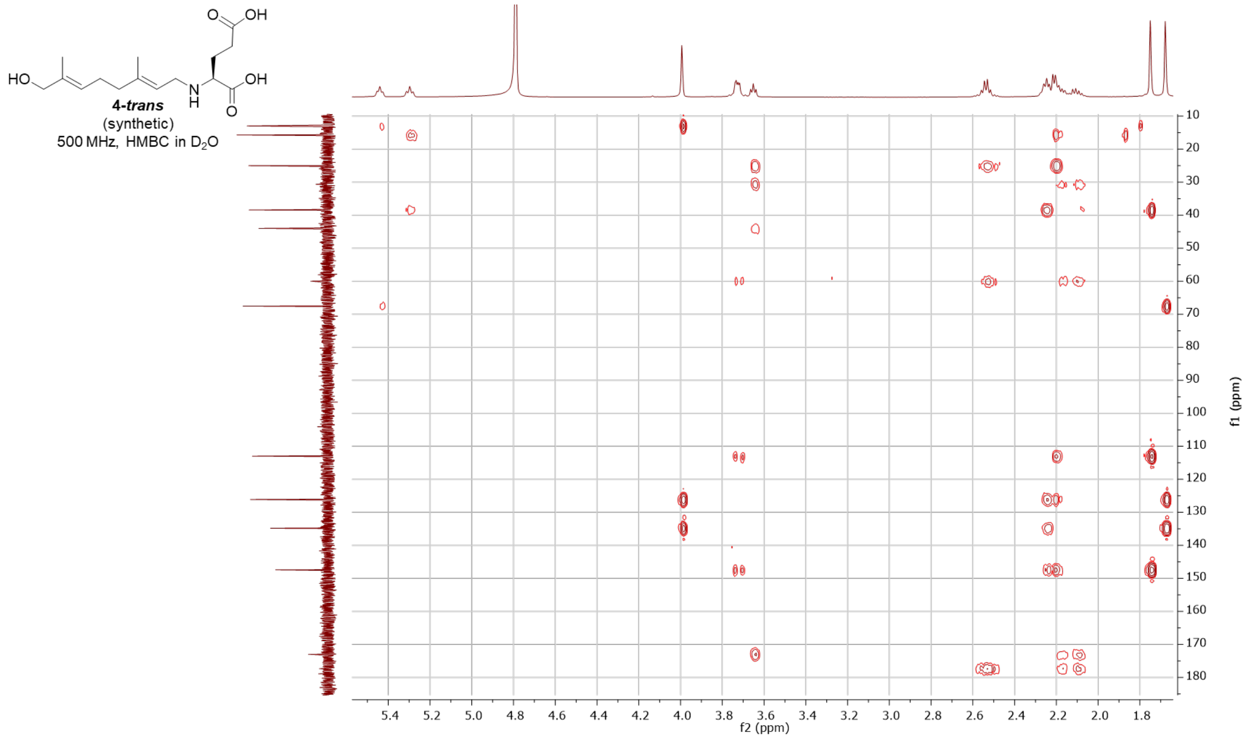
4-trans
(synthetic)
500 MHz, HMBC in $D_2O$
f2 (ppm)
f1 (ppm)

7’-hydroxy-L-NNG- synthetic (**4-*cis***)

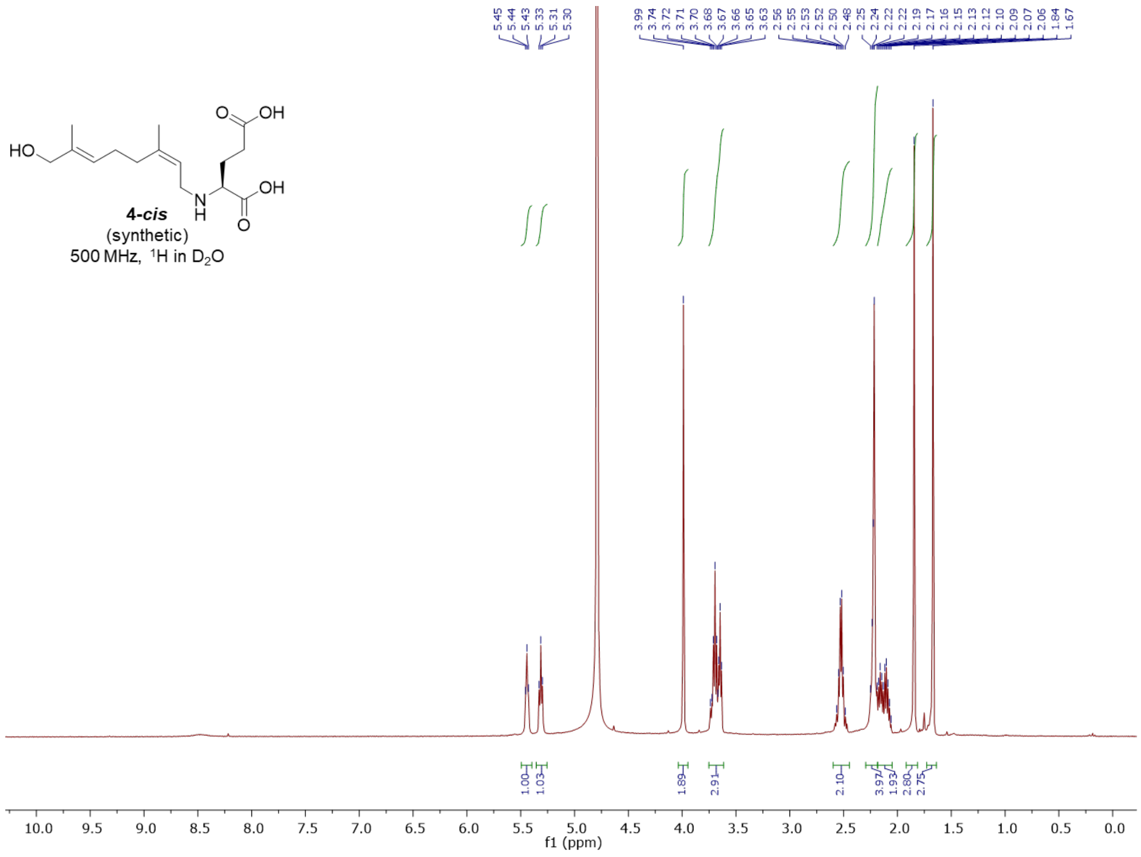


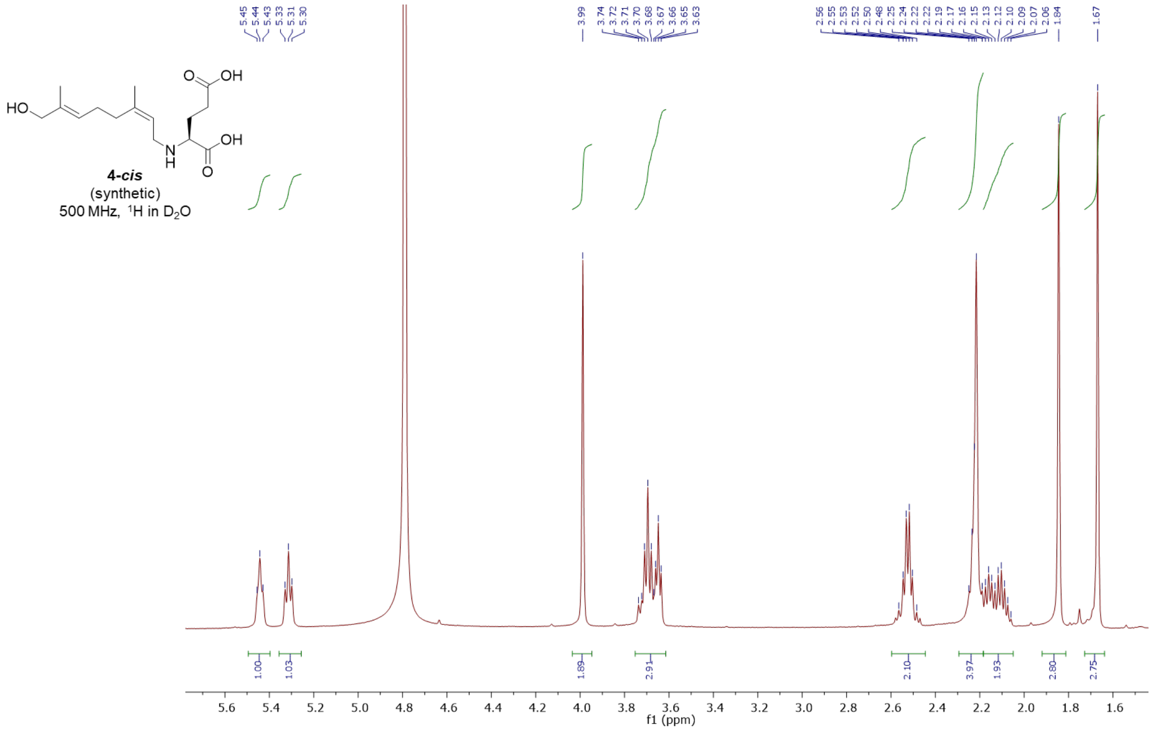

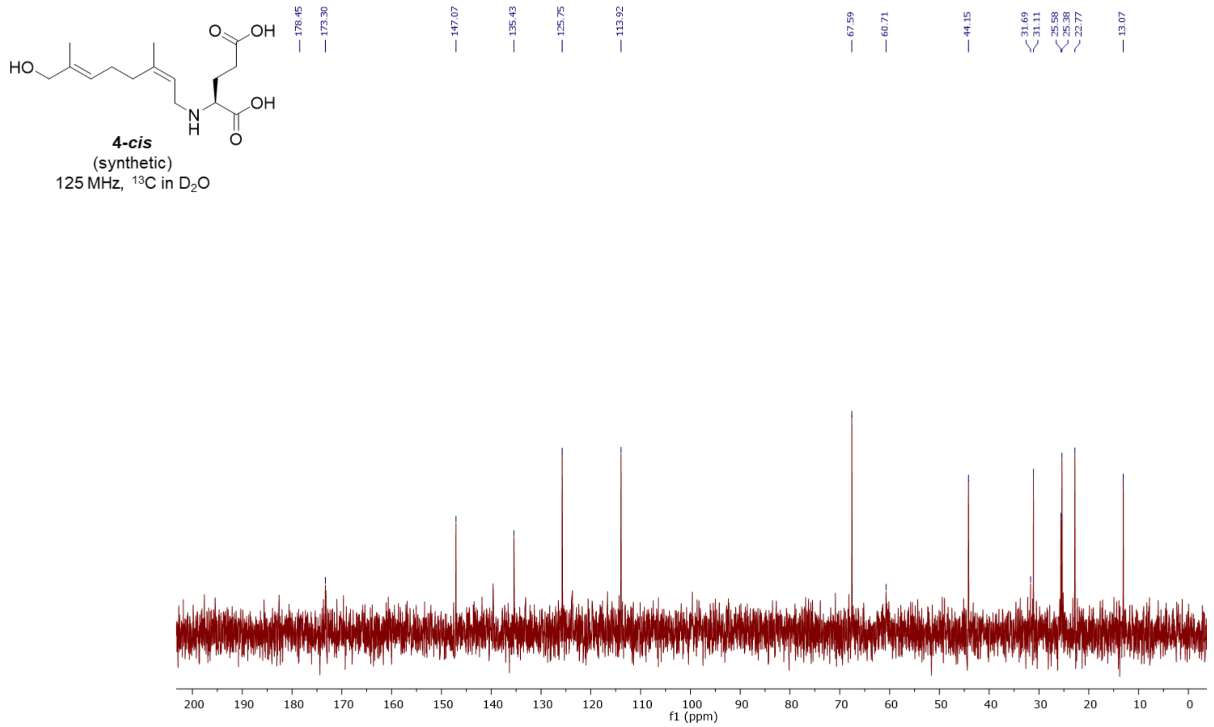
HO
O
OH
N
H
OH
O
4-cis
(synthetic)
125 MHz, $^{13}C$ in $D_2O$
178.45
173.30
147.07
135.43
125.75
113.92
67.59
60.71
44.15
31.69
31.11
25.58
25.38
22.77
13.07
f1 (ppm)

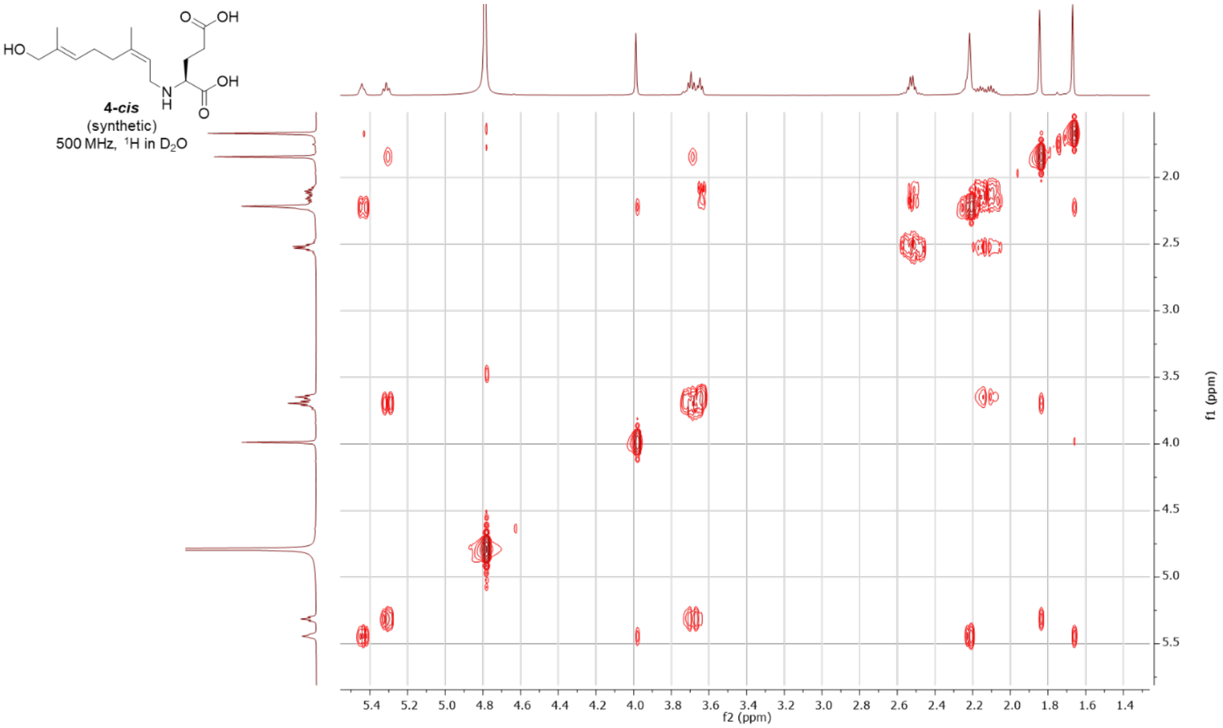
HO
O
OH
N
H
OH
O
4-cis
(synthetic)
500 MHz, $^{1}H$ in $D_2O$
f2 (ppm)
f1 (ppm)

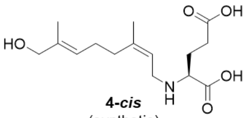

***4-cis***
(synthetic)
500 MHz, HSQC in $D_2O$

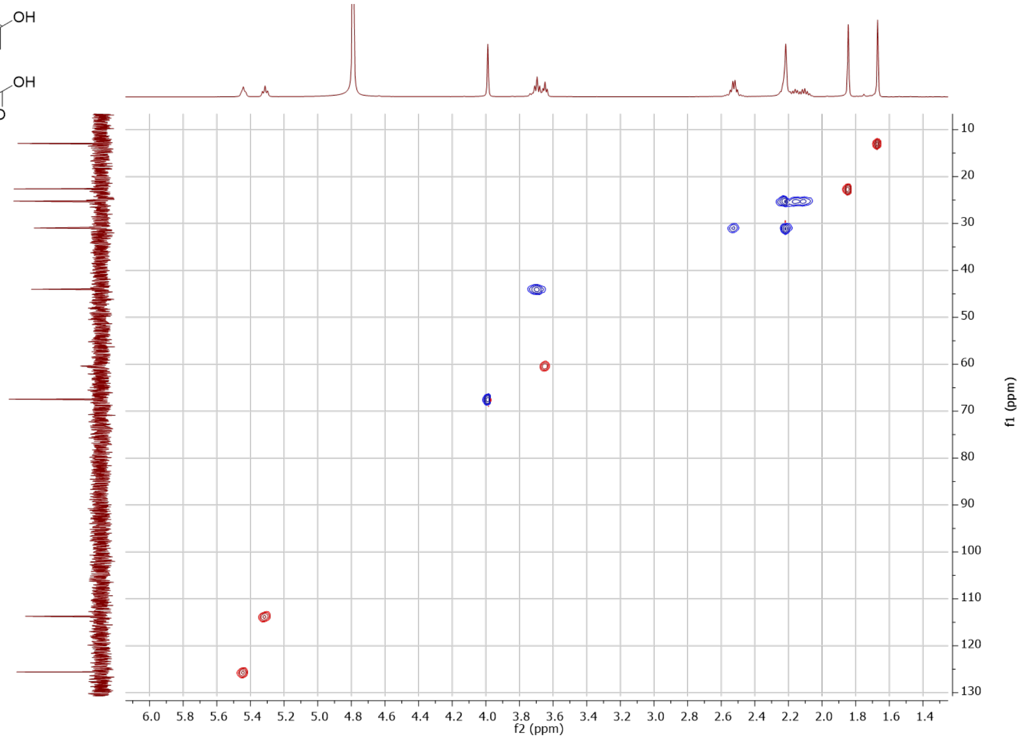


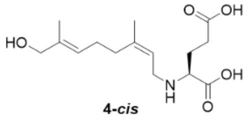

***4-cis***
(synthetic)
500 MHz, HMBC in $D_2O$

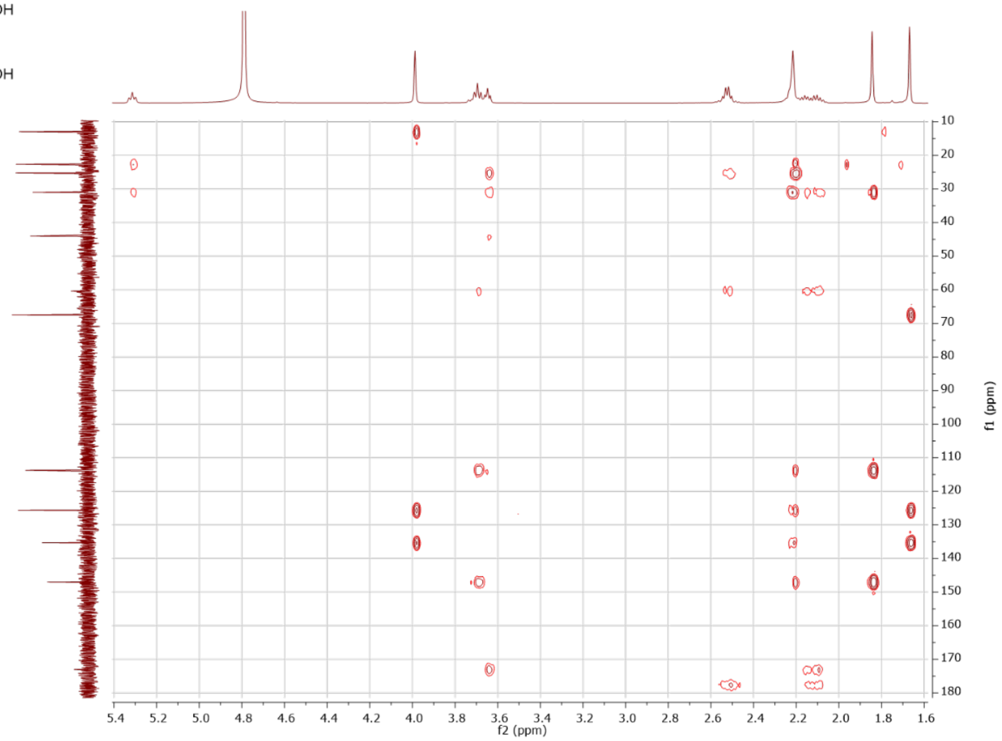

L-NGG - synthetic **(5-*trans*)**

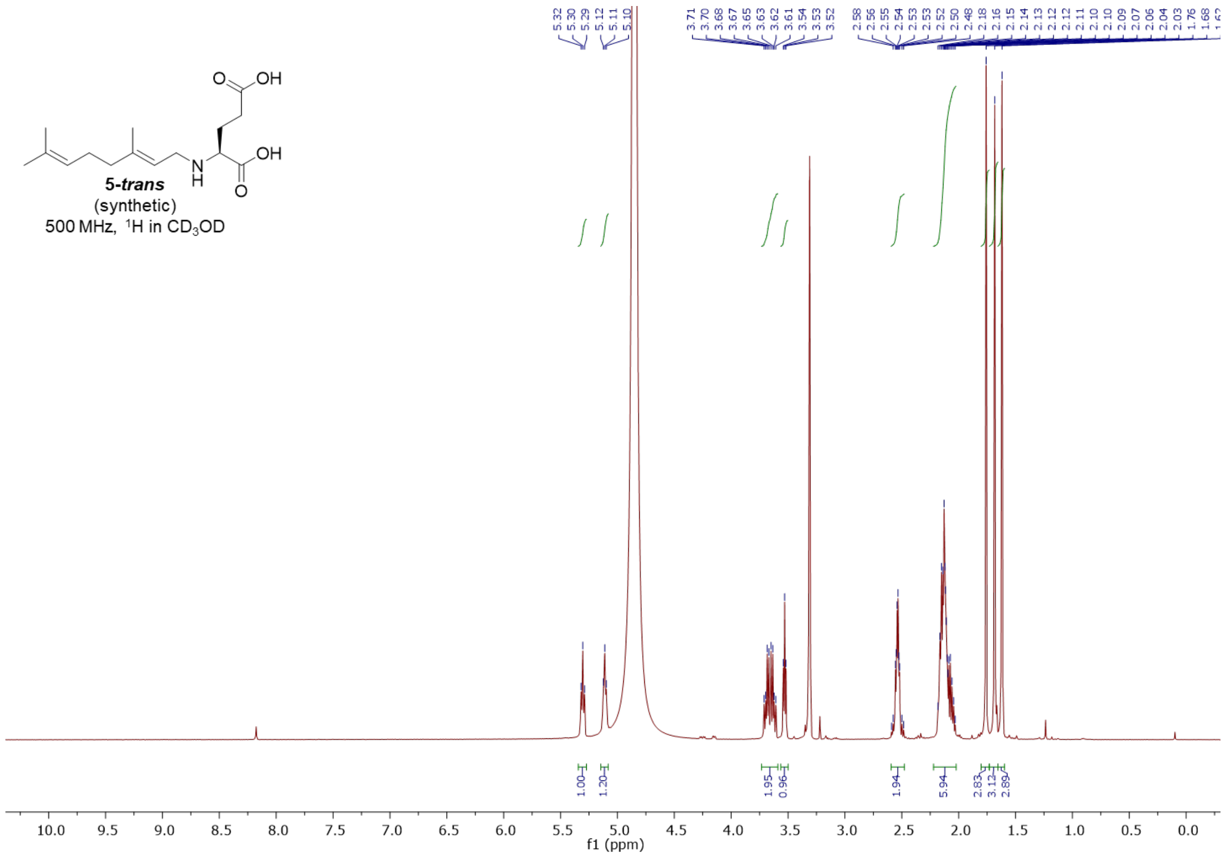


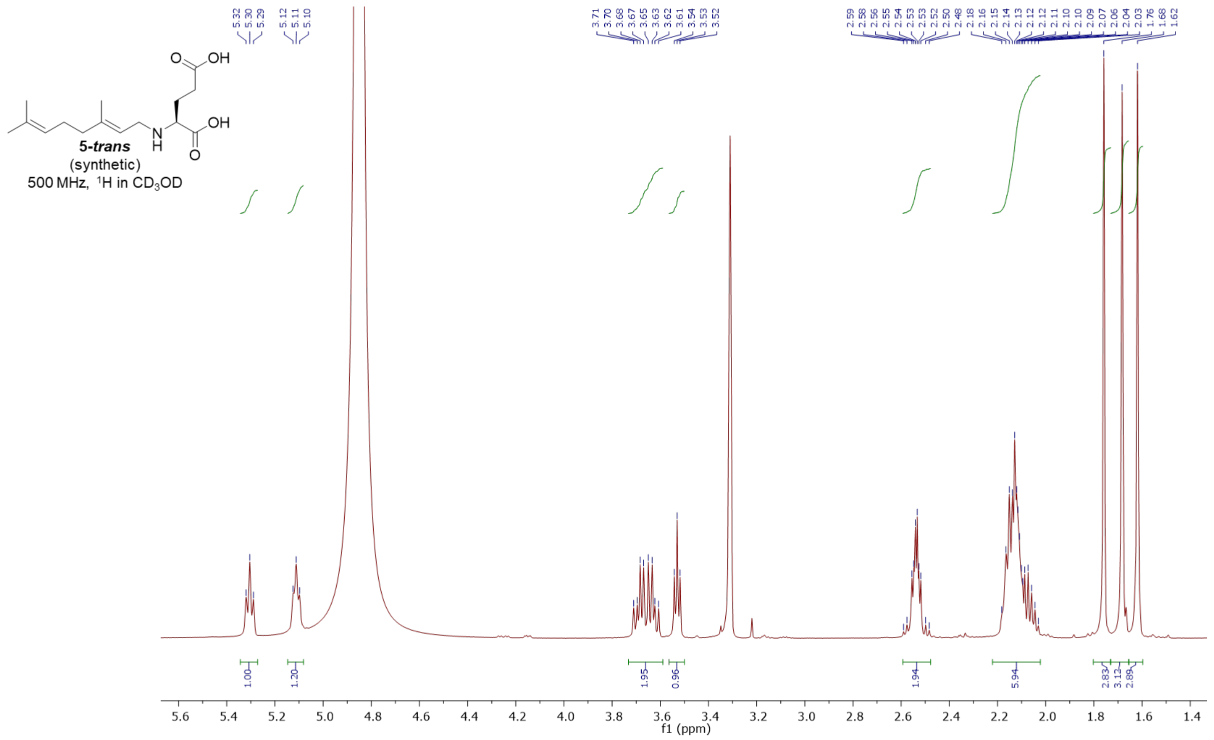

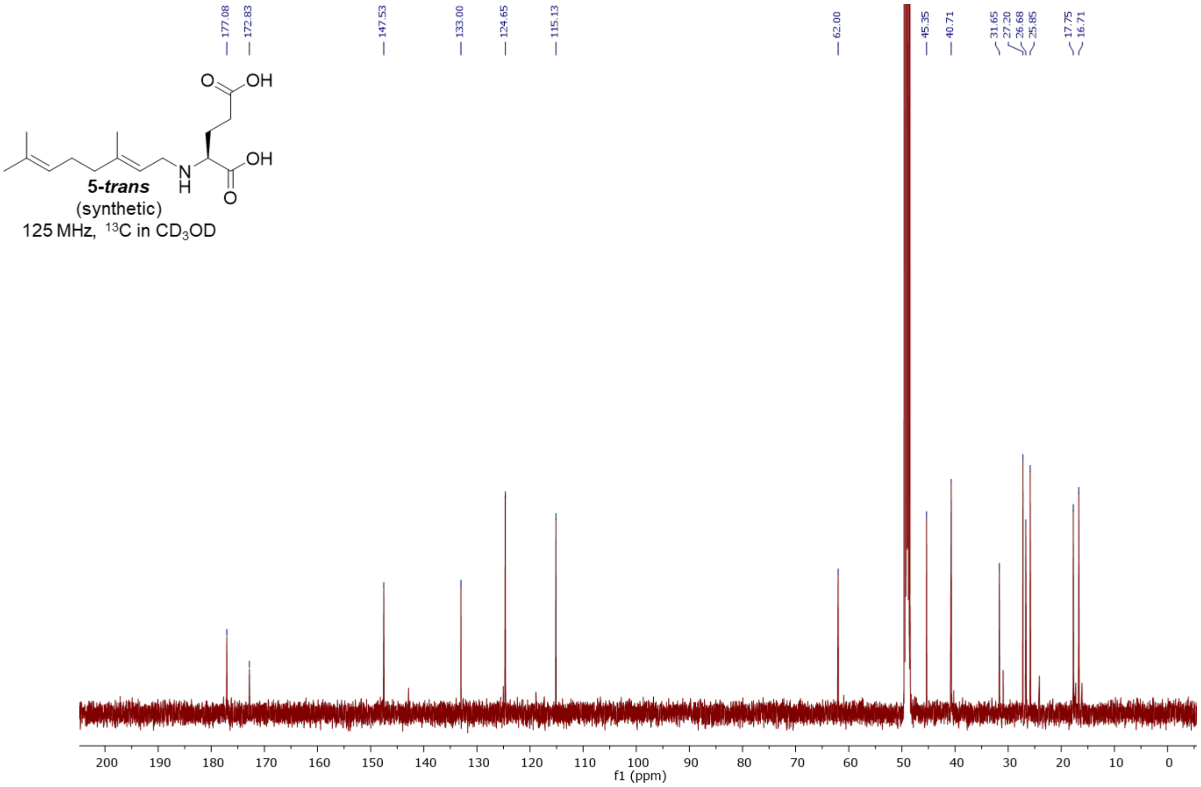
177.08
172.83
147.53
133.00
124.65
115.13
62.00
45.35
40.71
31.65
27.20
26.68
25.85
17.75
16.71
5-trans
(synthetic)
125 MHz, 13C in CD3OD
f1 (ppm)

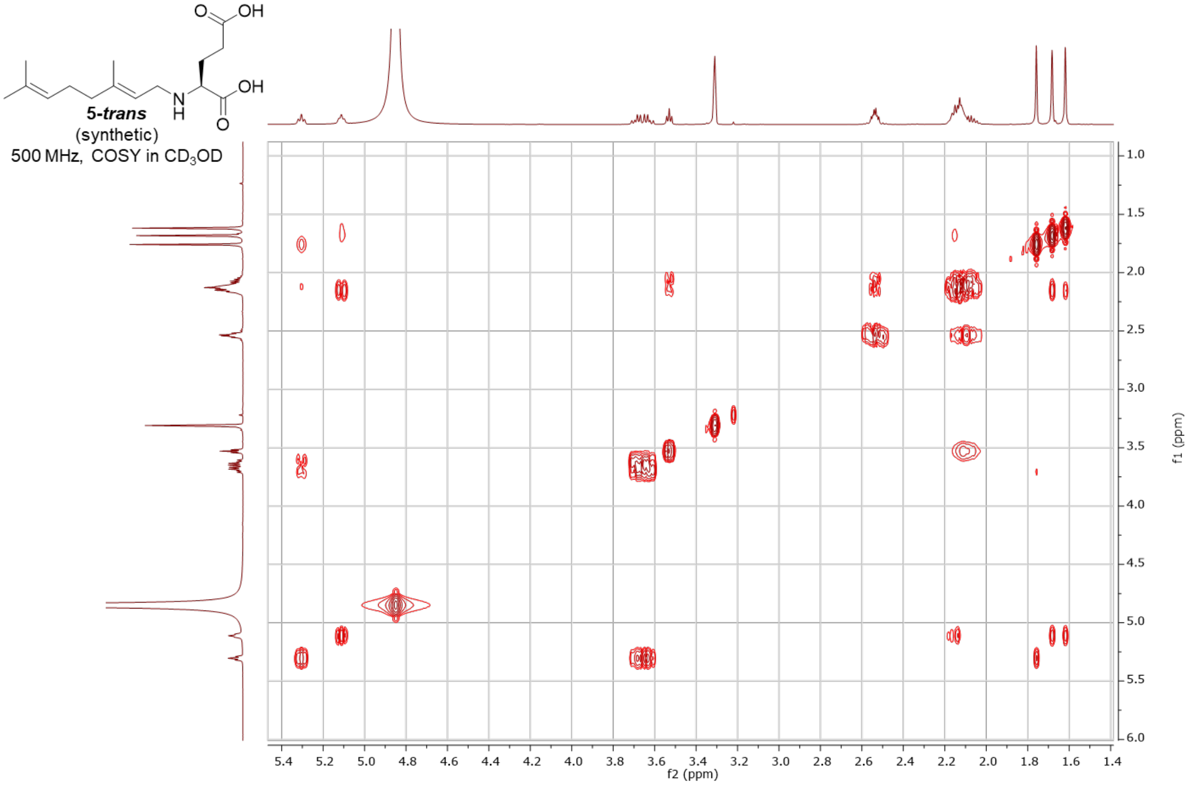
5-trans
(synthetic)
500 MHz, COSY in CD3OD
f2 (ppm)
f1 (ppm)

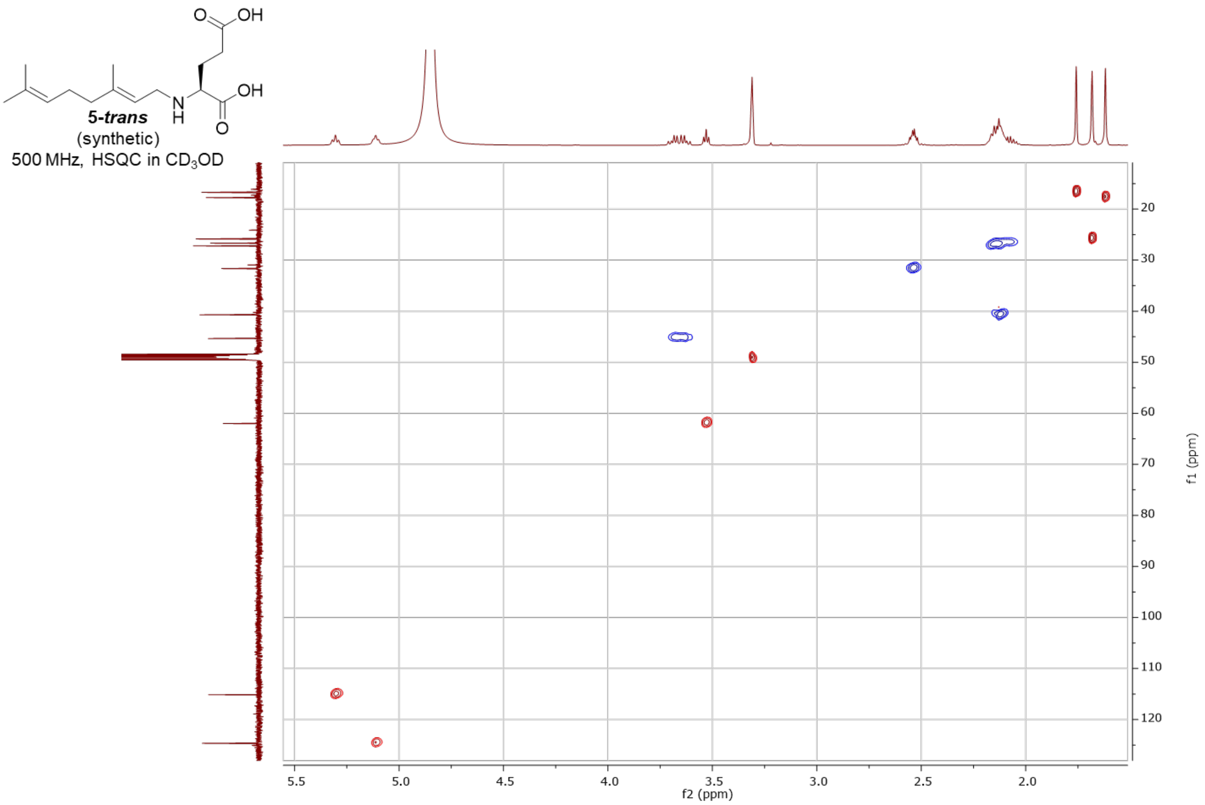
5-*trans*
(synthetic)
500 MHz, HSQC in $CD_3OD$
f2 (ppm)
f1 (ppm)

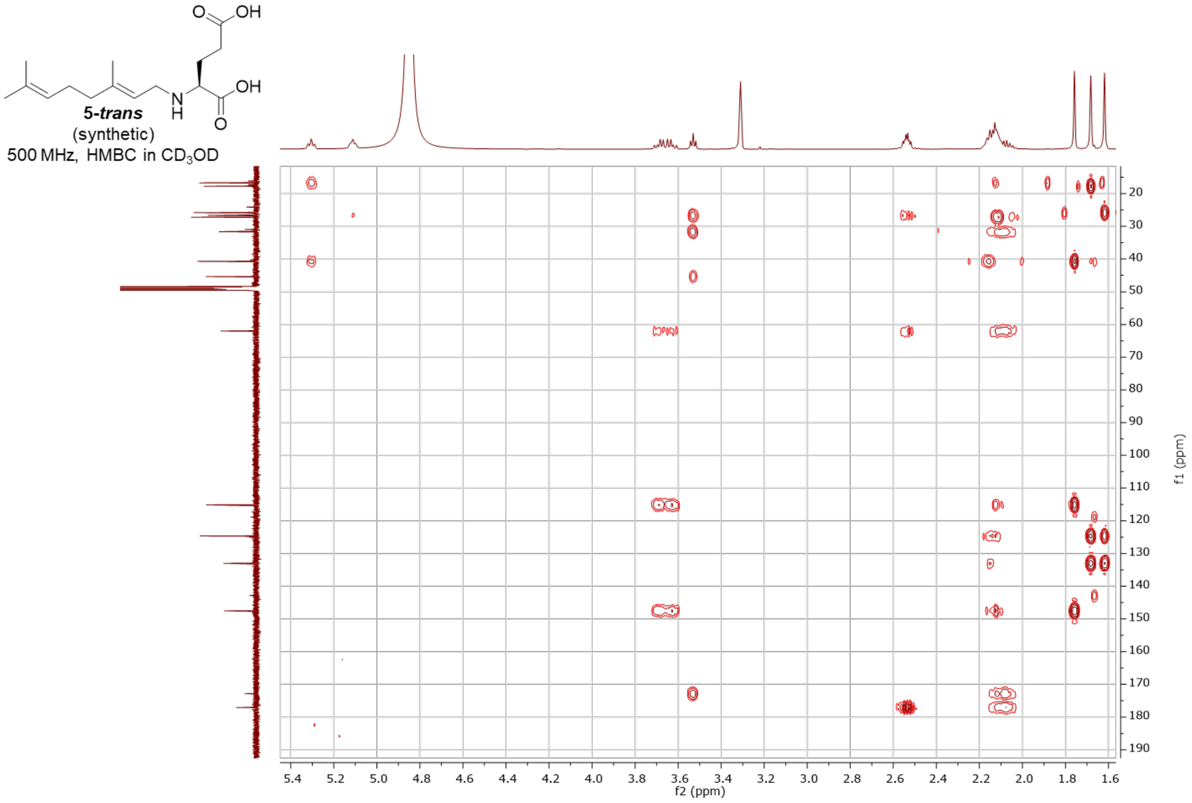
5-*trans*
(synthetic)
500 MHz, HMBC in $CD_3OD$
f2 (ppm)
f1 (ppm)

### L-NNG – synthetic (***5-cis***)

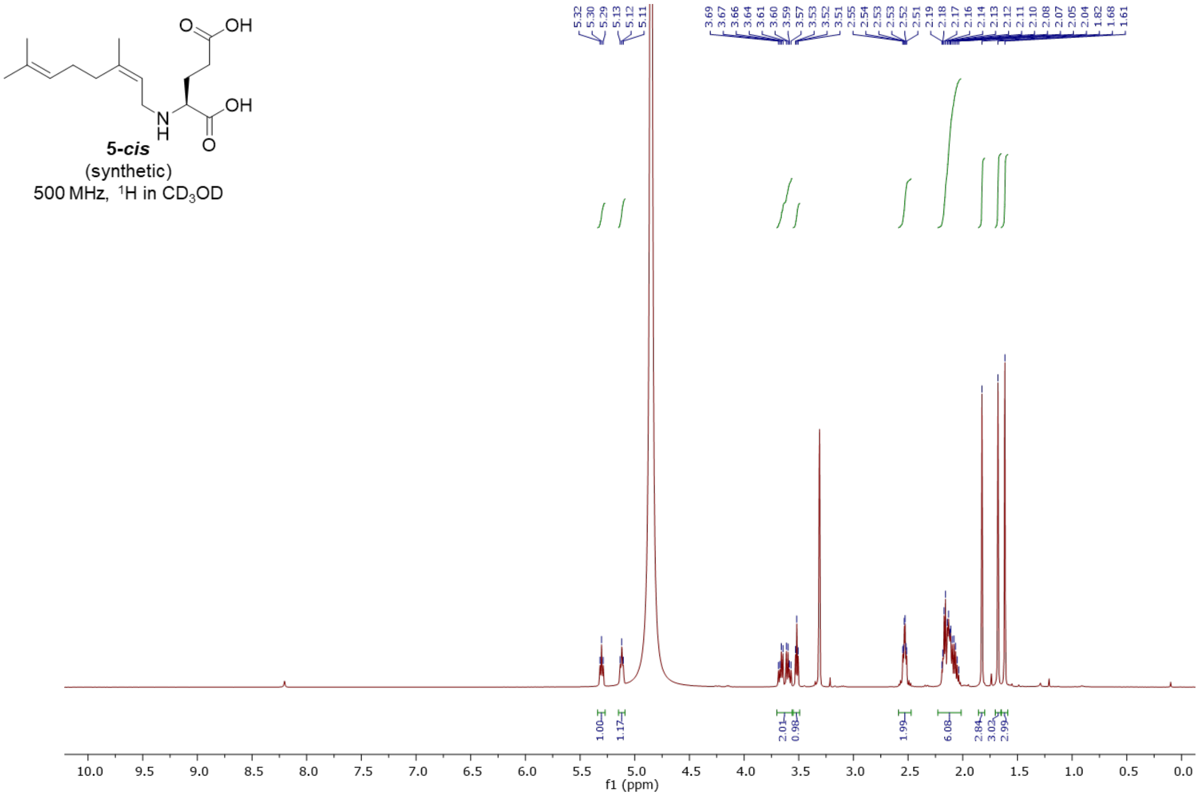


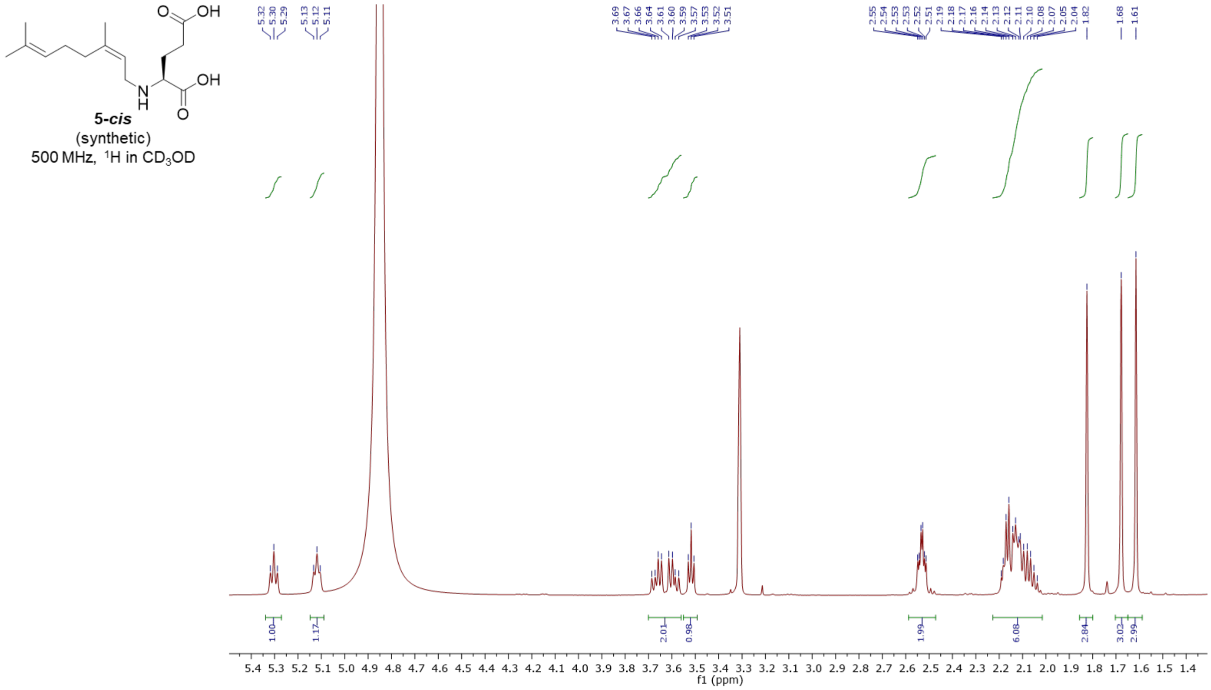

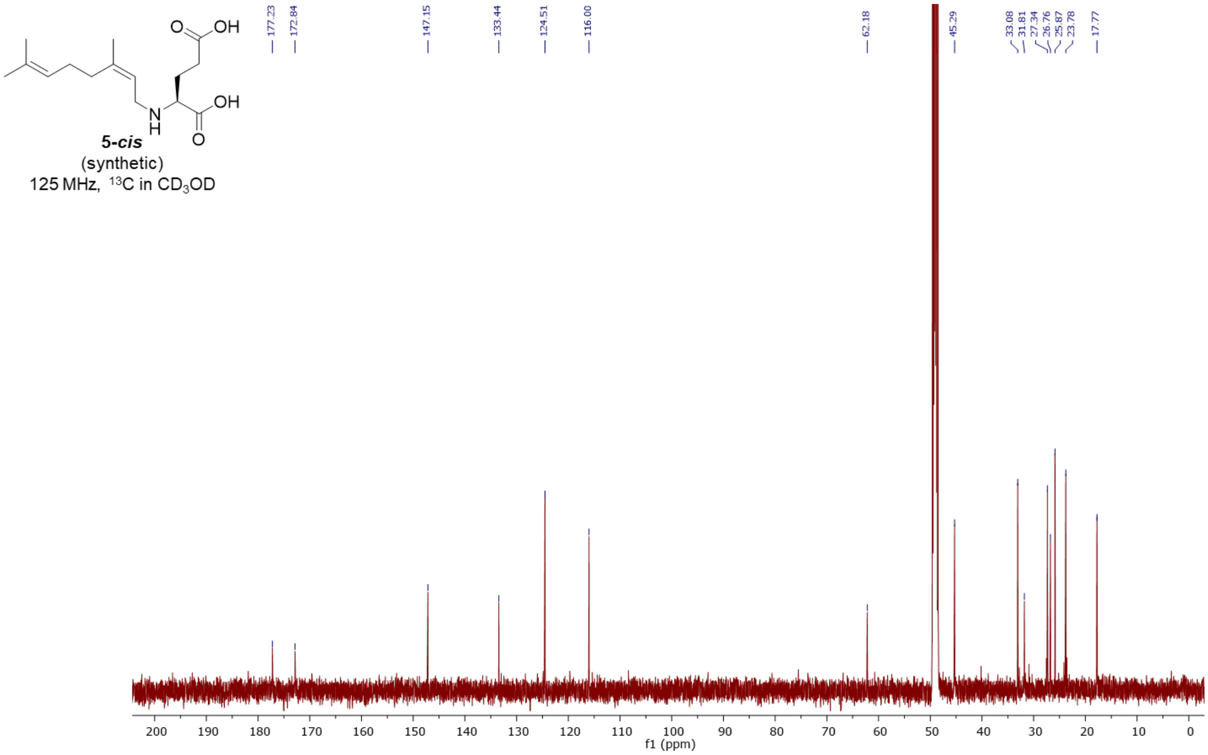

5-cis
(synthetic)
125 MHz, $^{13}C$ in $CD_3OD$
177.23
172.84
147.15
133.44
124.51
116.00
62.18
45.29
33.08
31.81
27.34
26.76
25.87
23.78
17.77
f1 (ppm)


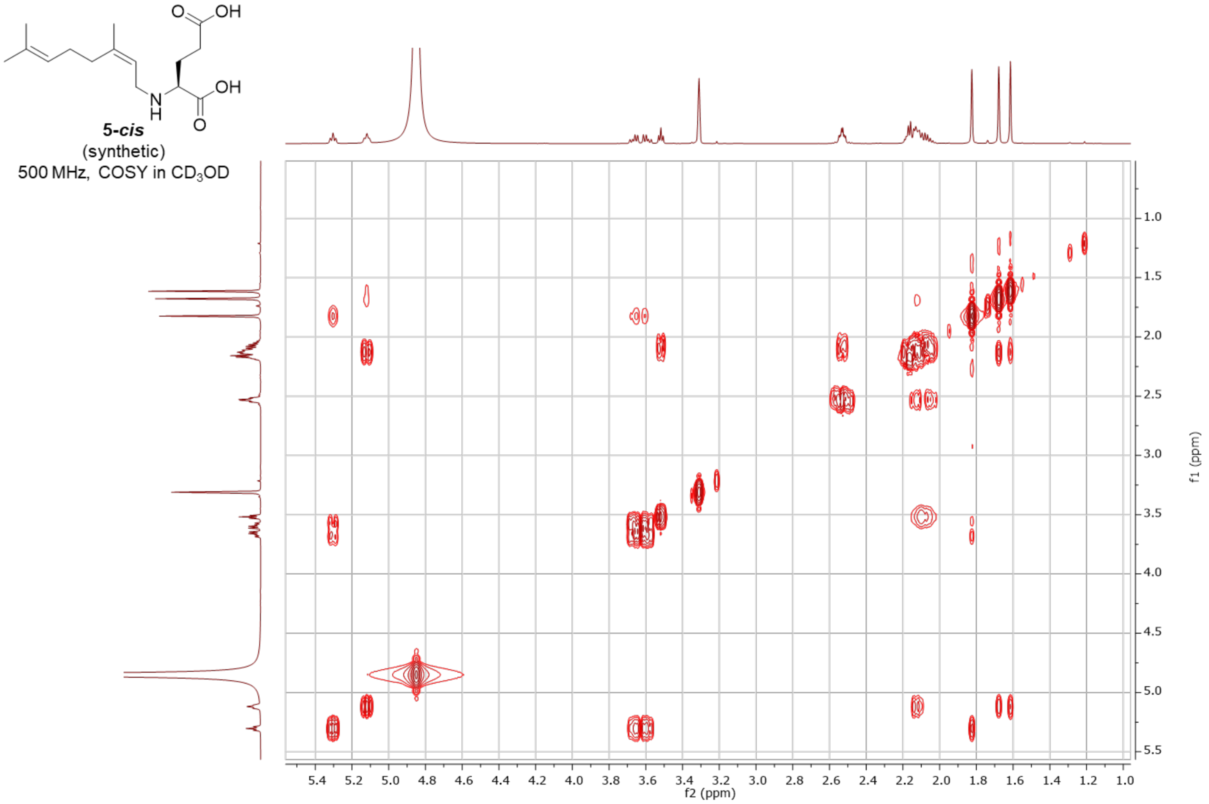

5-cis
(synthetic)
500 MHz, COSY in $CD_3OD$
f2 (ppm)
f1 (ppm)

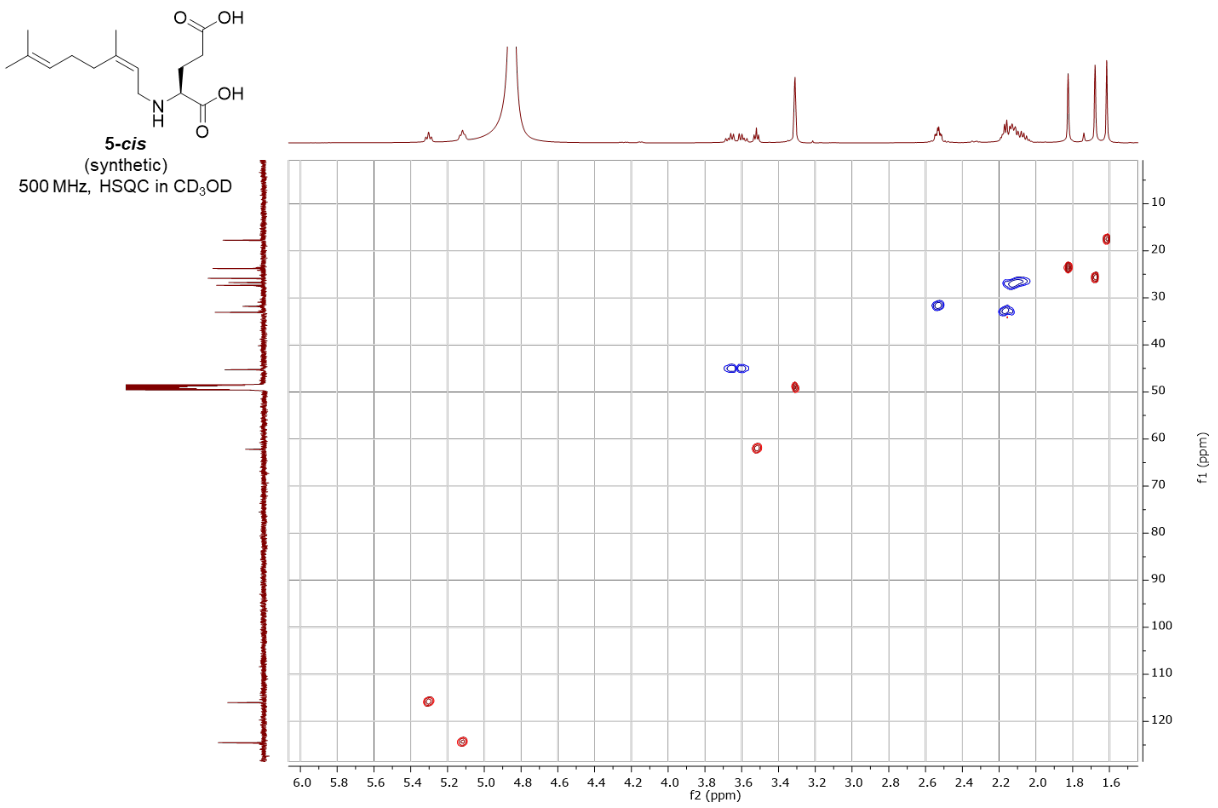

5-*cis*
(synthetic)
500 MHz, HSQC in $CD_3OD$
f2 (ppm)
f1 (ppm)


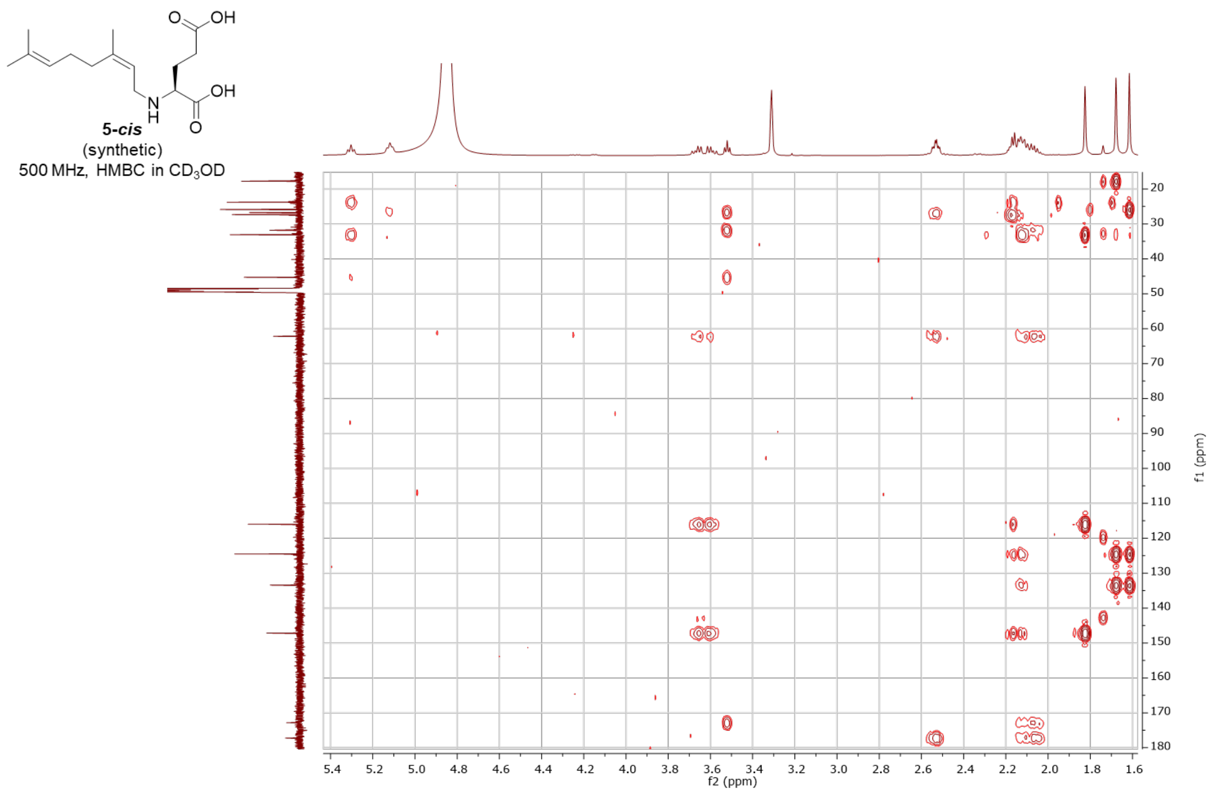

5-*cis*
(synthetic)
500 MHz, HMBC in $CD_3OD$
f2 (ppm)
f1 (ppm)

7’-carboxy-*N*-(3’*S*)-citronellyl-L-glutamic acid - synthetic (**6**)

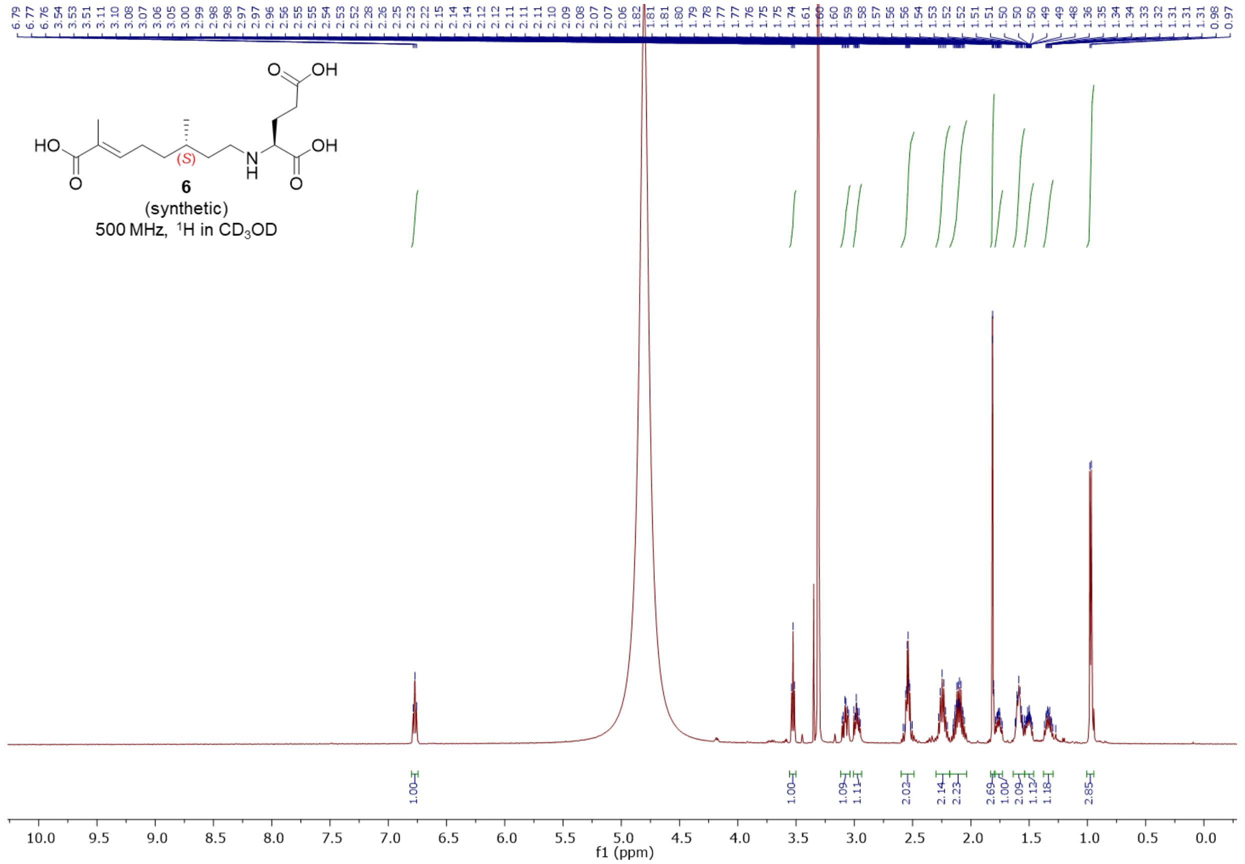


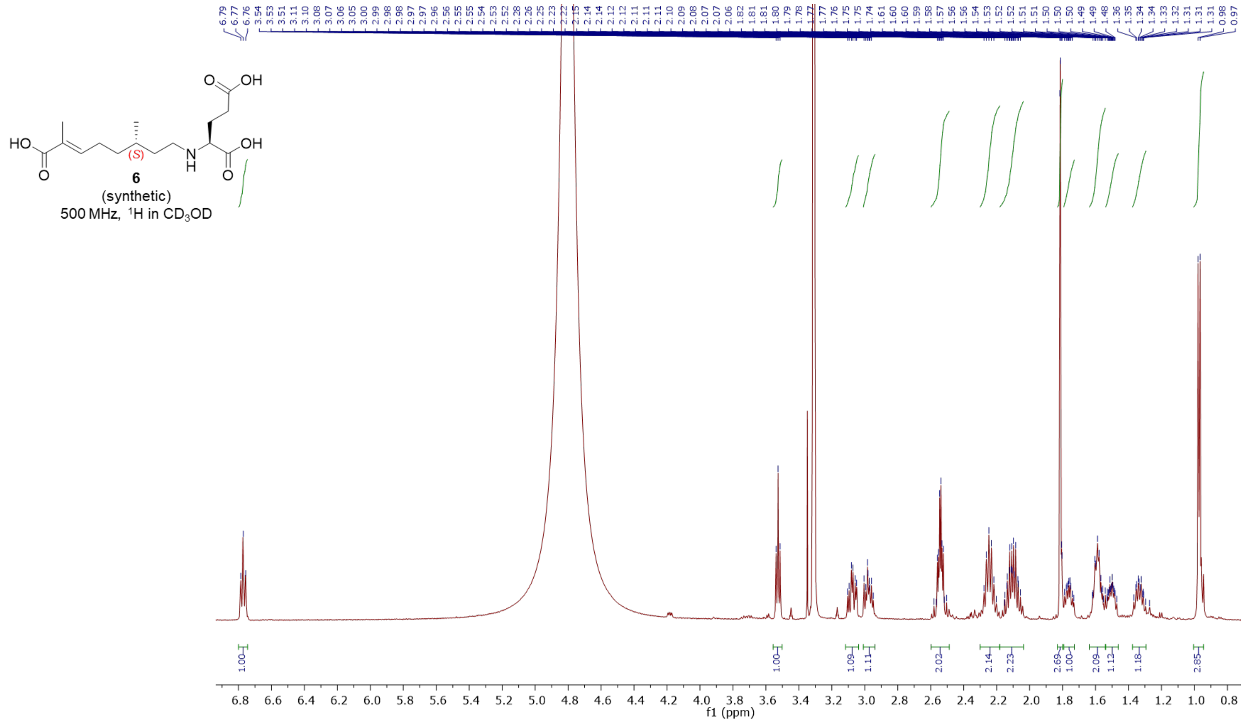

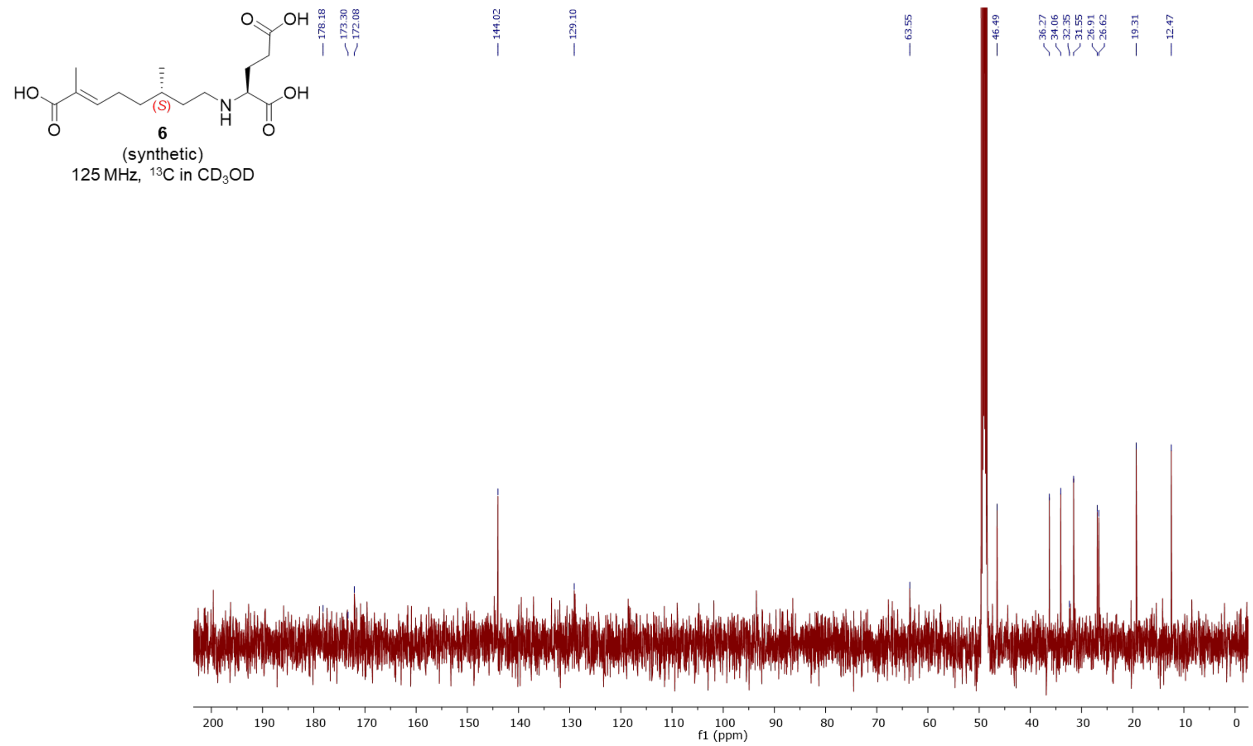
6
(synthetic)
125 MHz, $^{13}C$ in $CD_3OD$
178.18
173.30
172.08
144.02
129.10
63.55
46.49
36.27
34.06
32.35
31.55
26.91
26.62
19.31
12.47
f1 (ppm)

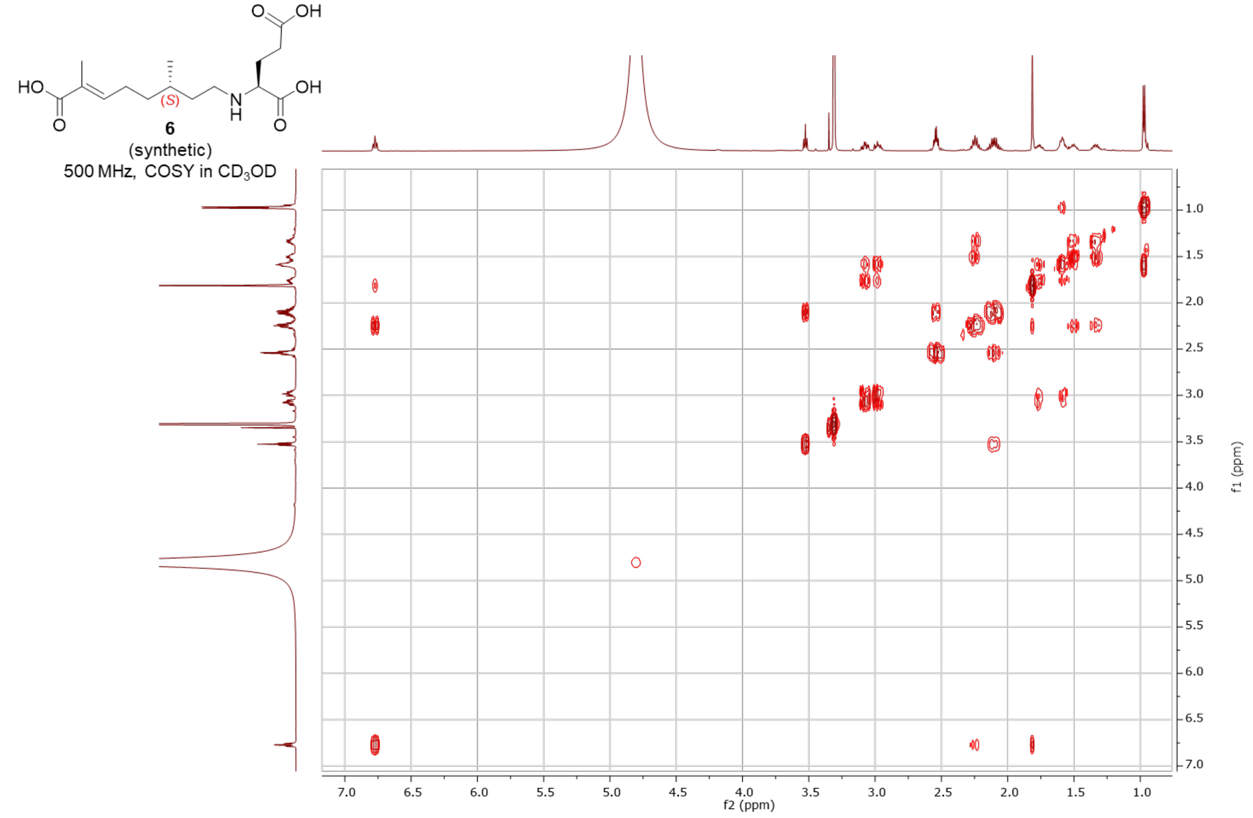
6
(synthetic)
500 MHz, COSY in $CD_3OD$
f2 (ppm)
f1 (ppm)

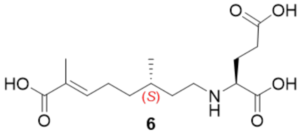


**6**
(synthetic)
500 MHz, HSQC in $CD_3OD$

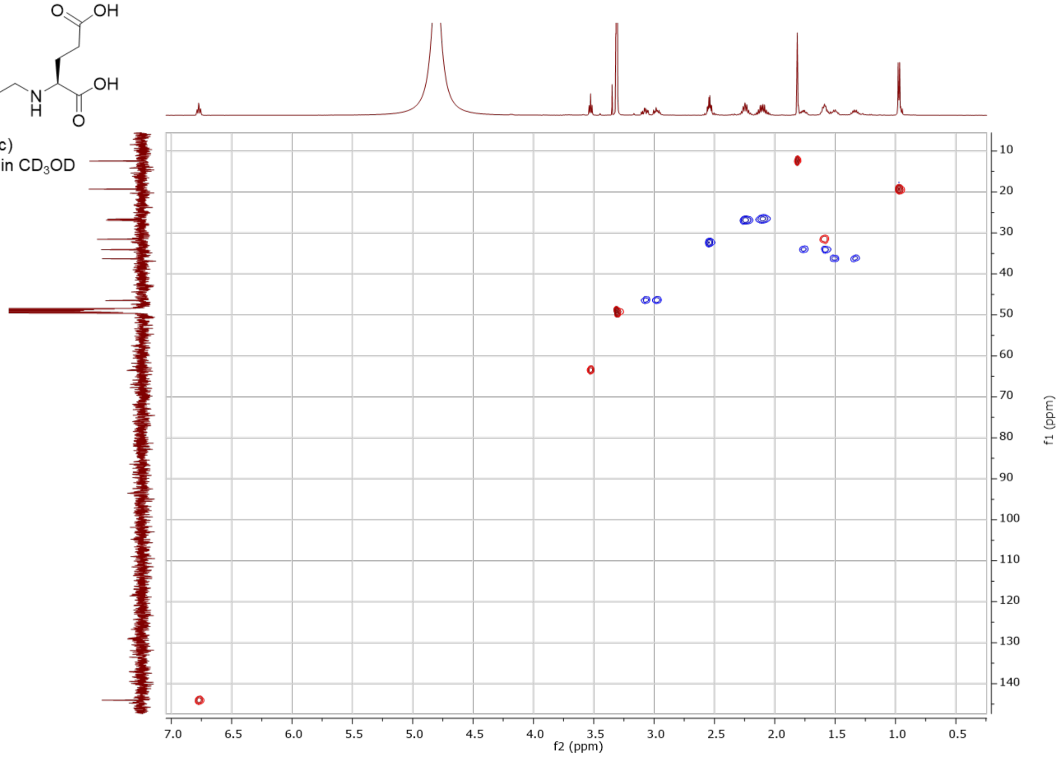


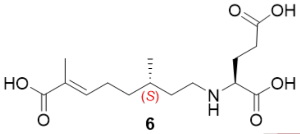


**6**
(synthetic)
500 MHz, HMBC in $CD_3OD$

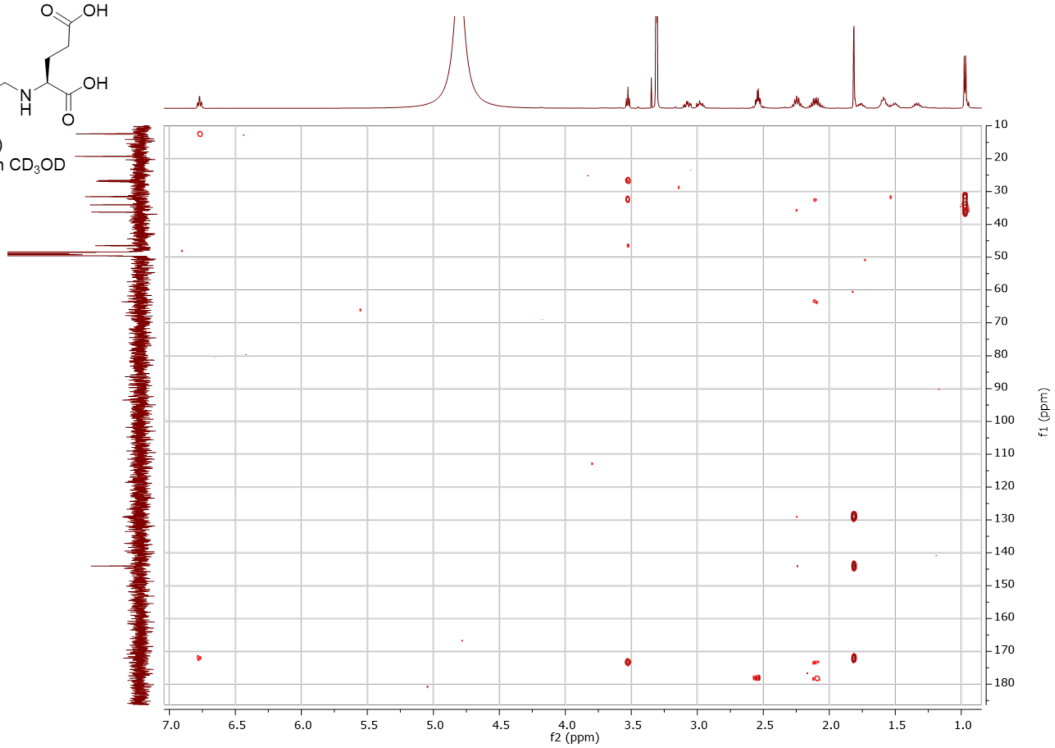

### 7’-carboxy-*N* -(3’*R*)-citronellyl-L-glutamic acid - synthetic (**7**)

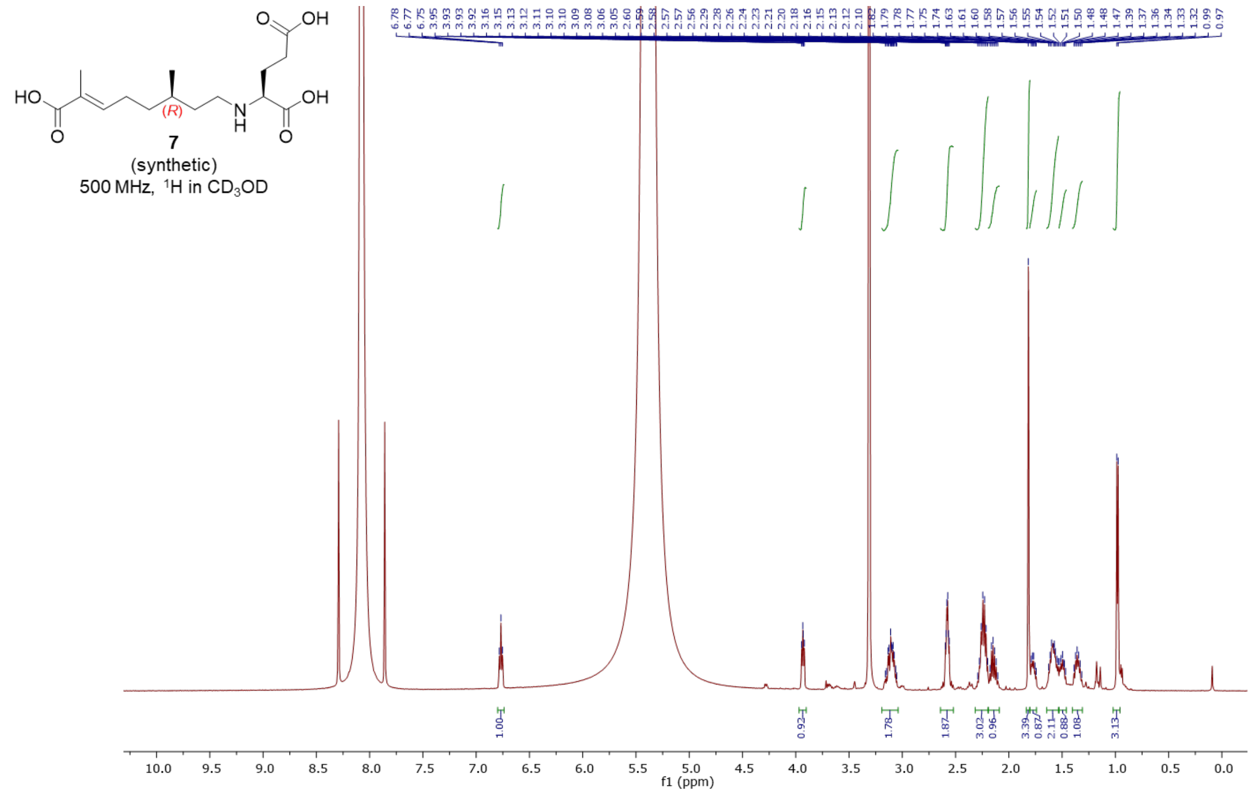


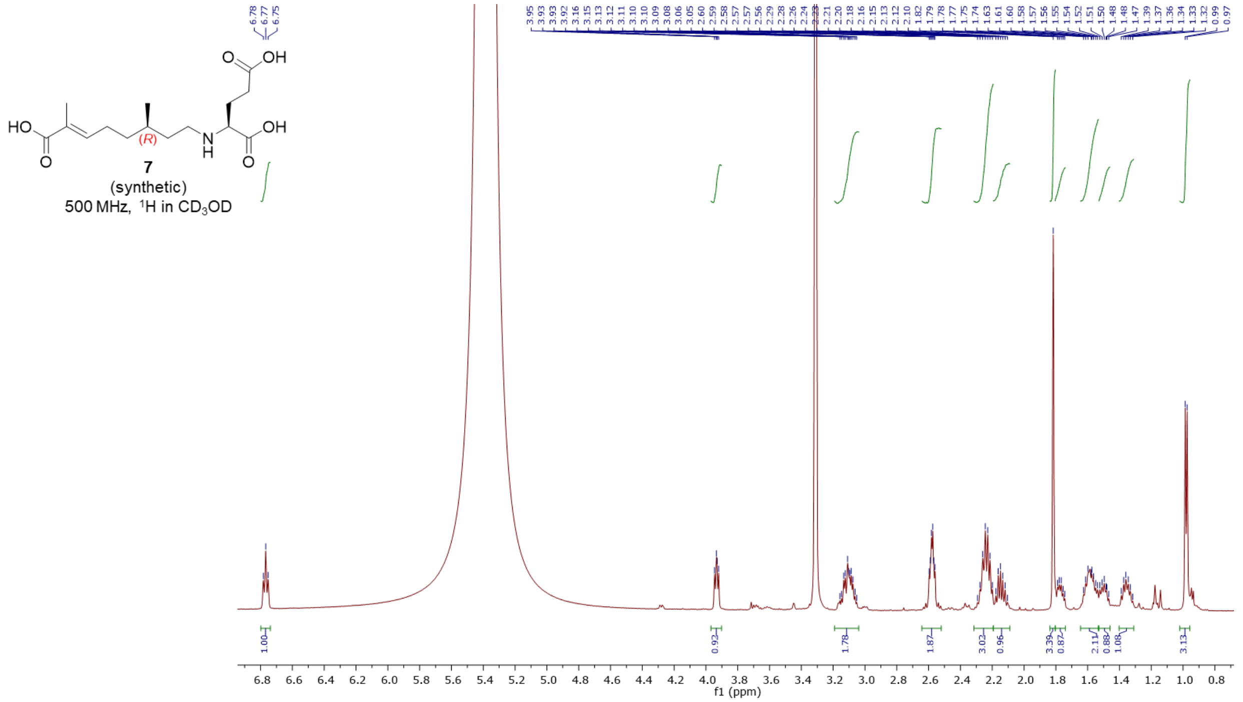

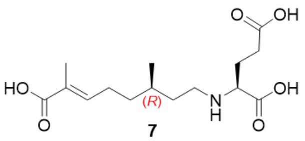

7

(synthetic)

125 MHz, $^{13}C$ in $CD_3OD$

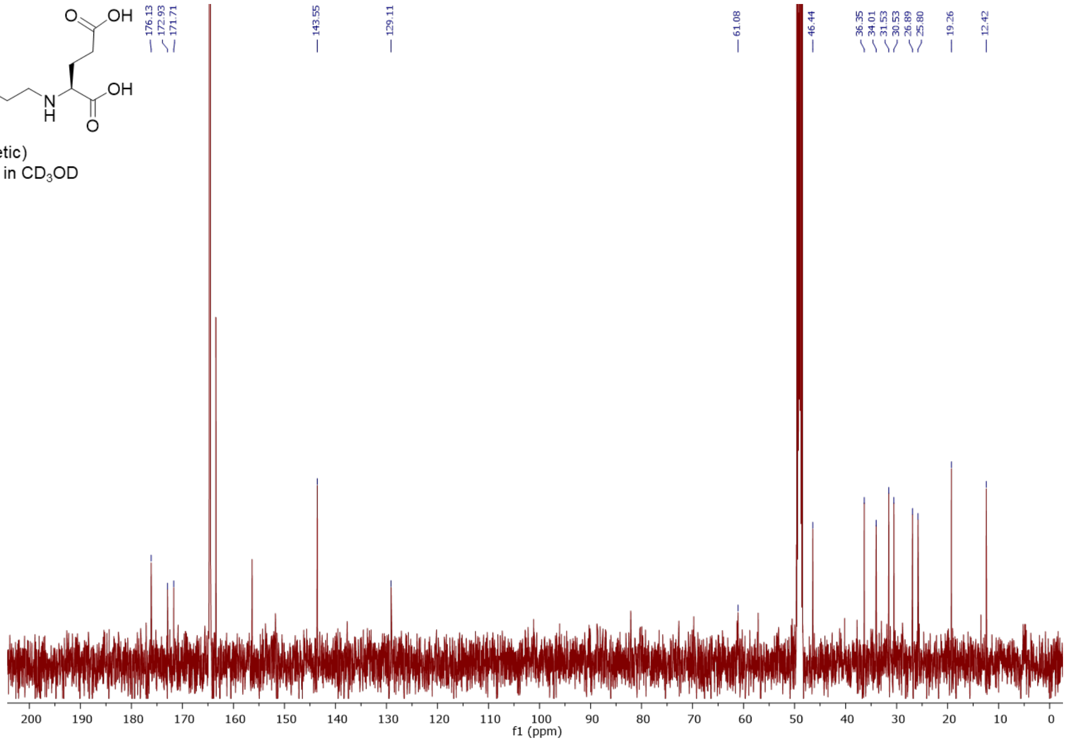


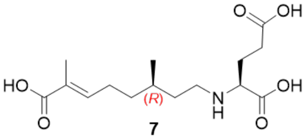

7

(synthetic)

500 MHz, COSY in $CD_3OD$

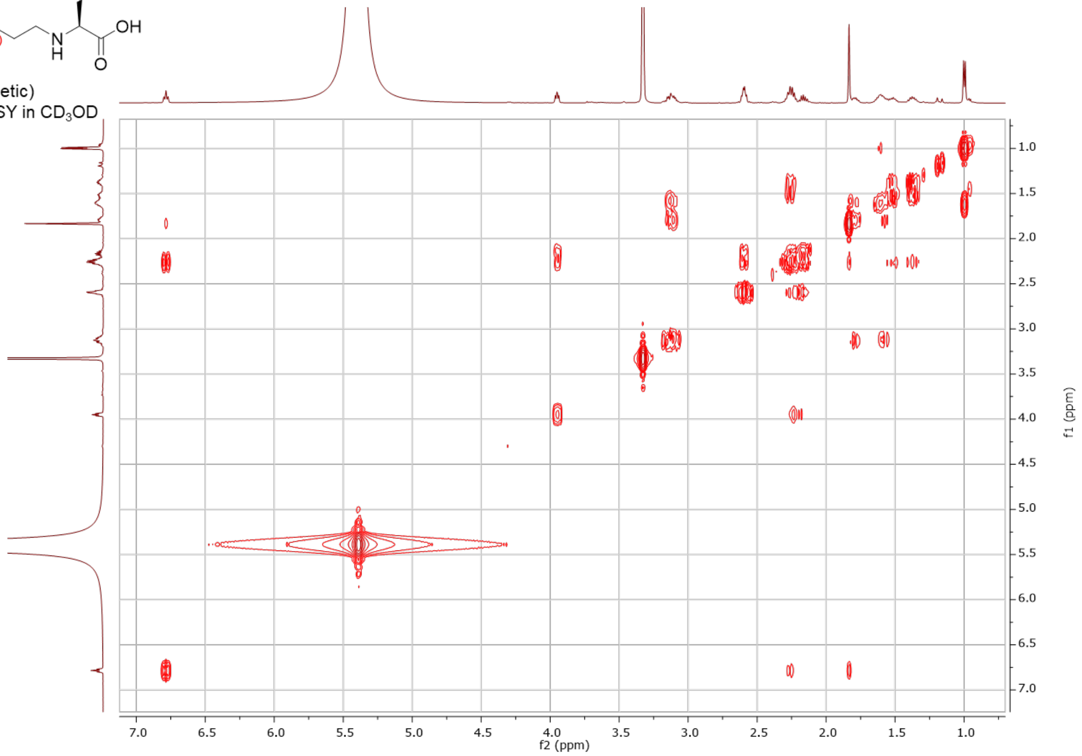

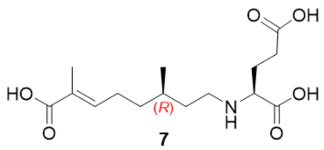


**7**
(synthetic)
500 MHz, HSQC in $CD_3OD$

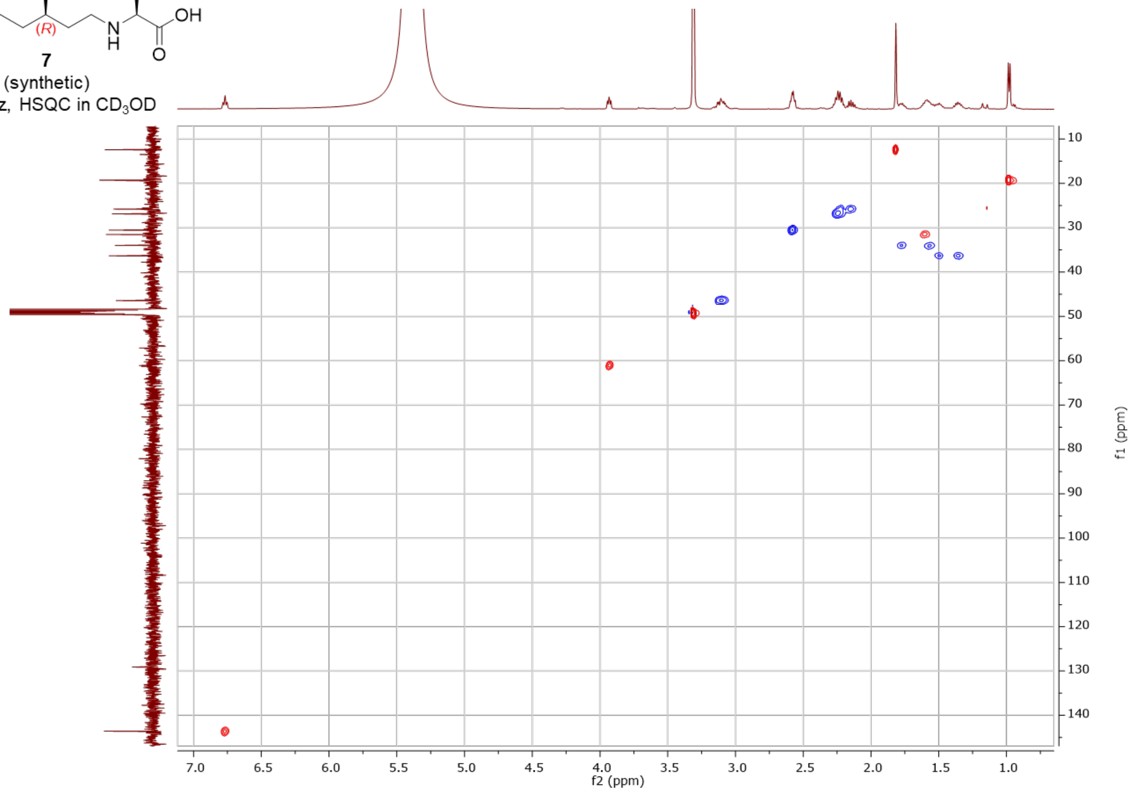


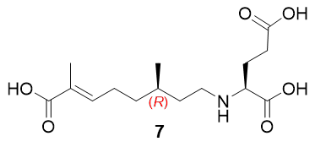


**7**
(synthetic)
500 MHz, HMBC in $CD_3OD$

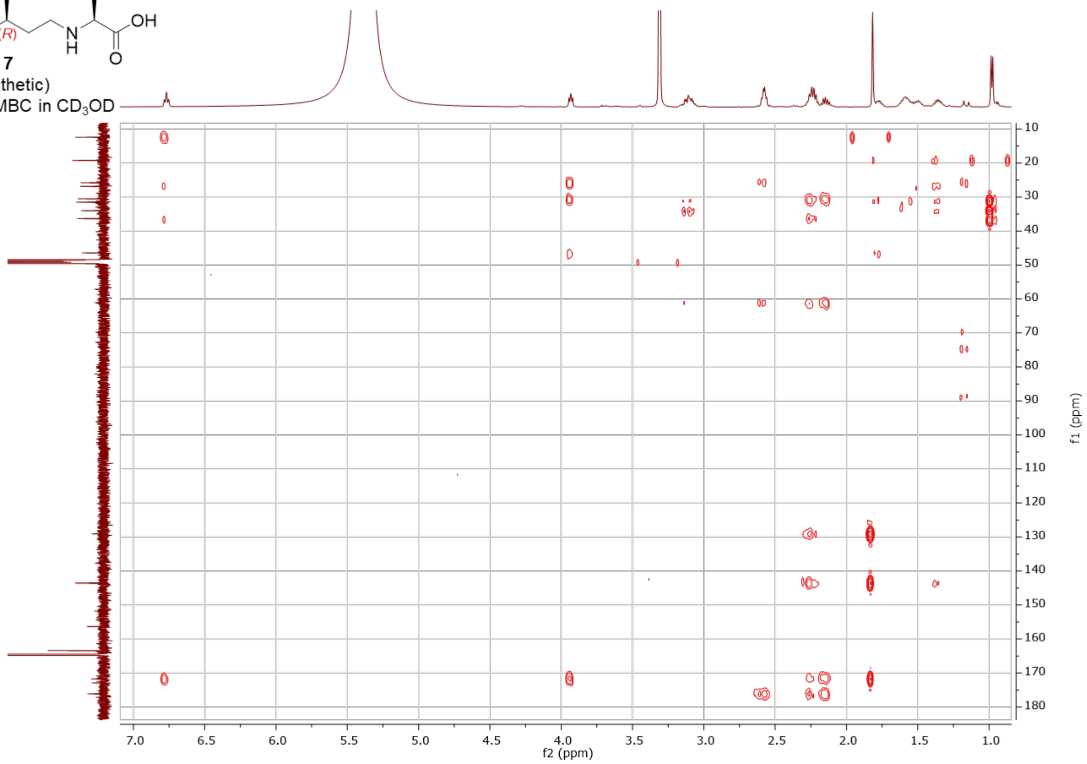

<u>7'-carboxy-*N*-(3'*S*)-citronellyl-3*R*-hydroxy-L-glutamic acid – enzymatic (**8**)</u>

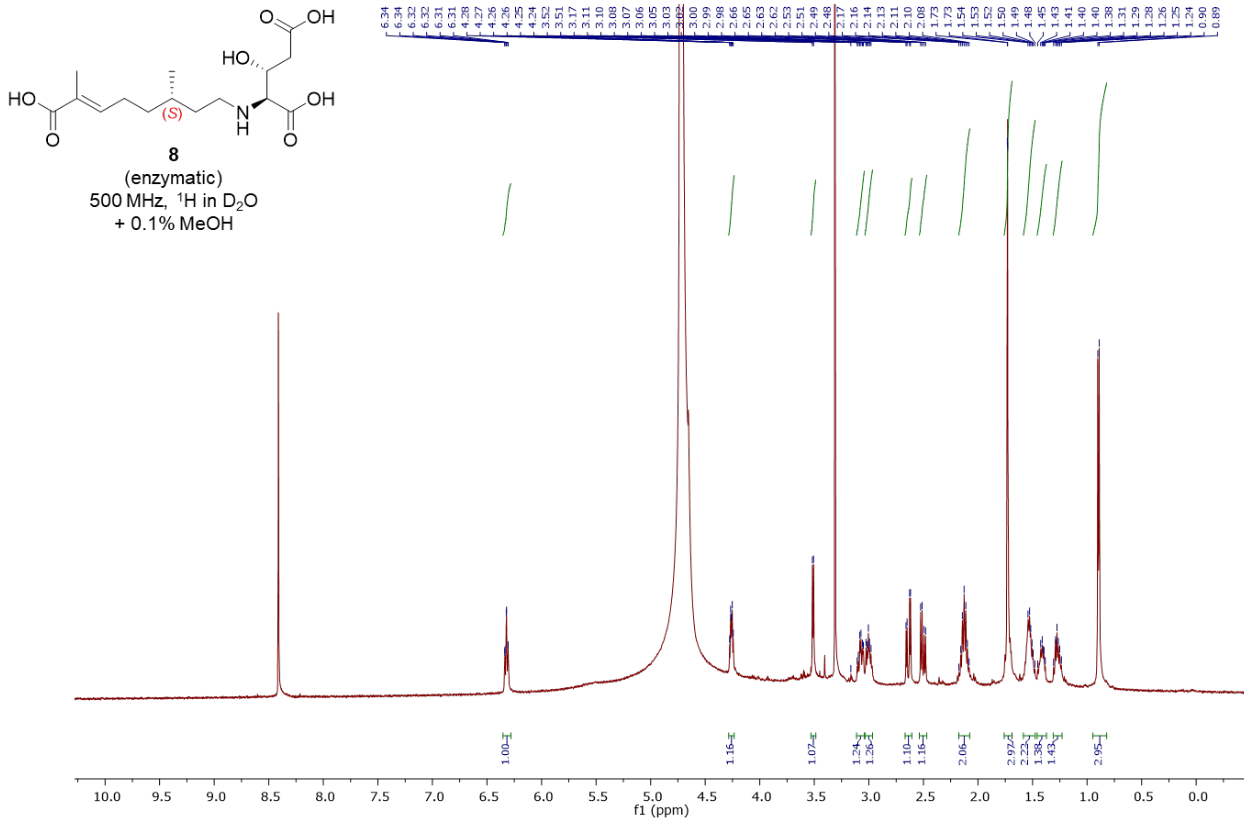


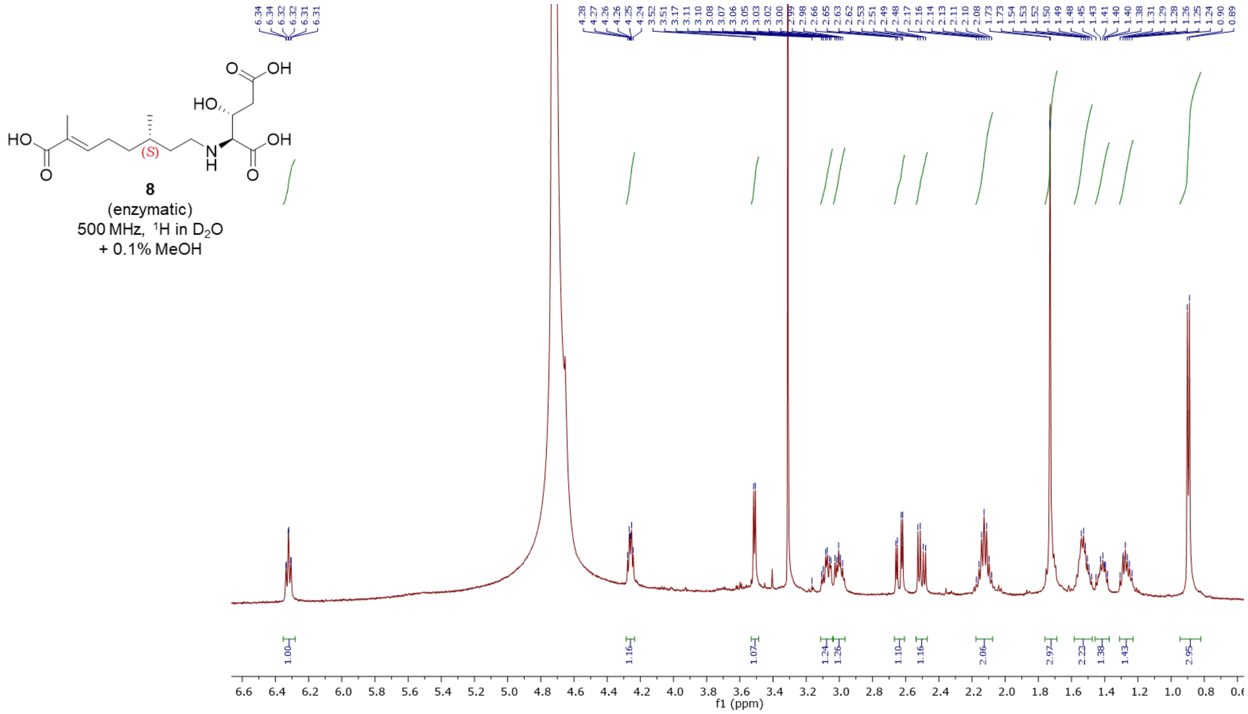

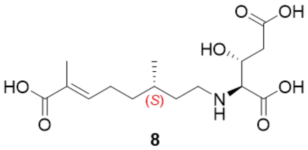


**8**
(enzymatic)
500 MHz, COSY in $D_2O$
+ 0.1% MeOH

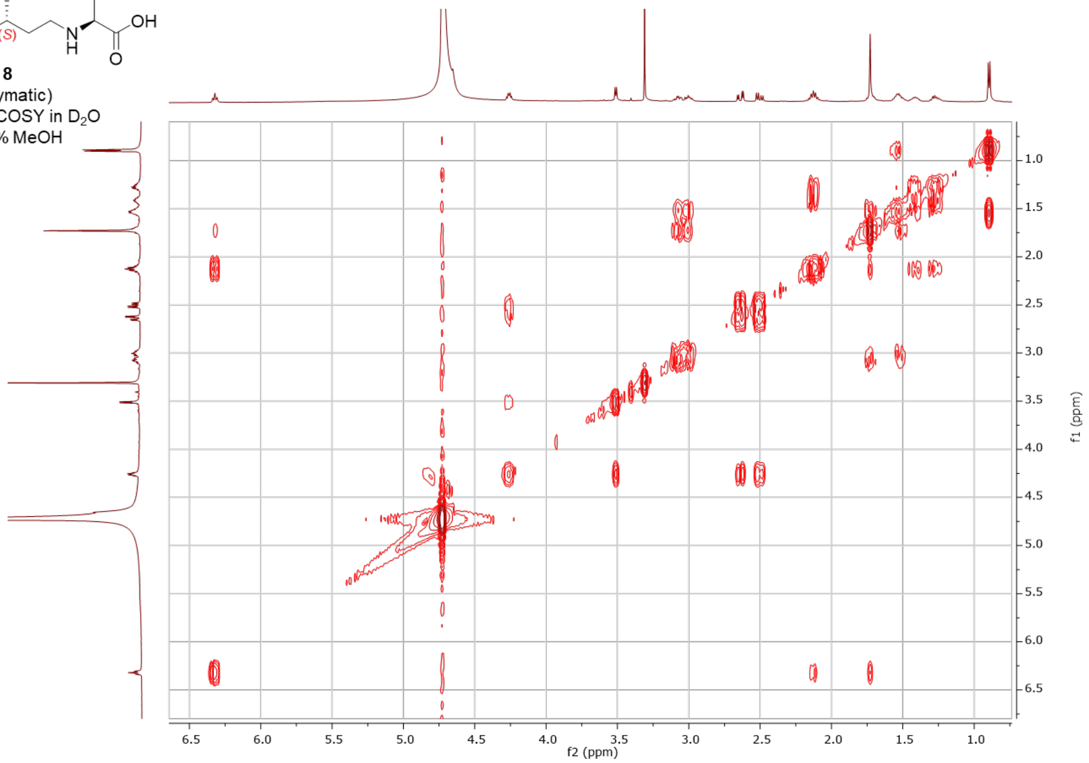


**8**
(enzymatic)
(500 MHz, HSQC in $D_2O$
+ 0.1% MeOH

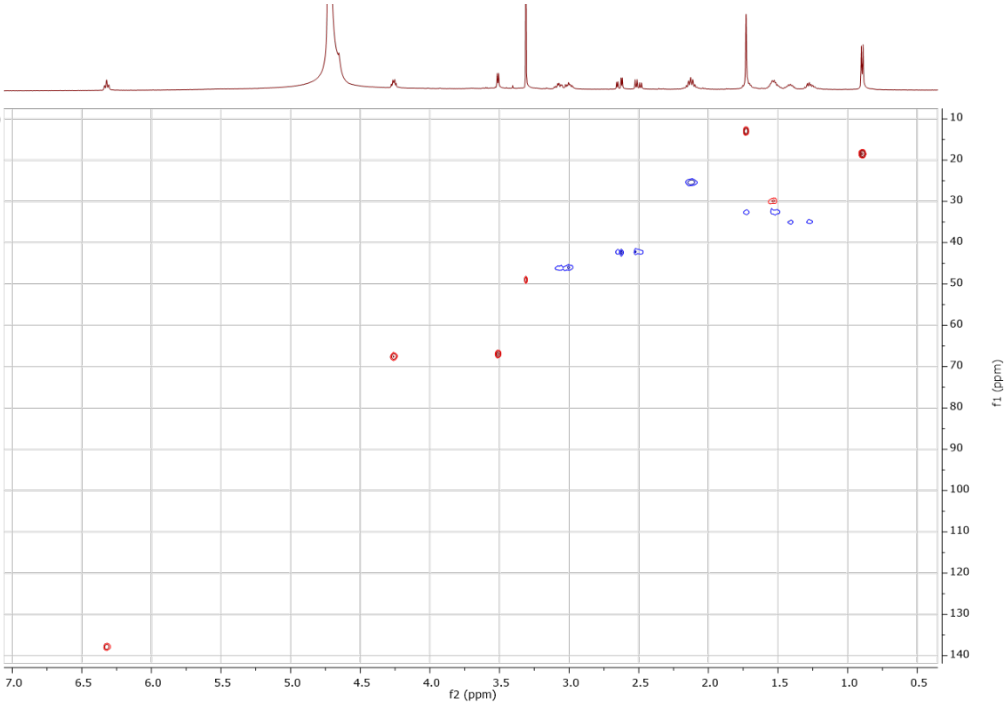

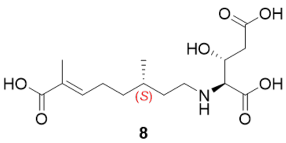

**8**
(enzymatic)
500 MHz, HMBC in $D_2O$
+ 0.1% MeOH

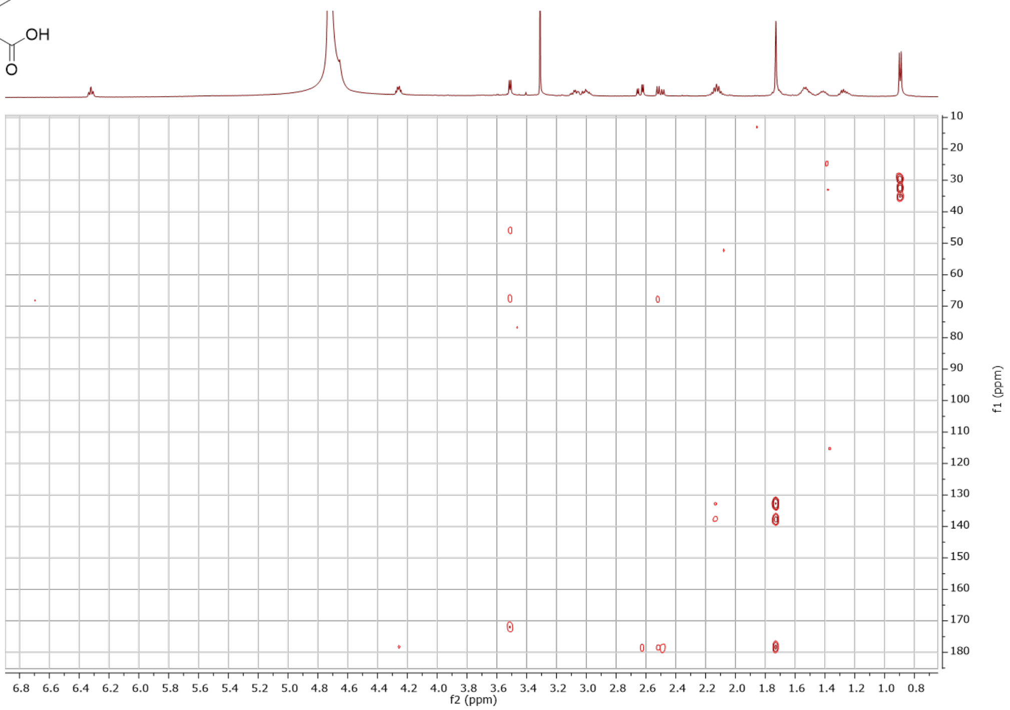

7’-hydroxy-*N*-(3’*S*)-citronellyl-L-glutamic acid - synthetic (**9**)

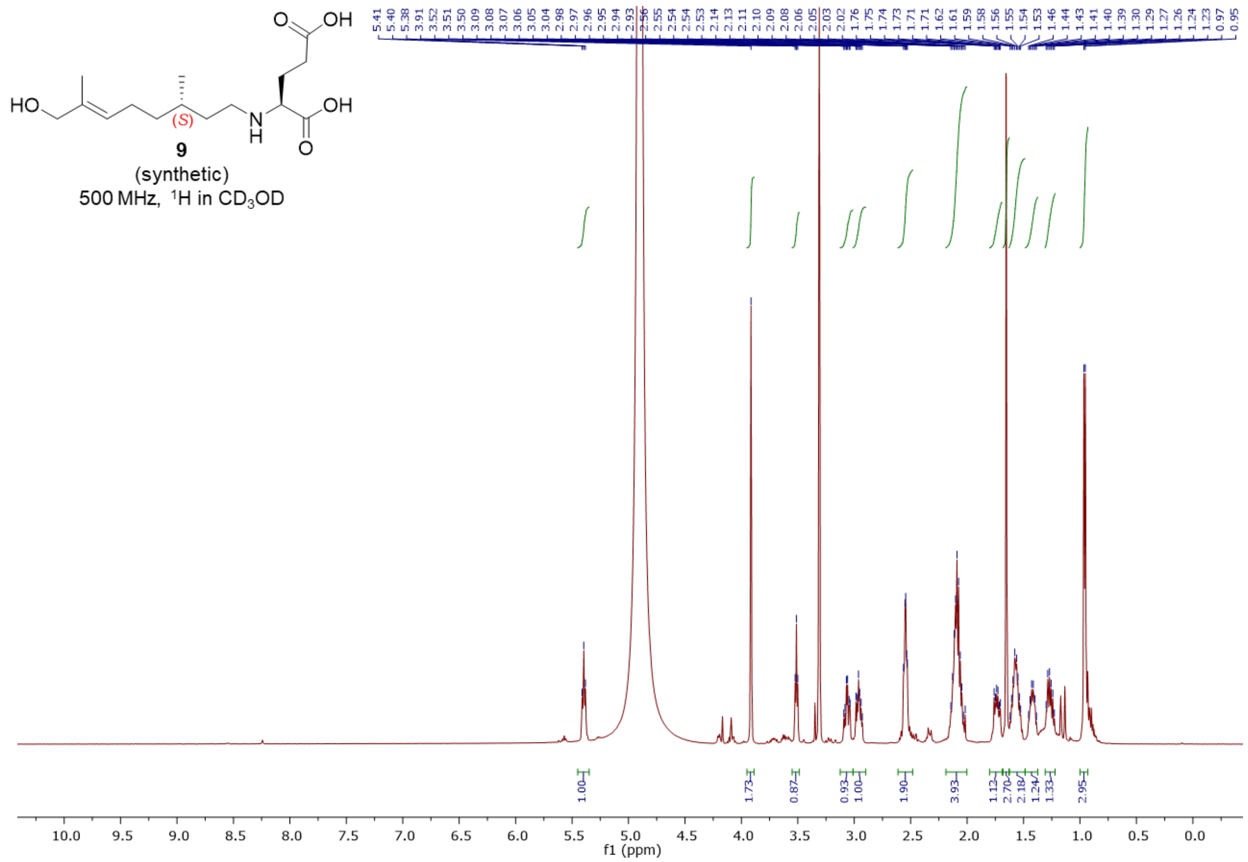


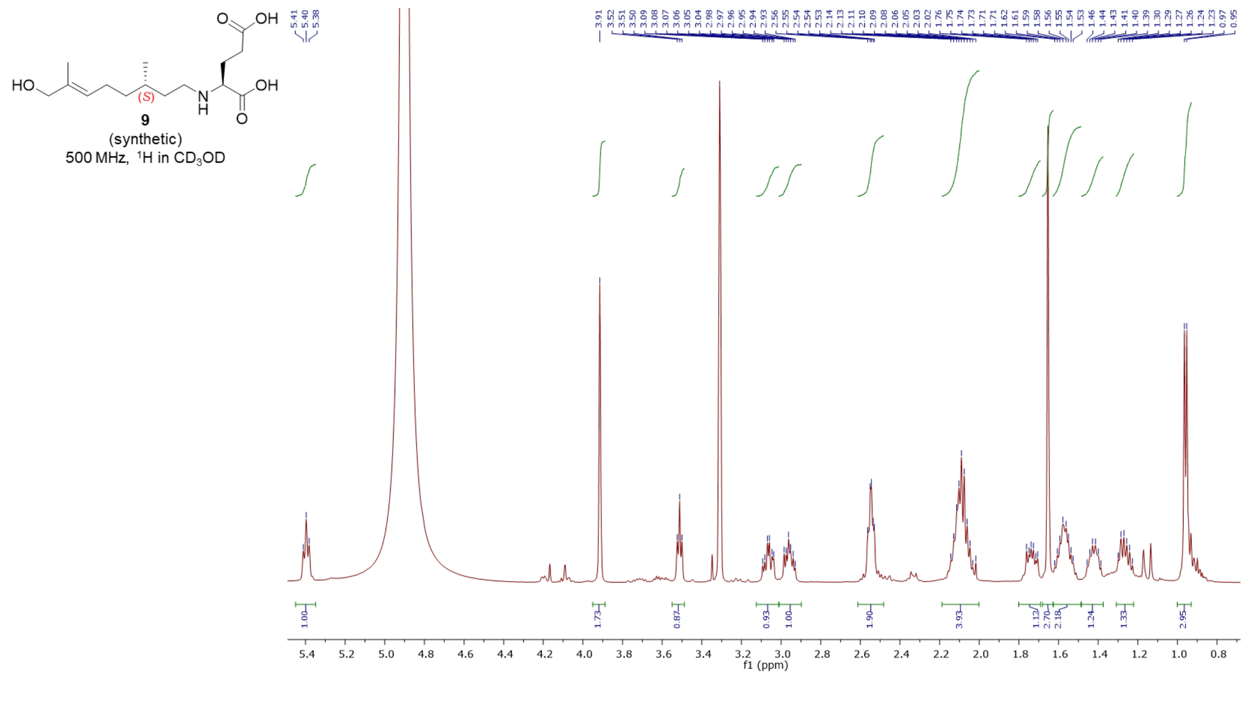

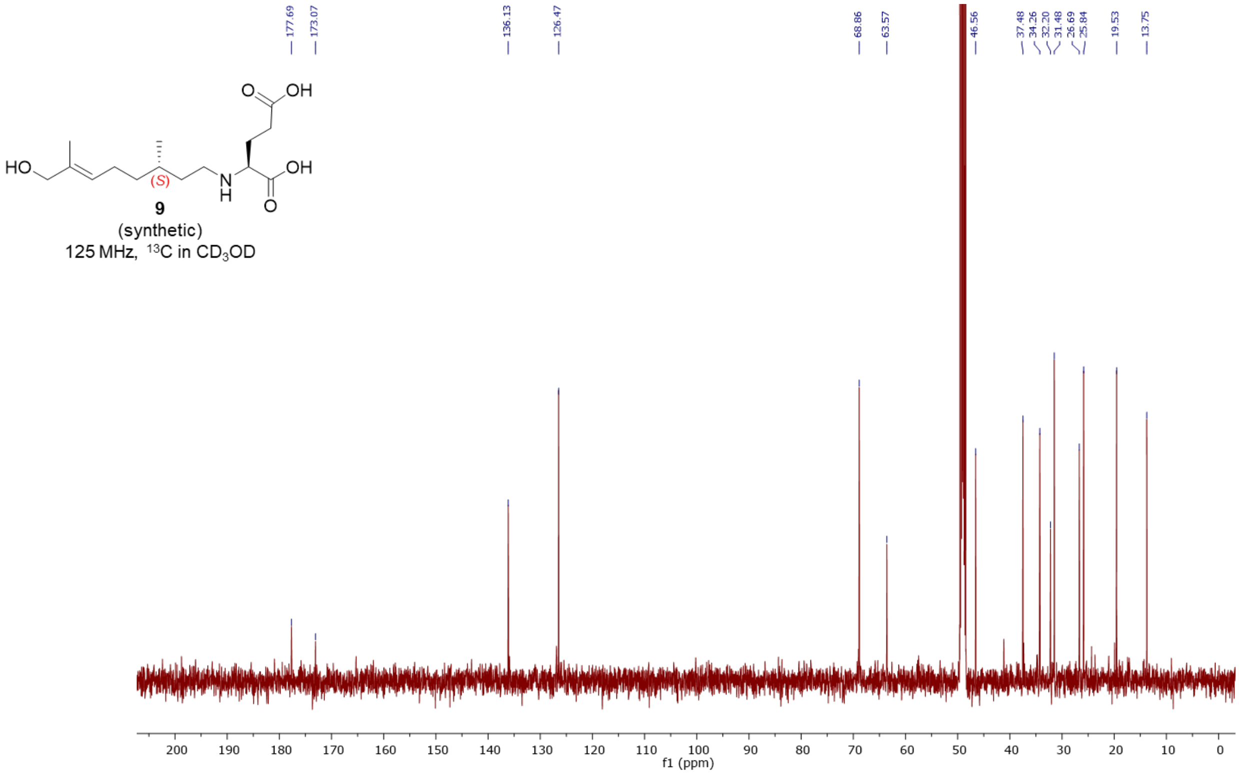

HO
OH
O
N
H
(S)
9
(synthetic)
125 MHz, 13C in CD3OD
177.69
173.07
136.13
126.47
68.86
63.57
46.56
37.48
34.26
32.20
31.48
26.69
25.84
19.53
13.75
f1 (ppm)


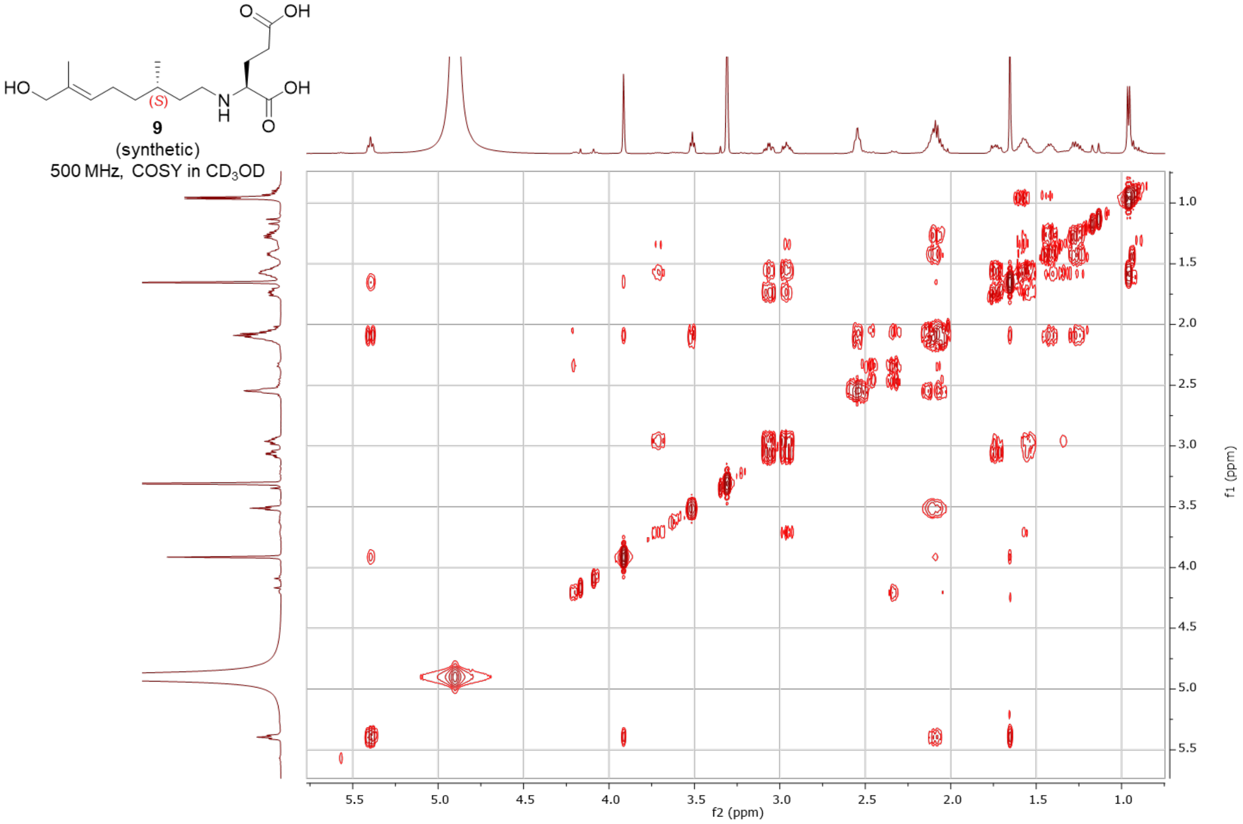

HO
OH
O
N
H
(S)
9
(synthetic)
500 MHz, COSY in CD3OD
f2 (ppm)
f1 (ppm)

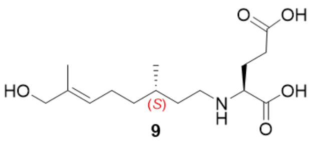


**9**
(synthetic)
500 MHz, HSQC in $CD_3OD$

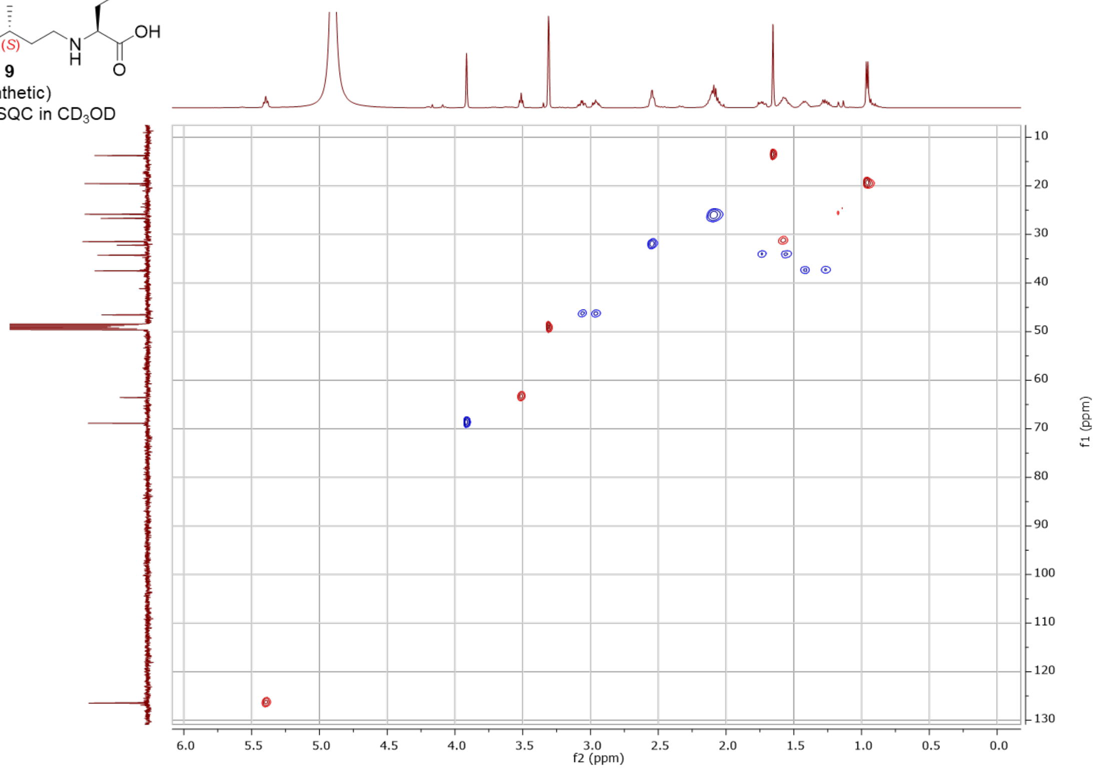


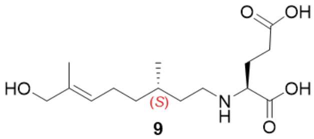


**9**
(synthetic)
500 MHz, HMBC in $CD_3OD$

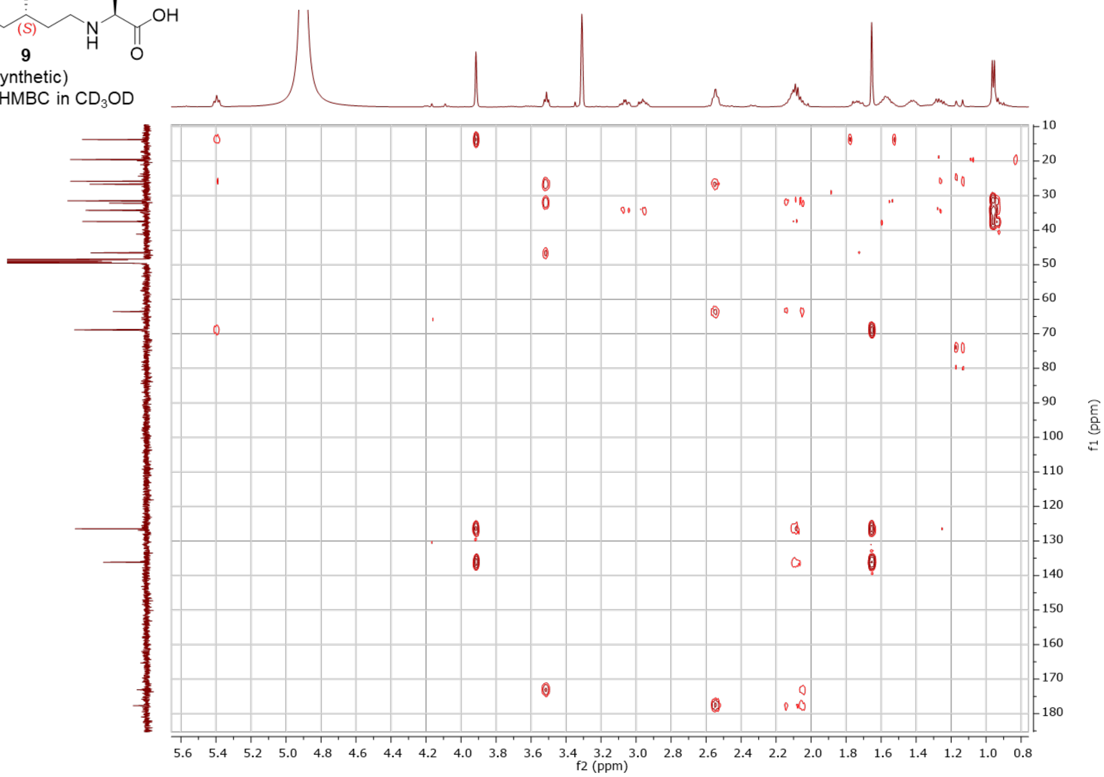

#### 7’-hydroxy-*N*-(3’*R*)-citronellyl-L-glutamic acid - synthetic (**10**)

**10**
(synthetic)
500 MHz, $^{1}$H in $CD_3OD$

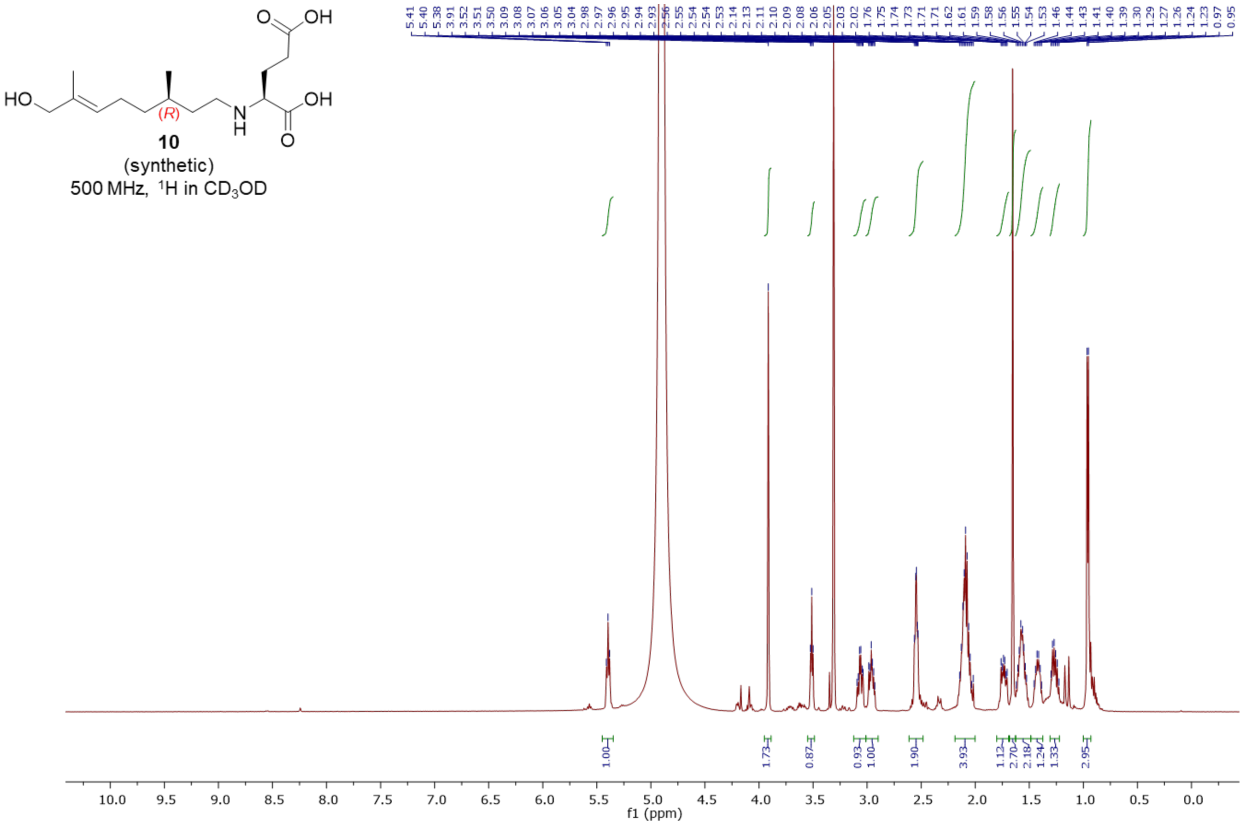


**10**
(synthetic)
500 MHz, $^{1}$H in $CD_3OD$

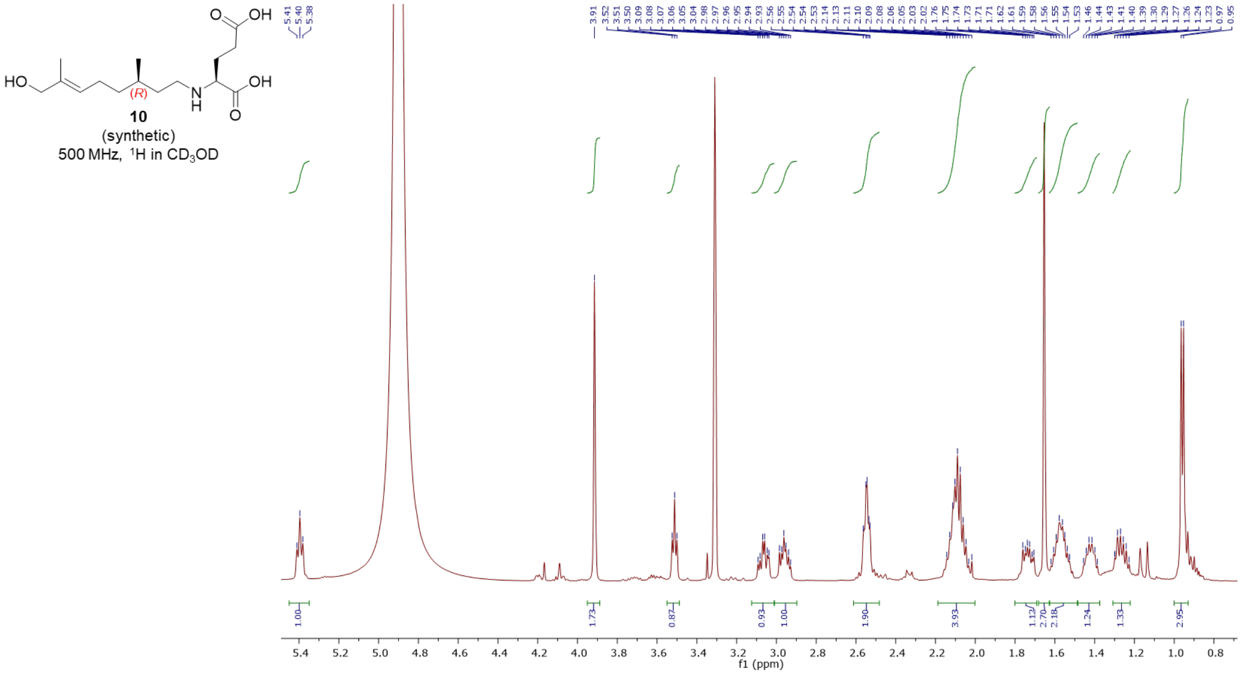

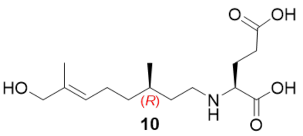


**10**
(synthetic)
125 MHz, $^{13}C$ in $CD_3OD$

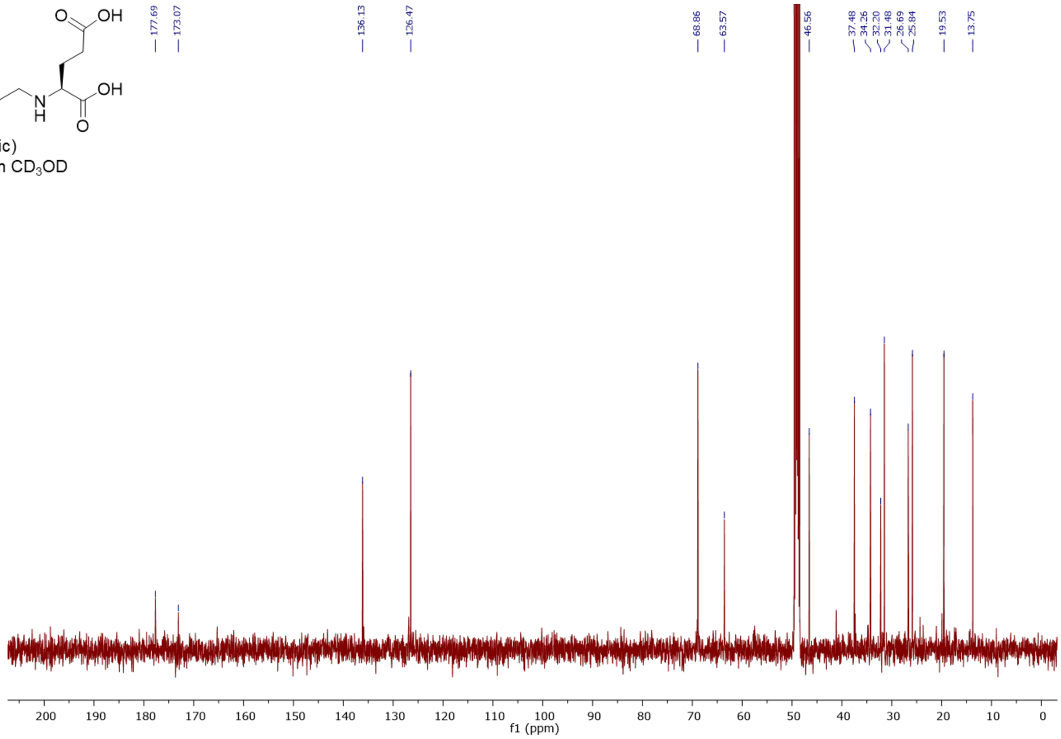


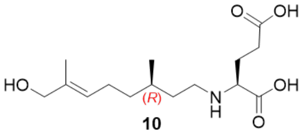


**10**
(synthetic)
500 MHz, COSY in $CD_3OD$

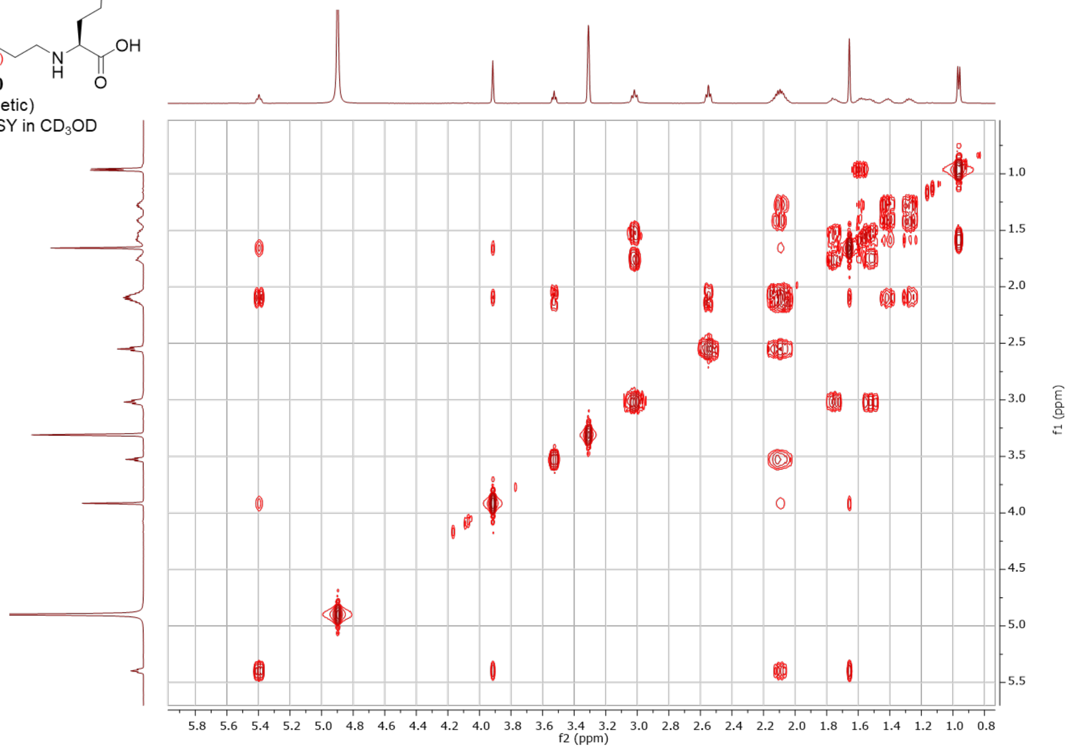

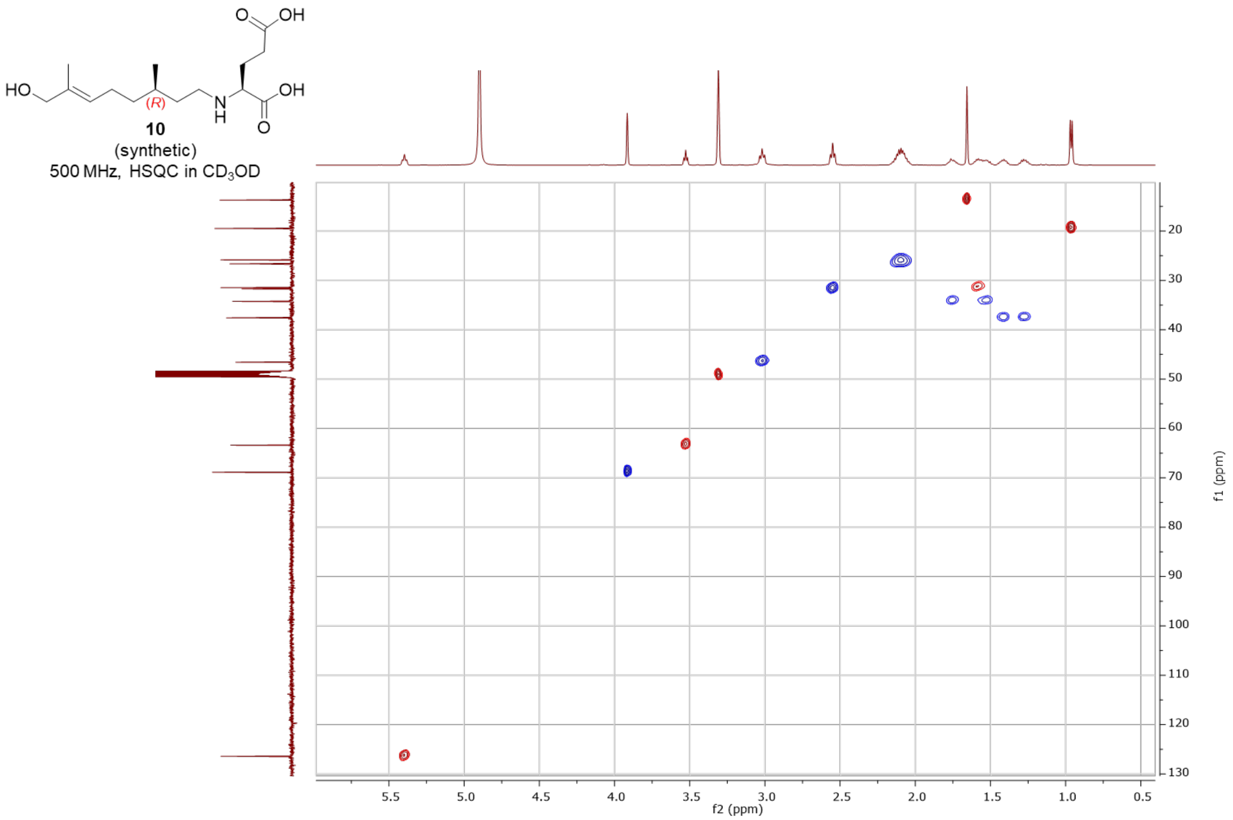


**10**
(synthetic)
500 MHz, HSQC in $CD_3OD$

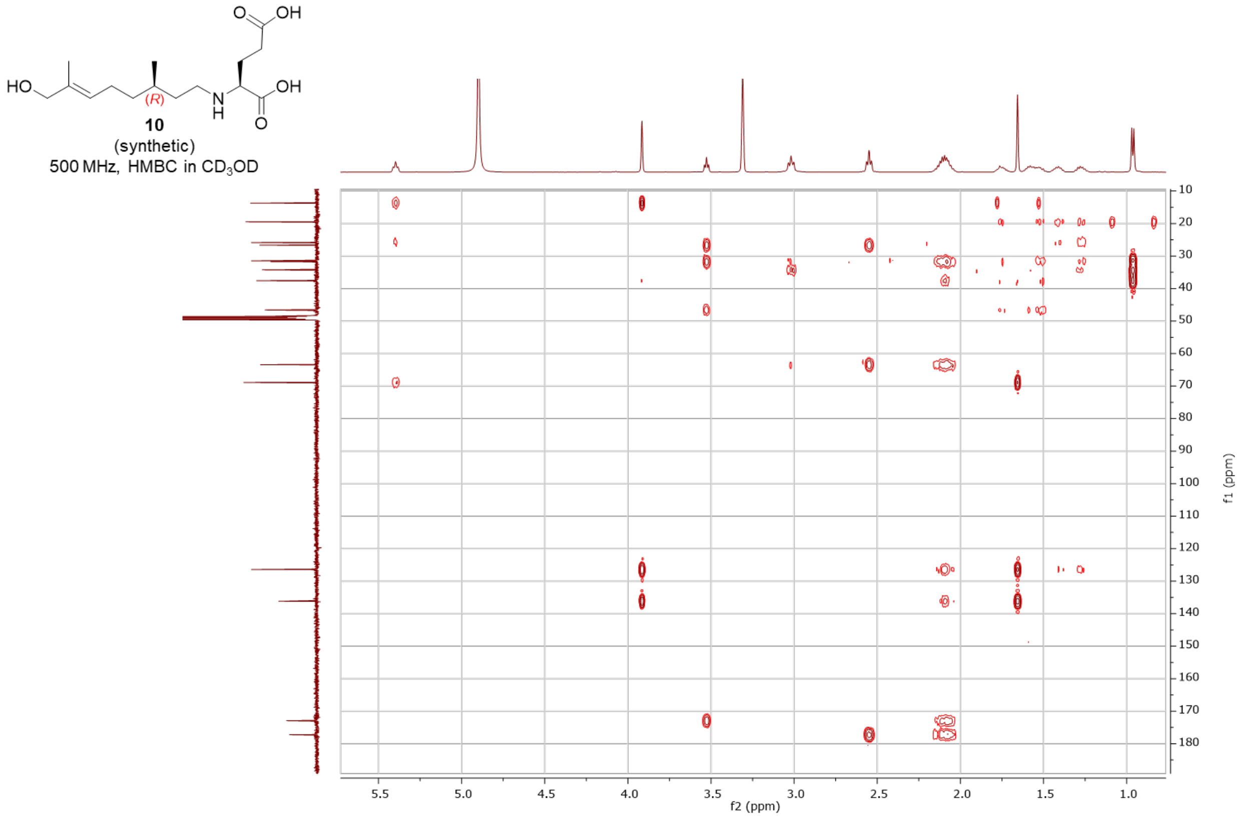


**10**
(synthetic)
500 MHz, HMBC in $CD_3OD$

*N*-(*S*)-citronellyl- L-glutamic acid - synthetic (**11**)

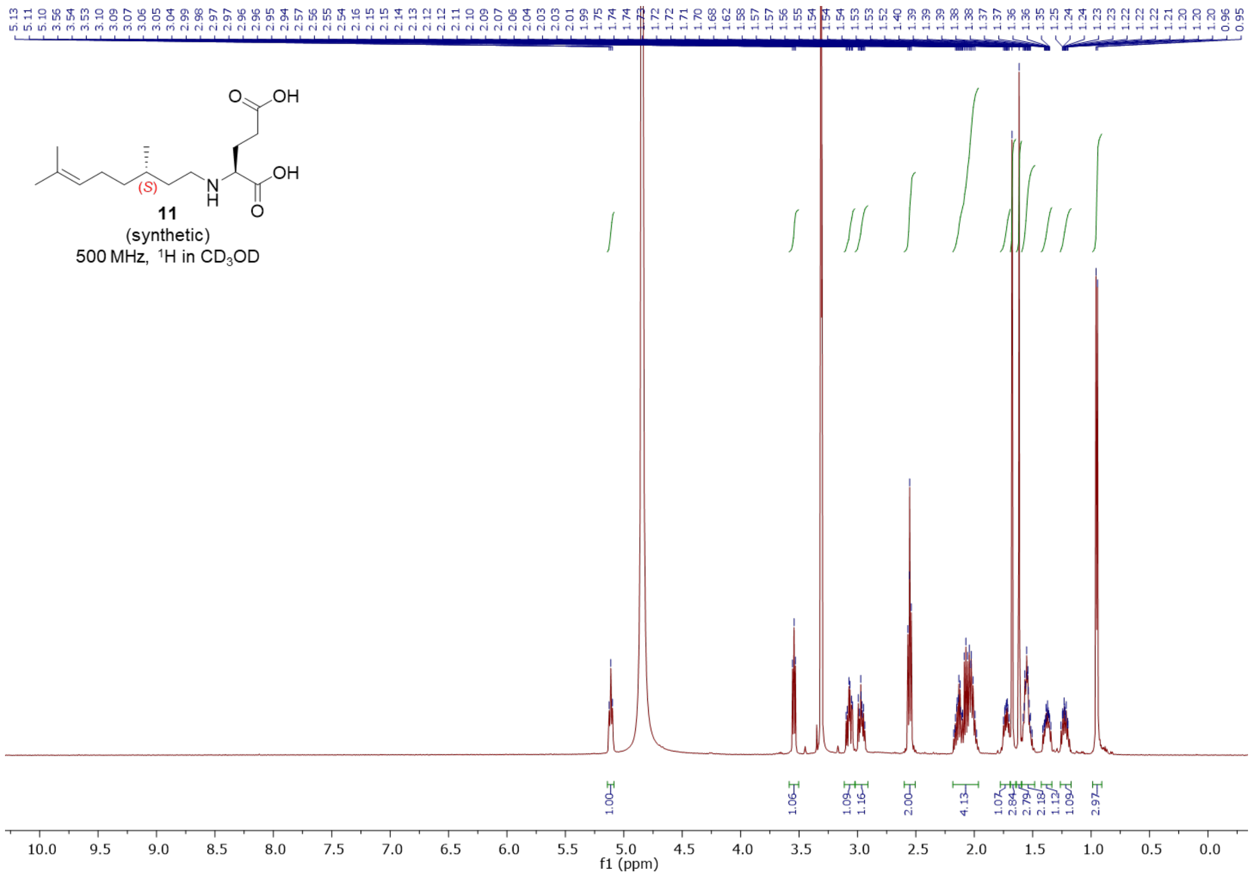


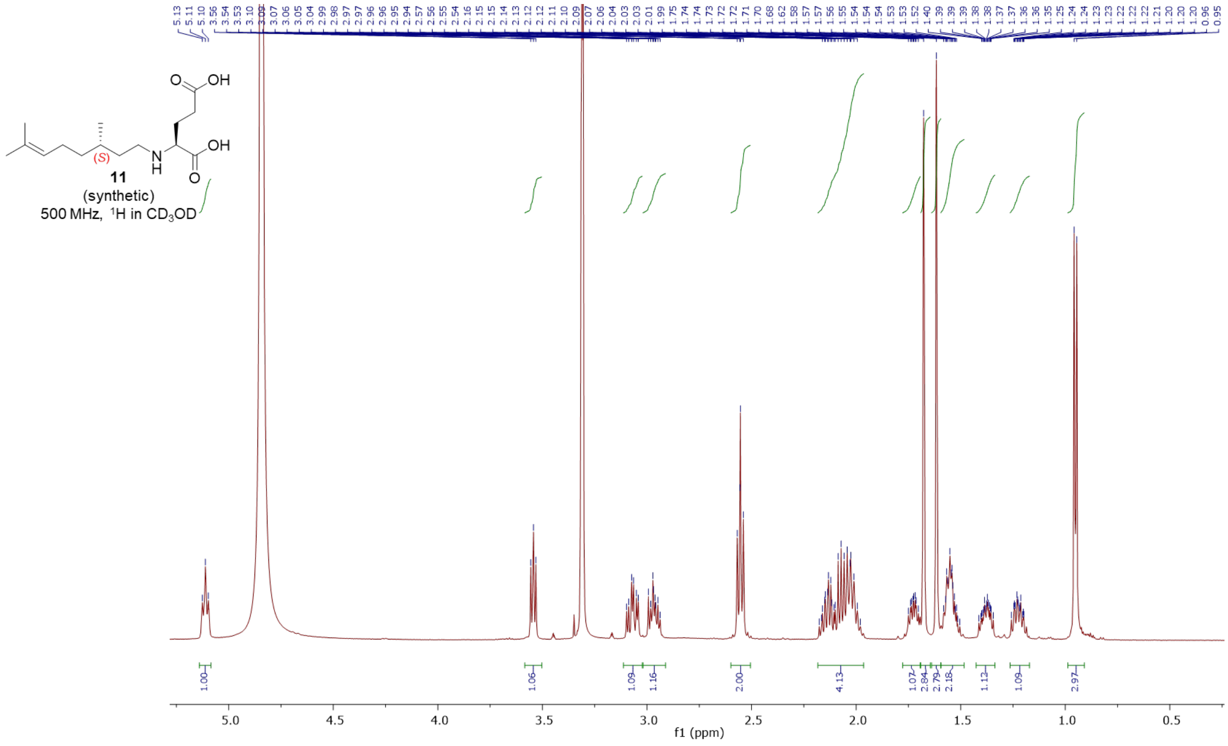

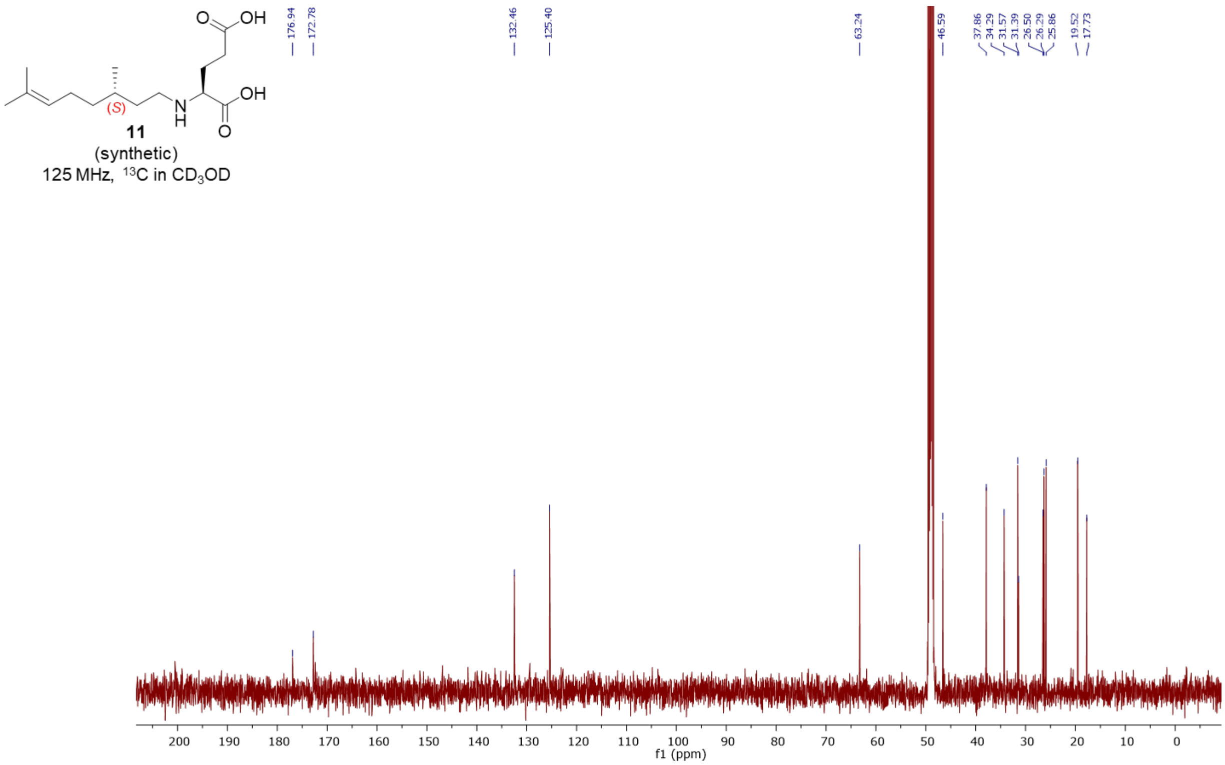
11
(synthetic)
125 MHz, 13C in CD3OD
f1 (ppm)
176.94
172.78
132.46
125.40
63.24
46.59
37.86
34.29
31.57
31.39
26.50
26.29
25.86
19.52
17.73

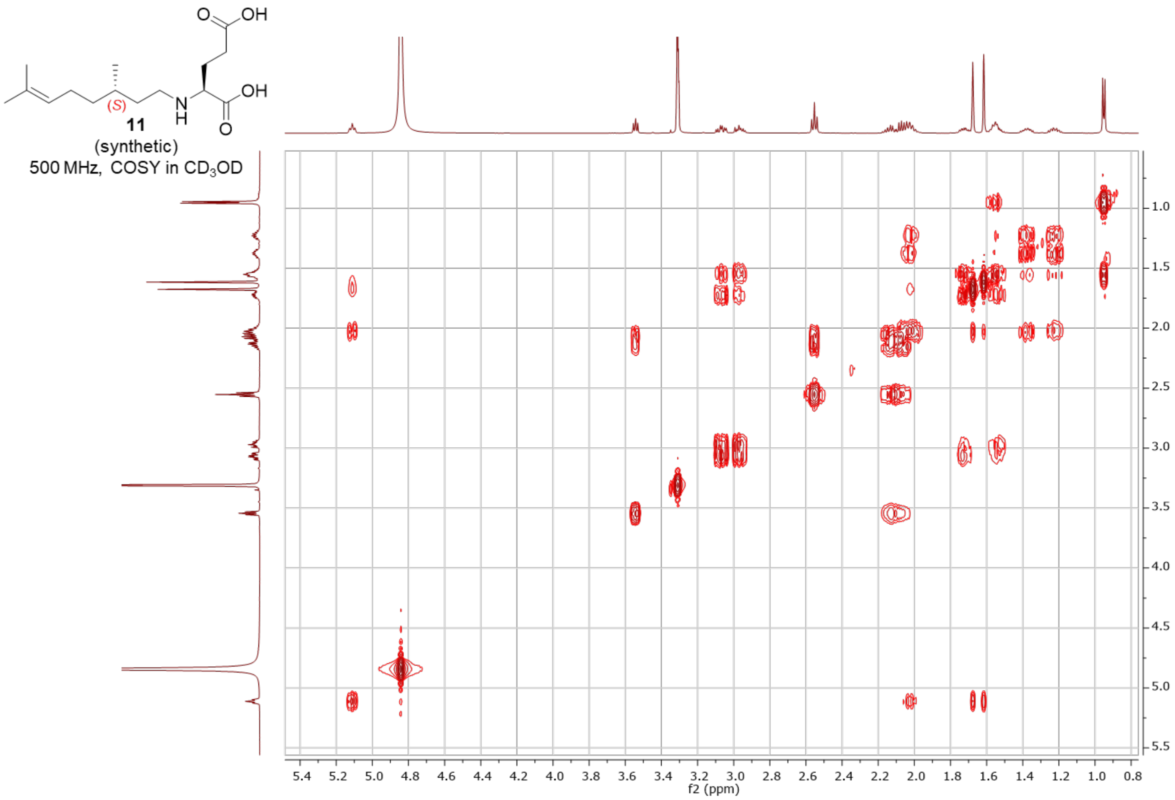
11
(synthetic)
500 MHz, COSY in CD3OD
f2 (ppm)
f1 (ppm)

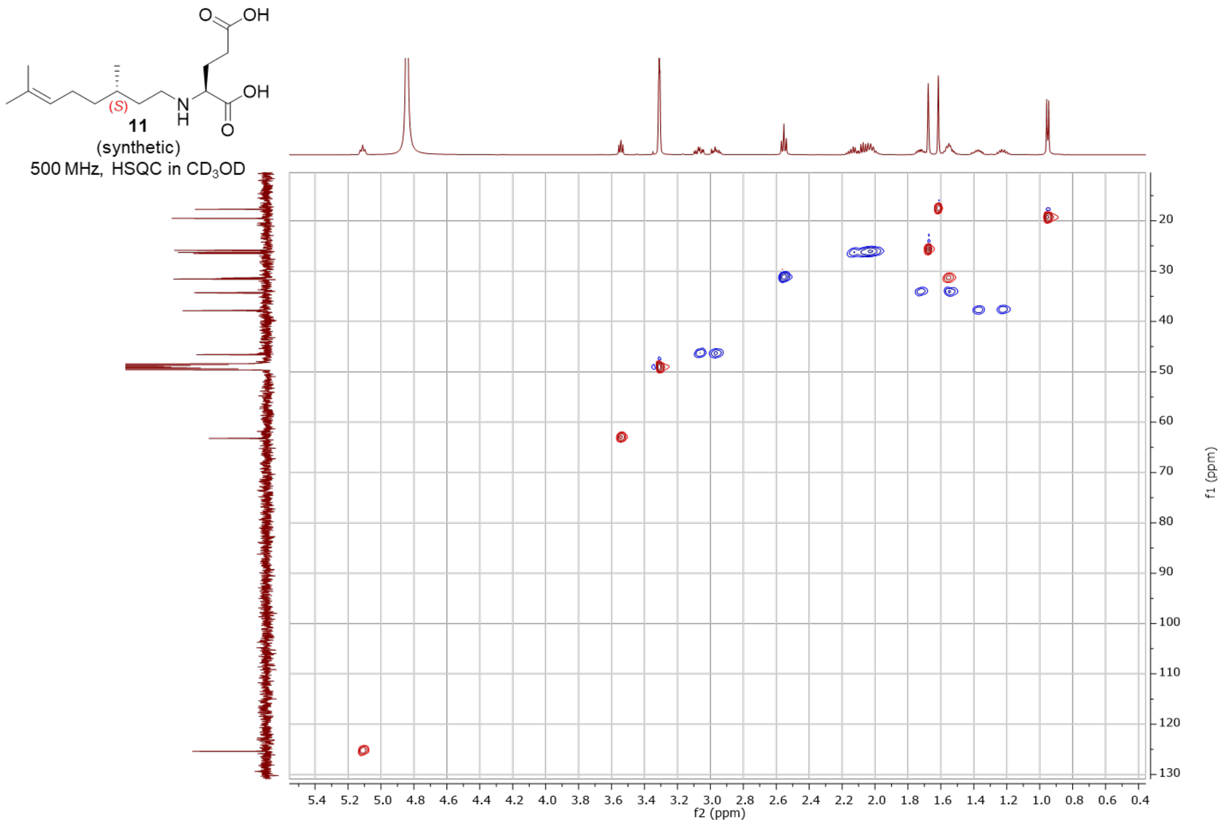
OH
O
OH
O
N
H
(S)
11
(synthetic)
500 MHz, HSQC in $CD_3OD$
f2 (ppm)
f1 (ppm)

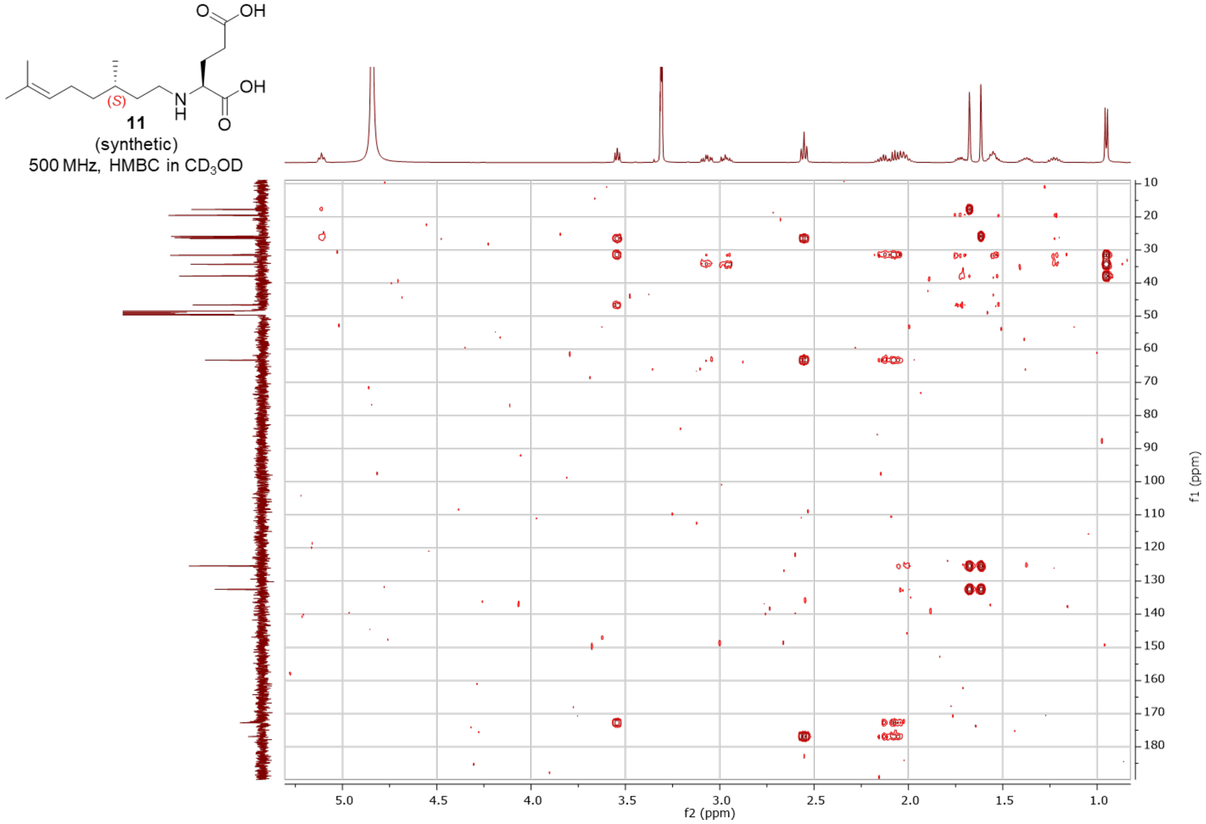
OH
O
OH
O
N
H
(S)
11
(synthetic)
500 MHz, HMBC in $CD_3OD$
f2 (ppm)
f1 (ppm)

*N*-(*R*)-citronellyl-L-glutamic acid - synthetic (**12**)

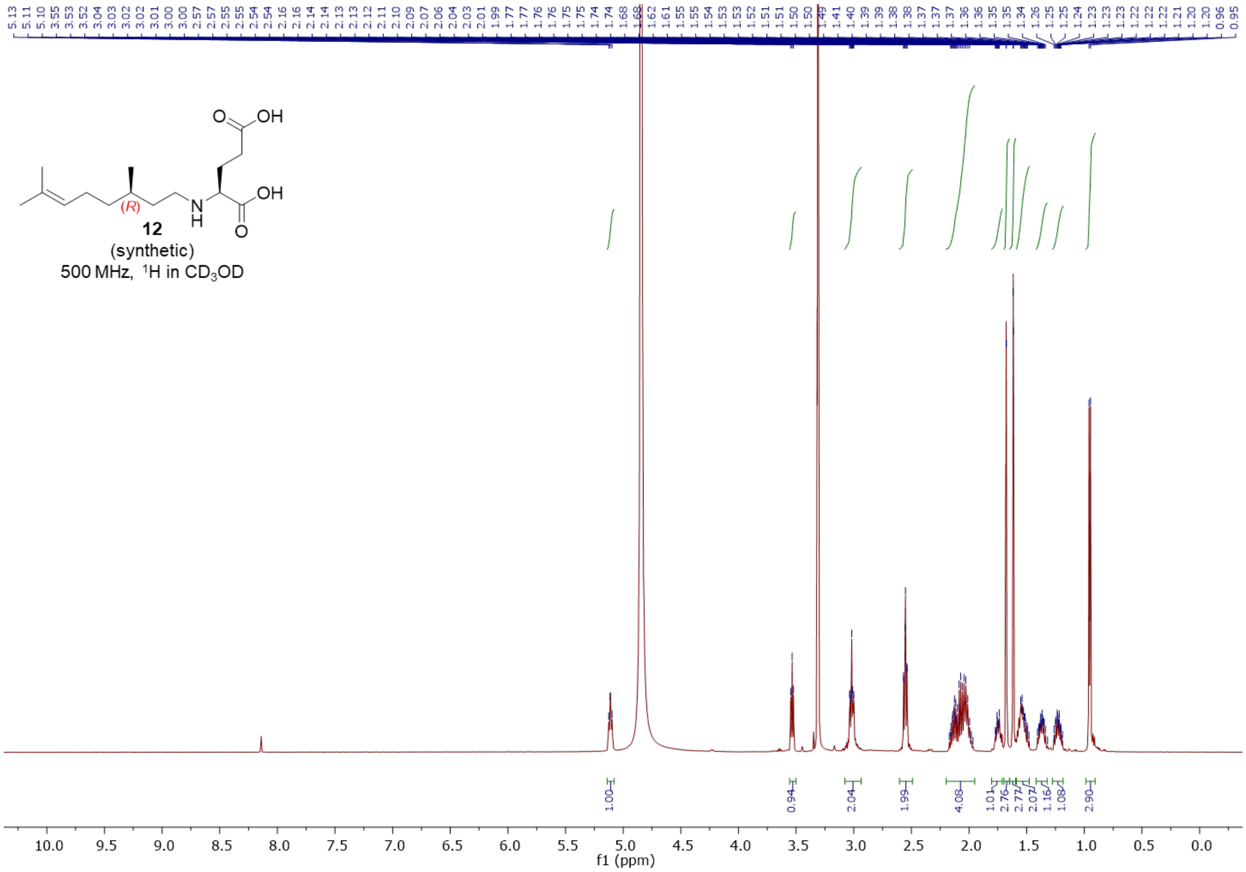


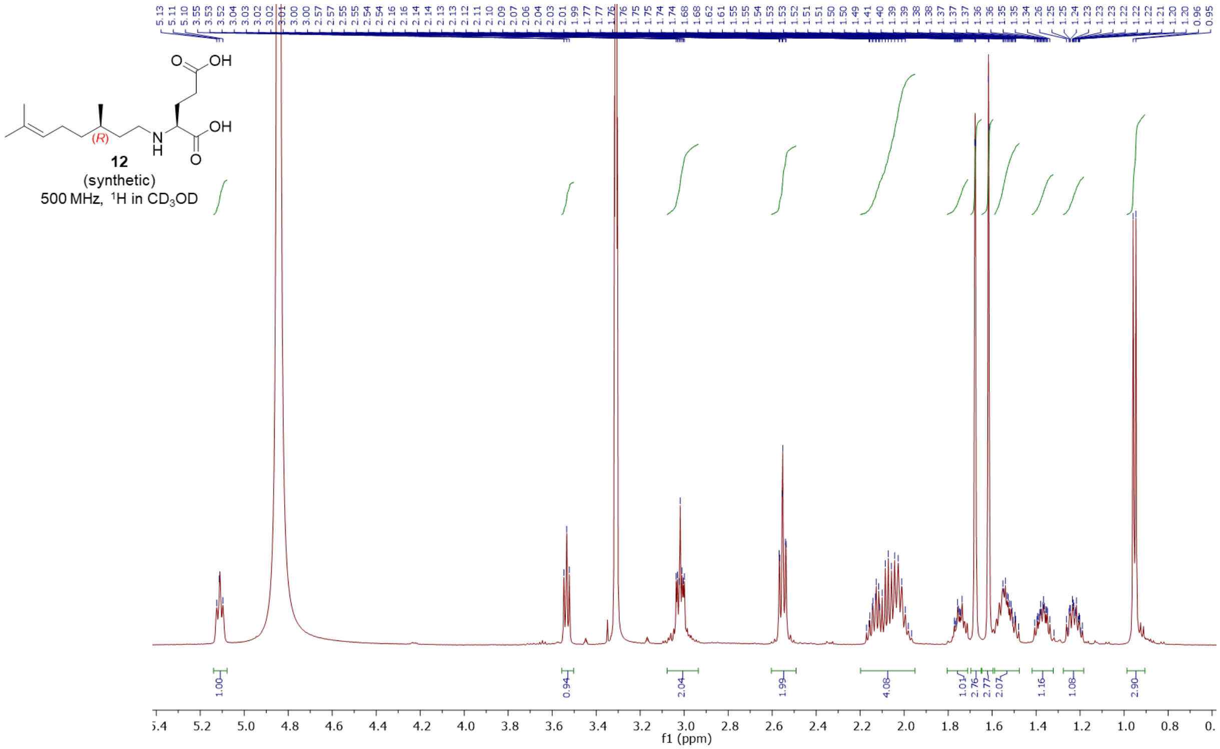

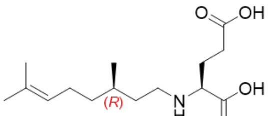


12
(synthetic)
125MHz, $^{13}C$ in $CD_3OD$

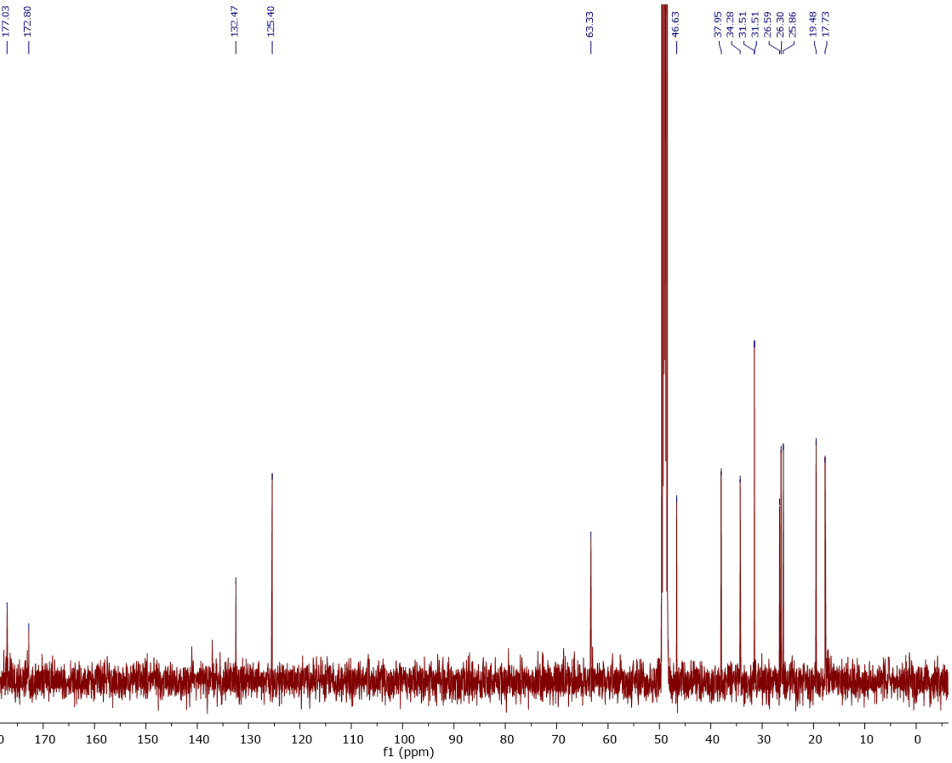


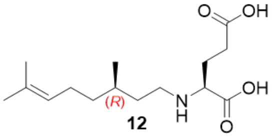


12
(synthetic)
500 MHz, COSY in $CD_3OD$

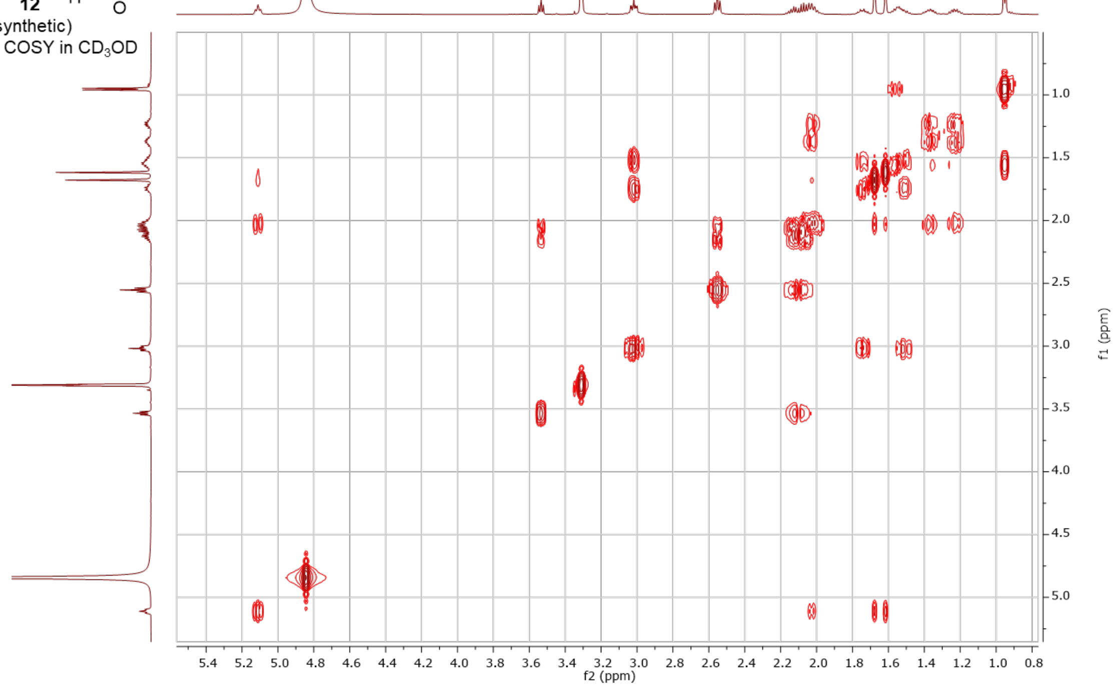

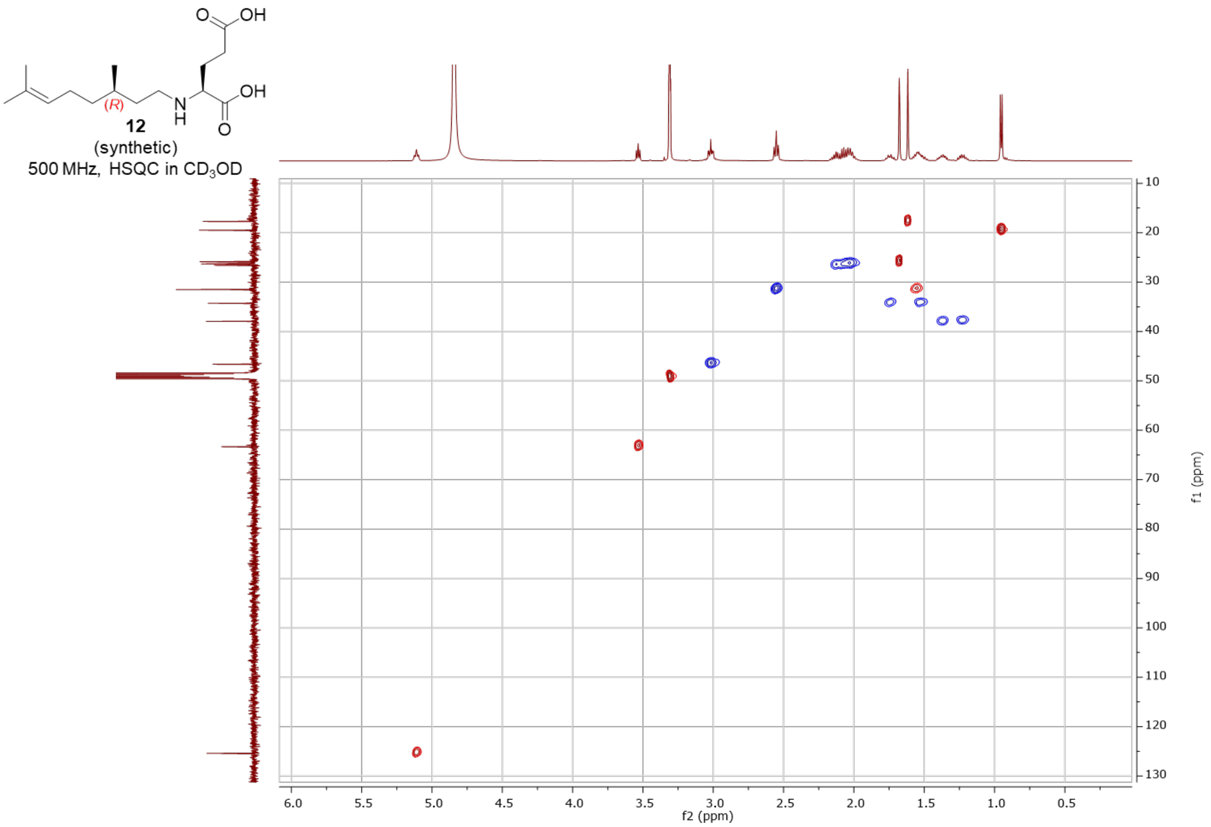

12
(synthetic)
500 MHz, HSQC in $CD_3OD$
f2 (ppm)
f1 (ppm)


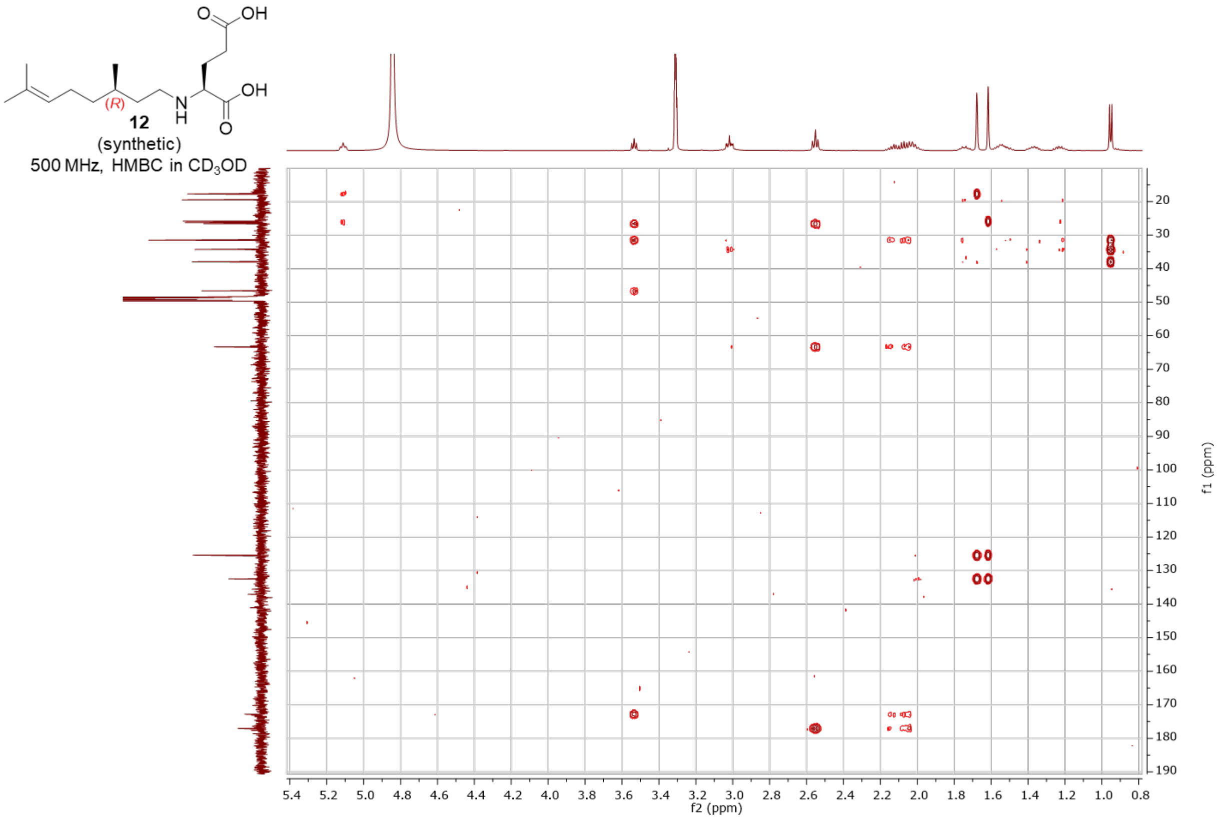

12
(synthetic)
500 MHz, HMBC in $CD_3OD$
f2 (ppm)
f1 (ppm)

<u>*N*-isopentyl-L-glutamic acid – synthetic (**13**)</u>

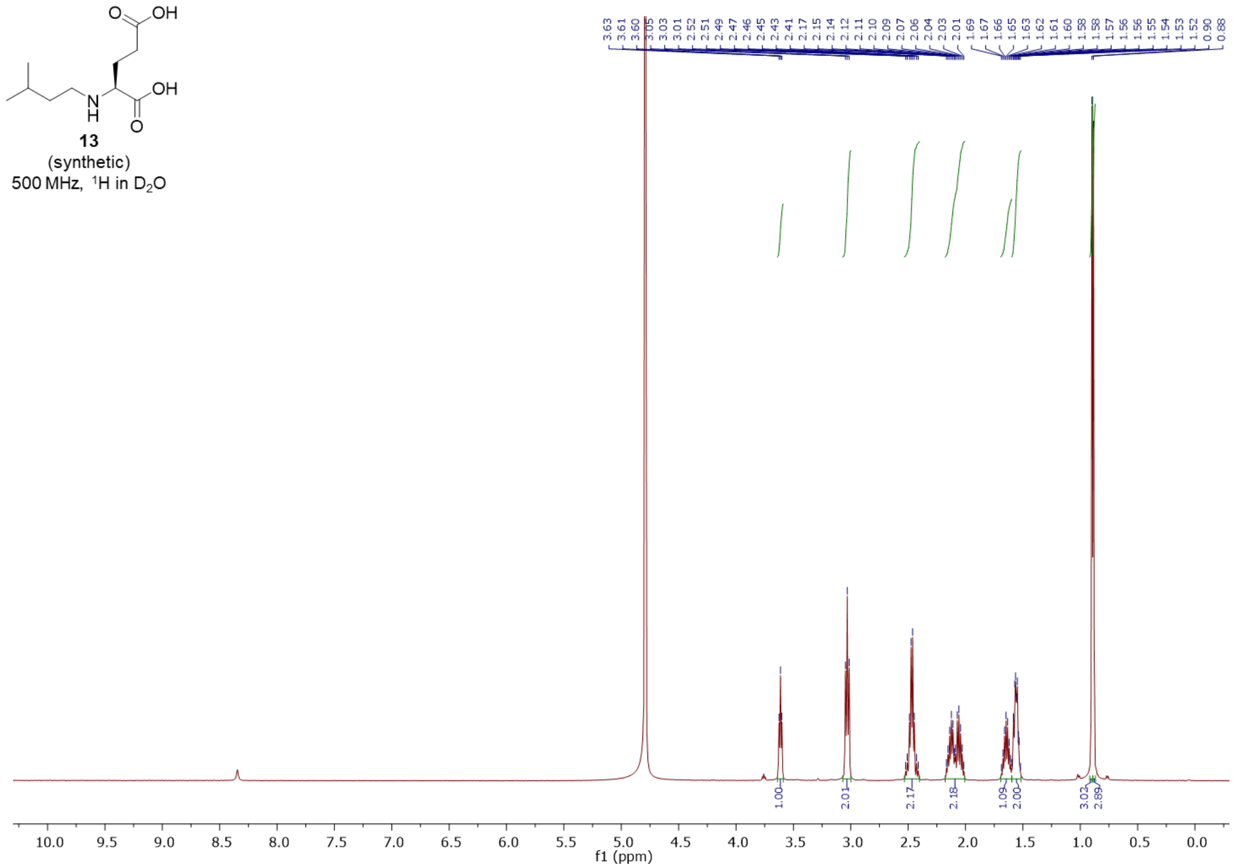


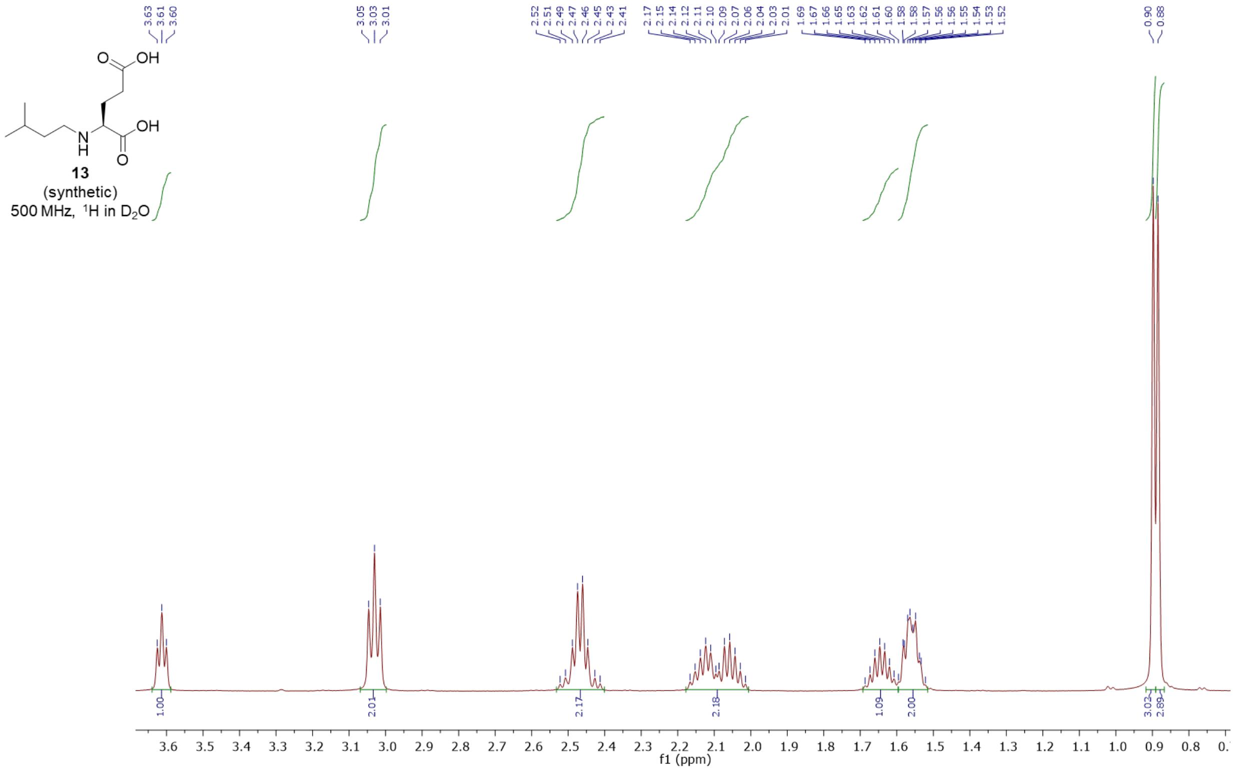

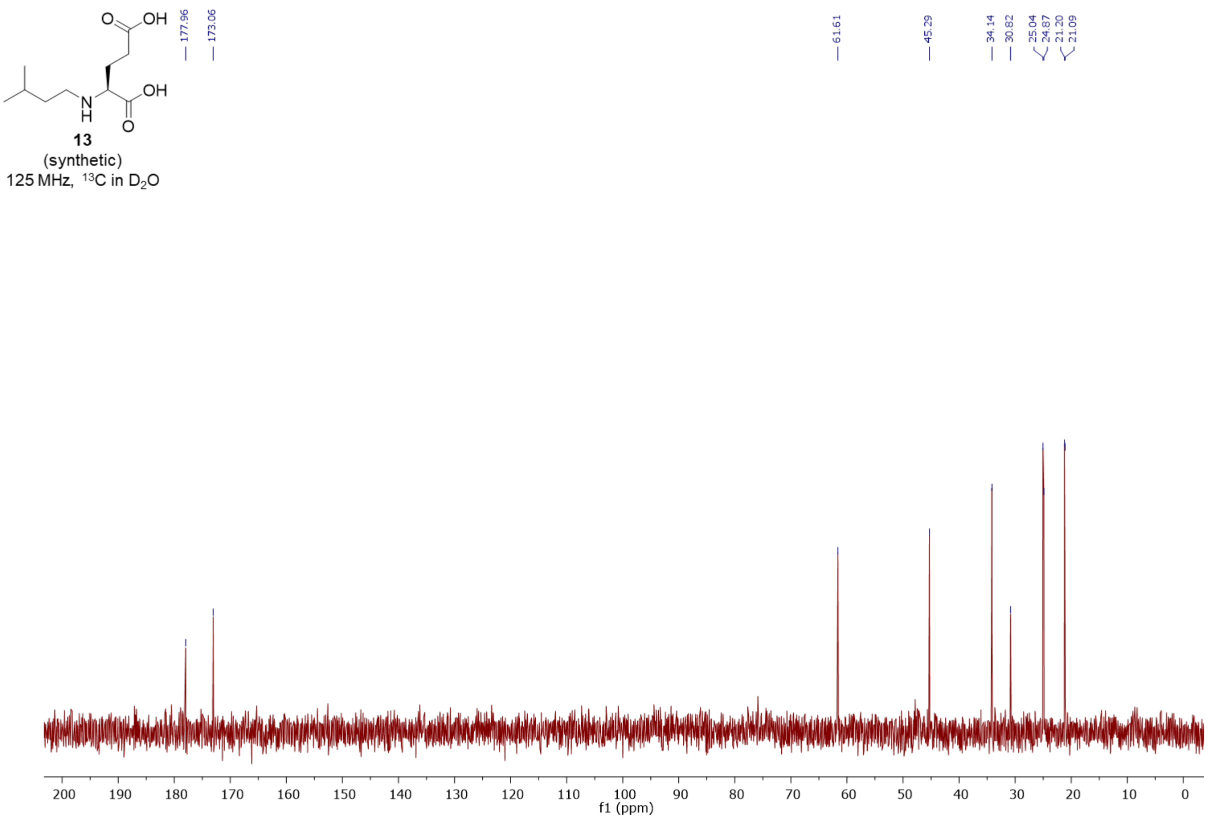
13
(synthetic)
125 MHz, ¹³C in D₂O
177.96
173.06
61.61
45.29
34.14
30.82
25.04
24.87
21.20
21.09
f1 (ppm)

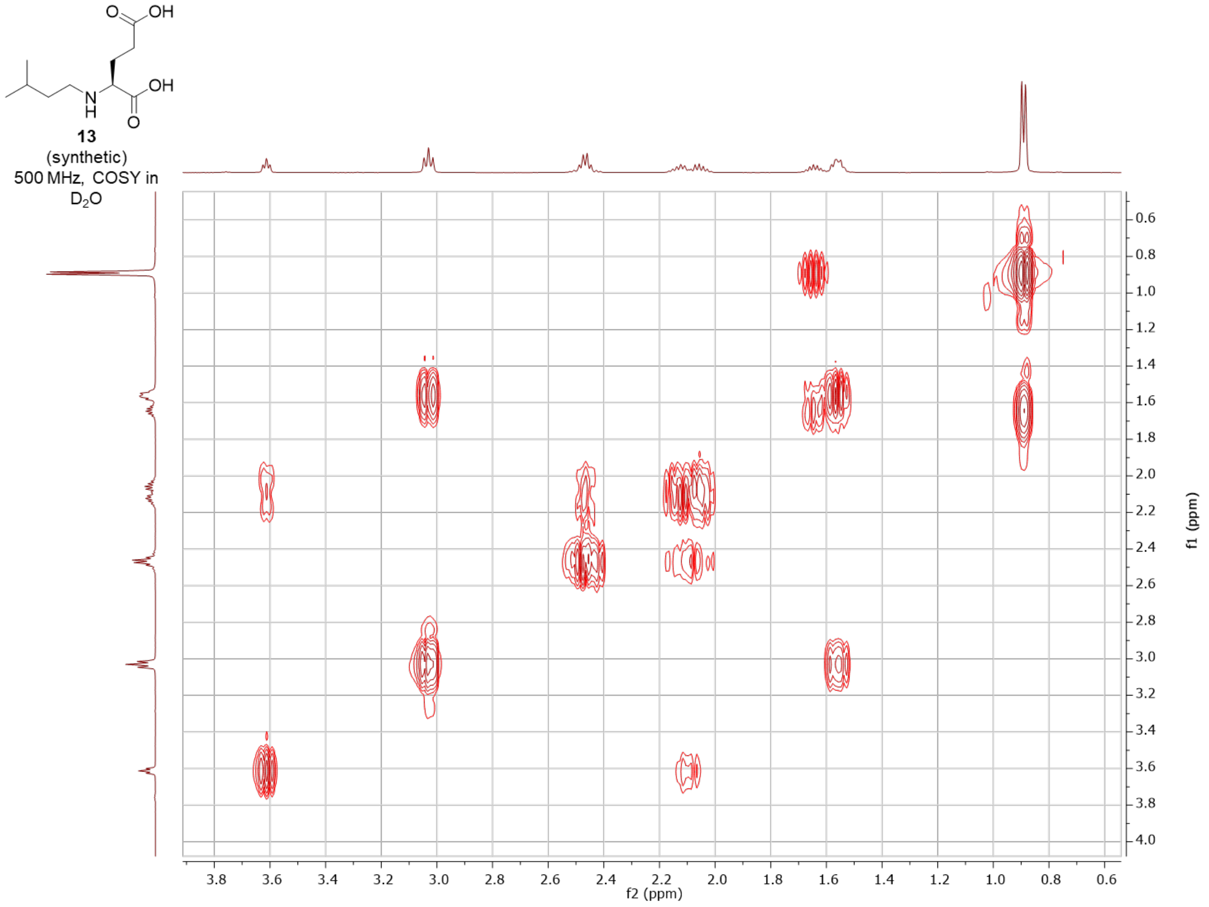
13
(synthetic)
500 MHz, COSY in D₂O
f2 (ppm)
f1 (ppm)

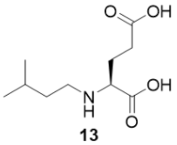

**13**
(synthetic)
500 MHz, HSQC in $D_2O$

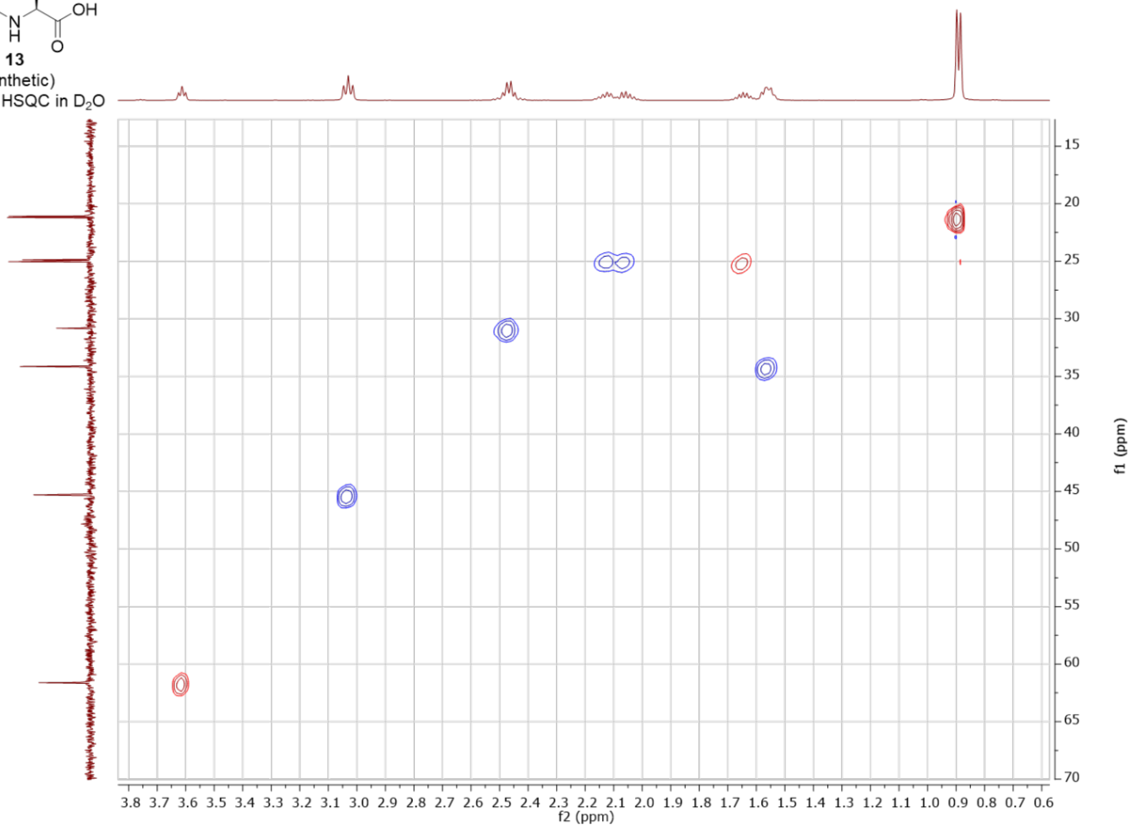


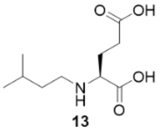

**13**
(synthetic)
500 MHz, HMBC in $D_2O$

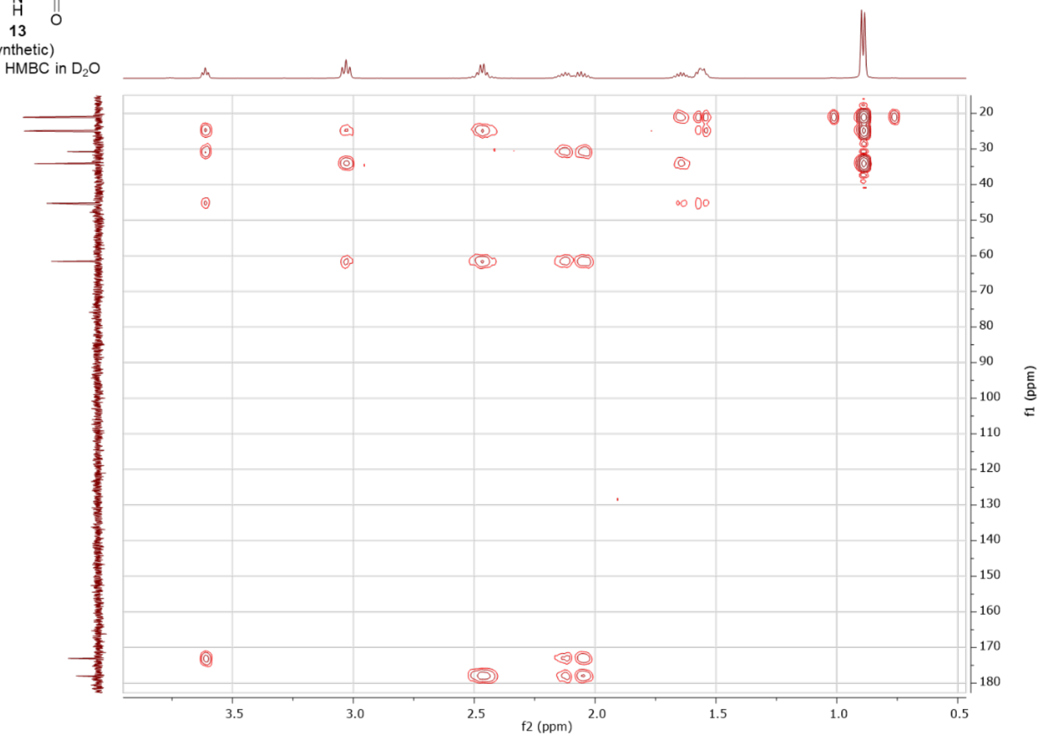

#### *N*-isopentyl-(3*R*)-hydroxy-L-glutamic acid – enzymatic (**14**)

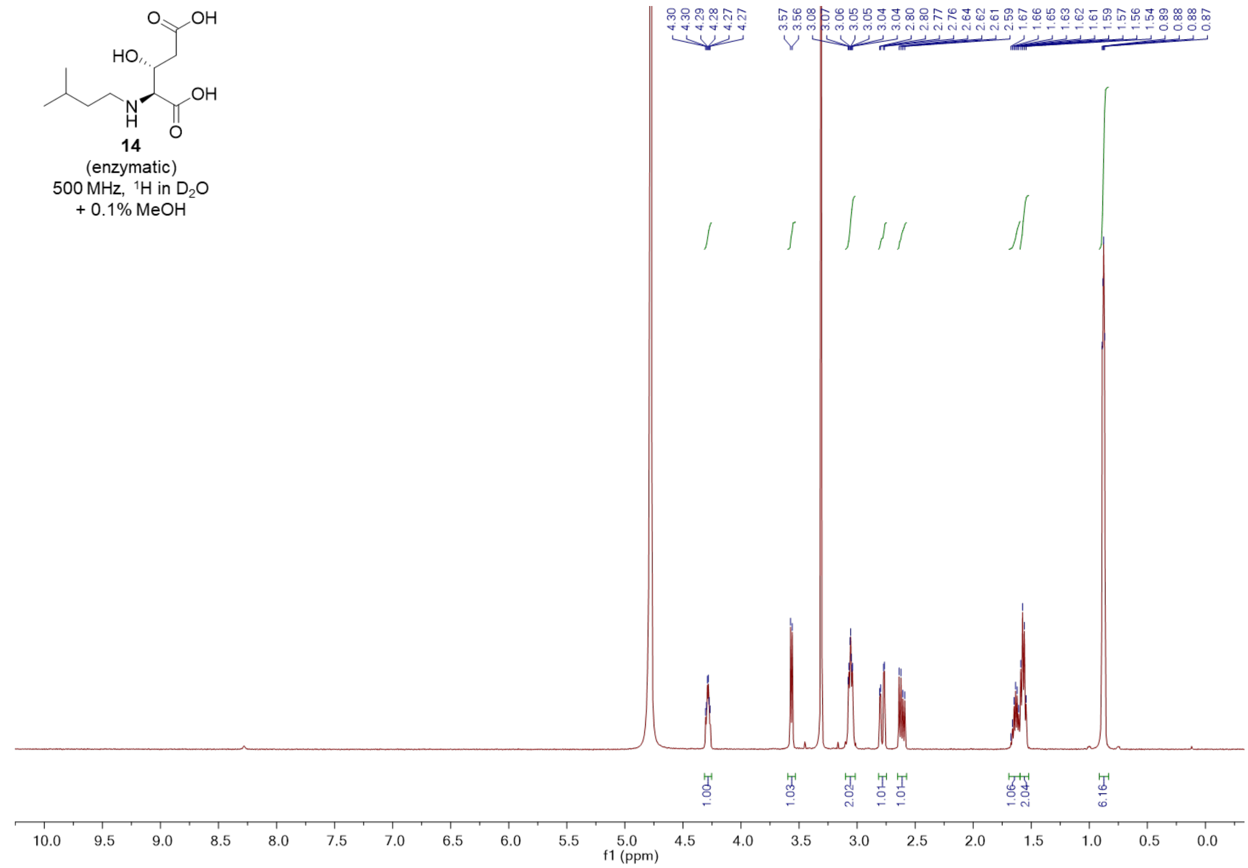


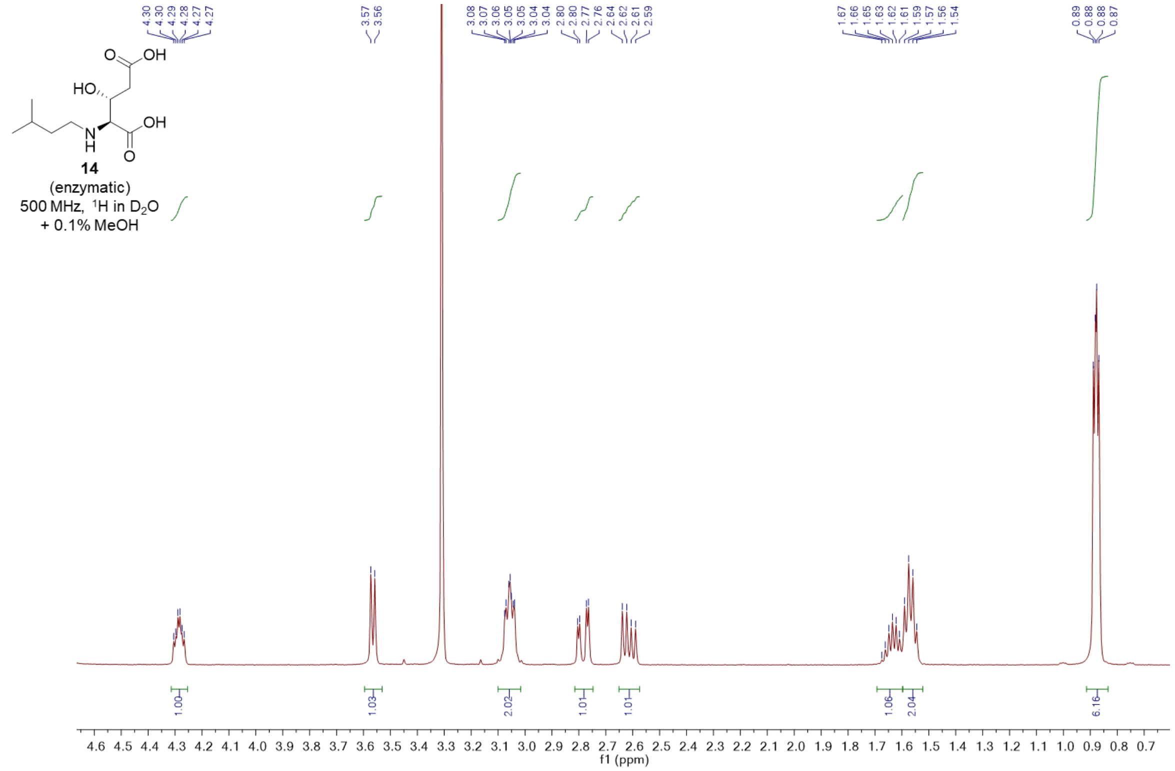

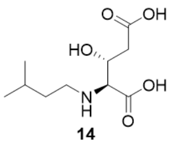

**14**
(enzymatic)
125 MHz, $^{13}C$ in $D_2O$
+ 0.1% MeOH

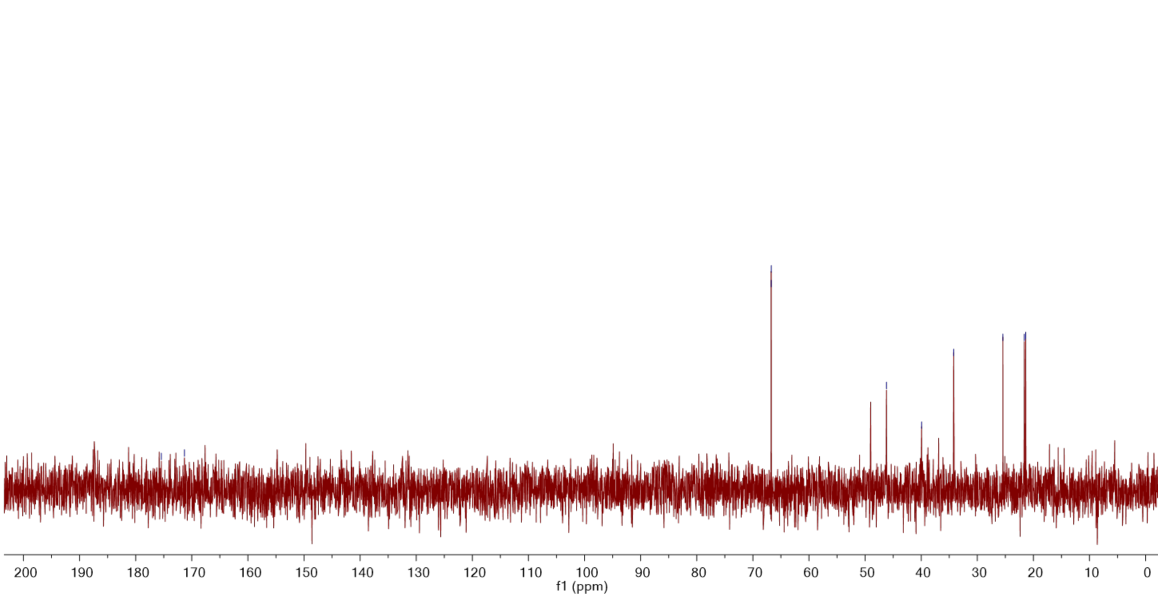


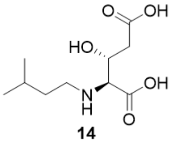

**14**
(enzymatic)
500 MHz, COSY in $D_2O$
+ 0.1% MeOH

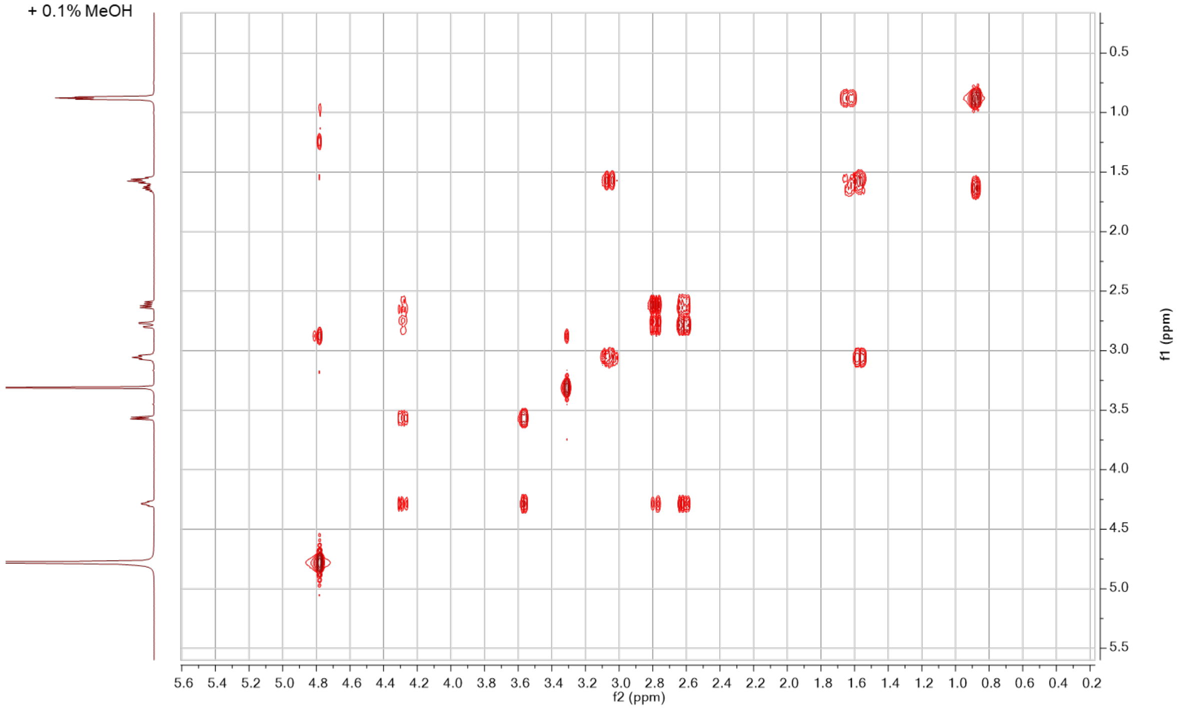

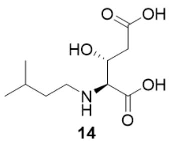


**14**
(enzymatic)
500 MHz, HSQC in $D_2O$
+ 0.1% MeOH

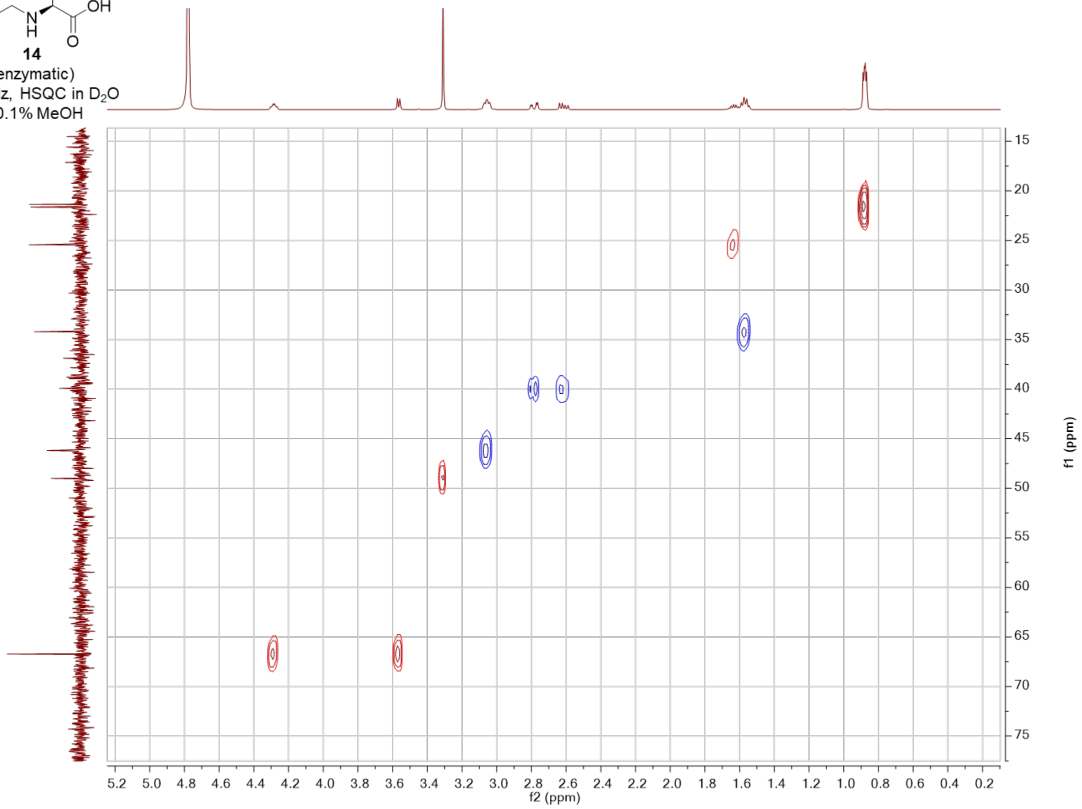


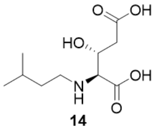


**14**
(enzymatic)
500 MHz, HMBC in $D_2O$
+ 0.1% MeOH

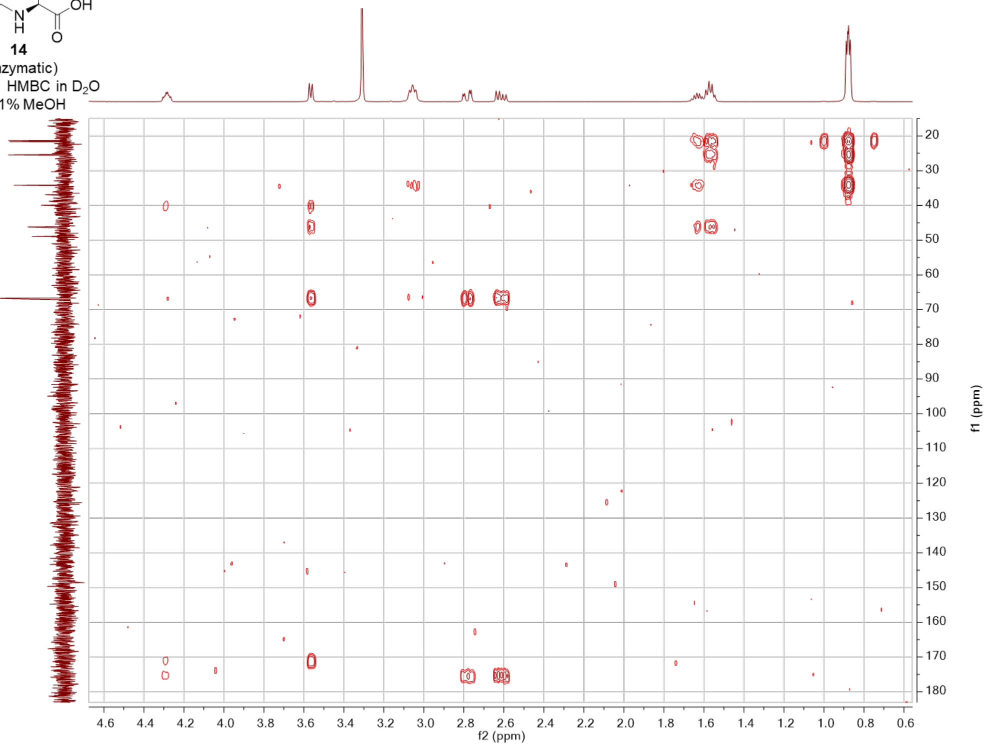

3*R*-hydroxy-L-glutamic acid - synthetic (**SI-9**)

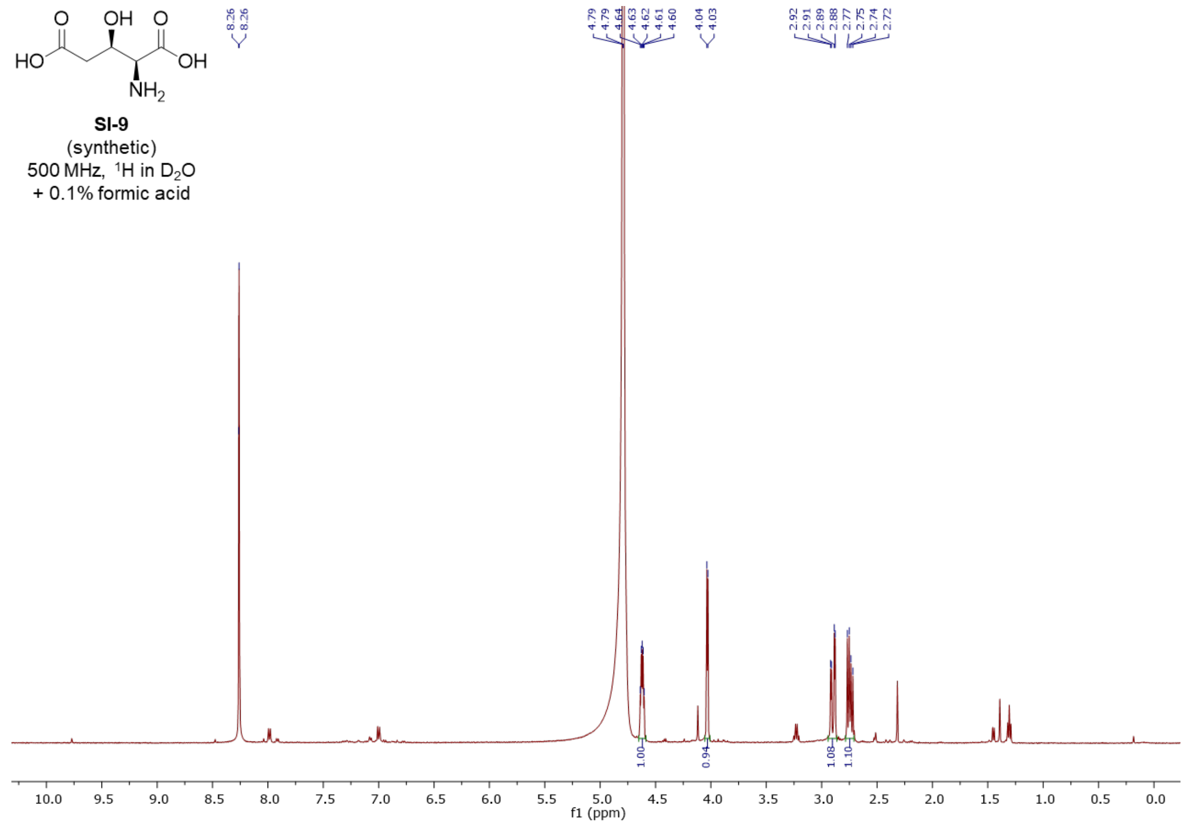


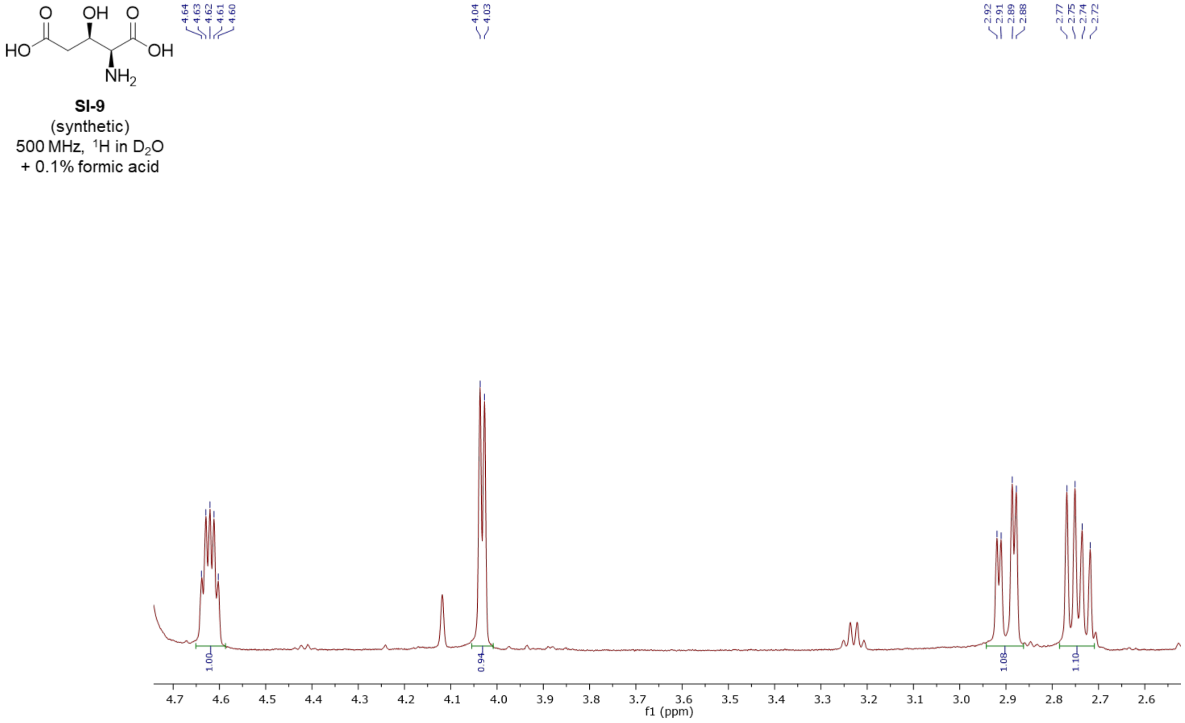

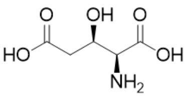


**SI-9**
(synthetic)
125 MHz, $^{13}C$ in $D_2O$
+ 0.1% formic acid

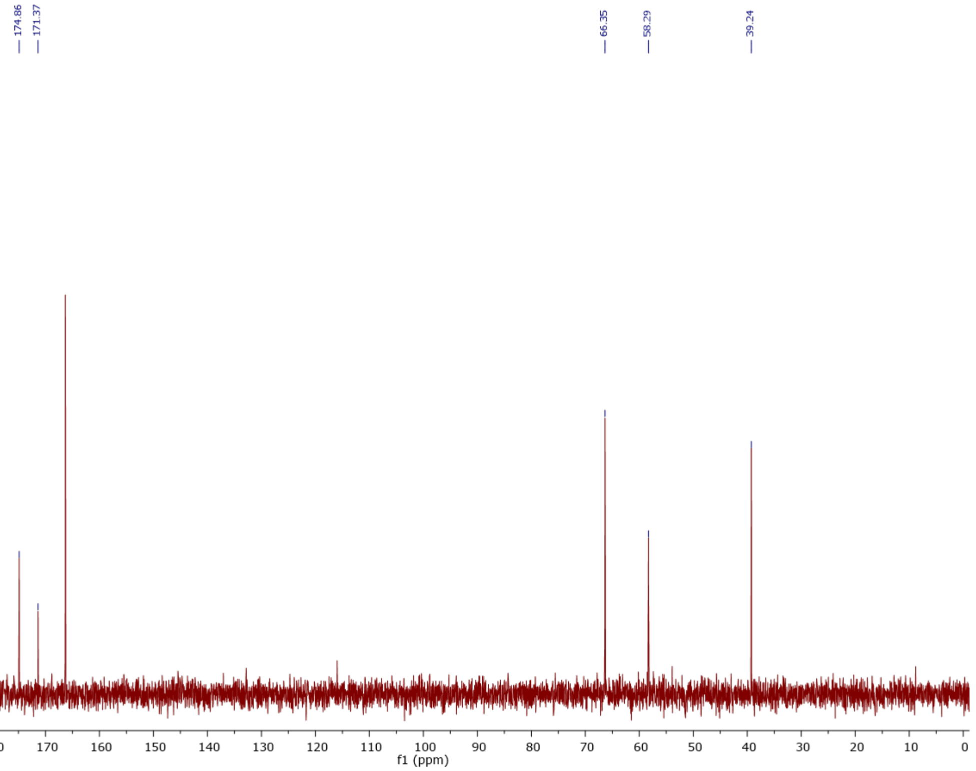


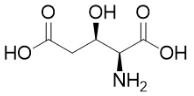


**SI-9**
(synthetic)
500 MHz, COSY in $D_2O$
+ 0.1% formic acid

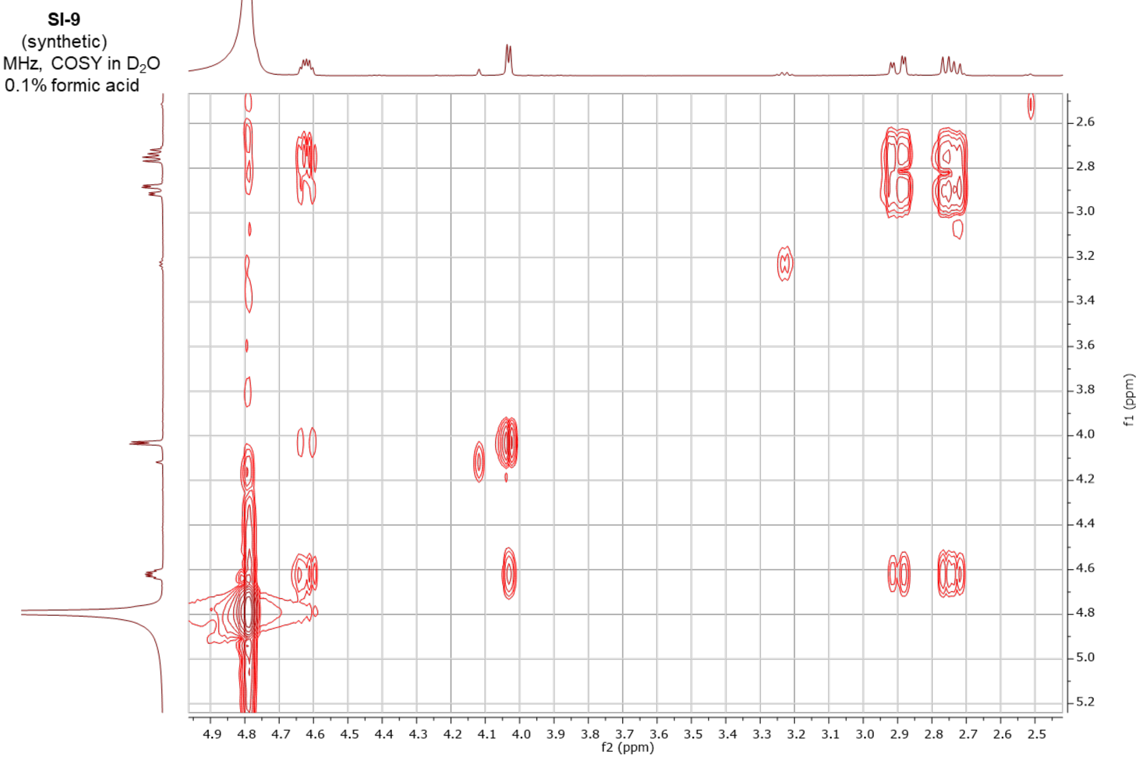

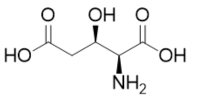


**SI-9**
(synthetic)
500 MHz, HSQC in $D_2O$
+ 0.1% formic acid

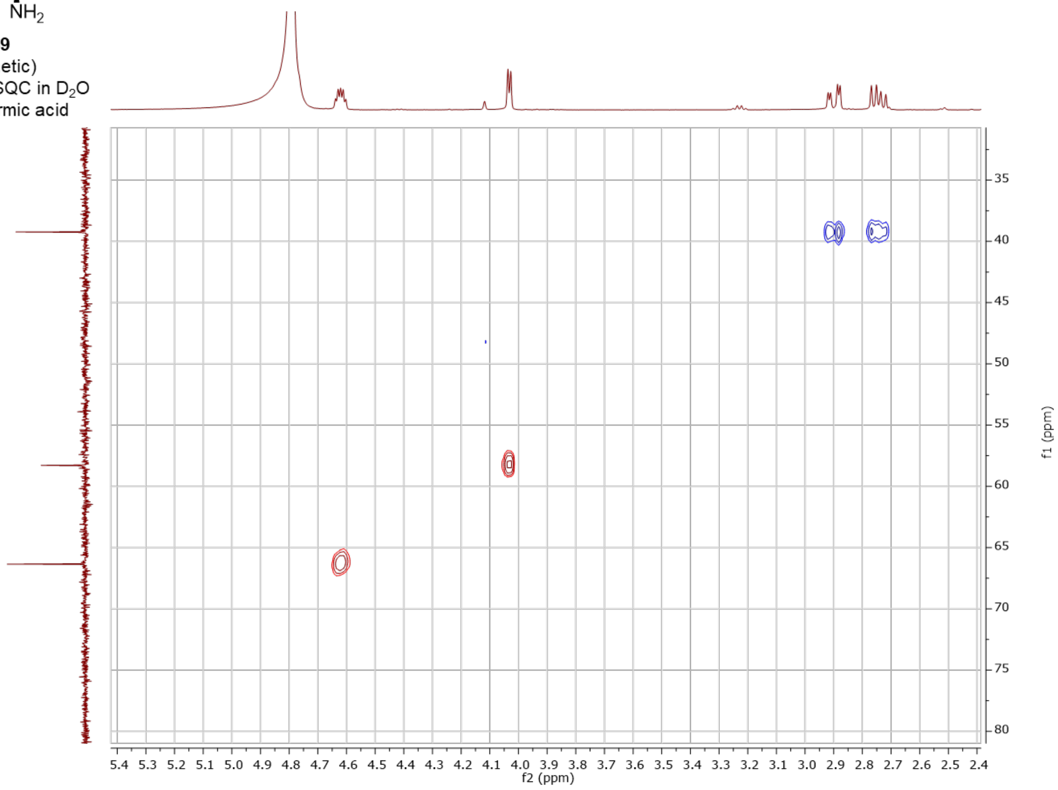


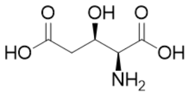


**SI-9**
(synthetic)
500 MHz, HMBC in $D_2O$
+ 0.1% formic acid

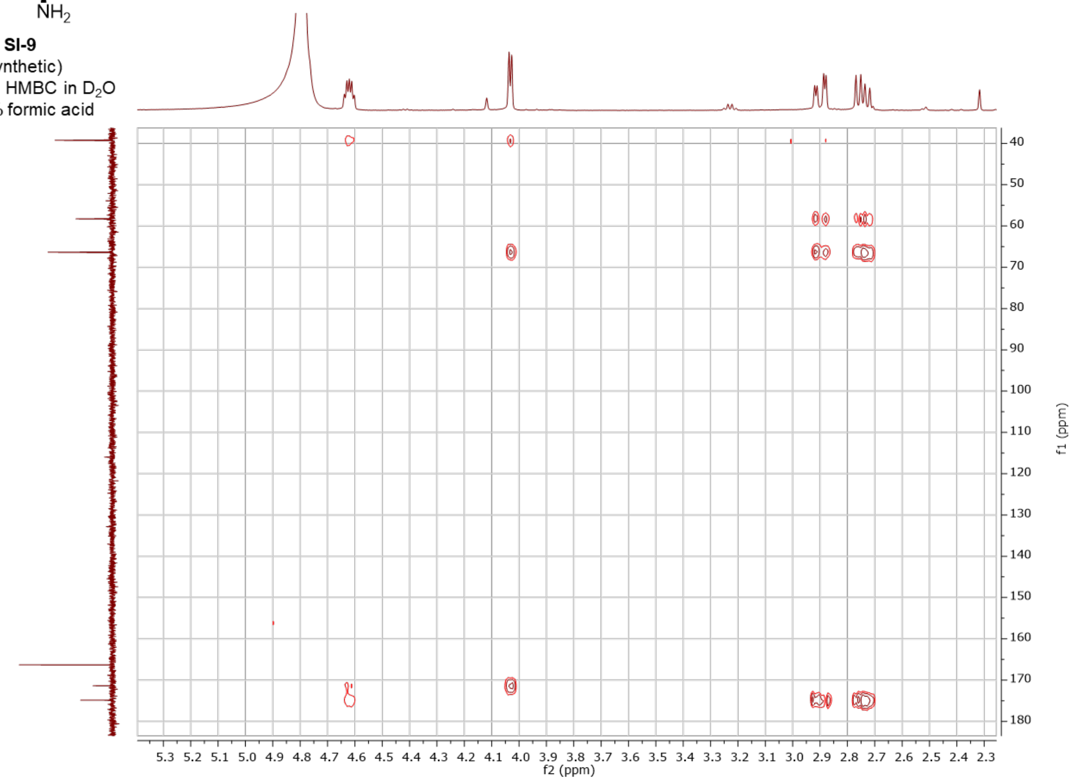
